# Termite diet rather than geographical origin determines the microbiome composition and functional genetic structure of nests from South American and African representatives, as revealed by a multiomics approach

**DOI:** 10.1101/2022.08.13.503768

**Authors:** Juan José González Plaza, Jaromír Hradecký, Jan Šobotník

## Abstract

Termites represent one of the most important insect groups worldwide due to their key role as plant decomposers and proxy of carbon recycling in the tropical rainforest ecosystems. Besides, high relevance in research has been given to these social insects due to a prominent role as urban pests. However, one of the most fascinating aspects of termites are their defence strategies that prevent the growth of detrimental microbiological strains on their nests. One success factor is the key role of the nest allied microbiome. Understanding how beneficial microbial strains aid termites in pathogen biocontrol strategies could provide us with an enhanced repertoire for fighting antimicrobial resistant strains or mine for genes for bioremediation purposes.

We carried out a multiomics approach for dissecting the nest microbiome in a wide range of termite species, covering several feeding habits and three geographical locations at two tropical sides of the Atlantic Ocean, and an African savanna. Our experimental approach included untargeted volatile metabolomics, targeted evaluation of volatile naphthalene, taxonomical profile for bacteria and fungi through amplicon sequencing, and further dive into the genetic repertoire through a metagenomic sequencing approach.

Volatile naphthalene was present in species belonging to the genera *Nasutitermes* and *Cubitermes*. We further assessed the apparent differences in terms of bacterial community structure, having found a stronger influence from feeding habits and genera, rather than the geographical location. Lastly, our metagenomic analysis revealed that the gene content provides both soil feeding genera with similar functional profiles, while the wood feeding genus shows a different one. These results seem to be independent of the geographical location, indicating that the nest functional profile is heavily influenced by the diet of the termite inhabiting, building, and maintaining the nest.

## 1 Introduction

Termites represent one of the most successful stories of adaptation in the evolution of animal kingdom. One of the main ingredients for biological success is the effective use and exploitation of a specific niche and its resources, what allows for the transference of genetic information to the next generations. Termites have adapted to use plant dead material in order to obtain their energetic resources. In the process of doing so, they have achieved a key role in the recycling of carbon in one of the most important ecosystems on Earth, the tropical rainforest. The use of lignocellulosic sources depends largely on the symbiotic relationships with a wide range of symbiotic microorganisms (Brune, 1998; Ohkuma and Brune, 2010). Not only interact termites with these microorganisms for energetic purposes, but are indeed deeply related with them because they live on their surface or at their nest material. Nests are the Achilles heel of termites, since their whole social structure turns around their central place of existence. That is so because termites live in populated colonies with a series of intricated galleries, were the conditions of humidity (Wiltz et al., 1998) and temperature create an ideal setting for the growth of pathogenic organisms, such as fungi. Not exclusively because of these two, but also due to the high content of nutrients in termite nests in comparison with the nutrient-poor surrounding soil environment (López-Hernández, 2001; van der Sande et al., 2018), yet rich in nutrient cycling (Jordan and Herrera, 2015). Among the reasons for the high content of nutrients in nests and immediate surrounding areas is the way in which nests are built. Nests are a product of a considerable amount of faeces as cementing tool (Jose et al., 1989; Chouvenc et al., 2018), and soil particles that are transported from the surrounding environment (Mizumoto et al., 2021). Considering these limited availabilities of food substrates in the surrounding soil, the conditions are set for the competitive race for survival in the microbial moiety, with a continuous input of external strains, potentially detrimental and threatening the existence of the whole colony. These external microorganisms thrive for an environment which could limit the presence of grazing organisms (Clarholm, 1981; Rønn et al., 2002), and high availability of nutrients provided by the input of termite faeces when they maintain and build their nests (Chouvenc et al., 2013). However, termites are very successful in their strategies for limiting the growth of detrimental microorganisms. Among them, it has been appointed that fumigation with certain type of compounds, such as naphthalene, aids in controlling pathogens (Chen et al., 1998a; Wright et al., 2000).

In this context, we have carried out a research effort to better characterize the microbial communities living in termite nests among several termite species across different geographical locations. We were interested in termites with two feeding strategies, wood and soil, originating at both sides of the Atlantic, including rainforest and savanna representatives. To evaluate the presence of volatile compounds, we used a coupled untargeted gas chromatography–mass spectrometry (GC-MS) analyte identification strategy, followed by a targeted naphthalene analysis. These metabolomics approaches were complemented with a taxonomic characterization of the microbial communities existing on the same material, in order to try to understand the potential co-occurrences of certain identified metabolites with bacterial or fungal genera. Lastly, we further performed shotgun metagenomic sequencing from selected termite colonies for a deeper understanding of the genetic microbial contribution to defence and naphthalene production.

Our results showed first a much lower amount of naphthalene in termite nests than those previously reported in literature. A very interesting observation found throughout the three omics used approaches (metabolomic, amplicon sequencing, metagenomic sequencing) is that genera and especially the feeding habits determine the relationships between the different microbial communities, regardless of the geographical origin.

## 2 Material and methods

### 2.1 Sample collection and initial processing

Nests from twenty-eight termite colonies were collected during 2019 and 2020 at different sampling campaigns in South America and Africa. The first sampling site is located in French Guiana, Petit-Saut Dam area, which is located in Sinnamary river (5°03’ N, −53°02.76’ E), and is characterized by a hot and humid weather throughout the year with two dry (February-March, July-November) and two rainy seasons (Colas et al., 2020).

In Africa we collected samples in the area of Ebogo village (Cameroon, 3°23.86’N, 11°28.19’E), located in the Mbalmayo Forest Reserve. This area is characterized by a bimodal precipitation pattern (Knoben et al., 2019), with a dense evergreen forest subjected to perturbations on approximately a third of the territory due to agricultural practices (Mey and Gore, 2021). The dry periods span from December to February, and July to August (Zapfack and Engwald, 2008). The second African sampling site was Nyika National Park (Malawi, - 10°32.92’N, 33°53.47’E), characterized mostly by miombo woodland, and in minor degree by montane dambos and grasslands, where mean average temperatures are of 23 °C (Allingham and Harvey, 2013), and the dry season lasts mostly from June to October, while rainfalls occur chiefly during November-March (van Velden et al., 2020). Two of the samples originate in the breeds at Faculty of Forestry and Wood Sciences (Czech University of Life Sciences Prague, Czech Republic), maintained in humid constant temperatures of 28 °C.

Nest samples were processed in clean hexane-wiped metal trays under laminar flow hood to avoid contamination. Large pieces of nests devoid of termites were broken in small fragments and stored at −80 °C until processing. Samples were further homogenized in a Retsch oscillatory mill. The hexane-wiped steel 10 mL chambers were filled with nest material, closed, submerged into liquid nitrogen until stabilized. Samples contained in steel chambers were then homogenized to fine powder for 5 minutes at 30 Hz frequency, recovered, and stored at −80 °C until further analysis.

### 2.2 Metabolite profiling

For initial solid phase microextraction (SPME) volatiles profiling, 200 mg of previously homogenized samples were placed into different 10 mL screw-top vials for headspace analysis, and were sealed with a magnetic cap. Sample incubation and the following metabolite extraction, were performed at 50 °C in agitator of Gerstel MPS2 autosampler (Gerstel, SUI) for 10 and 30 minutes respectively. For separation and detection gas chromatograph coupled with time of flight mass spectrometer (GC-TOF-MS) (Leco Pegasus 4D, Leco, USA) was used. Temperature programmed injector was operated in splitless mode at 275 °C. For separation, a 30 m (0.25 mm i.d., 0.25 μm film thickness) Rxi-5Sil MS (Restec, USA) column was used. The temperature program for the 1D oven was as follows: 40 °C for 1 min, then ramped at a rate of 10 °C/min to 210 °C, then at 20 °C/min to 300 °C with a hold time of 3 minutes. The total GC run time was 26 min. The mass spectrometer was operated in mass range 35-500 m/z with acquisition speed of 10 Hz.

For targeted naphthalene analysis, an SPME method based on Cao (2012) and Tsimeli et al. (2008) was used, employing standard addition quantification. One hundred mg of homogenized sample were placed into 10 mL HS vial and 0.5 mL of water (UHPLC-MS purity, Supelco, USA) was added before sealing with magnetic screwcap. Four vials per sample were prepared. Thru septum, to first two vials, 10 μL of methanol (HPLC Plus purity, Sigma-Aldrich, USA) were injected, while in resting two, 10 μL of methanol containing different amount of naphthalene were injected. Sample was heated under agitation at 50 °C for 10 minutes and then volatiles from headspace were collected onto a SPME fibre for 5 minutes.

Linear response of naphthalene was observed, when blank nest sample was spiked on levels from 1-1100 ng/g. Method accuracy and reproducibility were checked by measuring a certified reference material (CRM) of polyaromatic hydrocarbons in soil (CRM170, Lot: LRAC8900, Sigma-Aldrich, USA). Measuring six replications of CRM, relative standard deviation (RSD) of entire determination was 12%, resulting in uncertainty (U=2xRSD) of 25%. Comparing obtained naphthalene content of 563 ng/g ± 140 ng/g (average ±U) to certified value of 573 ± 45 ng/g showed good accuracy of the presented method. The only modification for CRM measurement was that due to high naphthalene concentration, sample weight was reduced to 10 mg per vial.

### 2.3 Species identification

DNA from termites belonging to each colony were isolated using the Qiagen Blood and Tissue kit according to manufacturer’s instructions. Isolated DNA was used for amplification of the Cytochrome Oxidase II gene (COII) with the following program: 1 minute at 94 °C, 30 cycles of 94 °C for 15 sec, 61 °C for 1 minute, and 72 °C for 35 sec, and a final step of elongation at 72 °C for 10 minutes. PCR products were submitted to Microsynth AG (Switzerland) for Sanger sequencing. Results were compared to NCBI nr database for retrieving BLAST results (Altschul et al., 1990). The used primers were previously described by Benjamino and Graf (Benjamino and Graf, 2016).

### 2.4 Microbial DNA isolation

Powder homogenized samples were used for isolation of DNA. One hundred mg of soil were used as starting material using the NucleoSpin Soil DNA kit (Macherey-Nagel, Germany). The whole procedure was carried out under laminar flow, using sterile material, and kit components were exclusively opened and manipulated under sterile conditions. Samples were quantified by Qubit 2.0 Fluorometer (Thermo Scientific, USA) and quality checked through spectrophotometric methods. Samples were submitted to Novogene (Hong-Kong, China) for sequencing of the regions 16S V34 and ITS2 in order to retrieve the taxonomic structure of the bacterial and fungal communities of the samples. DNA was subjected to quality control checks through DNA purity (optical density 260/280), agarose gel electrophoresis to evaluate carryover of contaminants, and Qubit (Thermo Scientific, USA) concentration evaluation.

### 2.5 Sequencing

#### 2.5.1 16S rRNA gene sequencing

Amplicon sequencing for the 16S V4 region with primers 515F (GTGCCAGCMGCCGCGGTAA) and 806R (GGACTACHVGGGTWTCTAAT) was carried out following standard company procedures (Novogene, Hong Kong, China). In brief, a 292 bp amplicon was obtained through PCR with the above-mentioned specific primers. PCR products were quantified, purified (Qiagen Gel Extraction Kit, Qiagen, Germany), and used for library preparation (NEBNext Ultra DNA Library Pre-Kit, Illumina, San Diego, CA, USA). Libraries were sequenced following a 250 bp sequencing strategy.

#### 2.5.2 Internal Transcribed Spacer (ITS) Sequencing

ITS amplicon sequencing was carried out by Novogene (Hong Kong, China), following standard company procedures. In brief, 1 ng of DNA was used to amplify fungal ITS genes from ITS2 region with a specific set of primers ITS3 (GCATCGATGAAGAACGCAGC), and ITS4 (TCCTCCGCTTATTGATATGC), with an amplicon size of 386 bp. The overall procedure with amplicons follows the one explained for 16S.

#### 2.5.3 Metagenomic sequencing

We selected two genera, comprising four samples each, from those where naphthalene was produced (*Cubitermes* and *Nasutitermes*). For comparison purposes a third group with no naphthalene presence in nests was selected (*Anoplotermes*). DNA was isolated as above-mentioned, and submitted for Illumina PE150 sequencing at Novogene (Hong Kong, China). After quality control in standard agarose gels and quantity concentration estimation with Qubit 2.0 (ThermoFisher, USA), DNA was randomly sheared through sonication. Fragments were end-polished, A-tailed, and then the Illumina adapters (5’ Adapter: 5’-AGATCGGAAGAGCGTCGTGTAGGGAAAGAGTGT-3’; 3’ Adapter: 5’-GATCGGAAGAGCACACGTCTGAACTCCAGTCAC-3’) were ligated. A PCR amplification was carried out using P5 and P7 indexed oligonucleotides, and products purified with AMPure XP system. Size distribution of libraries was evaluated at Agilent 2100 Bioanalyzer (Agilent Technologies, CA, USA), and quantified through real-time PCR for achieving equimolar concentrations. Sequencing was subsequently carried out at Illumina instruments.

### 2.7 Microbiome data processing and analysis

#### 2.6.1 Amplicon sequencing data: 16S and ITS

The assignment of paired-end reads to samples was based on their unique barcodes, afterwards truncated by cutting off the barcode and primer sequence. Overlaps of paired-end reads served for merging reads using FLASH (Magoč and Salzberg, 2011). Reads were quality filtered to obtain high-quality clean tags (Bokulich et al., 2013) using QIIME (V1.7.0). Reads were then compared to a reference database (Gold DB, http://drive5.com/uchime/uchime_download.html) with the UCHIME algorithm (Edgar et al., 2011) for detection and removal of chimera sequences. Clean reads were stored in individual files in fastq format. These files were used as input in a common tool used in microbiome studies for bioinformatic treatment of clean demultiplexed reads, QIIME2 (release 2021.4) (Bolyen et al., 2019). This wrap-up tool was used to import (q2-import, SingleEndFastqManifestPhred33V2) the clean demultiplexed single fastq files corresponding to our samples with the import plugin. Denoising was carried out with DADA2 (via q2-dada2-denoise single) (Callahan et al., 2016), what allows for identification of all observed amplicon sequence variants. ASVs were aligned using mafft (Katoh et al., 2002) (via q2-alignment), and used to build a phylogeny (fasttree2), via q2-phylogeny (Price et al., 2010).

After DADA2, sequences that were found in the negative control “.fastq” file were removed from the project through quality-control exclude-seqs plugins.

Core metrics were calculated with core-metrics-phylogenetic plugin, where rarefaction was performed using a sampling depth of 10,000 for 16S data, and 20,000 for ITS reads. Among metrics, we calculated Faiths’s Phylogenetic Diversity (Faith, 1992), beta diversity such as UniFrac (Lozupone et al., 2007), unwheighted UniFrac (Lozupone and Knight, 2005), Jaccard Distance and Bray-curtis dissimilarity. Plugins were core-metrics-phylogenetic, alpha-group-significance, and beta-group-significance, where the metadata column of choice was “geography”, “genus”, or “feeding-group”.

Taxonomy was assigned with the q2-feature-classifier classify-sklearn naïve Bayes taxonomy classifier (Bokulich et al., 2018; Kaehler et al., 2019), for the 16S we used the weighted Greengenes 13_8 99% OTUs full-length sequences (DeSantis et al., 2006), while for ITS we used silva-138-99-nb-classifier (Pruesse et al., 2012; Quast et al., 2013; Yilmaz et al., 2014). Taxonomy plots were drawn in R through the ampvis2 package, using the tutorials available at: https://sites.google.com/a/ciad.mx/bioinformatica/home/metagenomica/visualizacion/ampvis2, and https://madsalbertsen.github.io/ampvis2/articles/ampvis2.html, where we used the Heillinger transformation in order to represent the PCA (Legendre and Gallagher, 2001).

Once taxonomy was assigned and frequency calculated per sample, we performed a Least Discriminant Analysis (LDA) Effect Size (LEfSe) (Segata et al., 2011). This algorithm discovers biomarkers that identify features on the tested biological conditions. A text tab separated file in metadata-columnar format was loaded at a Galaxy instance (https://huttenhower.sph.harvard.edu/galaxy/). The OTU table was filtered in the case of 16S data to include only those that satisfied frequency > 0.001 in at least one of the samples.

### 2.8 Metagenomics bioinformatics

Sequenced reads were first cleaned with trimmomatic (Bolger et al., 2014), and then subjected to a quality control process (FASTX Toolkit 0.0.14. FASTQ/A short-reads pre-processing tools), as shown in **Supplementary material S1**. We followed then the Kalamazoo Protocol for metagenome assembly khmer 0.8.4 (Brown et al., 2013). We used the digital normalization protocol as stated (Brown et al., 2012, 2015), and used the normalized reads as an input for SqueezeMeta v1.4.0, v. May 2021 (Tamames and Puente-Sánchez, 2019), in order to carry out a first assembly via Megahit (Li et al., 2015). We carried out that first assembly using the *seqmerge* mode, but did not use the statistics or any other results. After carrying out the assembly, the generated contigs were used as external assembly reference (extassembly) in SqueezeMeta, and thus, no assembly was performed at this second step. At this second round, the input consisted on trimmed, quality control passed, no spaces (in house script) files containing separately forward and reverse pairs, gzipped. The pipeline followed the usual procedure by SqueezeMeta, with the following tools:

Assembly carried out with Megahit (Li et al., 2015), contig statistics with prinseq (Schmieder and Edwards, 2011), and redundant contig removal by cd-hit (Schmieder and Edwards, 2011). Merging of contigs was carried out with Minimus2 (Treangen et al., 2011), RNA prediction with Barrnap (Seemann, 2014), taxonomy classification of 16S rRNA sequences by RDP classifier (Wang et al., 2007), tRNA/tmRNA prediction by Aragorn (Laslett and Canback, 2004), and Prodigal for Open Rreading Frames (ORF) prediction (Hyatt et al., 2010). Diamond (Buchfink et al., 2014) served as the tool to search for similar patterns at GenBank (Clark et al., 2016), eggNOG (Huerta-Cepas et al., 2016), KEGG (Kanehisa and Goto, 2000). Then HMM homology searches were performed by HMMER3 (Eddy, 2011) for the Pfam database (Finn et al., 2016). Reads were mapped against contigs with Bowtie2 (Langmead and Salzberg, 2012), and binning was done using MaxBin2 (Wu et al., 2016) and Metabat2 (Kang et al., 2019), where binning results were combined using DAS Tool (Sieber et al., 2018). MiniPath (Ye and Doak, 2009) was used against KEGG (Kanehisa and Goto, 2000) and MetaCyc (Caspi et al., 2018) databases for pathway prediction.

At the end of the pipeline, tables for results processing in R were created by means of the smq2tables.py script provided in the SqueezeMeta package. Results were loaded in R through the use of SQMtools (Puente-Sánchez et al., 2020), and charts produced according to the SQMtools manual. Some of the R packages used were: “SQMtools”, “ggplot2”, “reshape2”, “pathview”, “data.table”.

Non-metric multidimensional scaling (NMDS) was carried out in RStudio (RStudio Team, 2020), using Transcripts per Million (TPM) calculated values obtained from SqueezeMeta pipeline, for those genes with KEGG annotation, through “vegan” package. This ordination collapses the information from all genes versus the samples, to visualize the relationships between samples.

We further used diamond (Buchfink et al., 2014) in order to search for genes and patterns of interest in our final contig list with Open Reading Frames, where the databases of reference were: AnHyDeg (Callaghan and Wawrik), AromaDeg (Duarte et al., 2014), BacMet (Pal et al., 2014), bactibase (Hammami et al., 2010), CARD (protein_homolog_model) (Alcock et al., 2020), and CAZy (Carbohydrate Active Enzymes database, http://www.cazy.org/) (Lombard et al., 2014). We used by default an e-value threshold of 0.005, an identity percentage threshold of 40, and manually curated the results discarding those hits with lower than 50 as bit-score. We also compared our assembled contigs with a plasmid database plsdb (Galata et al., 2019) through blastn (Altschul et al., 1990) with e-value threshold of 5 × 1^-10^ and an identity percentage of 50.

## 3 Results

### 3.1 Identification of termite species belonging to each collected colony

Alive termites belonging to each of the collected nests used in this study were taxonomically identified through molecular methods (COII amplification and sequencing). Results are shown in **Table 1**.

**Table 1.**
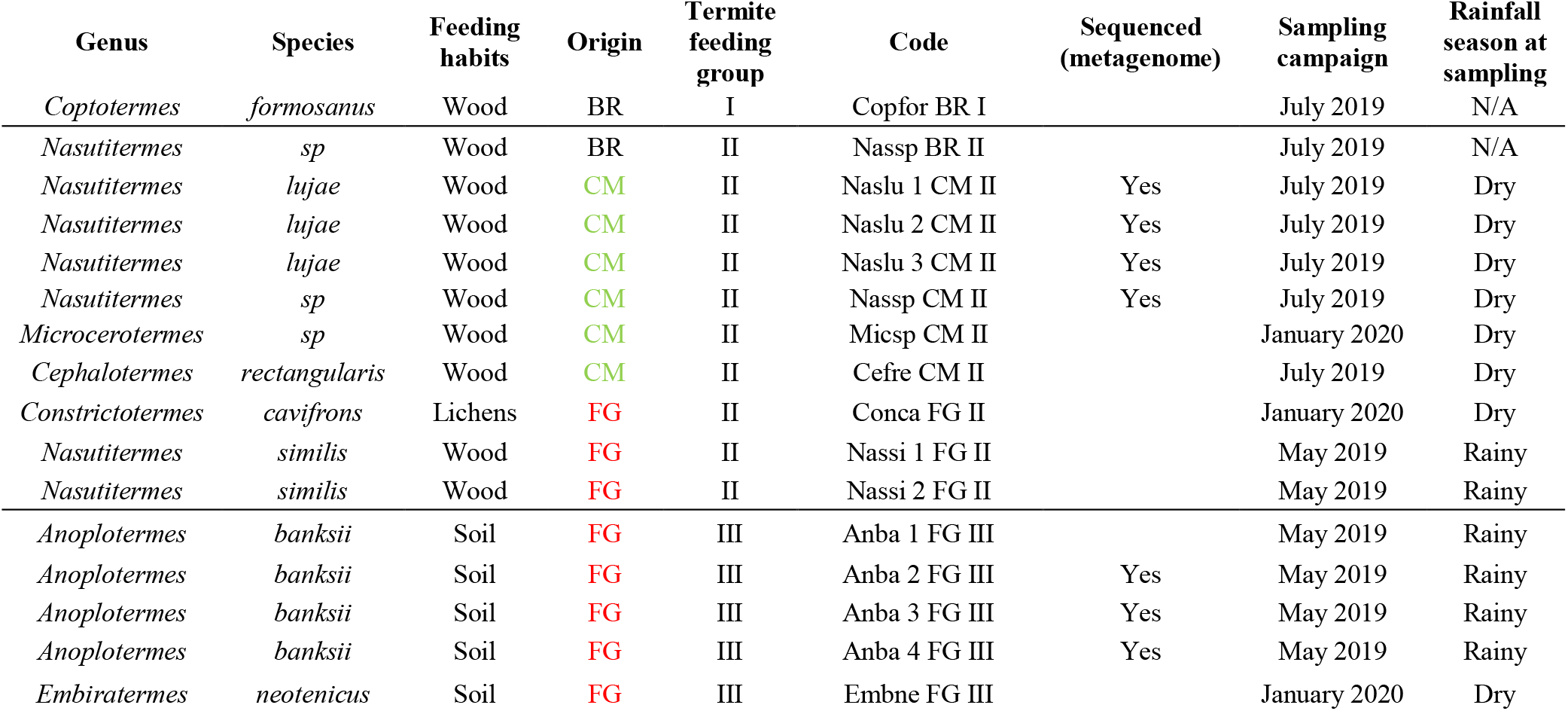
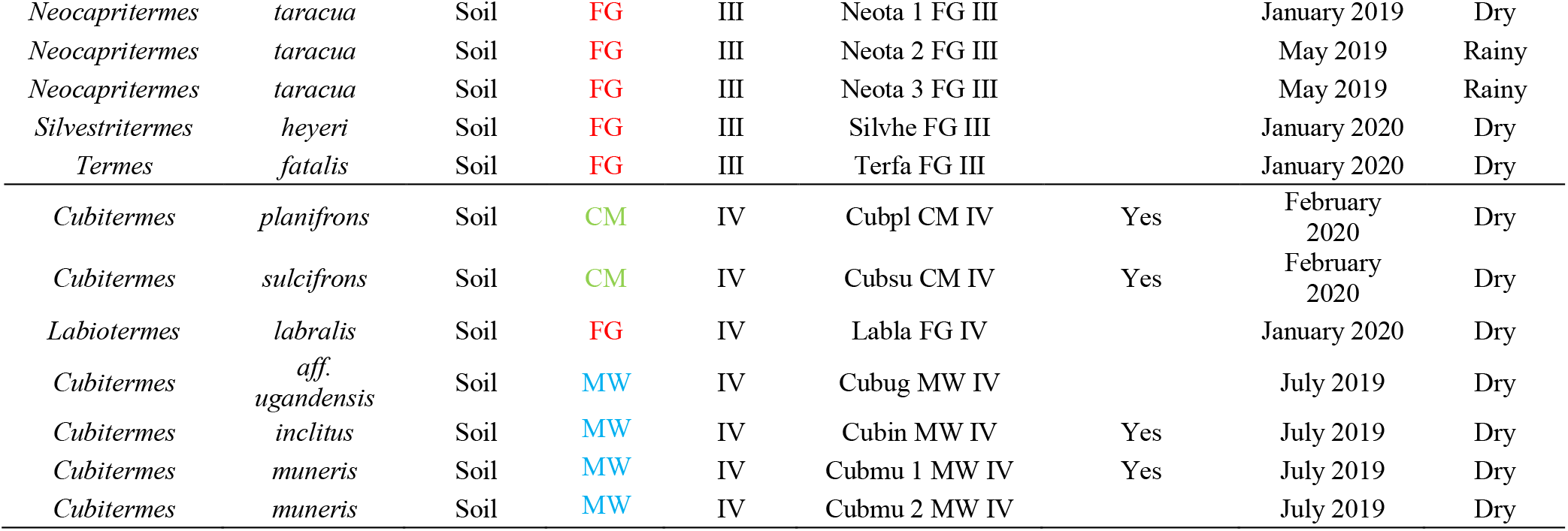
Species identification after COII method. Feeding habits of termites, their geographical origin, and classification according to termite feeding group are shown as well. Note: For metagenomic data analysis after sequencing, we have used the generic Ban1, Ban2, and Ban3 for simplicity purposes, and have the correspondence: Ban1 = Anba 2 FG III; Ban2 = Anba 3 FG III; Ban3 = Anba 4 FG III. Note: Breeds samples were collected in our termite collection at Czech University of Life Sciences, Prague, Czech Republic. N/A: Not applicable; BR: Breeds; CM: Cameroon; FG: French Guiana; MW: Malawi.

| Genus | Species | Feeding habits | Origin | Termite feeding group | Code | Sequenced (metagenome) | Sampling campaign | Rainfall season at sampling |
| --- | --- | --- | --- | --- | --- | --- | --- | --- |
| *Coptotermes* | *formosanus* | Wood | BR | I | Copfor BR I |  | July 2019 | N/A |
| *Nasutitermes* | *sp* | Wood | BR | II | Nassp BR II |  | July 2019 | N/A |
| *Nasutitermes* | *lujae* | Wood | CM | II | Naslu 1 CM II | Yes | July 2019 | Dry |
| *Nasutitermes* | *lujae* | Wood | CM | II | Naslu 2 CM II | Yes | July 2019 | Dry |
| *Nasutitermes* | *lujae* | Wood | CM | II | Naslu 3 CM II | Yes | July 2019 | Dry |
| *Nasutitermes* | *sp* | Wood | CM | II | Nassp CM II | Yes | July 2019 | Dry |
| *Microcerotermes* | *sp* | Wood | CM | II | Micsp CM II |  | January 2020 | Dry |
| *Cephalotermes* | *rectangularis* | Wood | CM | II | Cefre CM II |  | July 2019 | Dry |
| *Constrictotermes* | *cavifrons* | Lichens | FG | II | Conca FG II |  | January 2020 | Dry |
| *Nasutitermes* | *similis* | Wood | FG | II | Nassi 1 FG II |  | May 2019 | Rainy |
| *Nasutitermes* | *similis* | Wood | FG | II | Nassi 2 FG II |  | May 2019 | Rainy |
| *Anoplotermes* | *banksii* | Soil | FG | III | Anba 1 FG III |  | May 2019 | Rainy |
| *Anoplotermes* | *banksii* | Soil | FG | III | Anba 2 FG III | Yes | May 2019 | Rainy |
| *Anoplotermes* | *banksii* | Soil | FG | III | Anba 3 FG III | Yes | May 2019 | Rainy |
| *Anoplotermes* | *banksii* | Soil | FG | III | Anba 4 FG III | Yes | May 2019 | Rainy |
| *Embriatermes* | *neotenicus* | Soil | FG | III | Embne FG III |  | January 2020 | Dry |
| *Neocapritermes* | *taracua* | Soil | FG | III | Neota 1 FG III |  | January 2019 | Dry |
| *Neocapritermes* | *taracua* | Soil | FG | III | Neota 2 FG III |  | May 2019 | Rainy |
| *Neocapritermes* | *taracua* | Soil | FG | III | Neota 3 FG III |  | May 2019 | Rainy |
| *Silvestritermes* | *heyeri* | Soil | FG | III | Silvhe FG III |  | January 2020 | Dry |
| *Termes* | *fatalis* | Soil | FG | III | Terfa FG III |  | January 2020 | Dry |
| *Cubitermes* | *planifrons* | Soil | CM | IV | Cubpl CM IV | Yes | February 2020 | Dry |
| *Cubitermes* | *sulcifrons* | Soil | CM | IV | Cubsu CM IV | Yes | February 2020 | Dry |
| *Labiotermes* | *labralis* | Soil | FG | IV | Labla FG IV |  | January 2020 | Dry |
| *Cubitermes* | *aff. ugandensis* | Soil | MW | IV | Cubug MW IV |  | July 2019 | Dry |
| *Cubitermes* | *inclitus* | Soil | MW | IV | Cubin MW IV | Yes | July 2019 | Dry |
| *Cubitermes* | *muneris* | Soil | MW | IV | Cubmu 1 MW IV | Yes | July 2019 | Dry |
| *Cubitermes* | *muneris* | Soil | MW | IV | Cubmu 2 MW IV |  | July 2019 | Dry |

We have classified the nests according to the termite species habiting them. Ten other colonies were classified in Group II from three locations: our laboratory breeds (BR), Cameroon (CM), and French Guiana (FG). Another group comprising ten colonies originating from French Guiana were classified as Group III. Lastly, Group IV comprised colonies originating from Cameroon, French Guiana, and Malawi (MW). Most of the species from these colonies belong to the *Cubitermes* genus, comprising five different species (**Table 1**).

### 3.2 Untargeted metabolite profiling

Principal component analysis (PCA) was used to find separation and grouping of samples, based on non-target volatiles analysis. Differentiation of samples according to feeding material was observed, while also geographical location seems to influence the distribution, according to the grouping of samples, which is better observed when samples with less than three representatives per genus are discarded (**Fig. 1**).

**Figure 1.**
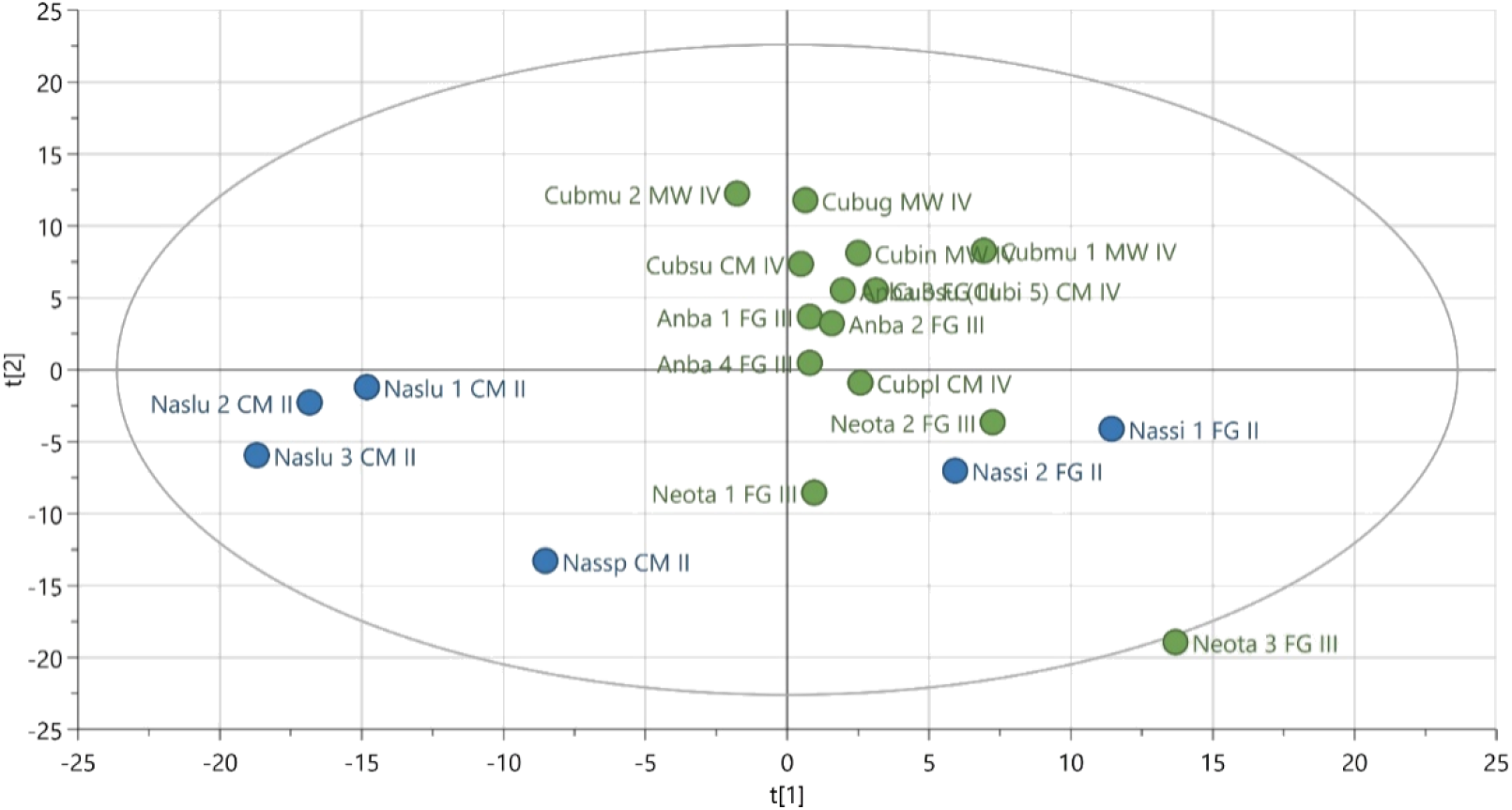
PCA plot using the complete untargeted profiling analysis. Samples are coloured according to feeding habits: green, soil; blue, wood

Among the wood-feeding representatives, we observe a clearer separation in the genus *Nasutitermes* between CM and FG. Three samples belonging to the nests from the same species (Naslu 1-3) grouped together, located at the same Quadrant as another CM wood feeder nest (Nassp CM II). The four nest samples belonging to the soil feeding species (*Anoplotermes banksii,* FG) grouped together.

Samples originating from MW grouped together, occupying the upper part of chart, together with most of the representatives from feeding groups III and IV.

To further investigate differences in feeding, genus, and geographical origin, a Partial least squares discriminant analysis (PLS-DA) was used.

We have observed a clear stratification according to the food origin (**Fig. 2A**) where all the wood-feeding species nests occupy the I and IV quadrants, and the soil feeding species nests are on the II and III. Besides, a second degree of separation is again the geography, where FG wood feeders are on top of Quadrant I, and CM wood feeders lie grouped on Quadrant II. The inverse trend can be observed in samples from soil feeding groups, with MW samples mostly on top, and a gradient of other samples from CM in the middle towards all the FG samples at the bottom. The genus component has been represented in **Fig. 2B** and we find defined groups according to their group, with some overlaps in the soil feeding samples with some influence of the geographical origin (**Fig. 2C**).

**Figure 2.**
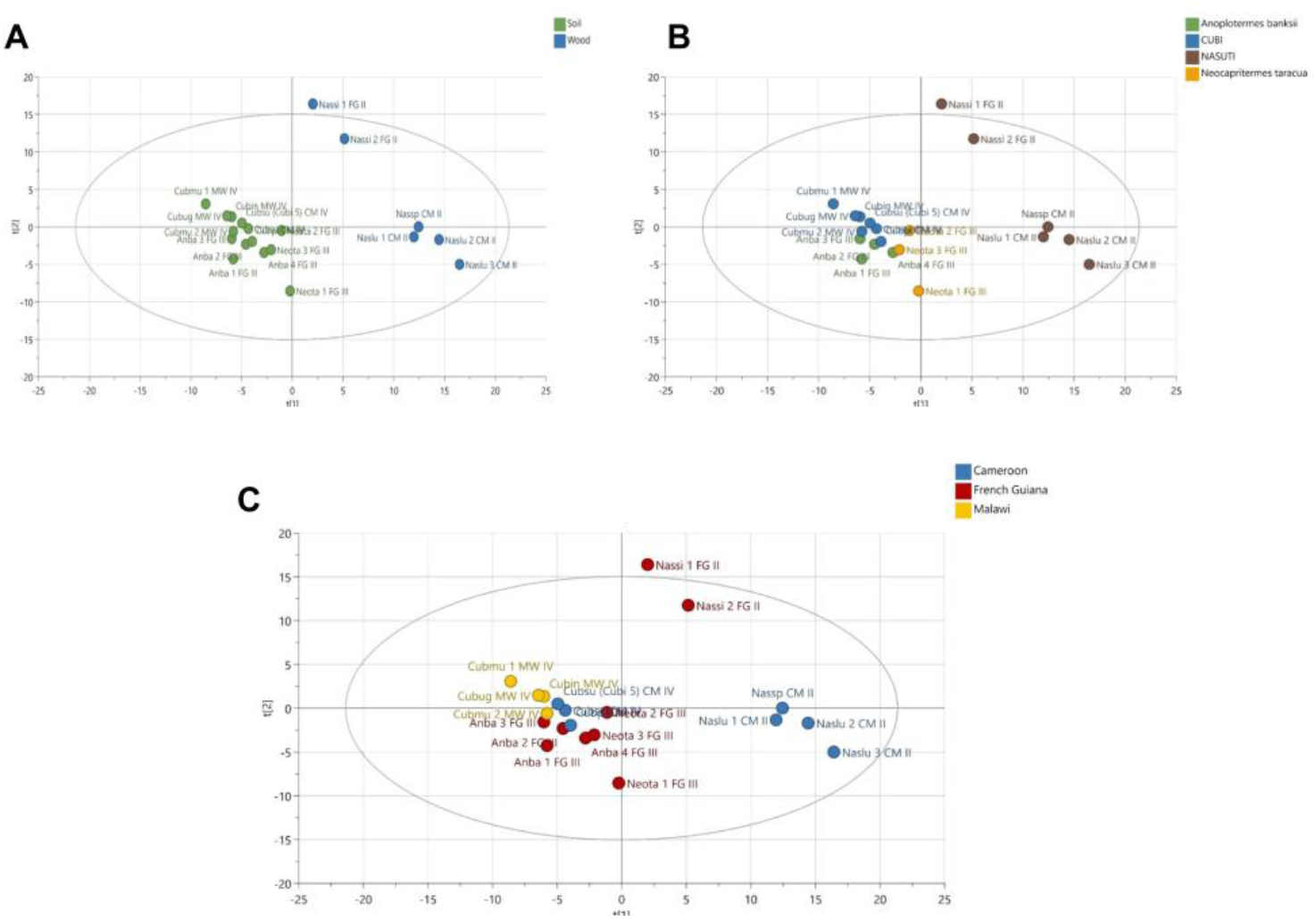
PLS-DA scores plot with classes wood feeders and soil feeders. The scores plot was further coloured according to A) type of food substrate, B) genera, and C) geographical origin.

Lastly, we represented the same OPLS-DA according to the geographical origin, and we observe in a clearer manner the overlaps in the soil feeding groups, where we can still observe a certain stratification according to the location. This stratification is very clear in the case of the wood feeding samples with the clear division between both FG originating samples, and the rest within this genus coming from CM.

The 10 most decisive compounds for separation presented on **Fig. 2** are identified in **Table 2**, where we displayed either their contribution to feeding separation, or genera.

**Table 2.**
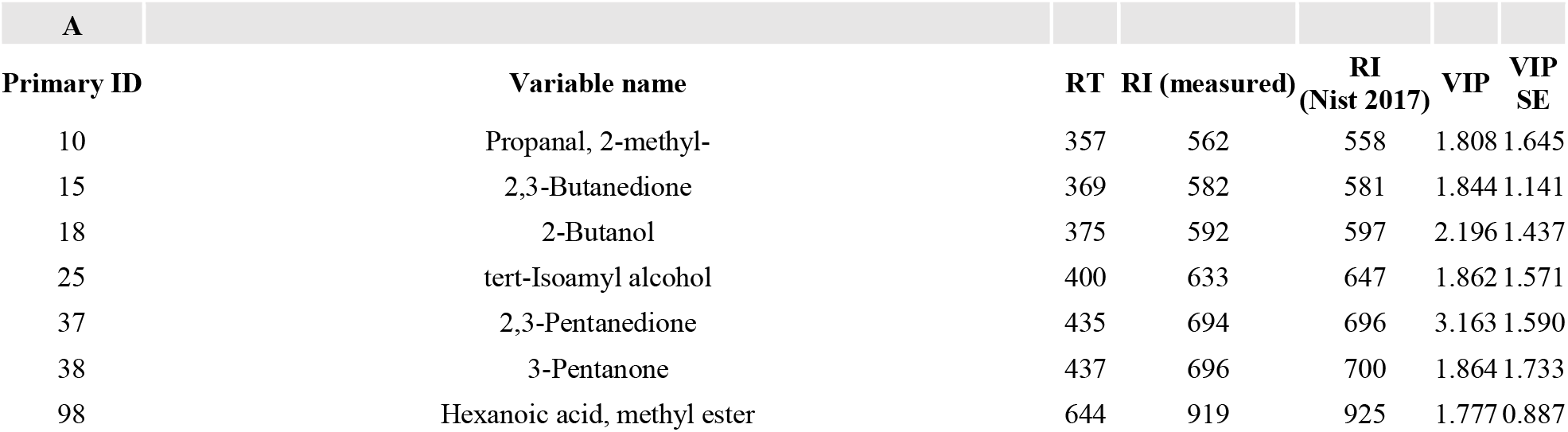
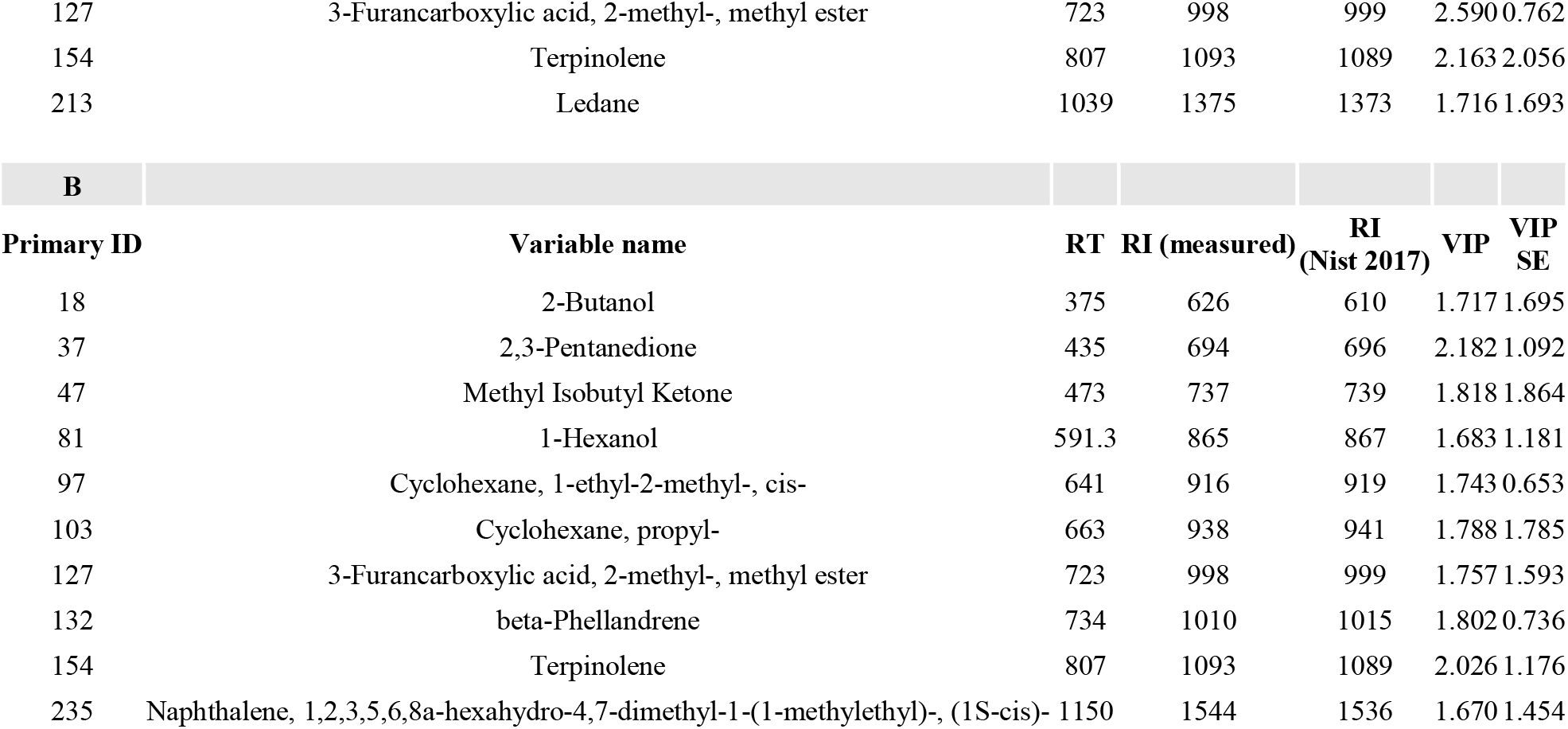
Ten most decisive analytes when considering: A) feeding habits, and B) genera. This table represents the 10 first variables on the VIP analysis. VIP: Variable Importance in Projection; RT: Retention Time; RI: Retention Index; VIP SE: VIP standard error.

### 3.3 Targeted metabolite profiling: Naphthalene content in samples

Naphthalene is polyaromatic hydrocarbon that was observed in the volatile profiling during the non-targeted metabolite analysis (**Fig. 3A**). The highest abundancies were recorded in samples from the genus *Nasutitermes*, *Coptotermes* from BR, and some of the *Cubitermes*. There was certainly a weak presence in many of the samples belonging to Group III, especially the species *Anoplotermes banksii* and the three samples from *Neocapritermes taracua*.

**Figure 3.**
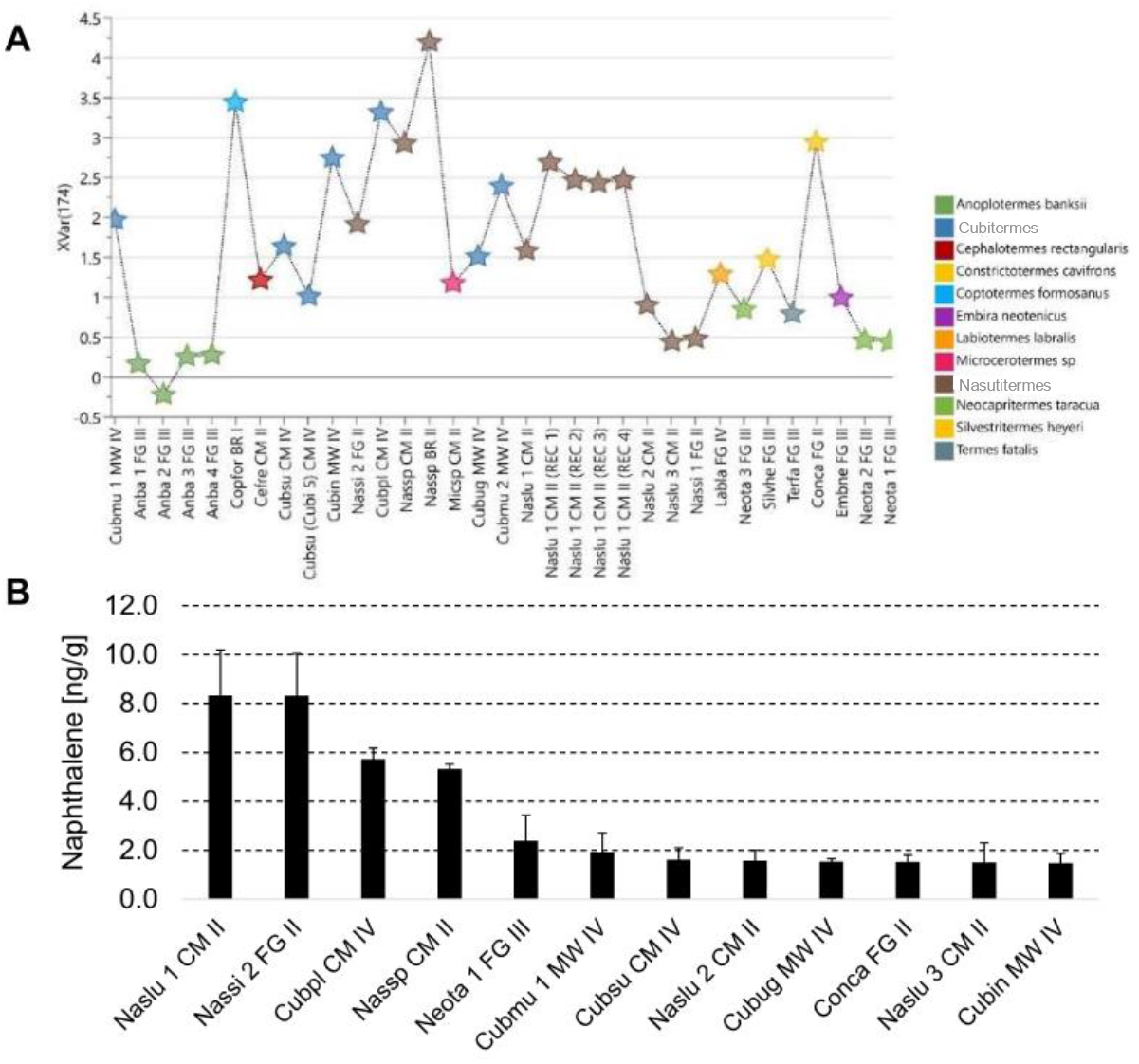
Naphthalene analysis. A) Results from untargeted metabolite profiling. B) Results from targeted metabolite profiling, naphthalene concentration expressed in ng per gram of nest material. Naslu: *Nasutitermes lujae;* Nassi: *Nasutitermes similis;* Cubpl: *Cubitermes planifrons;* Nassp: *Nasutitermes sp.;* Cubmu: *Cubitermes muneris;* Cubsu: *Cubitermes* sulcifrons; Cubug: *Cubitermes aff. ugandensis;* Conca: *Constrictotermes cavifrons;* Cubin: *Cubitermes inclitus;* CM: Cameroon; FG: French Guiana; MW: Malawi.

A quantitative method was applied to determine the exact concentration of this compound in nest material, where the naphthalene concentration ranged from 1.5 to 8 μg/kg (**Fig 3B**). These targeted metabolite results indicate that volatile naphthalene is not generally present in the studied species belonging to Group III, except for the nest material from one of the colonies of *Neocapritermes taracua* (colony No. 1). In keeping with the untargeted results, we have observed a general presence in species belonging to feeding Groups II and IV. In detail, these included colonies belonging to three samples from the genus *Nasutitermes* (Group II, both Cameroon and French Guiana), and the five species belonging to the genus *Cubitermes* (Group IV) that were included within this study (Cameroon and Malawi). Naphthalene was found as well in the colony nest material from *Constrictotermes cavifrons,* belonging to Group II (French Guiana).

When we quantified naphthalene in a targeted approach with a known standard, we observe a similar trend (**Fig. 3B**). The samples belonging to group III do not show detectable amounts of naphthalene (with the exception of one of the *Neocapritermes* samples). There are two main genera where we could detect naphthalene, mainly *Nasutitermes* and *Cubitermes*. We mostly observe samples from the African locations. We observe the presence of this compound markedly in some species of Group II and IV, while in groups I and III is very weak.

After correction with blank, volatile naphthalene was only found in one nest of the FG samples belonging to group III. These results indicate that volatile naphthalene is not generally present in the studied species belonging to Group III, except for the nest material from one of the colonies of *Neocapritermes taracua* (colony No. 1). In contrast, we have observed a general presence in species belonging to feeding Groups II and IV. This observed presence of volatile naphthalene does not seem to be dependant of geographical location, as we have observed it in samples from all localities. There is a notable presence of this compound in colonies belonging to three species from the genus *Nasutitermes* in Group II (both Cameroon and French Guiana) and in the five species belonging to the genus *Cubitermes* in Group IV that were included within this study (Cameroon and Malawi). Naphthalene was found as well in the colony nest material from *Constrictotermes cavifrons*, belonging to Group II (French Guiana).

### 3.4 Bacterial community structure

The 16S amplicon sequencing results of the V4 region have been represented in **Table S1**. We have calculated an average of 131,154 effective tags, after removal of chimera sequences for our samples. These values range from 121,959 tags (*Coptotermes*, BR) to 142,192 non-chimeric tags (NasLu3, CM). In our DNA isolation process, which was carried out with sterility-opened reagents inside of a laminar flow-hood during the entire isolation process, we also included a negative DNA extraction control. In this case, after chimera removal we obtained a total of 36,406 tags.

After the data treatment in Qiime2, we did a comparison of the assigned OTUs according to their representation on each, which can be accessed in **Table S2** (OTUs on each sample) and **Table S3** (feature table). There was a total of 531 OTUs assigned to the negative extraction control, from a total of 19,753 OTUs in our project. The overlap with other samples has been calculated, and from these 532 OTUs, we found overlap with other samples in a total of 106 OTUs (**Table S3**).

We then filtered the sequences after “dada2” step. Practically, all of the sequences were removed, having remained a total of 112 tags (from a total of 36,406 prior to filtering). We have also observed a decrease of sequences in the rest of samples, where the minimum amount of tags belongs to Neota2FG.III with 38,777 tags.

#### 3.4.1 Whole set of samples

We did not find a clear stratification of samples when carrying out a PCOA plot using the Bray-Curtis dissimilarity metric. We then evaluated the α-diversity by the Abundance Based Coverage Estimator (ACE) Index. We found that at a Kruskal-Wallis pairwise comparison there are differences in terms of geography (**Fig. 6**) (MW vs FG, p-value = 0.0146; and BR vs FG, p-value = 0.026). Comparing feeding groups, we found some significant differences (II vs III, p-value = 0.005; and III and IV, p-value = 0.006). When we consider genera, we found also some differences in ACE Index values (*Anoplotermes* vs *Neocapritermes,* p-value = 0.034; *Cubitermes* vs *Neocapritermes,* p-value = 0.02; and *Nasutitermes* vs *Neocapritermes,* p-value = 0.03). When using all groups, Kruskal-Wallis tests indicated differences when considering feeding groups (p-value = 0.007) and geography (p-value = 0.012).

For β-diversity calculation, among other measurements we highlight the results obtained at the Unweighted Unifrac distance. We obtained significant differences between feeding groups III and IV (p-value = 0.039), and the geographical locations FG and MW (p-value = 0.039).

#### 3.4.2 Subset of samples with at least three representatives per genus

We carried out a selection of samples where at least three representatives per genus were present. The calculation of α-diversity was performed with the ACE Index. From a statistical point of view, we found that at a Kruskal-Wallis pairwise comparison there are differences in terms of geography between FG and MW, p-value = 0.031), although this is not significant when taking all groups together (p-value = 0.077). Comparing feeding groups, we found significant differences when comparing II vs III, p-value = 0.045; and III vs IV, p-value = 0.045, and these differences are statistically significant among all groups (p-value = 0.028). When we consider genera, we found also some differences in ACE Index values (*Anoplotermes* vs *Neocapritermes*, p-value = 0.034; *Cubitermes* vs *Neocapritermes,* p-value = 0.02; and Nasutitermes vs *Neocapritermes,* p-value = 0.034), being as well significant among all groups (p-value = 0.03).

The β-diversity was calculated by the Unweighted Unifrac distance, testing the significance for feeding groups through a pairwise permanova (differences between groups II and III, p-value = 0.029; groups III and IV, p-value = 0.005; differences among all groups p-value = 0.001). In the case of geography, the differences were observed between FG and MW (p-value = 0.045), while differences among all groups were of p-value = 0.02. Data with at least three representatives per genus could be compared by genera as well. In this case, we found statistical differences between *Anoplotermes* and *Cubitermes* (p-value = 0.006), and also between *Anoplotermes* and *Nasutitermes* (p-value = 0.006). Lastly, the differences between groups had a p-value = 0.001.

We used then the OTU annotated data in R through the ampvis2 package to create PCA analysis (**Fig. 4**). Results show a clear stratification of samples both by genera and feeding strategy (**Fig. 4A**), while the stratification is less clear when grouping samples by geographical location (**Fig. 4B**). The two principal components of the analysis explain 24.6% and 17.7 % of the variance.

**Figure 4.**
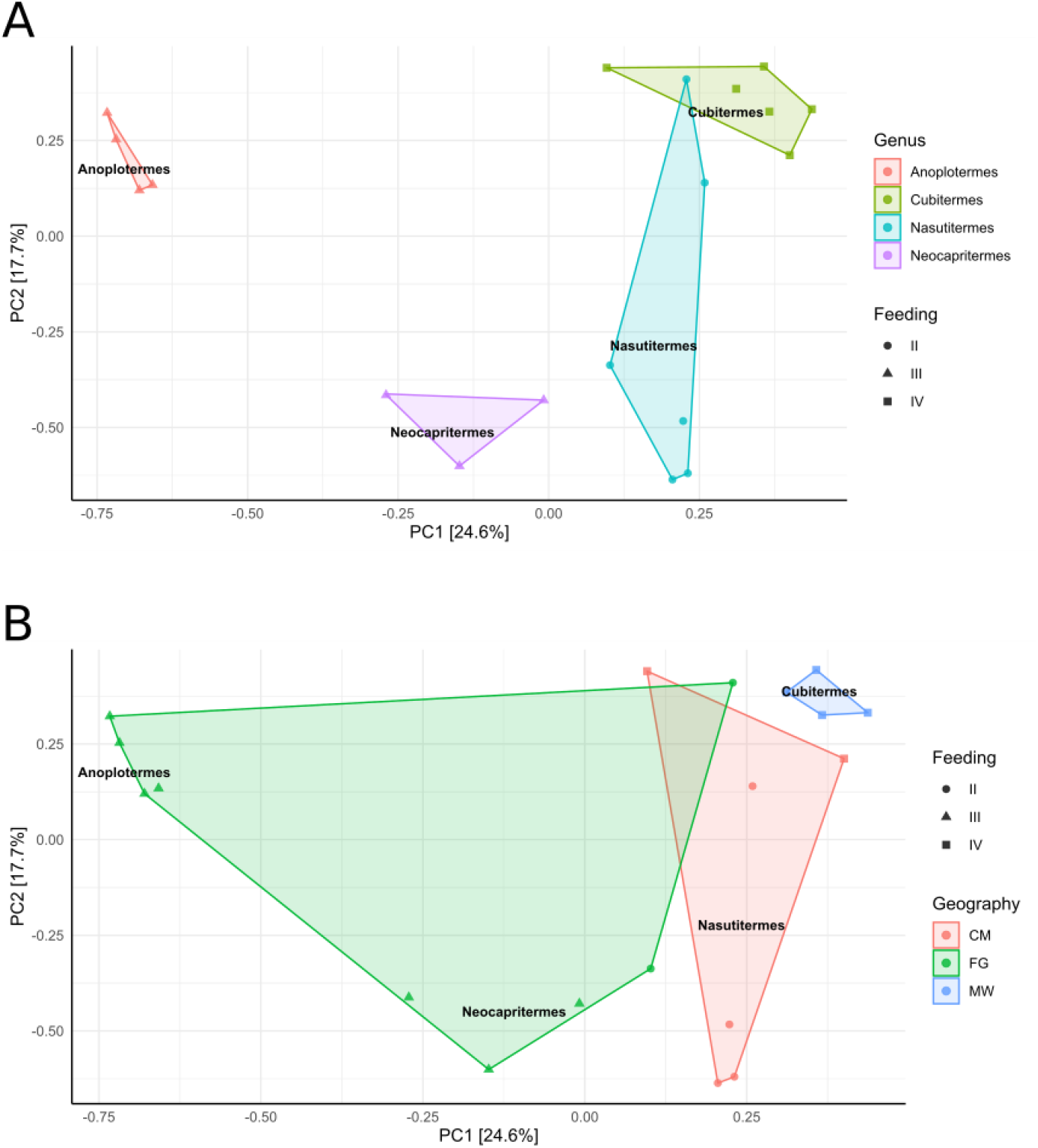
PCA analysis with 16S data after elimination of samples with less than 3 representatives per genus, and the tags from the negative sample control. A) Samples were subjected to Hellinger transformation, sample constrain and colour were “Genus”, shape by Feeding Strategy. B) Samples were subjected to Hellinger transformation, sample constrain was “Genus”, while sample colour was Geography, shape by Feeding Strategy. Genera indications were kept as shown in Fig. 4A.

The phylogenetic assignment of taxa has been represented as a table with the most abundant orders where the samples have been grouped by termite genera with distinction of geographies (**Table 3**). We observe an overall representation of *Actinomycetales, Acidimicrobiales, Clostridiales, Gemmatales, Solirubrobacterales,* being *Spirochaetales* another category well represented in many samples.

**Table 3.** Taxonomic overview of the samples retained for further analyses (with more than 2 representatives per genus). Taxa with over 5% representation have been shaded and bolded. FG: French Guyana; CM: Cameroon; MW: Malawi; Anba: *Anoplotermes banksii;* Cub: *Cubitermes;* NasLu: *Nasutitermes lujae;* Nas: *Nasutitermes;*NasSp: *Nasutitermes* sp.; Neo: *Neocapritermes taracua*.

|  | Anba | Cub<br>(CM) | Cub<br>(MW) | NasLu<br>(CM) | Nas<br>(FG) | NasSp | Neo |
| --- | --- | --- | --- | --- | --- | --- | --- |
| p_Actinobacteria;c_Actinobacteria; o_Actinomycetales | **32.79** | **39.88** | **31.61** | **37.50** | **24.00** | **44.50** | **34.61** |
| Unidentified Bacteria | **19.22** | **6.42** | **4.69** | **6.13** | **11.55** | 3.82 | **6.64** |
| p_Actinobacteria;c_Thermoleophilia; o_Solirubrobacterales | 2.35 | **5.09** | **5.07** | **7.10** | 3.50 | 4.95 | **6.78** |
| p_Actinobacteria;c_Acidimicrobiia; o_Acidimicrobiales | 3.11 | **5.87** | 3.13 | **7.32** | 3.21 | **9.86** | **8.58** |
| p_Firmicutes;c_Clostridia; o_Clostridiales | **10.50** | **5.61** | 4.98 | 4.73 | **5.60** | **10.46** | 2.49 |
| p_Planctomycetes;c_Planctomycetia; o_Gemmatales | 2.74 | 2.66 | **6.16** | 1.16 | **5.26** | 0.31 | **5.35** |
| p_Spirochaetes;c_Spirochaetes; o_Spirochaetales | 3.07 | 2.71 | 1.81 | **5.97** | **8.58** | 1.79 | 1.45 |
| p_Proteobacteria;c_Alphaproteobacteria; o_Rhodospirillales | 2.80 | 4.50 | 3.67 | 4.73 | 4.98 | 3.28 | **5.72** |
| p_Chloroflexi;c_Ktedonobacteria; o_Ktedonobacterales | 0.82 | 0.78 | 3.48 | 0.65 | 0.71 | 0.00 | 0.08 |
| p_Firmicutes;c_Bacilli; o_Bacillales | 1.19 | 0.93 | 1.58 | 0.81 | 0.74 | 1.00 | 2.76 |
| p_Proteobacteria;c_Alphaproteobacteria; o_Sphingomonadales | 0.03 | 1.33 | 1.62 | 1.64 | 0.66 | 0.31 | 0.44 |
| p_Actinobacteria | 1.02 | 0.74 | 0.11 | 0.07 | 0.08 | 0.13 | 0.10 |
| p_Proteobacteria;c_Alphaproteobacteria; o_Rickettsiales | 3.57 | 1.08 | 2.13 | 0.82 | 4.16 | 0.11 | 1.23 |
| p_Firmicutes;c_Bacilli; o_Turicibacterales | 1.71 | 0.87 | 0.78 | 0.40 | 1.14 | 0.66 | 1.18 |
| p_Proteobacteria;c_Gammaproteobacteria; o_Aeromonadales | 0.00 | 0.00 | 0.00 | 0.00 | 0.00 | 2.27 | 0.00 |
| p_Chloroflexi;c_Ktedonobacteria; o_Thermogemmatissporales | 0.08 | 0.26 | 1.86 | 0.06 | 0.11 | 0.00 | 0.21 |
| p_Proteobacteria;c_Betaproteobacteria; o_Burkholderiales | 1.39 | 0.54 | 1.28 | 1.38 | 1.98 | 0.57 | 2.81 |
| p_Acidobacteria;c_Acidobacteriia; o_Acidobacteriales | 1.50 | 1.67 | 2.07 | 1.16 | 2.99 | 0.76 | 2.36 |
| p_Firmicutes;c_Bacilli; o_Lactobacillales | 0.31 | 0.56 | 0.42 | 0.78 | 0.33 | 0.61 | 0.48 |
| p_Verrucomicrobia;c_[Spartobacteria]; o_[Chthoniobacterales] | 0.13 | 0.25 | 0.63 | 0.10 | 0.25 | 0.01 | 0.16 |
| p_Proteobacteria;c_Gammaproteobacteria; o_Xanthomonadales | 0.27 | 1.49 | 1.99 | 1.80 | 2.50 | 0.78 | 1.56 |
| p_Proteobacteria;c_Alphaproteobacteria; o_Ellin329 | 0.23 | 0.28 | 0.27 | 0.22 | 0.50 | 0.13 | 0.49 |
| p_Synergistetes;c_Synergistia; o_Synergistales | 0.53 | 0.13 | 0.18 | 0.18 | 0.24 | 0.09 | 0.09 |
| p_Bacteroidetes;c_Bacteroidia; o_Bacteroidales | 2.13 | 1.40 | 1.01 | 1.13 | 1.34 | 1.02 | 0.33 |
| p_Acidobacteria;c_Solibacteres; o_Solibacterales | 0.38 | 0.73 | 0.81 | 0.55 | 1.40 | 0.62 | 0.95 |
| p_Proteobacteria;c_Alphaproteobacteria; o_Rhizobiales | 0.19 | 0.94 | 1.97 | 1.63 | 1.31 | 0.69 | 1.91 |

LEfSe results are shown in **Fig. 5**. Samples from genus *Neocapritermes* were characterized for the presence of the class Betaproteobacteria, order IS_44. *Nasutitermes* samples was characterized by representatives from the genus *Streptomyces,* down to species *lanatus*. In the case of *Cubitermes,* presence of genus *Rhodococcus,* and members of the family *Intrasporangiaceae*. Lastly, *Anoplotermes* was characterized for the presence of *Chitinophaga*.

**Figure 5.**
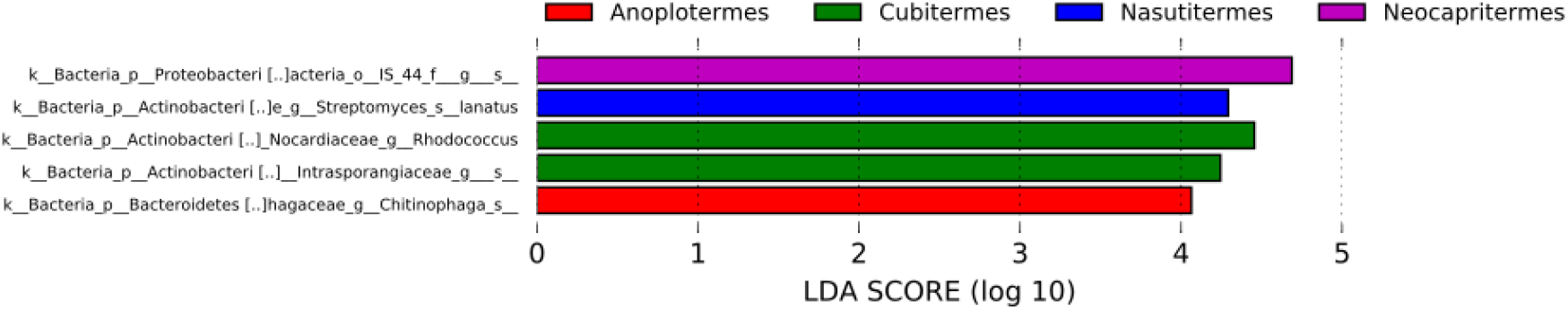
LEfSe results. LDA Score (log 10) for each category per genus is shown.

**Fig. 6.**
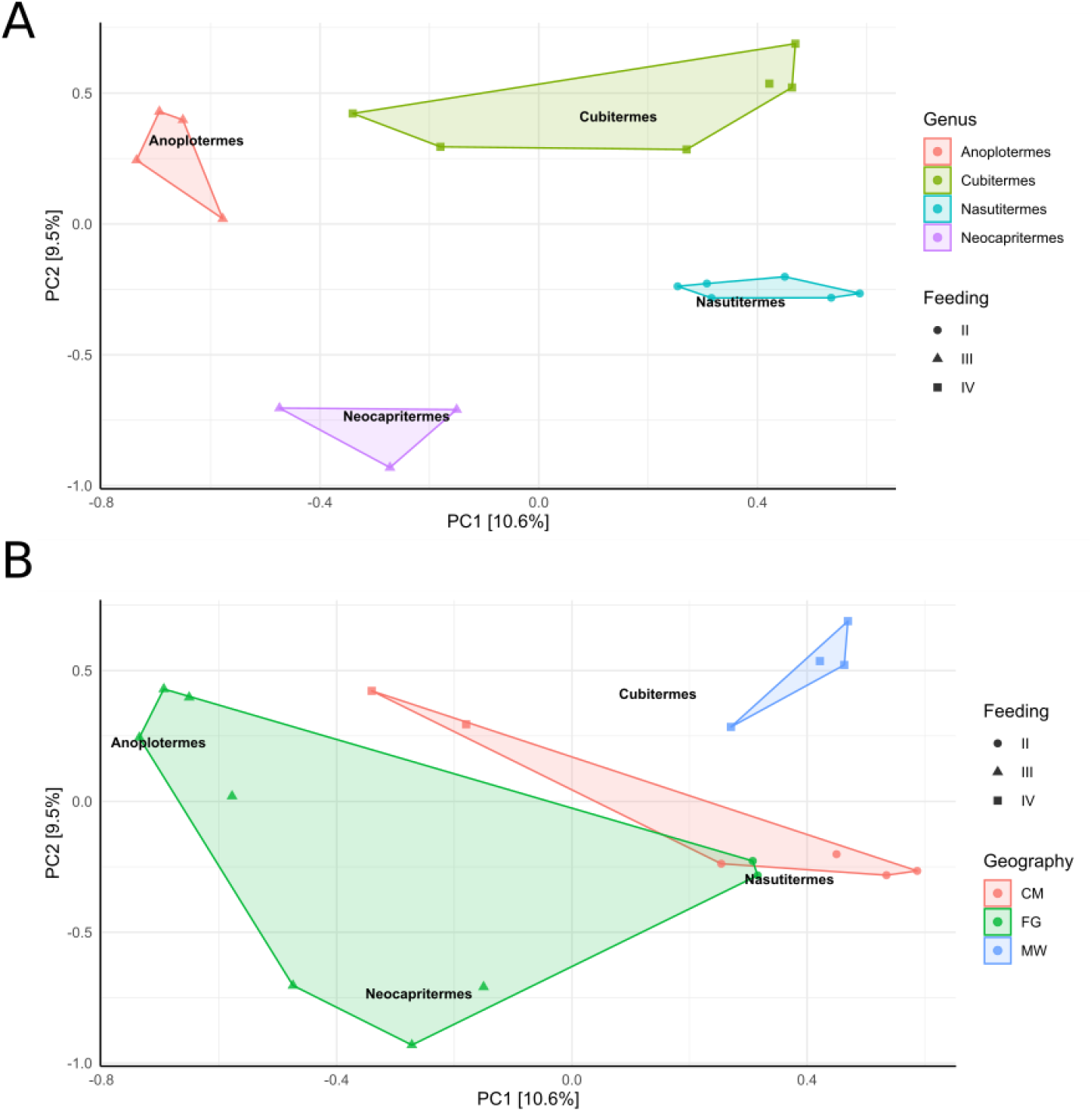
PCA analysis with ITS data after elimination of samples with less than 3 representatives per genus, and the tags from the negative sample control. A) Samples were subjected to Hellinger transformation, sample constrain and colour were “Genus”, shape by Feeding Strategy. B) Samples were subjected to Hellinger transformation, sample constrain was “Genus”, while sample colour was Geography, shape by Feeding Strategy. Genera indications were kept as shown in Fig. 6A.

### 3.5 Fungal community structure

#### 3.5.1 Whole set of samples

The first results when considering all samples are not conclusive (data not shown). We then evaluated the alpha diversity by the ACE Index. We found that at a Kruskal-Wallis pairwise comparisons there are no differences in terms of geography, either as pairwise comparisons or between all groups. Comparing feeding groups, we found some significant differences (II vs III, p-value = 0.045; II and IV, p-value = 0.003; III and IV, p-value = 0.032; between groups p-value =0.007). When we consider genera, we found also some differences in ACE Index values (*Anoplotermes* vs *Nasutitermes*, p-value = 0.018; *Cubitermes* vs *Nasutitermes*, p-value = 0.01; and *Nasutitermes* vs *Neocapritermes,* p-value = 0.017), but not between groups (p-value = 0.05).

The beta diversity was evaluated by the Unweighted Unifrac distance, found significant differences for feeding groups (II and III, p-value = 0.001; II and IV, p-value = 0.006; III and IV, p-value = 0.012), and geography (CM and FG, p-value = 0.005; FG and MW, p-value = 0.001).

#### 3.5.2 Subset of samples with more representatives per genus

We proceeded according to the previous section, and have selected only those samples where more than two representatives are present per genus.

We evaluated the alpha diversity by the ACE Index. We found that at a Kruskal-Wallis pairwise comparison in terms of geography there were no differences among any of the locations, at a level of significance α ≤ 0.05. Comparing feeding groups, we found significant differences when comparing II vs III (p-value = 0.004), between groups III vs IV (p-value = 0.01), and lastly at the whole groups’ comparison (p-value = 0.006). When we consider genera, we found also some differences in ACE Index values (*Anoplotermes* vs *Nasutitermes*, p-value = 0.019; *Cubitermes* vs *Nasutitermes,* p-value = 0.01; *Nasutitermes* vs *Neocapritermes,* p-value = 0.02; and all groups comparison, p-value = 0.012).

The beta diversity was calculated by the Unweighted Unifrac distance, testing the significance for Feeding groups through a pairwise permanova, where we found differences between groups II and III (p-value = 0.003), groups II and IV (p-value = 0.006), and groups III and IV (p-value = 0.003).

In the case of geography, the differences in β-diversity were observed between CM and FG (p-value = 0.025), and FG vs MW (p-value = 0.002). Data with at least three representatives per genus could be compared by genera as well. In this case, we found statistical differences between *Anoplotermes* and *Nasutitermes* (p-value = 0.007), and also between *Anoplotermes* and *Neocapritermes* (p-value = 0.030). Additional differences exist between *Cubitermes* and *Nasutitermes* (p-value = 0.005), and *Cubitermes* vs *Neocapritermes* (p-value = 0.009). Lastly, there were as well statistical differences between *Nasutitermes* and *Neocapritermes* (p-value = 0.013).

A PCA plot was built in R through the ampvis2 package (**Fig. 6**). Results show a clear stratification of samples especially by genera, and then and feeding strategy (**Fig. 6A**), while the stratification is less clear when grouping samples by geographical location (**Fig. 6B**). The two principal components of the analysis explain 10.6% and 9.5 % of the variance.

The phylogenetic assignment of taxa has been represented as a table with the most abundant orders where the samples have been grouped by termite genera with distinction of geographies (**Table 4**). We observe an overall representation of the phylum *Ascomycota,* among them the class *Sordariomycetes* with the order *Hypocreales,* and the class *Dothideomycetes*. Chlorophyceae were relevant in *Cubitermes* (CM). OTUs belonging to *Basidiomycota* were also found but in much minor proportion than *Ascomycota*.

**Table 4.** Taxonomic overview of the samples retained for further analyses (with more than 2 representatives per genus). Taxa with over 5% representation have been shaded and bolded. FG: French Guyana; CM: Cameroon; MW: Malawi; Anba: *Anoplotermes banksii;* Cub: *Cubitermes;* NasLu: *Nasutitermes lujae;* Nas: *Nasutitermes;*NasSp: *Nasutitermes* sp.; Neo: *Neocapritermes taracua*.

|  | Anba | Cub<br>(CM) | Cub<br>(MW) | Nas<br>(FG) | NasLu<br>(CM) | NasSp | Neo |
| --- | --- | --- | --- | --- | --- | --- | --- |
| p_*Ascomycota*; c_*Sordariomycetes*;__ | **31.30** | 3.74 | **6.20** | 3.18 | **9.74** | **81.75** | 0.98 |
| p_*Ascomycota*; c_*Sordariomycetes*; o_*Hypocreales* | **16.43** | **28.30** | **25.57** | **31.53** | **53.69** | **6.01** | **7.66** |
| p_*Ascomycota*; c_*Dothideomycetes*;__ | **33.94** | **40.02** | **30.97** | **12.03** | 2.79 | 3.75 | **37.95** |
| p_*Ascomycota* | **6.45** | **6.26** | **12.87** | **28.92** | **5.69** | **5.91** | **23.25** |
| Unidentified Eukaryote | **7.82** | **11.24** | **10.01** | **7.10** | **22.79** | 1.68 | **16.94** |
| p_*Chlorophyta*; c_*Chlorophyceae*; o_*Chlorophyceae* | 0.55 | **5.03** | 0.55 | 0.03 | 0.00 | 0.00 | 0.00 |
| p_*Phragmoplastophyta*; c_*Embryophyta*; o_*Magnoliophyta* | 0.07 | 0.00 | 0.00 | **5.08** | 0.00 | 0.00 | 0.02 |
| p_*Basidiomycota*; c_*Exobasidiomycetes* | 0.02 | 0.01 | 0.02 | 0.00 | 0.01 | 0.01 | **6.47** |
| p_*Basidiomycota*; c_*Agaricomycetes*; o_*Incertae_Sedis* | 0.02 | 0.00 | 0.01 | 4.81 | 0.00 | 0.00 | 0.00 |
| p_*Ascomycota*; c_*Dothideomycetes*; o_*Pleosporales* | 0.12 | 0.21 | 4.77 | 0.08 | 0.02 | 0.00 | 3.22 |
| p_*Ascomycota*; c_*Dothideomycetes*; o_*Capnodiales* | 1.04 | 1.90 | 1.94 | 4.20 | 0.21 | 0.00 | 1.38 |
| Unassigned_ | 0.61 | 0.66 | 1.94 | 0.11 | 4.83 | 0.86 | 0.72 |
| p_*Basidiomycota*; | 1.08 | 0.76 | 0.74 | 2.61 | 0.03 | 0.01 | 0.43 |
| p_*Chlorophyta*; | 0.15 | 1.20 | 2.92 | 0.12 | 0.00 | 0.00 | 0.00 |
| p_*Ascomycota*; c_*Eurotiomycetes*; o_*Chaetothyriales* | 0.03 | 0.31 | 0.57 | 0.00 | 0.00 | 0.00 | 0.90 |

The LEfSe analysis was carried out, and we found no significant taxa.

### 3.6 Metagenomic sequencing

#### 3.6.1 Cleaning, assembly, SqueezeMeta pipeline

We obtained a total of 625,536,948 raw reads (**Table 5**). Clean quality control passed reads were digitalized as described in methodology, and used as input in SqueezeMeta for the first assembly. This assembly yielded a 5,632,566 contigs, totalling 4,093,561,329 bp, with a minimum of 200 bp, and maximum length of 146,620 bp, average size of 726.77 bp, and N50 of 776 bp.

**Table 5.** Statistics on metagenomic sequencing. Raw reads: total amount of reads of raw data, each four lines taken as one unit. For paired-end sequencing, it equals the amount of read1 and read2, otherwise it equals the amount of read1 for single-end sequencing. Seq.: sequencing; Geo.: Geographical. Raw data: (Raw reads) * (sequence length), calculating in G. For paired-end sequencing like PE150, sequencing length equals 150, otherwise it equals 50 for sequencing like SE50. Effective: (Clean reads/Raw reads) * 100. Error: base error rate. Q20, Q30: (Base count of Phred value > 20 or 30) / (Total base count). GC: (G & C base count) / (Total base count)

| Geo. location | Species | Code | Sample seq. code | Data treatment code | Raw reads | Raw data | Effective (%) | Error (%) | Q20 (%) | Q30 (%) | GC (%) |
| --- | --- | --- | --- | --- | --- | --- | --- | --- | --- | --- | --- |
| FG | *Anoplotermes banksii* | Anba 2 FG III | Ban2 | Ban1 | 53,288,002 | 8 | 99.67 | 0.02 | 98.3 | 95.44 | 66 |
|  |  | Anba 3 FG III | Ban3 | Ban2 | 55,905,810 | 8.4 | 99.7 | 0.02 | 98.4 | 95.59 | 65.8 |
|  |  | Anba 4 FG III | Ban4 | Ban3 | 66,039,800 | 9.9 | 99.59 | 0.03 | 97.6 | 93.83 | 68.3 |
| CM | *Nasutitermes lujae* | Naslu 1 CM II | Nal1 | NasLu1 | 63,479,926 | 9.5 | 99.68 | 0.02 | 98 | 94.97 | 68.3 |
|  |  | Naslu 2 CM II | Nal2 | Naslu2 | 53,196,396 | 8 | 99.67 | 0.02 | 98.1 | 95.17 | 68.3 |
|  |  | Naslu 3 CM II | Nal3 | Naslu3 | 63,131,466 | 9.5 | 99.66 | 0.02 | 98.1 | 95.14 | 67.5 |
|  | *Nasutitermes sp.* | Nassp CM II | NasSp | NasSp | 56,112,754 | 8.4 | 99.7 | 0.02 | 98.2 | 95.34 | 68.1 |
|  | *Cubitermes planifrons* | Cubpl CM IV | CubPL | CubPl | 49,692,682 | 7.5 | 99.74 | 0.02 | 98.2 | 95.35 | 67.1 |
|  | *Cubitermes sulcifrons* | Cubsu CM IV | CubSu | CubSu | 55,052,860 | 8.3 | 99.71 | 0.02 | 98.2 | 95.25 | 67 |
| MW | *Cubitermes muneris* | Cubmu 1 MW IV | MLW3 | CubMu | 56,193,314 | 8.4 | 99.75 | 0.02 | 98.2 | 95.24 | 66.4 |
|  | *Cubitermes inclitus* | Cubin MW IV | MLW2 | CubIn | 53,443,938 | 8 | 99.7 | 0.02 | 98.2 | 95.34 | 69.1 |
| Total |  |  |  |  | 625,536,948 |  |  |  |  |  |  |

The reads were assigned taxonomically, and the most abundant hits per sample are shown in **Table 6**, while the statistics on Open Reading Frames (ORFs) is presented on **Table 7**.

**Table 6.**
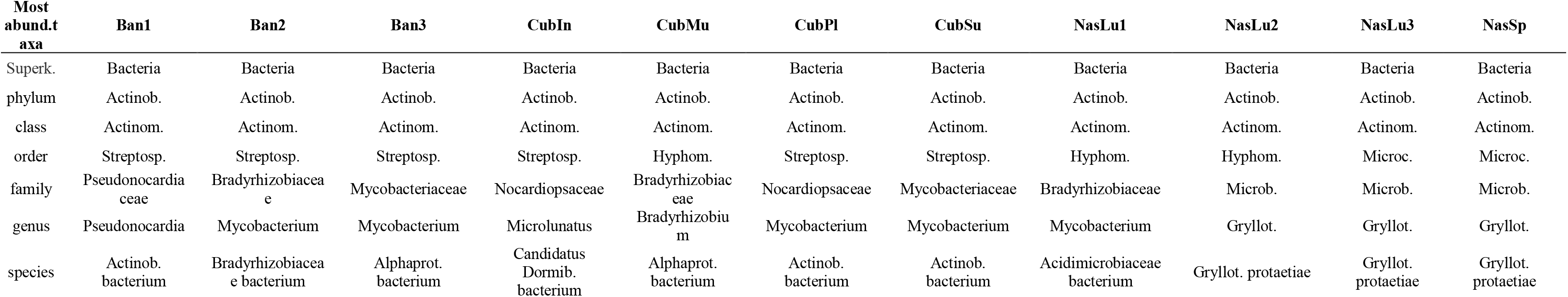
Statistics on metagenomic sequencing. Abund.: abundant; Superk: superkingdom; Actinob.: Actinobacteria; Actinom.: Actinomycetia; Streptosp.: Streptosporangiales; Hyphom.: Hyphomicrobiales; Microc.: Micrococcales.; Microb.: Microbacteriaceae.; Gryllot.: Gryllotalpicola.; Dormib.: Dormibacteraeota; Alphaprot.: Alphaproteobacteria. Note: “Data treatment code” (Table 4) has been used to name samples, for simplicity purposes.

| Most abund.t axa | Ban1 | Ban2 | Ban3 | CubIn | CubMu | CubPI | CubSu | NasLu1 | NasLu2 | NasLu3 | NasSp |
| --- | --- | --- | --- | --- | --- | --- | --- | --- | --- | --- | --- |
| Superk. | Bacteria | Bacteria | Bacteria | Bacteria | Bacteria | Bacteria | Bacteria | Bacteria | Bacteria | Bacteria | Bacteria |
| phylum | Actinob. | Actinob. | Actinob. | Actinob. | Actinob. | Actinob. | Actinob. | Actinob. | Actinob. | Actinob. | Actinob. |
| class | Actinom. | Actinom. | Actinom. | Actinom. | Actinom. | Actinom. | Actinom. | Actinom. | Actinom. | Actinom. | Actinom. |
| order | Streptosp. | Streptosp. | Streptosp. | Streptosp. | Hyphom. | Streptosp. | Streptosp. | Hyphom. | Hyphom. | Microc. | Microc. |
| family | Pseudonocardia<br>ceae | Bradyrhizobiacea<br>e | Mycobacteriaceae | Nocardiopsaceae | Bradyrhizobiac<br>eae | Nocardiopsaceae | Mycobacteriaceae | Bradyrhizobiaceae | Microb. | Microb. | Microb. |
| genus | Pseudonocardia | Mycobacterium | Mycobacterium | Microlunatus | Bradyrhizobiu<br>m | Mycobacterium | Mycobacterium | Mycobacterium | Gryllot. | Gryllot. | Gryllot. |
| species | Actinob.<br>bacterium | Bradyrhizobiacea<br>e bacterium | Alphaprot.<br>bacterium | Candidatus<br>Dormib.<br>bacterium | Alphaprot.<br>bacterium | Actinob.<br>bacterium | Actinob.<br>bacterium | Acidimicrobiaceae<br>bacterium | Gryllot. protaetiae | Gryllot.<br>protaetiae | Gryllot.<br>protaetiae |

**Table 7.**
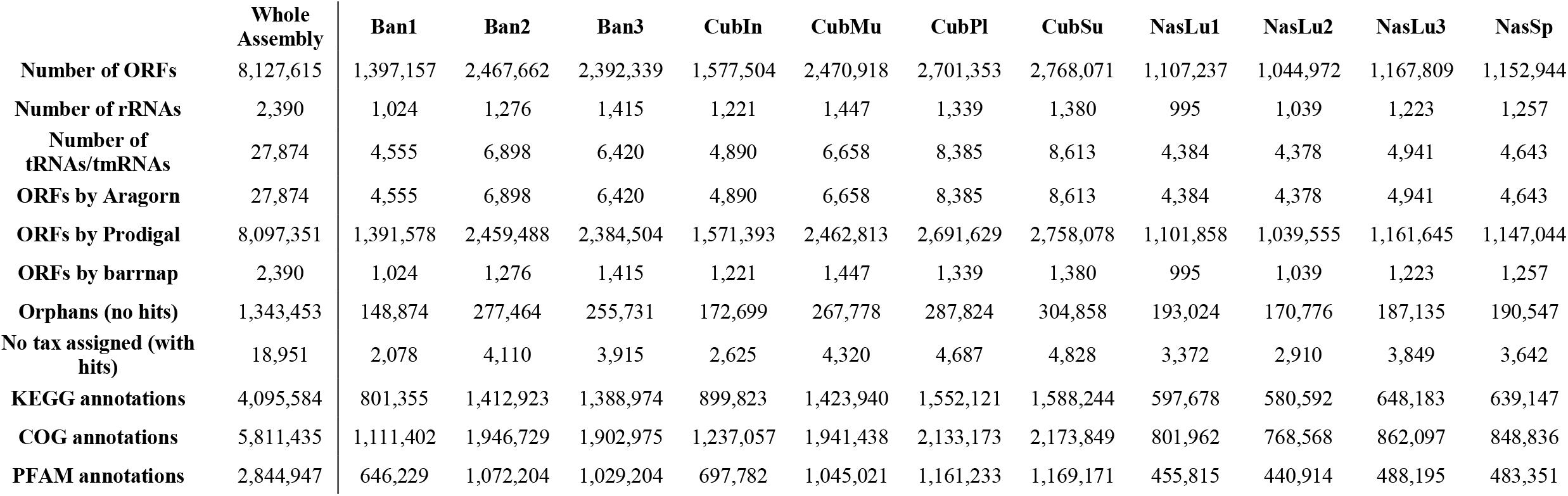
Statistics on ORFs after SqueezeMeta pipeline.

Binning rendered a total of 47 bins (**Table 8**), with 33 over 50% completeness, from which 16 ≥75%, and finally 2 over 90% completeness. Twenty-two bins had a contamination lower than 10% and 3 over 50%.

**Table 8.** Statistics on bins after SqueezeMeta pipeline. Qual.: quality; contam.: contamination.

|  |  |
| --- | --- |
| Number of bins | 47 |
| Complete $\geq 50\%$ | 33 |
| Complete $\geq 75\%$ | 16 |
| Complete $\geq 90\%$ | 2 |
| Contamination $< 10\%$ | 22 |
| Contamination $\geq 50\%$ | 3 |
| Congruent bins | 7 |
| Disparity $>0$ | 40 |
| Disparity $\geq 0.25$ | 21 |
| Hi-qual. bins ($>90\%$ complete, $<10\%$ contam.) | 1 |
| Good-qual. bins ($>75\%$ complete, $<10\%$ contam.) | 3 |

#### 3.6.2 Overall characterization of samples

We used the TPM values for all KEGG annotated genes, to evaluate the general structure of samples and their relationships through non-metric multidimensional scaling (NMDS), which was carried out in R (**Fig. 7**). We observe that all *Nasutitermes* samples cluster together, while the other two groups of nests belonging to genera having a soil feeding strategy are in the same area, although having two dissimilar patterns.

**Fig. 7.**
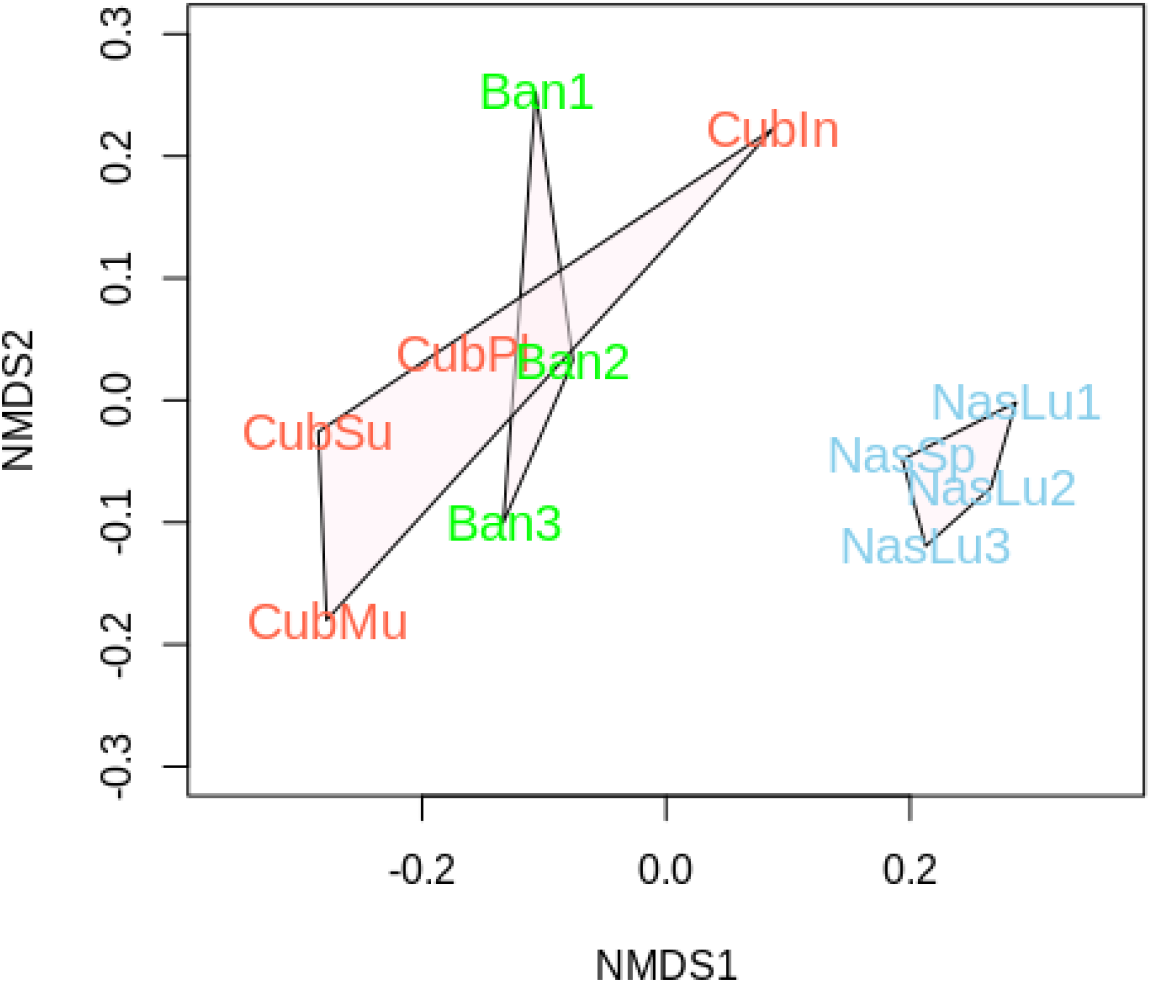
Non-metric multidimensional analysis considering all the samples sequenced in this study (metagenomic shotgun sequencing).

We then analysed the taxonomic signature of samples after metagenomic sequencing. The level Phylum/genus was represented as a bar chart in **Fig. 8**.

**Fig. 8.**
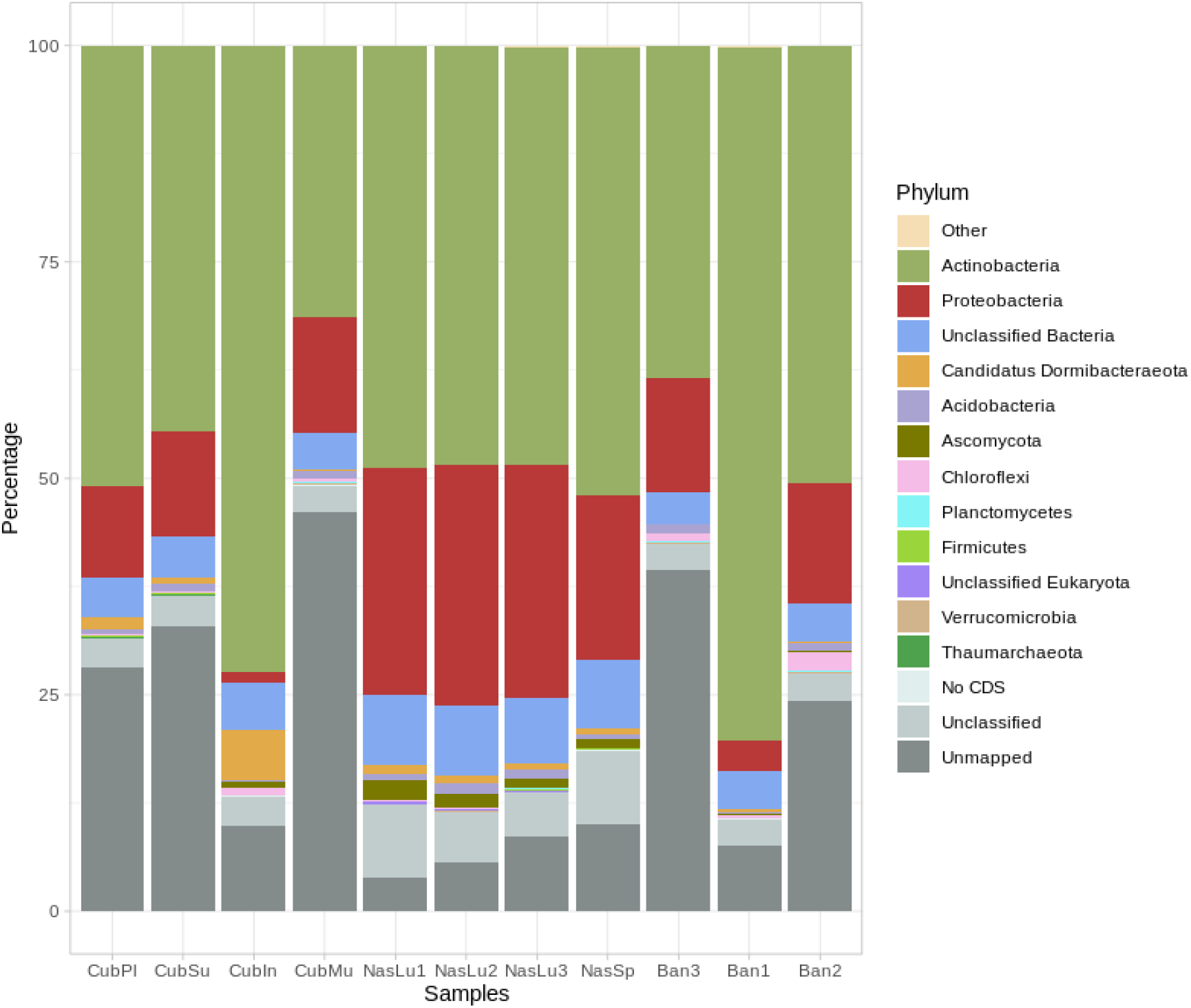
Taxonomic representation at the phylum level in those samples submitted for metagenomic sequencing. Percentages were calculated in through SQMtools after SqueezeMeta pipeline. Phylum levels were obtained after annotation of sequences.

We can observe that the four samples belonging to the genus *Nasutitermes* (all originating in CAM) have a similar profile at the genus level, while the two soil-feeding genera (*Cubitermes*, from MW and CAM; *Anoplotermes*, FG), despite differences at the species level, present a closer profile. An outstanding exception is the biological replicate Ban1, with a lower representation of Proteobacteria than the rest of samples from this genus (all samples as well belonging to the species *Anoplotermes banksii*). For *Nasutitermes*, we find differences between the three *Nasutitermes lujae* replicates, and *Nasutitermes* sp. The samples belonging to the genus *Cubitermes* have a more variable profile. Among those, CubPl and CubSu belong to the CM sampling site, while CubIn and CubMu originate in MW.

We further carried out a subset of the present genera belonging to the “aromatic AA” representation (“Phenylalanine, tyrosine, and tryptophan biosynthesis”) (**Fig. 9**). The most important genera for soil feeders seem to lie within an unclassified *Streptosporangiales* genus with higher abundance for *Cubitermes* and *Anoplotermes*, while *Nasutitermes* had higher representation of *Microlunatus*, an unclassified genus within *Microbacteriaceae*, and *Gryllotalpicola*.

**Fig. 9.**
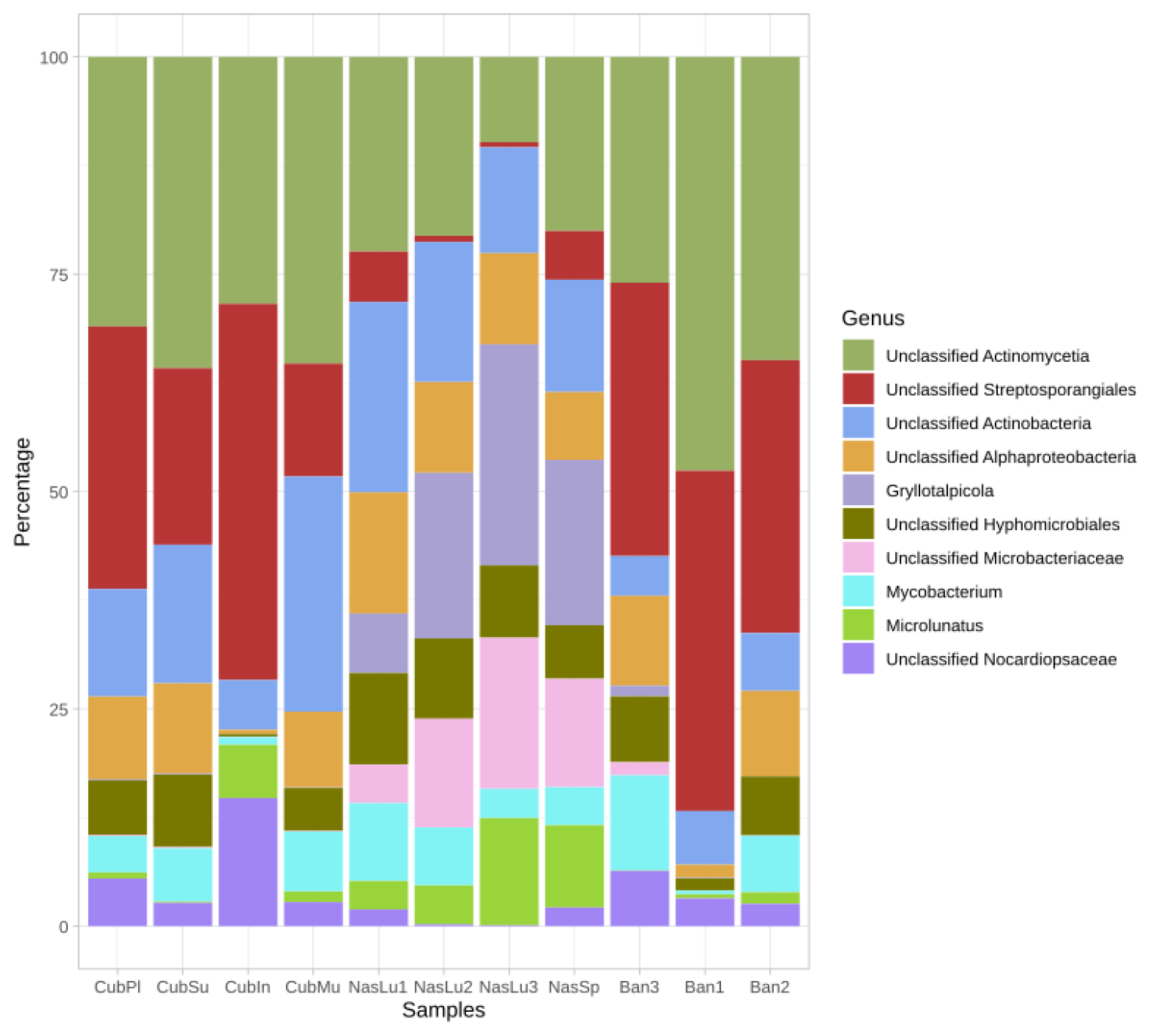
Taxonomic representation at the genus level in those samples submitted for metagenomic sequencing, which belong to the subset “aromatic_aa” (‘Phenylalanine, tyrosine and tryptophan biosynthesis’). Percentages were calculated in through SQMtools after SqueezeMeta pipeline. Genus levels were obtained after annotation of sequences.

A LDA Effective Size (LEfSe) was calculated using the PFAM annotation, and differences were found among the three represented genera. We obtained 61 classes of annotated genes that are specific biomarkers for each of the three (**Fig. 10, Table 9**). *Anoplotermes* was characterized by several biomarkers, among the first PF00196: Bacterial regulatory proteins luxR family, PF13424: Tetratricopeptide repeat, PF03704: Bacterial transcriptional activator domain, and PF13191: AAA ATPase domain. *Cubitermes* had PF00872: Transposases, Mutator family, and PF0084: sulfatase as the only biomarker categories found. One of the most represented biomarker categories in the genus *Nasutitermes* was PF00296: Luciferase-like monooxygenase followed by PF02515: CoA transferase family III, PF00873: AcrB/AcrD/AcrF family, and PF00171: Aldehyde dehydrogenase family.

**Fig. 10.**
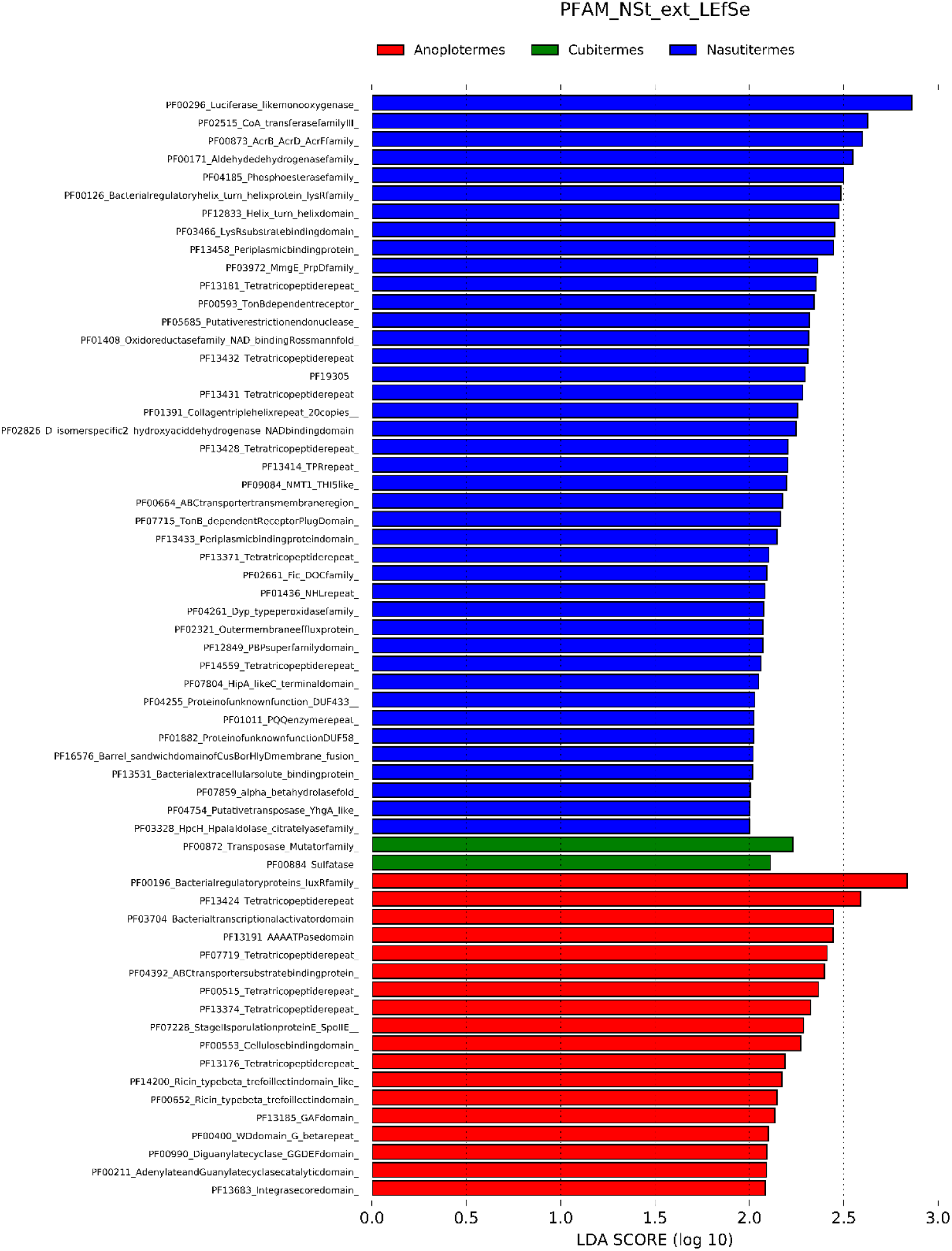
LEfSe analysis for the contigs annotated in PFAM at. LDA Score (log_10_) for each category per genus.

**Table 9.**
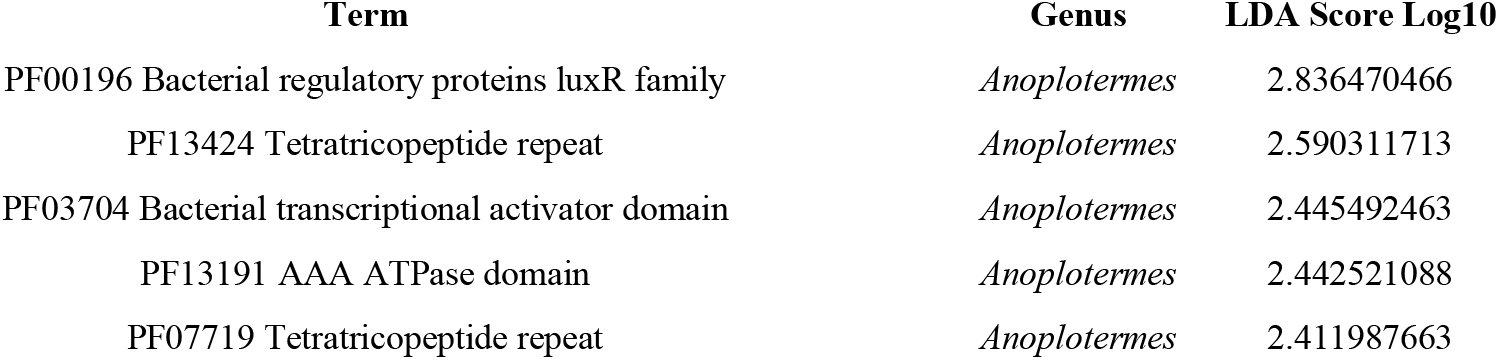
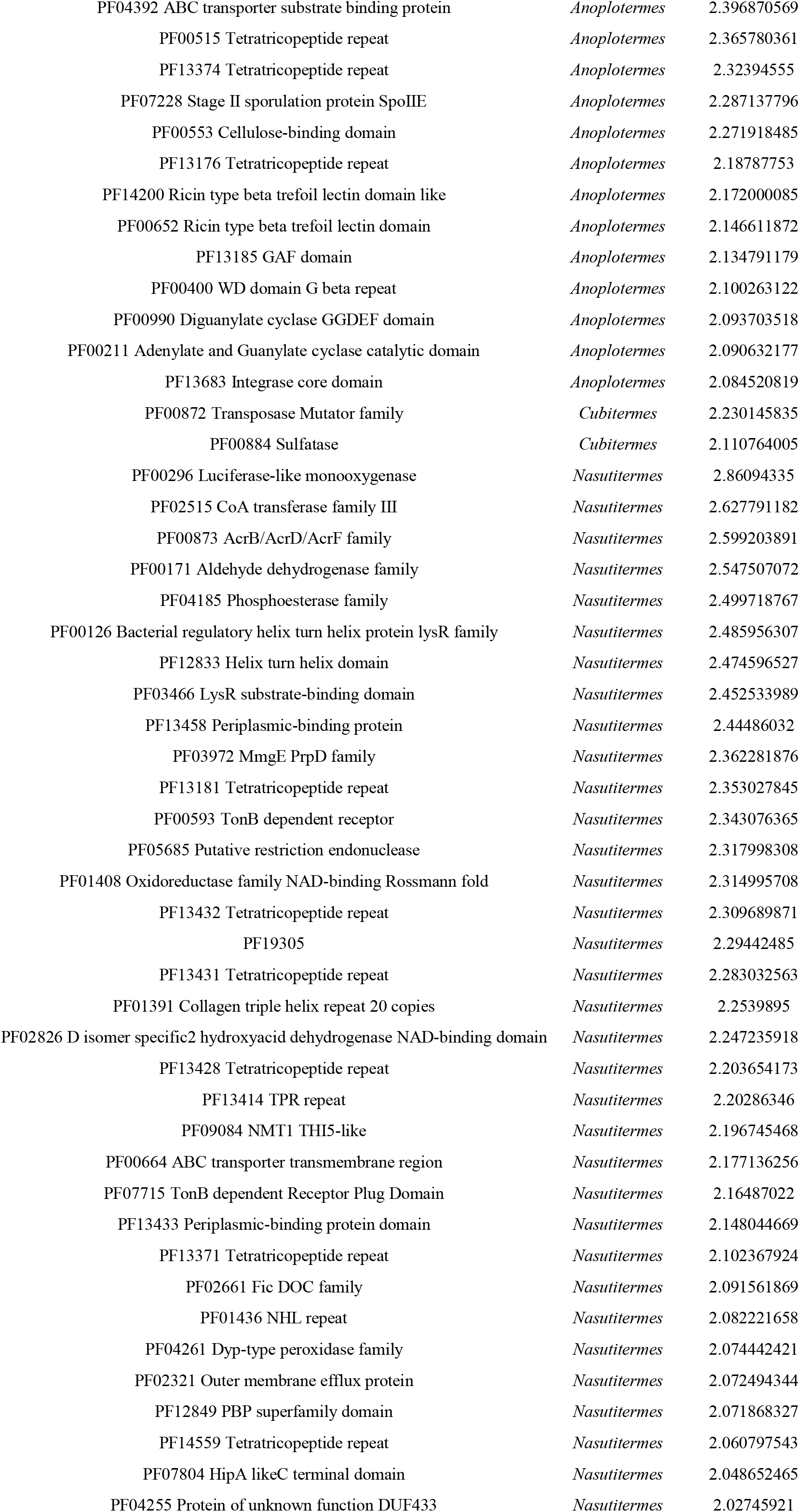
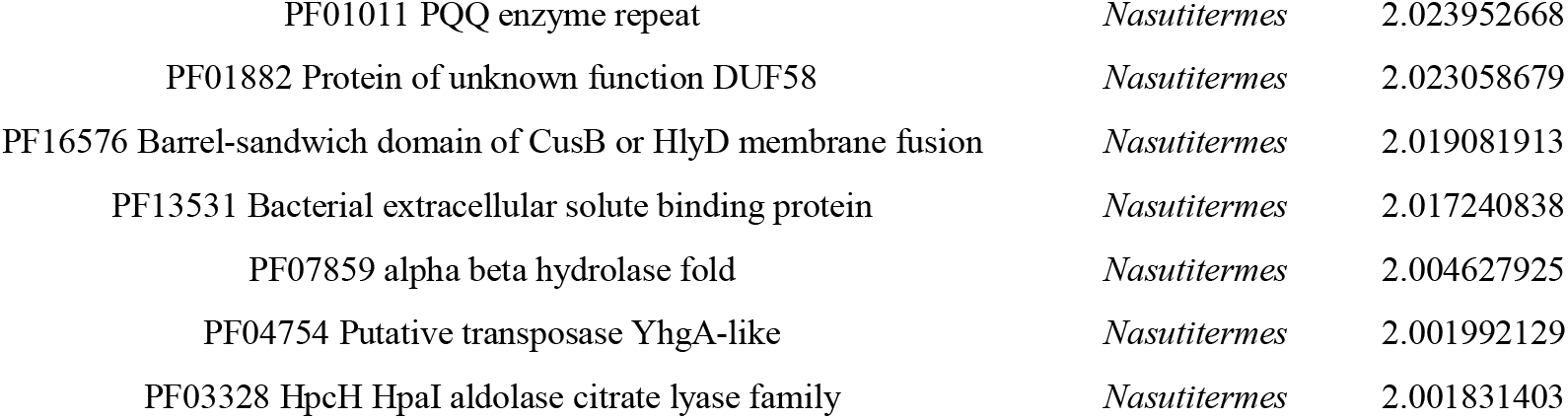
LEfSe biomarker categrories for the three genera included in the metagenomic sequencing.

#### 3.6.3 Differential gene content analysis

We further evaluated the differences among samples in a similar way to transcriptomics Differentially Expressed Genes. In this case, we termed them “Differentially Present Genes” since we are studying the metagenome. Firstly, we approached the structure of the data through a hierarchical clustering using Euclidean distance (**Fig. 11A**). We find two clearly separated nodes, one with all *Nasutitermes* samples (belonging to nests of wood feeding termites from the genus *Nasutitermes*, CAM), and those belonging to the soil feeding genera forming the second group. In the second group, there is as well a subdivision in two nodes. One comprises Ban1 and Cubin. The other is subdivided into a branch where we find the other two *Anoplotermes* samples (Ban2 and Ban3), and another branch by the rest of the *Cubitermes* samples. Among them, at an inner position we find the *Cubitermes* samples originating in CAM. These differences can be clearly observed through a correlation heatmap (**Fig. 11B**), and the overall gene content can be observed at **Fig. S2**, for PFAM annotation.

**Fig. 11.**
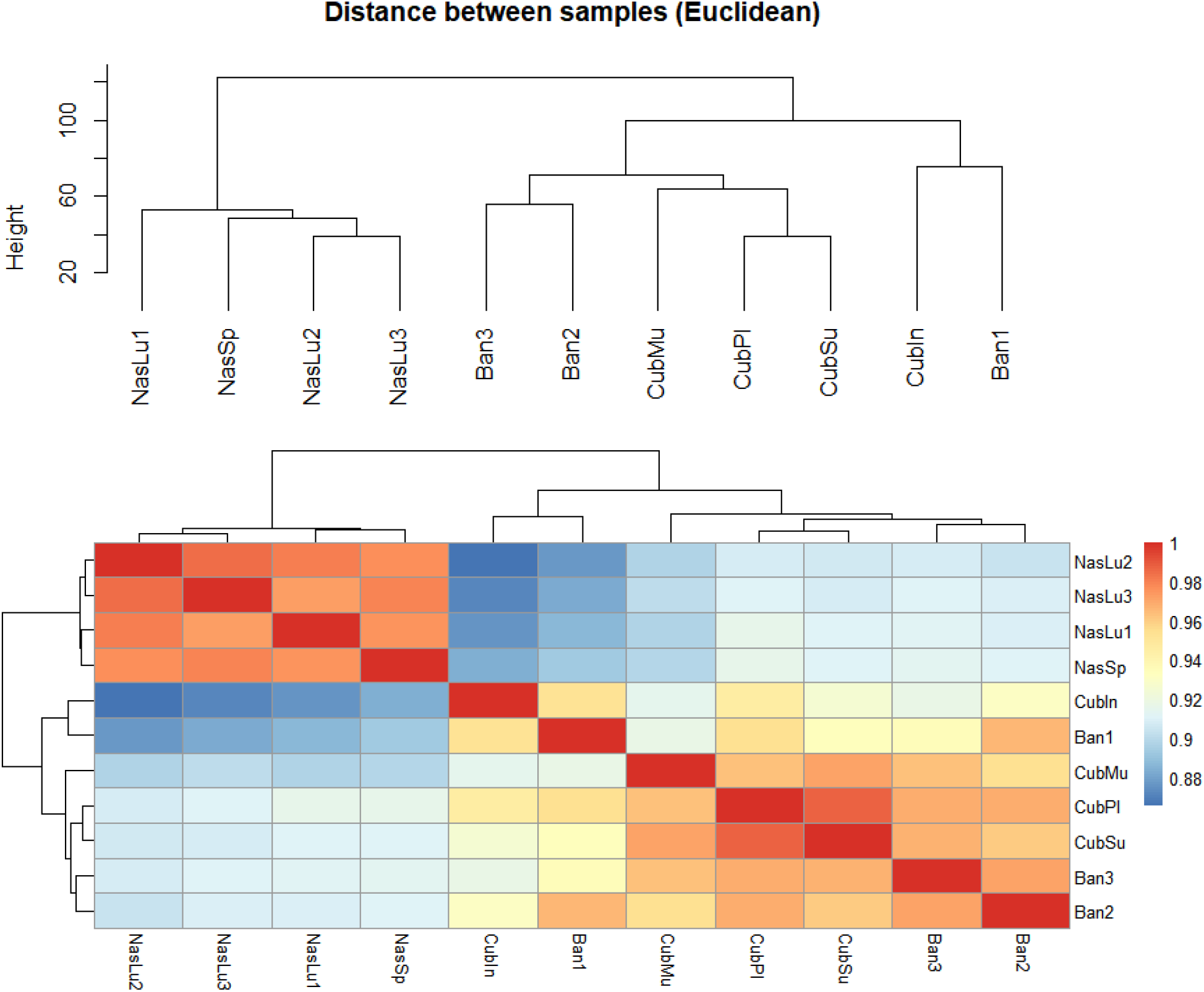
Hierarchical clustering of samples, A) Euclidean distance. B) Hierarchical clustering represented by means of a heatmap.

The main differences existing between pair of samples in terms of Differentially Present Genes (DPG) have been displayed through volcano plots (**Fig. 12**). When using COG annotation, at a fold change of 2 (|log2FoldChange| ≥ 1; DESeq2 padj ≤ 0.05) we found from a total of 2,099 DPGs, 1,147 upregulated genes (more present in *Cubitermes*) when comparing the two soil feeders (*Cubitermes* vs. *Anoplotermes*), and 952 COG annotated genes with higher presence in *A. banksii* samples (**Fig. 12A**). When comparing *Nasutitermes* and *Cubitermes*, we found 6,771 upregulated DPGs, and 7,871 downregulated ones (statistically significant higher presence in the soil feeder) (**Fig. 12B**). the last comparison was carried out between *Nasutitermes* and *Anoplotermes* (**Fig. 12C**). We found 7,152 upregulated genes, while there were 7,478 downregulated DPGs.

**Fig. 12.**
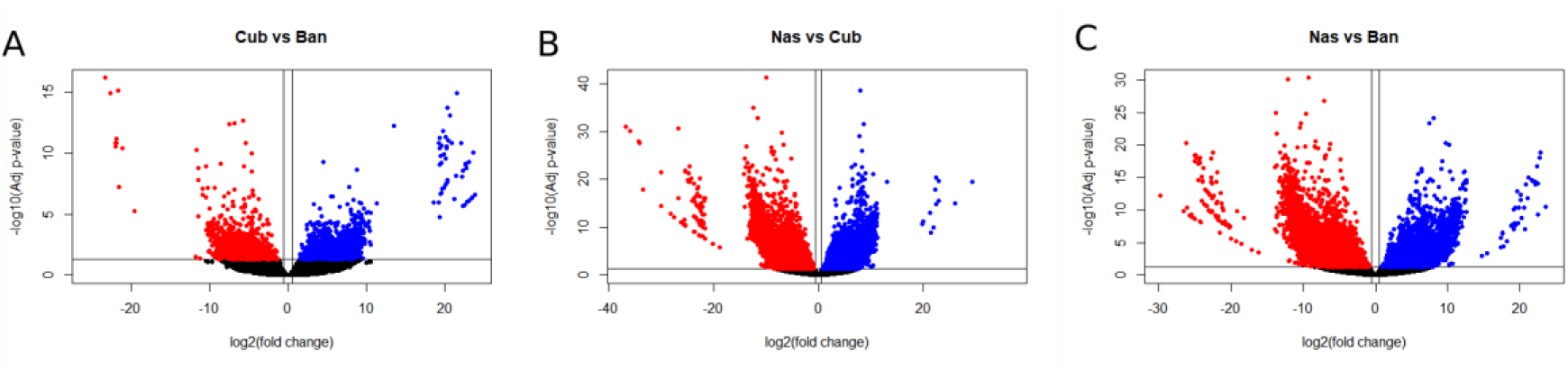
Volcano plot for the DPGs in the comparisons between microbial nests metagenomic samples, when using COG annotated genes as a reference. A) *Cubitermes* vs. *Anoplotermes*. B) *Nasutitermes* vs. *Cubitermes*. C) *Nasutitermes* vs *Anoplotermes*. Cub.: *Cubitermes;* Ban.: *Anoplotermes;* Nas.: *Nasutitermes*. Blue: genes with higher presence on the first component of the comparison (up-“regulated”); Red: genes with lower presence on the first component of the comparison (down-”regulated”).

We then carried out a similar analysis using the PFAM annotated genes (heatmap showing the overall in **Fig. S2**). For the comparison between the functional profiles of microbial communities present in the nests from both soil feeding genera, we found a total of 194 PFAM annotated contigs with statistically significant over-presence in *Cubitermes*, while *Anoplotermes* had 265 PFAM-annotated contigs with over-presence. We could equally state that these 294 genes have a decreased presence in the microbial communities from *Cubitermes* genus, in comparison with the microbial communities on those nests from *Anoplotermes*. We found some genes with extreme differences in presence, such as PF03945: delta endotoxin, N-terminal domain (FC = 16,326,654.29; p-adj = 1.55 10^-07^) and PF14470: Bacterial PH domain (FC =5,562,101.72; p-adj =1.34 10^-15^). On the other side, with higher presence in *Anoplotermes* PF02183: Homeobox associated leucine zipper (FC =3,509.92; p-adj =9.10^-14^).

The comparisons between microbial communities from the two soil feeding termites with the wood feeding termite nest provided again a higher number of DPGs, as previously observed with the COG annotation. The first comparison was carried out between *Nasutitermes* and *Cubitermes*. We found a total of 2,002 PFAM annotated contigs with statistically significant over-presence in *Nasutitermes*, while *Cubitermes* had 1,114 PFAM-annotated contigs with over-presence. We could equally state that these genes have a decreased presence in the microbial communities from *Nasutitermes* genera, in comparison with the microbial communities on those nests from *Cubitermes* genera. Among the functions most present in *Nasutitermes* we find PF16937: Type III secretion system translocator protein, HrpF (FC =4,569.92; p-adj =7.20 10^-13^), PF17667: Fungal protein kinase HrpF (FC =2,032.43; p-adj =2.26 10^-07^), and PF05420: Cellulose synthase operon protein C C-terminus (BCSC_C) (FC= 1,943.41; p-adj =1.01 10^-15^). The other direction, higher presence in *Cubitermes,* provides a number of highly present PFAM functions (Table S1), for instance PF03945: delta endotoxin, N-terminal domain (FC= 299,812,460.54, p-adj = 8.66 10^-14^).

The comparison between *Nasutitermes* and *Anoplotermes* yielded 2,034 upregulated genes, while 1,207 were downregulated.

#### 3.6.4 Annotation of ORFs in specialized databases

We carried out further annotation of the ORFs retrieved by SqueezeMeta with several specialized databases of interest (view Mat. Methods section) (Suppl. material S2 to S5). The conditions that had to be satisfied for being included in our list was a minimum identity ≥ 40%, bit-score ≥ 50. An arbitrary threshold of ≥ 90% identity (diamond blastx) was selected firstly for considering genes as known or already deposited in databases, while under this threshold genes were considered novel.

##### 3.6.4.1 Differential gene content: AromaDeg

We used the annotation obtained through diamond blastx (AromaDeg DB) as an input for differential gene content analysis. We firstly curated the results to eliminate those that were duplicated, where the total AromaDeg annotated genes counted for a total of 8,132. The comparison *Cubitermes* vs *Anoplotermes* yielded 14 upregulated DPGs, and 21 downregulated DPGs. The comparison *Nasutitermes* vs *Cubitermes* was carried out, and 468 genes were upregulated or significantly more present in *Nasutitermes* genus, while 82 were downregulated or with higher presence in *Cubitermes*. For the comparison *Nasutitermes* vs *Anoplotermes* we found 224 upregulated genes, and 12 downregulated ones.

##### 3.6.4.2 Differential gene content: CAZy

We carried out the analysis with the CAZy DB annotated genes. In the comparison between both soil feeders we found no statistical differences.

*Nasutitermes* vs *Cubitermes* yielded 287 genes upregulated, while 2 were downregulated. We found a larger number of genes for the *Nasutitermes* vs *Anoplotermes* comparison. This one provided 519 upregulated genes, while 218 were downregulated.

##### 3.6.4.3 Differential gene content: CARD

The comparison between the soil feeders yielded 54 upregulated genes, and 241 genes with higher presence in *Anoplotermes*. The correlation among samples and hierarchical clustering divided clearly all samples by genera. The comparison *Nasutitermes* vs *Cubitermes* resulted in 1,587 CARD annotated genes with higher presence in the wood feeder nest community, while the soil feeder nest community had 988 upregulated genes.

When *Nasutitermes* was compared with *Anoplotermes*, we found 1,403 upregulated genes, while 1,745 genes were downregulated (higher presence in the soil feeder).

## 4 Discussion

A very important aspect of termite ecological success is their ability to maintain a pathogen-free nest. This is of paramount importance since they live in colonies with very high density of individuals, where conditions are humid, nutrients are widespread, and temperatures can favour the growth of microbial strains. Among different strategies, fumigation with organic compounds such as naphthalene can aid (Chen et al., 1998b). We wanted to evaluate through an untargeted metabolomic approach the presence of volatile compounds, including naphthalene. We studied termite nests originating in three geographical locations, displaying different feeding strategies (**Table 1**). Some of the difficulties that we have faced came as a consequence of the taxonomic identification of samples. While we for instance thought that *Cubitermes* samples from MW were the same species, it was further confirmed to be three different ones. This fact creates an additional constraint since the samples are unique and sampling campaigns are costly.

We have then carried out a multi-omics approach with metabolite untargeted and targeted analysis, being the second omic level the taxonomic microbial profile, and lastly a metagenomic sequencing of selected samples with detected naphthalene presence.

One of the most important findings, is that the metabolite untargeted approach was able to stratify the samples according to feeding strategies, with a certain influence of the geographical location. Samples with one representative were eliminated from the analysis, especially due to bias from samples originating in artificial environments (breeds). The 10 most decisive analytes were several organic compounds, most of them of type alcohol. Naphthalene was also found when we consider the separation between genera (**Table 2B**).

One of these compounds (**Table 2**), 2-butanol, has resemblances with 2-methyl-butanol, previously reported to have antifungal activity (Matsuura and Matsunaga, 2015). We found it important for separation of samples according to feeding habits and genera.

Terpenes are a group of aromatic compounds that have been related to mediation of information transmission (Gershenzon and Dudareva, 2007), or have been tested specifically as a toxic compound against termites (Seo et al., 2014). Among them, terpinolene was found as a decisive analyte in our study. Terpenes have being reported repeatedly in termite literature as a chemical component of the defensive secretions in *Nasutitermes* soldiers (Prestwich et al., 1981), not related with feeding habits as explained by its absence in workers. Besides, terpenes have been appointed as an antimicrobial substance, as confirmed by Mitaka and collaborators (2017). Terpinolene was firstly identified in the cephalic secretion from *Amitermes herbertensis* (Moore, 1968), and later in small quantities of a Nasutitermitinae, *Grallatotermes africanus* (Prestwich, 1979), in the defensive secretion from *Nasutitermes ephratae* soldiers (Valterová et al., 1986), soldier frontal glands in *Termitogeton planus* (Dolejšová et al., 2014), and significant amounts in *Constrictotermes cyphergaster* soldiers (De Mello et al., 2021). According to our results, this compound is mostly found in the nests from *Nasutitermes,* with neglectable concentrations in the nests from the other genera. Considering literature, we believe that this compound is used as an antimicrobial defensive mechanism, and that the secretions may be carried out by soldiers, as extensively reported in several studies.

A PLS-DA analysis showed a better distinction according to the feeding habit and genera. Most of the soil feeding genera cluster together. Thus, we hypothesized that the geographical origin has a lower effect on the nest microbial communities than the feeding habits or the genera, and a great distinction among metabolite content in the nest material from our analysed termite species reflects the termite dietary habits.

At the targeted approach, our data indicated a lower level of naphthalene in comparison with published literature (Chen et al., 1998b; Wilcke et al., 2000). In overall, two genera concentrate the readings of naphthalene (*Nasutitermes* and *Cubitermes*), while the rest of samples had a scattered representation in some genera. We did not record naphthalene in the soil feeding nests from *Anoplotermes* from FG. We believe that the type of sample preparation could have caused partly these readings, although a soil standard indicated correctness in our method. In some cases, the results could have been affected by the time elapsed from sampling to processing, matter of a few days due to travelling and transportation. Even if that is the case, we could still observe clear differences between samples at the volatile level, which were further confirmed at the other omic levels.

### 4.2 Sequencing

#### 4.2.1 Bacteria

Our analyses included a negative extraction control (Eisenhofer et al., 2019). Bacteria are ubiquitous and even dust on a tube can produce a signal after PCR amplification, while many times there is as well an input of DNA in the reagents (Salter et al., 2014). We have deployed specific measurements for avoiding contamination inputs, for example carrying out the nest sampling under laminar-flow conditions, equally as those put in practice when performing the DNA isolation protocol. The reagents were treated as if they were sterile microbial growth media, putting in practice all aseptic methods that are normally used when working in microbiological experiments. While reagents are not sterile, we only used them for this project, and all bottles were exclusively opened under aseptic conditions and using sterile filtered tips. These measures are of paramount importance when working with samples with potentially small quantities (Weyrich et al., 2019), but as well for any other type of projects, a cross contamination can lead to incorrectly establish a core microbiome among samples. Despite these precautions, stringent conditions, and wide experience working in aseptic conditions by the sole operator, we have observed a number of OTUs in negative samples. After initial analysis of these data, and we confirm that the OTUs do not overlap with the ones in our samples, except for a really minor amount which would not affect the downstream results. However, we carried out a filtration step according to common procedures in Qiime2, which resulted in the elimination of virtually all contaminating tags. This elimination was effective for the negative control sample (some residual tags remained with zero overlap with the rest of samples), and for the rest of samples.

##### 4.2.1.1 Whole set of samples

We firstly analysed the microbial communities in the nests, with a quick overview on the bacterial community compositions through calculation of the Bray-Curtis dissimilarity metric (Sorensen, 1948; Bray and Curtis, 1957). This allowed us to obtain a quick classification of the studied colonies given the broad species origin. This index provides an idea of similarity/dissimilarity due to the product of present taxa and their relative abundances (van Rensburg et al., 2015). When studying the whole set of samples, we observed similar results than those obtained with the overall untargeted profile (Fig. S1), without a very clear stratification of microbial communities. The ACE index of α-diversity provides an overview of the microbial richness within one sample, but does not take in account the OTUs abundance (Qiao et al., 2017). We observed an effect of geographical origin in the richness of the microbial communities associated to different nests, where the samples with highest values were those from Breeds. This could be owed to an artificial input of different strains in several samples and colonies kept in similar laboratory conditions and close proximity. Besides, there is an input of human microbiome that could be at play. The second highest level was that of MW samples. We hypothesize that the savannah conditions could pose a higher degree of stress over soil microbial communities, which could find a better shelter in these nests.

In terms of feeding strategies, wood feeders showed statistical differences with soil feeders from group III. While this could be owed to the different type of diet, these differences were as well present between groups III (with origin in FG) and IV (only African representatives).

Genera was a factor with more differences among microbial communities, where we found differences mainly in those ones with at least three representatives. Among wood or soil feeders, we observed only ACE index differences between *Nasutitermes* and *Neocapritermes*. Kruskal-Wallis comparisons among all groups indicated differences when considering feeding groups and geography.

At β-diversity the weighted UniFrac distance can represent the differences between samples according to the evolutionary history (Lozupone and Knight, 2005; Lozupone et al., 2007, 2011), and how different are two microbial communities. Our results show significant statistical differences between feeding groups III and IV, and between FG and MW. That could be surprising considering that groups III and IV are both soil feeders, but can be explained because all of the samples from group III originate in FG, while the ones from group IV originate in Africa, with a great number of them from MW. These two locations are precisely the ones yielding differences when considering geography, and agree with the reported statistical differences among all groups observed at the α-diversity, which were observed on feeding strategies and geographical origin. For β-diversity, we could not compute distances based on genera, as some groups had only one representative.

##### 4.2.1.2 Samples with at least three representatives per genus

Unfortunately, we could not increase the number of samples from other genera than *Anoplotermes, Cubitermes, Nasutitermes*, and *Neocapritermes*., since we largely rely on bringing nests from distant and remote geographical locations, and not all times they survive transportation. Soil feeding species are especially sensitive. A long-time observation in our laboratory, is the fascinating fact that nests without the presence of termites are quickly populated by pathogenic fungi. At the moment we ignore if it is infection what drives the death of these colonies, or these species are especially sensitive to the artificial laboratory conditions. What is clear is that in matter of scarcely two weeks fungi dominates the colony and impairs sample processing or the use of the material. This is another factor that limits our potential for bringing a higher number of colonies to our home laboratory.

Their stratification in a PCOA plot using Bray Curtis values (plot not shown), indicate an important component of variability in bacterial microbial populations explained by the type of feeding habits, as most of the samples are clustering in proximal areas of the representation, with only one exception from one sample belonging to group II within group IV. Additionally, we got the idea that the genera can be another important factor in the explanation of variability. The potential lack of influence of the geographical location is indicated by the presence of *Cubitermes* samples in the same sector, despite originating in Malawi and Cameroon (with known differences between tropical rainforest and savannah). However, we cannot discard it completely, since some samples at the feeding group III (*Neocapritermes*) are closer to a representative of group II from FG. These results are indicative that feeding group and genus have a stronger role in stratification than geographical location. We cannot ascertain that naphthalene presence could be driving the stratification.

Among other biodiversity calculations we obtained the Faith Phylogenetic Diversity (Faith’s PD) index (Faith and Baker, 2006), indicating that among geographical locations exists a statistical difference between those samples from Malawi and those in French Guiana, as reported as well for the ACE index. This indicates that differences in diversity at the nest bacterial microbial communities have, until some extent, influences due to the different conditions at geographical locations. The lack of differences between the rest of groups, for instance CM and MW, could point towards a conservation of the diversity in Africa, despite the marked ecological differences between the two sites. These differences across the ocean are of higher entity, accounting for the statistically significant differences between FG (tropical rainforest) and MW (savannah), although not enough when comparing the two tropical sites at both sides of the Atlantic.

We found as well statistical differences in diversity at the genera level, accounting for differences between *Cubitermes* (MW-CM, SF) and *Neocapritermes* (FG, SF), and at the other hand between *Nasutitermes* (CM, WF) vs *Neocapritermes* (FG, SF), and among *Anoplotermes* (FG) and *Neocapritermes* (FG). Here we can observe as well that geography is not a strong factor determining richness differences (comparison between *Anoplotermes* and *Neocapritermes*), but genera seems to take a greater role. Furthermore, when considering feeding habits we must remember that *Cubitermes* and *Neocapritermes* are in two distinct feeding groups (Group IV and Group III respectively), were we also report significant statistical differences. We could point towards the feeding strategy as another difference to account for the bacterial diversity. In terms of β-diversity results are mainly in agreement with the α-diversity, except for the differences among genera.

The PCA plot based on the annotated OTUs further confirmed that the differences between microbial communities are determined strongly by the genera and then the feeding strategy (**Fig. 6A**). While *Anoplotermes* and *Neocapritermes* belong to the same feeding group, they occupy distant locations in the plot, what provides the idea that genus is a stronger component than feeding strategy. The second part of the plot (**Fig. 6B**) indicates that the geographical location is weak when explaining the variability among nest bacterial communities based on 16S amplicon data.

The results from our study agree with the published termite literature regarding the bacterial taxonomic composition of nests. *Actinomycetales* were reported as dominant in termite nests (Krishanti et al., 2018). These are Gram-positive bacteria capable of forming branching hyphae, and related to nutrient recycling roles (Goodfellow and Williams, 2003), and have been used for production of secondary metabolites, accounting for a very large fraction of all bioactive secondary metabolites in use at industry (Olano et al., 2008). This group has been also reported in the study from Chouvenc et al. (2018), where authors studied the origins of acquisition of *Streptomyces*, the largest genus within *Actinobacteria* (El-Naggar, 2021), in *Coptotermes* nests. Authors have reported a recruitment from surrounding soil. Similar results regarding *Streptomyces* were reported in nests of *Reticulitermes flavipes* (Aguero et al., 2021). Another group of bacteria notably described in literature is that of *Rhizobiales*, Gram-negative bacteria belonging to the *Alphaproteobacterial* class, and with many members capable of fixing nitrogen (Beeckmans and Xie, 2015) and associated with many plants, lichens, and mosses (Erlacher et al., 2015). This order was also reported as enriched in termite nests of *Coptotermes testaceus, Heterotermes tenuis,* and *Nasutitermes octopilis* (Soukup et al., 2021). In our case the proportion is limited.

*Clostridiales* are gram-positive anaerobic bacteria (Bowman, 2011) that have capability to form resistant spores (Paredes-Sabja et al., 2011), which have been described as thermophilic ethanologens capable of fermenting lignocellulosic biomass (Arora et al., 2019). This bacterial order includes species that are lignocellulolytic (Auer et al., 2017). This order has been associated with the termite gut community of soil feeders (Ohkuma and Brune, 2010), and also *Reticulitermes santonensis,* through the presence of *Clostridia* related clones within *Firmicutes* (Yang et al., 2005), and was appointed as one of the groups carrying out recycling of uric acid (Thong-On et al., 2012). In *Odontotermes yunnanensis* Long et al. (2010) report the presence of a wide representation of bacteria from *Clostridium* genus in fungus comb structures. We found that this group was of the highest proportion in two of the soil feeders (*Anoplotermes* and *Cubitermes*), but also in nests from *Nasutitermes*. Regarding *Acidimicrobiales*, we have found no specific information in termite literature.

*Solirubrobacterales*, an order with few species described as Gram-positive mesophilic bacteria growing in a moderate range of temperature (Whitman and Suzuki, 2015). Our results are in agreement with those described by Enagbonma et al. (2020), where they report this group at termite mounds from *Coptotermes* species. Uncultured representatives of *Solirubrobacterales* have been associated to the degradation of lignin in soils (Wilhelm et al., 2018; Silva et al., 2021), previously reported as well in sugarcane farm soils (Ogola et al., 2021).

*Spirochaetales* are Gram-negative that can comprise both anaerobic and strict aerobic bacteria (Smibert and Johnson, 1973). They have been reported in increased numbers when manure is applied to soil (Rieke et al., 2018). This group was also reported in *R. santonensis* gut (Yang et al., 2005) and gust from *Coptotermes gestroi* (Do et al., 2014), and have been appointed as active cellulose degraders (Xia et al., 2014), with abundance in wood-feeding termites (Tokuda et al., 2018; Hu et al., 2019).

*Gemmatales* are Gram-negative chemoheterotrophic bacteria (Dedysh et al., 2020), which could act as an indicator of elevated phosphorus content in soil (Mason et al., 2021), and in a complex microbial community dedicated to the utilization of cellulose could be providing endocellulases and β-glucosidases activities (McDonald et al., 2019).

The characterization of samples through biomarkers (LEfSe analysis, **Fig. 5**) indicated that certain genera were representative of specific samples when compared together. *Streptomyces* was associated with *Nasutitermes,* an observation that is in agreement with the study of Chouvenc (2018), and Aguero (2021), and could be indicative of a defensive role of this bacterial order in the termite nest.

*Neocapritermes* with Beta-proteobacteria, which have been described as characteristic cellulolytic bacteria in acidic forest soils in temperate areas (Štursová et al., 2012), and belonging to this class the genera *Burkholderia* and *Collimonas* have been characterized as capable of carrying out mineral-wheathering (Lepleux et al., 2012). Nests microbial communities from *Cubitermes* were associated through LDA with the genus *Rhodococcus,* comprising aerobic Gram-positive bacteria. Members from this genus have been associated with degradation of nitroaromatic compounds (Subashchandrabose et al., 2018), but more importantly with lignin degradation activities (Chong et al., 2018). This observation is aligned with the fact that lignin degradation is not important in wood feeders (Ohkuma, 2003; Griffiths et al., 2013), and could be an indication that members of this genus carry this important task in the *Cubitermes-* associated communities. and Intrasporangiaceae.

*Chitinophaga* has been described as a chitinolytic and lignolytic genus (Funnicelli et al., 2021), besides *Spirochaeta*, where one of the species was reported to degrade complex plant polysaccharides and depolymerize lignin (Pandit et al., 2016). That observation agrees with the type of diet in *Anoplotermes,* which will find lignin enriched sources in the soil, and by construction of the nest this type of bacteria can be enriched. Additionally, we could speculate that this bacteria is in charge of eliminating the growth of hyphae from fungi due to its ability to grow on fungal material, as shown by McKee and collaborators (McKee et al., 2019) when studying different roles of *Chitinophaga pinensis*. Perhaps, the role of this genus in Anoplotermes nest could be related to recycling of the dead material from fungi.

#### 4.2.2 Amplicon sequencing: fungi

##### 4.2.2.1 Whole set of samples

PCoA Emperor plots based on Bray Curtis diversity metric explains lower variability than in the case of 16S data. In overall, a few explanations emerge from this representation. We observe a grouping of samples according to feeding strategy, and geographical location, since most of the samples from each sampling site cluster in the proximity. There is still a clustering of samples according to genera, for *Anoplotermes*, *Neocapritermes*, *Nasutitermes*, while *Cubitermes* are divided by geographical location.

ACE index calculations informed of a lack of differences in terms of OTU richness when we compared samples based on their geographical origin. Differences existed at the genus and feeding strategy levels. At this occasion, the feeding groups where we found differences were II vs III or IV, but also III vs IV. This indicates that the feeding strategy is capable of driving a differential richness in the fungal communities present in the termite nest. Furthermore, this seems to be reflected as well for the differences between genera, since *Anoplotermes*, *Cubitermes*, and *Neocapritermes* have each one differences towards *Nasutitermes* in ACE metrics.

In terms of β-diversity, we found similar differences at statistical level as those observed in ACE metrics (same categories in feeding groups and genera), while in this case there were as well statistical differences in the pairwise comparisons at geographical level in CM-FG, and FG-MW. These results could indicate that the fungal communities are more susceptible of the geographical location, especially for the found differences between CM or MW vs FG at both sides of the Atlantic. The observed differences seem to be stronger in overall for fungi than for bacteria, and in this case driven both by the feeding strategies, especially wood vs soil feeding, and geographical origin (S. America vs Africa).

##### 4.2.2.2 Samples with at least three representatives per genus

We also proceeded to eliminate all those nest microbial communities with less than three representatives per genus, and as it happened with the 16S data, we got very clear results.

The PCoA representation provides clearer results than those for 16S, although with less explanation of variability. The factors that drive variability and stratification of samples are mainly feeding strategy, but especially genera. The representation of the latter stratifies the samples in four sectors. This is an indication that the fungal profile has a marked accent in the differentiation of fungal microbial communities.

For the ACE metrics differences were reduced, while those in terms of genera remained. The β-diversity showed again these differences between group II and III or IV, between III and IV. We have here the case where the richness seems to be less affected by the feeding groups, but these strategies are driving differences in the composition of the communities, or how different are one from each other.

Richness was affected by the genera, somewhat affected by the termite diet. The β-diversity was as well patent among the different genera. Because we also observed statistical significance of the location, we can conclude that differences are a complex product of the genera, feeding strategy, and ultimately also geographical origin with certain differences at both sides of the Atlantic, but not within Africa.

The PCA plot built in R (**Fig. 6**) was indicative of the stronger effect of genera and feeding strategy. We believe that genera were even stronger in sample stratification than type of diet, because even two genera in the same feeding group are clearly separated (*Anoplotermes* and *Neocapritermes*). Lastly, while existing, the geographical influence was weaker than the other two. These results are in overall agreement with our observations for the bacterial communities.

At the observed taxon profiles, we found that nests are dominated by the presence of OTUs belonging to the phylum *Ascomycota*. Within this phylum, the class *Sordariomycetes* and *Dothideomycetes* are the most abundant, as reported for nest walls from *Cornitermes cumulans* (Menezes et al., 2018). From the phylum *Basidiomycota* we found the appearance of two categories, the classes *Exobasidiomycetes* and *Agaricomycetes*. In keeping with our observations, these two phyla were the most frequent and represented in fungal isolates from *O. formosanus* (Xu et al., 2020). Similar results have been also reported by other recent studies (Větrovský et al., 2020)

Two phyla that belong to the kingdom Plantae appear as well in the first positions, but with distinct distribution among samples, down to the orders Chlorophyta and Magnoliophyta. We can explain the appearance of these organisms belonging to a different kingdom due to the reference used to annotate the ITS through QIIME2, in this case the Silva database which also contains Eukaryotic entries. Algae and other eukaryotes that contain 18S rRNA gene can be detected through amplification (Nakai et al., 2012; Banerji et al., 2018) even being used for classification (Buchheim et al., 2012; Mikhailyuk et al., 2020). We decided not to filtrate it as it could provide important information for our study.

There are some differences between samples belonging to the same genus but at different locations. According to what we described above for the different diversity metrics, it seems that geography is also a factor to have in account when considering differences between nest fungal microbial communities.

#### 4.2.3 Metagenomic sequencing

There was a further selection on the samples used at metagenomic sequencing, as reported in the results section. For the case of *Anoplotermes*, all three samples belong to the same species: *Anoplotermes banksii*. For *Nasutitermes*, we had three representatives of *Nasutitermes lujae*, and one from *Nasutitermes* sp. Lastly, for *Cubitermes* we had representatives of four different species (**Table 1**). While it could be argued that we should have included only representatives of the same species on each sequenced group, that is very difficult when we talk about termites. There are constraints due to the inherent issues bringing the material in adequate conditions from the tropics to our home laboratory. Not every colony is able to arrive in good conditions, and when termites die, the nest is quickly invaded by pathogenic fungi. If we take in account the suboptimal conditions that we face in the field, it is clear that in-situ processing is not feasible. We hope that in future sequencing portable devices currently existing can be applied in our field, as it could help enormously in termite research. One aspect where it can help researchers, is the identification of colonies. The case of *Cubitermes* reflects those emerging issues with clarity. While all colonies and termites look the same at the bare or even taxonomically trained eye, it is very difficult to make accurate decisions. That is even an arduous task for termite taxonomists with good equipment in the laboratory, and it is not always possible to have these trained scientists on the field. Even at that level, our originally thought 4 equal representatives of *Cubitermes*, turned out to be different species. That was until certain extent happening with the genus *Nasutitermes*.

Additional limitations that drove our selection was the computational power, since the predicted amount of data and information was vast, and later on it turned out to be of a very notable magnitude. It also presents an enormous potential for discussion and description of novel genes of great interest for different biotechnological purposes. Computational restrictions have to be also taken in account, and while we could carry out the amplicon sequencing bioinformatic analyses without issues, the whole metagenomic data treatment and analysis required significant resources and computing time. This process delayed notably our pace especially when any step had to be repeated in order to introduce modifications that were not possible to foresee. We found that the SqueezeMeta pipeline leverages the process, and produces a vast amount of information that could be interpreted through SQM tools (Puente-Sánchez et al., 2020). Among other data management, SQM tools allow for sample sub-setting.

We have used different ways of presenting this data due to the massive amount of information that could be extracted from it. In general, we observe an equal trend for samples to stratify or group according to the feeding group and genera, with limited influence from the geographical location (**Fig. 7**). Despite small variability between nests from species belonging to the same genus, the differences are surprisingly stable, and even communities from soil feeding termite nests are having closer genomic and taxonomic composition at both sides of the Atlantic, as revealed at the NMDS analysis, which takes in account the full metagenome of each sample.

At the phylum level (**Fig. 8**) we still observe a comparable situation, where all *Nasutitermes* have a closer profile between themselves than those of *Cubitermes* or *Anoplotermes*. There are yet differences between samples within the same genera (with the exception of *Nasutitermes*). CubPl and CubSu both originating in CM have a closer profile than the other two from MW, while CubMu does still resemble the previous ones with incorporation of some OTUs not apparently present in CM (*Chloroflexi*). One of the samples from *Anoplotermes banksii* shows large differences with the other two samples from this species, probably owed to a worst state of that sample when arriving to our laboratory. The three samples from *Nasutitermes lujae* are very stable in terms of taxonomic representation. We can conclude that the taxonomic composition of the nests reflects the genera (where it seems to be stable), and especially eating habits.

In agreement with our previous taxonomic metabarcoding through 16S amplicon sequencing and ITS sequencing, we observe here similar trends. For instance, among fungi presence of *Ascomycota*, which was the most represented phylum, but in this case seems to have higher representation in wood feeders. For bacterial annotations, we find as well a very notable presence of *Actinobacteria* as reported in the first omic level. However, for 16S amplicon data we did not find that relevant proportion of *Proteobacteria* as that obtained by metagenomic sequencing. We found that the most abundant order in many of the sequenced soil feeders was *Streptosporangiales*, while in Nasutitermes they belong to class *Actinomycetia*, either to genera *Gryllotalpicola* in three cases or *Mycobacterium* in the last one.

But metagenomics has greater capabilities than just being able to provide another overview of taxonomical structure of the community. The part with most interest is the possibility to evaluate the functional potential, the metagenome.

Given a certain function, we could observe if at that level exists a differential composition in terms of taxa. We carried out a subset of functions (Puente-Sánchez et al., 2020) related to “Phenylalanine, tyrosine and tryptophan biosynthesis”. These three aromatic α-amino acids are involved in the metabolism of secondary metabolites (Parthasarathy et al., 2018). We found apparent differences according to the feeding strategies (**Fig. 9**). Once more, both soil feeding genera resemble each other, while the *Nasutitermes* keep a closer profile among them. This indicates that the differences are not only at the metabolomic level, or at the overall taxonomic structure of the community (both amplicon and metagenomic sequencing data). The differences go beyond and certain functions, as the one exemplified here, are carried out by different microorganisms. While nests from soil feeding termites display similar profile at both sides of the Atlantic, wood vs soil feeding are more dissimilar. Considering that, the geographical component seems to have a weaker contribution than the feeding habits in driving the functional differences among communities. There seems to be though a small reflection of a component that relates communities from the African environment, as we observe a group of unclassified *Actinobacteria* with similar proportion between *Nasutitermes* and *Cubitermes*. Our explanation here is that the feeding habits of termites and possibly some termite-active antimicrobial strategies are creating a set of conditions that allow the growth of certain type of genera, which will carry the required set of community functions, in this case the metabolism of aromatic amino-acids. We mentioned the termite active strategies, due to the presence of terpene as a selective metabolite in our study, with antimicrobial properties and secreted by *Nasutitermes*, as both evidenced in literature. The defensive or protective component of faeces was greatly reviewed by Cole, Ceja-Navarro, and Mikaelyan (2021). In our study, we infer that the combination of active strategies and passive constraints imposed by the feeding strategy create a specific growth-media for selected microorganisms. We could establish a parallelism with microbiology work in the laboratory, and state that termites with different feeding habits have formulated a growth media for their allied microbiome to thrive. A number of iterations of this process through trial and error in evolutionary time have probably fine-tuned these strategies, where the most successful in ensuring the nest and colony survival have been fixed. Until such an extent that microbial communities associated to two different genera of soil feeders resemble in structure at two sides of the Atlantic Ocean, where the different flora composition impose a different soil community (Lima-Perim et al., 2016). However, despite these stated differences even at short distance, the differences between the *Cubitermes* from CM and MW are minor, even when belonging to two clearly distinct biotopes. Thus, this is one more observation that helps us to associate a lower discrimination power to the geographical location.

Among the communities with higher presence for this function in *Nasutitermes*, the genus *Gryllotalpicola* has been found in larval galleries from the pine beetle *Monochamus alternatus* collected in the Chinese Jiangxi province (Chen et al., 2020), and was previously isolated from guts of the wood-feeding *Reticulitermes chinensis* Snyder, sampled in Wuhan, China (Fang et al., 2015). The interesting connection is that members from this genus have been found in insects with wood-feeding habits, at distant locations.

*Streptosporangiales* is an order of bacteria that can obtain carbon from green waste (Cai et al., 2018), especially at the middle composting stage (Li et al., 2021). They can secrete cellulases and hemicellulases for lignocellulosic degradation, besides of being producers of antibiotics that can supress pathogens (Gomes et al., 2017; Waglechner et al., 2019). These organisms are highly abundant in soil (McCarthy and Williams, 1992). Thus, since soil feeding genera use soil as their substrate, *Streptosporangiales* may originate at their surrounding environment and be enriched due to the substrate type and nutrients present. They could provide a cleaner environment at the termite nest due to production of antimicrobial compounds. This strategy, which we cannot ascertain if it is an active strategy or derived from the termite activity, repeats itself at both sides of the Atlantic Ocean.

The genus *Microlunatus* (*Actinobacteria*) has been previously reported in *Mycocepurus smithii* ants (Kellner et al., 2015), a fungus-farming ant reported to acquire its microbiome from the surrounding environment. Bacteria from this genus have been found from deep-sea sediments (Jroundi et al., 2020) to rhizospheric soil (Wang et al., 2008), and it has been indicated that they have potential for pollution management (Zhang et al., 2017), especially for capabilities of phosphorus accumulation (Zhong et al., 2018).

*Microbacteriaceae* is a family of Gram-positive bacteria with members that have been reported among the dominant cellulase producing strains isolated from yellow stem borers (*Scirpophaga incertulas*) (Bashir et al., 2013), but much less proportion in the guts from termites at the same study identified as *Odontotermes hiananensis,* which were feeding on decomposed trees and leaf litter.

The LEFSE analysis on the ORFs annotated at PFAM, and have found that each of the genera is having a specific signature.

But because this information is vast, we carried out a different approach for gene characterization. Are these apparent differences in specific functions translated into differential content of certain genes beyond a LEFSE analysis? Has the microbial community gained or being enriched in certain functions?

In transcriptomics, statistical testing is applied for finding differentially expressed genes (DGEs), what provides a clear idea of which genes are used in given conditions, being a molecular reflection of the phenotype. If this is applied in the case of metagenomics, the reflection is not exactly about what genes are being used, but more of the potential. However, this is not as static as if we were to evaluate the genome of a single organism. Being true that not all present genes are going to be expressed, we can infer that a certain function elevating its content in comparison with the same function in a second community, is relevant and this is not product of a random event. For instance, in the field of environmental antibiotic resistance qPCRs are routinely used to quantify the differences in gene content due to pharmaceutical pollution among different sites (González-Plaza et al., 2019; Milaković et al., 2019). We can then bring this knowledge to our current case and adopt the term “Differentially Present Genes” (DPGs).

While TPM is a normalization generally used in RNAseq experiments and its data treatment, according to Puente-Sánchez et al. (2020), it can be very useful in metagenomic experiments. In that regard, we found it as a very important value that allows for sample comparison and allowed us to get the first sample overview (**Fig. 7**). However, because we understand the limitations on TPM use (Zhao et al., 2020), we have carried out additional normalizations for comparison purposes, using procedures that use raw count values and are normalized differently, methodology taken from transcriptomics research (González Plaza et al. 2022, in preparation).

Testing for significant statistical differences will illustrate if a certain function is highly abundant in a nest microbial community, when compared with a second one. When carrying out these analyses, we can firstly establish a comparison between all samples based on the normalized count values for the PFAM annotated ORFs, which clearly separates *Nasutitermes* from the soil feeders (**Fig. 11**), again a division of microbial nest communities by the termite diet. We used for this analysis that reduced list of ORFs due to limitations in computing power. We additionally suffered the lack of a mixed approach between shotgun sequencing and long-read sequencing that could have notably improved the process and decreased the number of final contigs from where the ORFs where drawn.

When the analysis was brought to the DPGs content, we found that the differences between microbial communities in wood feeding termites nests and those from soil feeding termite nests are much stronger, given the amount of up-regulated (in terms of presence) genes, than those existing differences between the two soil feeding genera. Up-regulated in this study does not mean “higher expression” since we are working with DNA material, but it means a higher presence on that community.

The previously addressed recruitment of external or surrounding strains reported in literature, can be different in terms of specific strains, but they will be fundamentally equivalent in the functions that they carry out. Soil feeders “will recruit” microorganisms that are performing or carrying the same genes. We want to highlight that “recruit” may not be an active termite program, but a consequence of the complex cross-talk between active antimicrobial strategies, and the effect of diet.

Besides, the differences due to species appear to be of a lower entity, due to the relatively lower number of DPGs between both soil feeding genera. Among both soil feeder there were more than 2,000 DPGs, but those numbers rose to more than 14,000 DPGs when *Nasutitermes* was compared with either *Anoplotermes* banksii, or the *Cubitermes* genus.

We have carried out differential gene content analysis with the results obtained in specialized databases. We believe that this is important as a source of novel genes that could have a really important application in the biotechnological industry. Furthermore, our analyses have shown further a similarity among the genomic profiles between soil feeders and great differences to the wood feeder. Furthermore, the rationale for this additional genetic prospecting is that these databases can reflect different aspects of the microbial communities, which can be manifold and related with the metabolism as a cornerstone (Schmidt et al., 2019). In our study not only genes annotated in databases such as AnHyDeg or AromaDeg could serve as protection purposes when organic antimicrobial compounds such as terpenes are used, but also as part of a metabolism that could drive to social microbial interactions. For instance, antibiotic resistance (CARD database) can be used among members of the community to defend themselves against antibiotics, either from those produced by them, or from competing organisms (Scherlach and Hertweck, 2020). Additionally, at the usual amounts of antibiotic concentrations found at natural communities, these genes can serve for communication purposes rather than survival (Romero et al., 2011).

A first characterization of these results was to count for those known genes over a 90% identity threshold. We used a blasting approach with a lower identity threshold down to 40% but bit-score ≥ 50, based on Pearson (Pearson, 2013) to discover homologous genes. However, this threshold does not appear to be an adequate filtering value. Cases with 100% identity, had a length of alignment under 50 aminoacids. When using the maximum bit-score values, e-value was zero in 28 hits at the BacMet list, with alignment lengths varying from 605 – 1,118 amino-acids for those 28 genes, and identity ranging from 55% to 99%. Therefore, we suggest that the use of bit-score is a much better indication of the homology to described genes, rather than just using the percentage of identity.

We found that mostly known genes belong to the CAZy database, probably owed to the great interest of the cellulose and lignocellulose degrading field, where many sequencing approaches have been directed towards mining the microbial communities associated to insects that use cellulose. We believe that the other tools are also important, since for example the antibiotic resistance genes harboured in wildlife associated microbial communities has not been properly addressed, in comparison with the development of the field regarding man-made environments (Dolejska and Literak, 2019). Furthermore, the low number of described genes at BactiBase or AnhyDeg indicates the potential of these termite nest microbial communities for being a source of novel genes that can be used in fighting antimicrobial resistance (bacteriocins). The results from AnhyDeg mining can relate as well to a lower use of anaerobic degradation of hydrocarbons, since these nests environments should be mainly oxygenic. However, we carried out a thorough homogenization of the nest material, and that could have brought as well microorganisms thriving in deeper layers of biofilm structures or within the nest material.

When using three of these lists, we still found differences in gene content, again sharply evident between wood and soil feeders. The differences in the use of CAZy annotated genes were of small volume, and interestingly those were absent between both soil feeders. However, we must note that this could be a resemblance of the smaller number of genes that we used at this analysis due to computing limitations. If proportions are kept in regard to what we observe in the other comparisons, always higher number of DPGs between *Nasutitermes* and each of the soil feeders, it is logical to think that we did not have enough number of “events” for statistical differences to appear. Thus, while each of the lists referring to different type of microbial functional features, they maintain the observed differences when using the general PFAM annotated profile.

## 5 Conclusion

In our study we have evaluated a wide range of termite nests originating at the two sides of Atlantic Ocean, including nests belonging to species including soil and wood feeders. Sampling locations encompassed tropical rainforest and savannah habitats. We have faced constraints in our study that have limited the resolution, as some samples were unique representatives. Therefore, we have first characterized them, but for the main conclusions have been discarded. These limitations were imposed by the type of material and its source. We believe that further studies with higher number of samples per genus will have equal statistical power and reach similar findings.

We firstly found differences in the overall metabolite profile that indicated a strong effect of the feeding habits of termites inhabiting those nests than the geographical location. Some of the most selective metabolites at VIP analysis were found to be reported in literature from cephalic glands in *Nasutitermes* wood feeders. These compounds may have an antimicrobial role in those nests. When we evaluated naphthalene content through a targeted approach, we were surprised to find much lower content than those previously reported in literature. The naphthalene presence seems to be prominent in few of the tested microbial communities, especially those related to *Cubitermes* and *Nasutitermes* genera.

The evaluation of the bacterial and fungal taxonomical profiles from those microbial communities had no indication of any effect of naphthalene on them, but rather pointed towards a stratification owed to the termite feeding habits. This is probably owed to the fact that nest material is composed of termite faeces and surrounding soil material. We could not assess differences due to the geographical location. While the differences between both tropical rainforest soils could be limited, the microbial communities’ differences between the savannah and tropical soil should be of a very strong nature. However, even the differences between the species in nests from soil feeding termites from both distinct locations in Africa were minor, probably owed to the effect of genus and again the diet.

We carried out a metagenomic profiling of two nest containing naphthalene from two distinct genera (soil vs wood feeding) and an additional soil feeding termite nest from French Guiana with no detected amount of naphthalene. The results from our analyses support the previous observations, where the major drive for genetic functional differences are the feeding habits of those termites inhabiting the studied nests. We could still establish genetic differences between both soil feeding termites, but these were of a more limited entity than those between each of the soil feeding communities and the wood feeding one. It is to be highlighted the vast genetic richness of functions and genes of biotechnological application, as observed by our mining approaches with specialized databases. This is only one example of the virtually unlimited genetic diversity that tropical rainforests harbour, and should bring attention on the necessity of preserving this resource that could be applied in fields such as bioremediation of organic compounds, or the improvement in the synthesis of biofuel or bioethanol.

Gene content analyses have provided an overview of the apparent changes between distinct feeding habits and even allow us to indicate that the feeding habit is a stronger component into the gene diversity and overall functions, rather than the geographical location. In that regard, the results at both sides of the Atlantic for soil feeding strategies (reflected in the microbial composition of the termite) indicate that the local surrounding microbial communities have less effect, although we in this study ignore their composition in comparison with the communities studied here. We found certain differences between the *Cubitermes* representatives at species level, although still clustering together. Even when half of the representatives originate from a humid tropical location, while the Malawi representatives belong to a savannah ecosystem. Those are closer to *Anoplotermes*, despite the geographical distance. This is happening both at the gene content level, but also at the taxonomical composition level. The taxonomy composition arises from metagenomic sequencing, but also from 16S and ITS amplicon sequencing. We found clearer associations between the feeding habits and bacterial microbial communities than fungal communities.

We conclude that the differences among microbial communities at termite nests depend largely on the type of diet of the termite species, and not on the surrounding environment. This observation was already presented by Hu and collaborators (Hu et al., 2019), as authors suggested the linkage between diet and microbiome in termite guts. This observation is widespread with other animal models (Colman et al., 2012; David et al., 2013; Baker et al., 2014; Flint et al., 2015; Otani et al., 2019; Leite-Mondin et al., 2021). However, we state here that these differences go beyond the gut to be even reflected at the nest community level, consequence of the type of building strategy. The nest community structure-feeding habits linkage is here as thoroughly stated through three omics approaches.

A wide range of published studies have indicated that termites recruiting microbial members from the environmental communities. Our combined results are not contradicting that, but indicate that the genetic functions that microbial communities belonging to soil feeding genera are carrying out are similar across the Atlantic, even in two different habitats, savannah and rainforest. While exists a recruitment of bacterial strains from the environment, what ultimately fixes the functional genomic content is the type of diet, and subsequently the nutrients that are available on each case. That observation has been repeated in our study at the three omic approaches.

Further studies considering the different microbial niches (gut, environment, food substrate, and nest) could shed more light on the relationship between the microbial communities under influence of termites and those in the surrounding environment.

## 6 Acknowledgements

This project has been funded by grants IGA No. B_19_04 from the Faculty of Forestry and Wood Sciences (FFWS, Czech University of Life Sciences, Prague), and “Advanced research supporting the forestry and wood-processing sector’s adaptation to global change and the 4^th^ industrial revolution,” OP RDE, Ministry of Education Youth and Sports of Czechia, Grant No. CZ.02.1.01/0.0/0.0/16_019/0000803. JJGP thanks Dr. María Suárez at Wageningen University (The Netherlands) for support with scripts, to SurfBio Project (EU Horizon 2020 grant agreement No. 952379) for the training material provided at the Online Training School, to the EMBO Practical Course Microbial “Metagenomics: A 360° Approach”, and ICCRAM-Universidad de Burgos (Spain). We thank the EVA4.0 management team at FFWS.

## 7 Authors contribution

JJGP and JŠ designed the study. JJGP and JŠ participated in sample collection. JJGP design the sampling strategy, isolated DNA and COII identification of many termites used in this study, processed nest material to powder, carried out all experiments regarding nest DNA isolation, sequencing data processing and analysis. JH performed metabolite profiling setup, experiments, data processing, and writing of the corresponding metabolite sections at the manuscript. JJGP wrote the main body of the manuscript, JH and JŠ participated in the article writing and data interpretation.

## 9 Ethical statement

Sampling was carried out following Nagoya guidelines for biodiversity sampling (Convention on Biological Diversity, 2010) with the required research permits to JŠ and other members of FFWS, and following ethical behaviour towards local communities.

