## Supplementary figures for "Termite diet rather than geographical origin determines the microbiome composition and functional genetic structure of nests from South American and African representatives, as revealed by a multiomics approach"

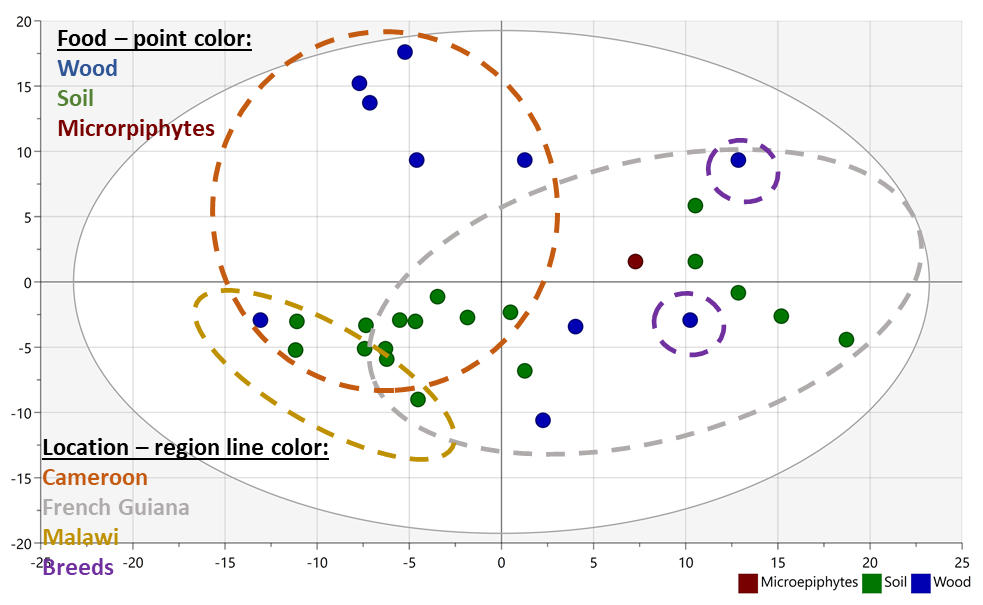

**Figure S1.** PCA plot using the complete untargeted profiling analysis. Samples are coloured according to feeding habits: green, soil; blue, wood; red, microepiphytes.

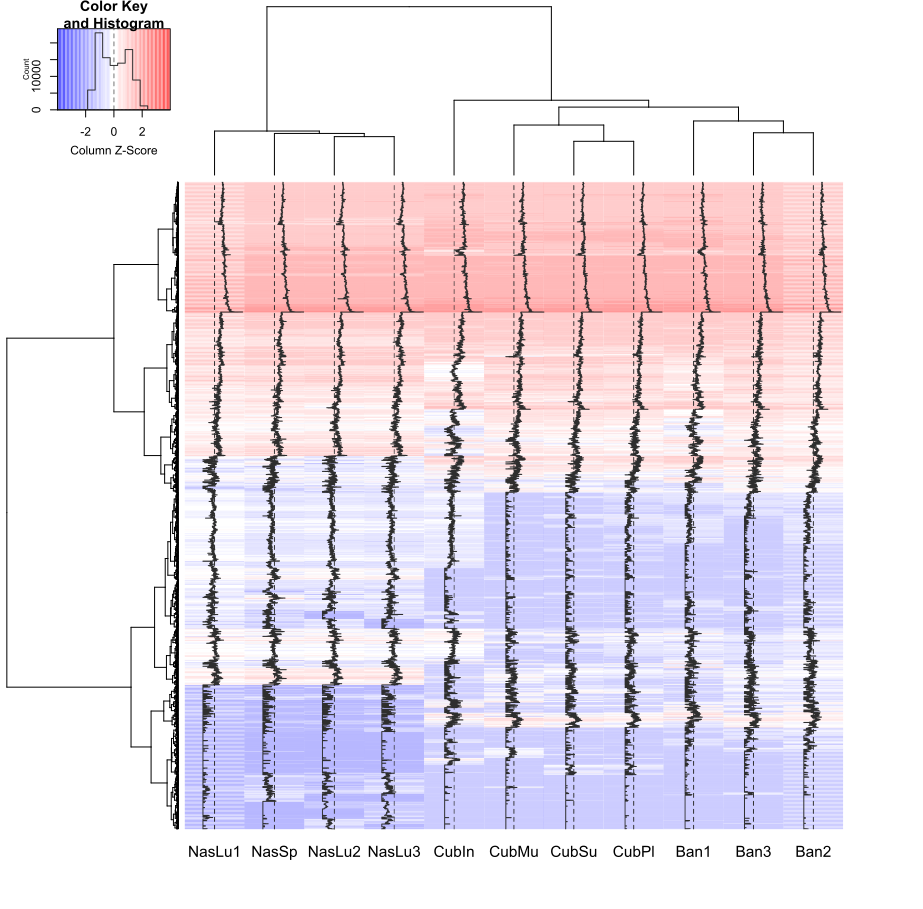

**Fig S2.** Heatmap showing the gene content differences and dendrograms of samples and genes, for all of those ORFs annotated to the PFAM database.

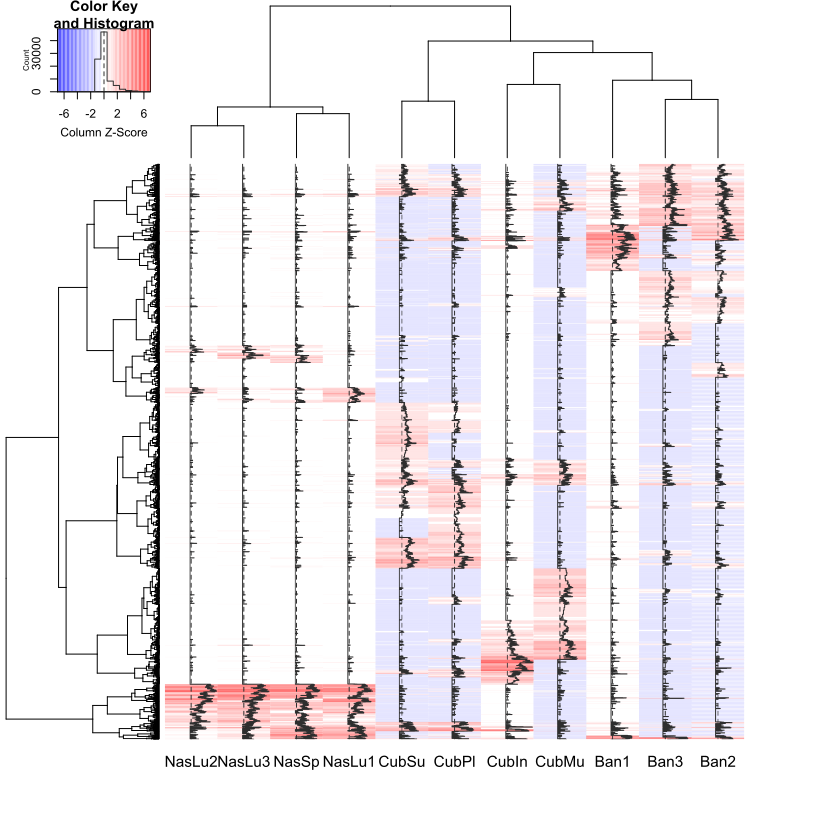

**Fig. S3.** Heatmap showing the gene content differences and dendrograms of samples and genes, for all of those ORFs annotated to the AromaDeg database.

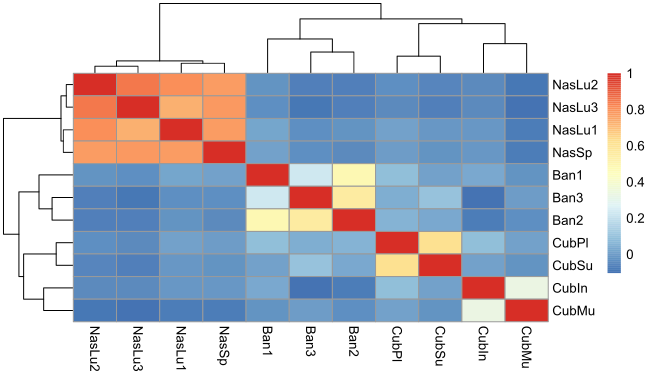

**Fig. S4.** Correlation heatmap of samples according to the genes annotated in CARD database.
