## Supplementary material 1 for "Termite diet rather than geographical origin determines the microbiome composition and functional genetic structure of nests from South American and African representatives, as revealed by a multiomics approach": common.js

```
// JavaScript Document
$(function(){
$('.main_menu li:eq(1)').live('mouseenter', function(){
	$('#menu_list').show();										
});
$('#menu_list').live('mouseleave', function(){
	$('#menu_list').hide();										
});
$('.main_menu li:eq(1)').live('mouseleave', function(){
	$('#menu_list').hide();										
});

$('.close').live('click',function(){
	$('.main_menu').fadeOut();
	$(this).removeClass("close").addClass("open");
})
$('.open').live('click',function(){
	$('.main_menu').fadeIn();
	$(this).removeClass("open").addClass("close");
})

});
//顶部导航

$(document).ready(function(){
	$(".img_show").each( function(i)
	{

		var currentPosition = 0;
		var slideHeight=110;
		var slides = $(this).find("li");
		//alert(slides.length);
		var numberOfSlides = slides.length;

		// Wrap all .slides with #slideInner div
		   slides
			.wrapAll('<div class="slideInner"></div>')
			// Float left to display horizontally, readjust .slides width
			.css({
			  //'float' : 'left',
			  'height' : slideHeight
			});

		// Insert controls in the DOM
		   $(this).find('.slideshow')
			.prepend('<input class="leftControl btn_up control" width="130" height="20" type="button"/>')
			.append('<input class="rightControl btn_down control" width="130" height="20" type="button"/>');
		// Hide left arrow control on first load
			  manageControls(currentPosition,$(this));

			  // Create event listeners for .controls clicks
			  $(this).find('.control')
				.bind('click',  {tou: $(this)}, function(event){
				  //alert(currentPosition);
				// Determine new position
				
				currentPosition = ($(this).attr('class')=='rightControl btn_down control') ? currentPosition+1 : currentPosition-1;
				// Hide / show controls
				manageControls(currentPosition, event.data.tou);
				// Move slideInner using margin-left
				//alert( $(this).parent().find('.slideInner') );
				event.data.tou.find('.slideInner').animate({
				  'marginTop' : slideHeight*(-currentPosition)
				});
			  });

			  // manageControls: Hides and Shows controls depending on currentPosition
			  function manageControls(position,lr){
				// Hide left arrow if position is first slide
				if(position==0){ $(lr).find('.leftControl').hide() } else{ $(lr).find('.leftControl').show() }
				// Hide right arrow if position is last slide
				if(position==numberOfSlides-4){ $(lr).find('.rightControl').hide() } else{ $(lr).find('.rightControl').show() }
			  }	

	});

  
  
});
//图片轮番
```
