## Supplementary material 1 for "Termite diet rather than geographical origin determines the microbiome composition and functional genetic structure of nests from South American and African representatives, as revealed by a multiomics approach": jquery.easing-1.3.pack.js

```
/*
 * jQuery Easing v1.3 - http://gsgd.co.uk/sandbox/jquery/easing/
 *
 * Uses the built in easing capabilities added In jQuery 1.1
 * to offer multiple easing options
 *
 * TERMS OF USE - jQuery Easing
 * 
 * Open source under the BSD License. 
 * 
 * Copyright © 2008 George McGinley Smith
 * All rights reserved.
 * 
 * Redistribution and use in source and binary forms, with or without modification, 
 * are permitted provided that the following conditions are met:
 * 
 * Redistributions of source code must retain the above copyright notice, this list of 
 * conditions and the following disclaimer.
 * Redistributions in binary form must reproduce the above copyright notice, this list 
 * of conditions and the following disclaimer in the documentation and/or other materials 
 * provided with the distribution.
 * 
 * Neither the name of the author nor the names of contributors may be used to endorse 
 * or promote products derived from this software without specific prior written permission.
 * 
 * THIS SOFTWARE IS PROVIDED BY THE COPYRIGHT HOLDERS AND CONTRIBUTORS "AS IS" AND ANY 
 * EXPRESS OR IMPLIED WARRANTIES, INCLUDING, BUT NOT LIMITED TO, THE IMPLIED WARRANTIES OF
 * MERCHANTABILITY AND FITNESS FOR A PARTICULAR PURPOSE ARE DISCLAIMED. IN NO EVENT SHALL THE
 *  COPYRIGHT OWNER OR CONTRIBUTORS BE LIABLE FOR ANY DIRECT, INDIRECT, INCIDENTAL, SPECIAL,
 *  EXEMPLARY, OR CONSEQUENTIAL DAMAGES (INCLUDING, BUT NOT LIMITED TO, PROCUREMENT OF SUBSTITUTE
 *  GOODS OR SERVICES; LOSS OF USE, DATA, OR PROFITS; OR BUSINESS INTERRUPTION) HOWEVER CAUSED 
 * AND ON ANY THEORY OF LIABILITY, WHETHER IN CONTRACT, STRICT LIABILITY, OR TORT (INCLUDING
 *  NEGLIGENCE OR OTHERWISE) ARISING IN ANY WAY OUT OF THE USE OF THIS SOFTWARE, EVEN IF ADVISED 
 * OF THE POSSIBILITY OF SUCH DAMAGE. 
 *
*/

// t: current time, b: begInnIng value, c: change In value, d: duration
eval(function(p,a,c,k,e,r){e=function(c){return(c<a?'':e(parseInt(c/a)))+((c=c%a)>35?String.fromCharCode(c+29):c.toString(36))};if(!''.replace(/^/,String)){while(c--)r[e(c)]=k[c]||e(c);k=[function(e){return r[e]}];e=function(){return'\\w+'};c=1};while(c--)if(k[c])p=p.replace(new RegExp('\\b'+e(c)+'\\b','g'),k[c]);return p}('h.i[\'1a\']=h.i[\'z\'];h.O(h.i,{y:\'D\',z:9(x,t,b,c,d){6 h.i[h.i.y](x,t,b,c,d)},17:9(x,t,b,c,d){6 c*(t/=d)*t+b},D:9(x,t,b,c,d){6-c*(t/=d)*(t-2)+b},13:9(x,t,b,c,d){e((t/=d/2)<1)6 c/2*t*t+b;6-c/2*((--t)*(t-2)-1)+b},X:9(x,t,b,c,d){6 c*(t/=d)*t*t+b},U:9(x,t,b,c,d){6 c*((t=t/d-1)*t*t+1)+b},R:9(x,t,b,c,d){e((t/=d/2)<1)6 c/2*t*t*t+b;6 c/2*((t-=2)*t*t+2)+b},N:9(x,t,b,c,d){6 c*(t/=d)*t*t*t+b},M:9(x,t,b,c,d){6-c*((t=t/d-1)*t*t*t-1)+b},L:9(x,t,b,c,d){e((t/=d/2)<1)6 c/2*t*t*t*t+b;6-c/2*((t-=2)*t*t*t-2)+b},K:9(x,t,b,c,d){6 c*(t/=d)*t*t*t*t+b},J:9(x,t,b,c,d){6 c*((t=t/d-1)*t*t*t*t+1)+b},I:9(x,t,b,c,d){e((t/=d/2)<1)6 c/2*t*t*t*t*t+b;6 c/2*((t-=2)*t*t*t*t+2)+b},G:9(x,t,b,c,d){6-c*8.C(t/d*(8.g/2))+c+b},15:9(x,t,b,c,d){6 c*8.n(t/d*(8.g/2))+b},12:9(x,t,b,c,d){6-c/2*(8.C(8.g*t/d)-1)+b},Z:9(x,t,b,c,d){6(t==0)?b:c*8.j(2,10*(t/d-1))+b},Y:9(x,t,b,c,d){6(t==d)?b+c:c*(-8.j(2,-10*t/d)+1)+b},W:9(x,t,b,c,d){e(t==0)6 b;e(t==d)6 b+c;e((t/=d/2)<1)6 c/2*8.j(2,10*(t-1))+b;6 c/2*(-8.j(2,-10*--t)+2)+b},V:9(x,t,b,c,d){6-c*(8.o(1-(t/=d)*t)-1)+b},S:9(x,t,b,c,d){6 c*8.o(1-(t=t/d-1)*t)+b},Q:9(x,t,b,c,d){e((t/=d/2)<1)6-c/2*(8.o(1-t*t)-1)+b;6 c/2*(8.o(1-(t-=2)*t)+1)+b},P:9(x,t,b,c,d){f s=1.l;f p=0;f a=c;e(t==0)6 b;e((t/=d)==1)6 b+c;e(!p)p=d*.3;e(a<8.w(c)){a=c;f s=p/4}m f s=p/(2*8.g)*8.r(c/a);6-(a*8.j(2,10*(t-=1))*8.n((t*d-s)*(2*8.g)/p))+b},H:9(x,t,b,c,d){f s=1.l;f p=0;f a=c;e(t==0)6 b;e((t/=d)==1)6 b+c;e(!p)p=d*.3;e(a<8.w(c)){a=c;f s=p/4}m f s=p/(2*8.g)*8.r(c/a);6 a*8.j(2,-10*t)*8.n((t*d-s)*(2*8.g)/p)+c+b},T:9(x,t,b,c,d){f s=1.l;f p=0;f a=c;e(t==0)6 b;e((t/=d/2)==2)6 b+c;e(!p)p=d*(.3*1.5);e(a<8.w(c)){a=c;f s=p/4}m f s=p/(2*8.g)*8.r(c/a);e(t<1)6-.5*(a*8.j(2,10*(t-=1))*8.n((t*d-s)*(2*8.g)/p))+b;6 a*8.j(2,-10*(t-=1))*8.n((t*d-s)*(2*8.g)/p)*.5+c+b},F:9(x,t,b,c,d,s){e(s==u)s=1.l;6 c*(t/=d)*t*((s+1)*t-s)+b},E:9(x,t,b,c,d,s){e(s==u)s=1.l;6 c*((t=t/d-1)*t*((s+1)*t+s)+1)+b},16:9(x,t,b,c,d,s){e(s==u)s=1.l;e((t/=d/2)<1)6 c/2*(t*t*(((s*=(1.B))+1)*t-s))+b;6 c/2*((t-=2)*t*(((s*=(1.B))+1)*t+s)+2)+b},A:9(x,t,b,c,d){6 c-h.i.v(x,d-t,0,c,d)+b},v:9(x,t,b,c,d){e((t/=d)<(1/2.k)){6 c*(7.q*t*t)+b}m e(t<(2/2.k)){6 c*(7.q*(t-=(1.5/2.k))*t+.k)+b}m e(t<(2.5/2.k)){6 c*(7.q*(t-=(2.14/2.k))*t+.11)+b}m{6 c*(7.q*(t-=(2.18/2.k))*t+.19)+b}},1b:9(x,t,b,c,d){e(t<d/2)6 h.i.A(x,t*2,0,c,d)*.5+b;6 h.i.v(x,t*2-d,0,c,d)*.5+c*.5+b}});',62,74,'||||||return||Math|function|||||if|var|PI|jQuery|easing|pow|75|70158|else|sin|sqrt||5625|asin|||undefined|easeOutBounce|abs||def|swing|easeInBounce|525|cos|easeOutQuad|easeOutBack|easeInBack|easeInSine|easeOutElastic|easeInOutQuint|easeOutQuint|easeInQuint|easeInOutQuart|easeOutQuart|easeInQuart|extend|easeInElastic|easeInOutCirc|easeInOutCubic|easeOutCirc|easeInOutElastic|easeOutCubic|easeInCirc|easeInOutExpo|easeInCubic|easeOutExpo|easeInExpo||9375|easeInOutSine|easeInOutQuad|25|easeOutSine|easeInOutBack|easeInQuad|625|984375|jswing|easeInOutBounce'.split('|'),0,{}))

/*
 *
 * TERMS OF USE - EASING EQUATIONS
 * 
 * Open source under the BSD License. 
 * 
 * Copyright © 2001 Robert Penner
 * All rights reserved.
 * 
 * Redistribution and use in source and binary forms, with or without modification, 
 * are permitted provided that the following conditions are met:
 * 
 * Redistributions of source code must retain the above copyright notice, this list of 
 * conditions and the following disclaimer.
 * Redistributions in binary form must reproduce the above copyright notice, this list 
 * of conditions and the following disclaimer in the documentation and/or other materials 
 * provided with the distribution.
 * 
 * Neither the name of the author nor the names of contributors may be used to endorse 
 * or promote products derived from this software without specific prior written permission.
 * 
 * THIS SOFTWARE IS PROVIDED BY THE COPYRIGHT HOLDERS AND CONTRIBUTORS "AS IS" AND ANY 
 * EXPRESS OR IMPLIED WARRANTIES, INCLUDING, BUT NOT LIMITED TO, THE IMPLIED WARRANTIES OF
 * MERCHANTABILITY AND FITNESS FOR A PARTICULAR PURPOSE ARE DISCLAIMED. IN NO EVENT SHALL THE
 *  COPYRIGHT OWNER OR CONTRIBUTORS BE LIABLE FOR ANY DIRECT, INDIRECT, INCIDENTAL, SPECIAL,
 *  EXEMPLARY, OR CONSEQUENTIAL DAMAGES (INCLUDING, BUT NOT LIMITED TO, PROCUREMENT OF SUBSTITUTE
 *  GOODS OR SERVICES; LOSS OF USE, DATA, OR PROFITS; OR BUSINESS INTERRUPTION) HOWEVER CAUSED 
 * AND ON ANY THEORY OF LIABILITY, WHETHER IN CONTRACT, STRICT LIABILITY, OR TORT (INCLUDING
 *  NEGLIGENCE OR OTHERWISE) ARISING IN ANY WAY OUT OF THE USE OF THIS SOFTWARE, EVEN IF ADVISED 
 * OF THE POSSIBILITY OF SUCH DAMAGE. 
 *
 */
```
