## Supplementary material 1 for "Termite diet rather than geographical origin determines the microbiome composition and functional genetic structure of nests from South American and African representatives, as revealed by a multiomics approach": jquery.fancybox-1.3.4.js

```
/*
 * FancyBox - jQuery Plugin
 * Simple and fancy lightbox alternative
 *
 * Examples and documentation at: http://fancybox.net
 *
 * Copyright (c) 2008 - 2010 Janis Skarnelis
 * That said, it is hardly a one-person project. Many people have submitted bugs, code, and offered their advice freely. Their support is greatly appreciated.
 *
 * Version: 1.3.4 (11/11/2010)
 * Requires: jQuery v1.3+
 *
 * Dual licensed under the MIT and GPL licenses:
 *   http://www.opensource.org/licenses/mit-license.php
 *   http://www.gnu.org/licenses/gpl.html
 */

;(function($) {
	var tmp, loading, overlay, wrap, outer, content, close, title, nav_left, nav_right,

		selectedIndex = 0, selectedOpts = {}, selectedArray = [], currentIndex = 0, currentOpts = {}, currentArray = [],

		ajaxLoader = null, imgPreloader = new Image(), imgRegExp = /\.(jpg|gif|png|bmp|jpeg)(.*)?$/i, swfRegExp = /[^\.]\.(swf)\s*$/i,

		loadingTimer, loadingFrame = 1,

		titleHeight = 0, titleStr = '', start_pos, final_pos, busy = false, fx = $.extend($('<div/>')[0], { prop: 0 }),

		isIE6 = $.browser.msie && $.browser.version < 7 && !window.XMLHttpRequest,

		/*
		 * Private methods 
		 */

		_abort = function() {
			loading.hide();

			imgPreloader.onerror = imgPreloader.onload = null;

			if (ajaxLoader) {
				ajaxLoader.abort();
			}

			tmp.empty();
		},

		_error = function() {
			if (false === selectedOpts.onError(selectedArray, selectedIndex, selectedOpts)) {
				loading.hide();
				busy = false;
				return;
			}

			selectedOpts.titleShow = false;

			selectedOpts.width = 'auto';
			selectedOpts.height = 'auto';

			tmp.html( '<p id="fancybox-error">无显著富集结果</p>' );

			_process_inline();
		},

		_start = function() {
			var obj = selectedArray[ selectedIndex ],
				href, 
				type, 
				title,
				str,
				emb,
				ret;

			_abort();

			selectedOpts = $.extend({}, $.fn.fancybox.defaults, (typeof $(obj).data('fancybox') == 'undefined' ? selectedOpts : $(obj).data('fancybox')));

			ret = selectedOpts.onStart(selectedArray, selectedIndex, selectedOpts);

			if (ret === false) {
				busy = false;
				return;
			} else if (typeof ret == 'object') {
				selectedOpts = $.extend(selectedOpts, ret);
			}

			title = selectedOpts.title || (obj.nodeName ? $(obj).attr('title') : obj.title) || '';

			if (obj.nodeName && !selectedOpts.orig) {
				selectedOpts.orig = $(obj).children("img:first").length ? $(obj).children("img:first") : $(obj);
			}

			if (title === '' && selectedOpts.orig && selectedOpts.titleFromAlt) {
				title = selectedOpts.orig.attr('alt');
			}

			href = selectedOpts.href || (obj.nodeName ? $(obj).attr('href') : obj.href) || null;

			if ((/^(?:javascript)/i).test(href) || href == '#') {
				href = null;
			}

			if (selectedOpts.type) {
				type = selectedOpts.type;

				if (!href) {
					href = selectedOpts.content;
				}

			} else if (selectedOpts.content) {
				type = 'html';

			} else if (href) {
				if (href.match(imgRegExp)) {
					type = 'image';

				} else if (href.match(swfRegExp)) {
					type = 'swf';

				} else if ($(obj).hasClass("iframe")) {
					type = 'iframe';

				} else if (href.indexOf("#") === 0) {
					type = 'inline';

				} else {
					type = 'ajax';
				}
			}

			if (!type) {
				_error();
				return;
			}

			if (type == 'inline') {
				obj	= href.substr(href.indexOf("#"));
				type = $(obj).length > 0 ? 'inline' : 'ajax';
			}

			selectedOpts.type = type;
			selectedOpts.href = href;
			selectedOpts.title = title;

			if (selectedOpts.autoDimensions) {
				if (selectedOpts.type == 'html' || selectedOpts.type == 'inline' || selectedOpts.type == 'ajax') {
					selectedOpts.width = 'auto';
					selectedOpts.height = 'auto';
				} else {
					selectedOpts.autoDimensions = false;	
				}
			}

			if (selectedOpts.modal) {
				selectedOpts.overlayShow = true;
				selectedOpts.hideOnOverlayClick = false;
				selectedOpts.hideOnContentClick = false;
				selectedOpts.enableEscapeButton = false;
				selectedOpts.showCloseButton = false;
			}

			selectedOpts.padding = parseInt(selectedOpts.padding, 10);
			selectedOpts.margin = parseInt(selectedOpts.margin, 10);

			tmp.css('padding', (selectedOpts.padding + selectedOpts.margin));

			$('.fancybox-inline-tmp').unbind('fancybox-cancel').bind('fancybox-change', function() {
				$(this).replaceWith(content.children());				
			});

			switch (type) {
				case 'html' :
					tmp.html( selectedOpts.content );
					_process_inline();
				break;

				case 'inline' :
					if ( $(obj).parent().is('#fancybox-content') === true) {
						busy = false;
						return;
					}

					$('<div class="fancybox-inline-tmp" />')
						.hide()
						.insertBefore( $(obj) )
						.bind('fancybox-cleanup', function() {
							$(this).replaceWith(content.children());
						}).bind('fancybox-cancel', function() {
							$(this).replaceWith(tmp.children());
						});

					$(obj).appendTo(tmp);

					_process_inline();
				break;

				case 'image':
					busy = false;

					$.fancybox.showActivity();

					imgPreloader = new Image();

					imgPreloader.onerror = function() {
						_error();
					};

					imgPreloader.onload = function() {
						busy = true;

						imgPreloader.onerror = imgPreloader.onload = null;

						_process_image();
					};

					imgPreloader.src = href;
				break;

				case 'swf':
					selectedOpts.scrolling = 'no';

					str = '<object classid="clsid:D27CDB6E-AE6D-11cf-96B8-444553540000" width="' + selectedOpts.width + '" height="' + selectedOpts.height + '"><param name="movie" value="' + href + '"></param>';
					emb = '';

					$.each(selectedOpts.swf, function(name, val) {
						str += '<param name="' + name + '" value="' + val + '"></param>';
						emb += ' ' + name + '="' + val + '"';
					});

					str += '<embed src="' + href + '" type="application/x-shockwave-flash" width="' + selectedOpts.width + '" height="' + selectedOpts.height + '"' + emb + '></embed></object>';

					tmp.html(str);

					_process_inline();
				break;

				case 'ajax':
					busy = false;

					$.fancybox.showActivity();

					selectedOpts.ajax.win = selectedOpts.ajax.success;

					ajaxLoader = $.ajax($.extend({}, selectedOpts.ajax, {
						url	: href,
						data : selectedOpts.ajax.data || {},
						error : function(XMLHttpRequest, textStatus, errorThrown) {
							if ( XMLHttpRequest.status > 0 ) {
								_error();
							}
						},
						success : function(data, textStatus, XMLHttpRequest) {
							var o = typeof XMLHttpRequest == 'object' ? XMLHttpRequest : ajaxLoader;
							if (o.status == 200) {
								if ( typeof selectedOpts.ajax.win == 'function' ) {
									ret = selectedOpts.ajax.win(href, data, textStatus, XMLHttpRequest);

									if (ret === false) {
										loading.hide();
										return;
									} else if (typeof ret == 'string' || typeof ret == 'object') {
										data = ret;
									}
								}

								tmp.html( data );
								_process_inline();
							}
						}
					}));

				break;

				case 'iframe':
					_show();
				break;
			}
		},

		_process_inline = function() {
			var
				w = selectedOpts.width,
				h = selectedOpts.height;

			if (w.toString().indexOf('%') > -1) {
				w = parseInt( ($(window).width() - (selectedOpts.margin * 2)) * parseFloat(w) / 100, 10) + 'px';

			} else {
				w = w == 'auto' ? 'auto' : w + 'px';	
			}

			if (h.toString().indexOf('%') > -1) {
				h = parseInt( ($(window).height() - (selectedOpts.margin * 2)) * parseFloat(h) / 100, 10) + 'px';

			} else {
				h = h == 'auto' ? 'auto' : h + 'px';	
			}

			tmp.wrapInner('<div style="width:' + w + ';height:' + h + ';overflow: ' + (selectedOpts.scrolling == 'auto' ? 'auto' : (selectedOpts.scrolling == 'yes' ? 'scroll' : 'hidden')) + ';position:relative;"></div>');

			selectedOpts.width = tmp.width();
			selectedOpts.height = tmp.height();

			_show();
		},

		_process_image = function() {
			selectedOpts.width = imgPreloader.width;
			selectedOpts.height = imgPreloader.height;

			$("<img />").attr({
				'id' : 'fancybox-img',
				'src' : imgPreloader.src,
				'alt' : selectedOpts.title
			}).appendTo( tmp );

			_show();
		},

		_show = function() {
			var pos, equal;

			loading.hide();

			if (wrap.is(":visible") && false === currentOpts.onCleanup(currentArray, currentIndex, currentOpts)) {
				$.event.trigger('fancybox-cancel');

				busy = false;
				return;
			}

			busy = true;

			$(content.add( overlay )).unbind();

			$(window).unbind("resize.fb scroll.fb");
			$(document).unbind('keydown.fb');

			if (wrap.is(":visible") && currentOpts.titlePosition !== 'outside') {
				wrap.css('height', wrap.height());
			}

			currentArray = selectedArray;
			currentIndex = selectedIndex;
			currentOpts = selectedOpts;

			if (currentOpts.overlayShow) {
				overlay.css({
					'background-color' : currentOpts.overlayColor,
					'opacity' : currentOpts.overlayOpacity,
					'cursor' : currentOpts.hideOnOverlayClick ? 'pointer' : 'auto',
					'height' : $(document).height()
				});

				if (!overlay.is(':visible')) {
					if (isIE6) {
						$('select:not(#fancybox-tmp select)').filter(function() {
							return this.style.visibility !== 'hidden';
						}).css({'visibility' : 'hidden'}).one('fancybox-cleanup', function() {
							this.style.visibility = 'inherit';
						});
					}

					overlay.show();
				}
			} else {
				overlay.hide();
			}

			final_pos = _get_zoom_to();

			_process_title();

			if (wrap.is(":visible")) {
				$( close.add( nav_left ).add( nav_right ) ).hide();

				pos = wrap.position(),

				start_pos = {
					top	 : pos.top,
					left : pos.left,
					width : wrap.width(),
					height : wrap.height()
				};

				equal = (start_pos.width == final_pos.width && start_pos.height == final_pos.height);

				content.fadeTo(currentOpts.changeFade, 0.3, function() {
					var finish_resizing = function() {
						content.html( tmp.contents() ).fadeTo(currentOpts.changeFade, 1, _finish);
					};

					$.event.trigger('fancybox-change');

					content
						.empty()
						.removeAttr('filter')
						.css({
							'border-width' : currentOpts.padding,
							'width'	: final_pos.width - currentOpts.padding * 2,
							'height' : selectedOpts.autoDimensions ? 'auto' : final_pos.height - titleHeight - currentOpts.padding * 2
						});

					if (equal) {
						finish_resizing();

					} else {
						fx.prop = 0;

						$(fx).animate({prop: 1}, {
							 duration : currentOpts.changeSpeed,
							 easing : currentOpts.easingChange,
							 step : _draw,
							 complete : finish_resizing
						});
					}
				});

				return;
			}

			wrap.removeAttr("style");

			content.css('border-width', currentOpts.padding);

			if (currentOpts.transitionIn == 'elastic') {
				start_pos = _get_zoom_from();

				content.html( tmp.contents() );

				wrap.show();

				if (currentOpts.opacity) {
					final_pos.opacity = 0;
				}

				fx.prop = 0;

				$(fx).animate({prop: 1}, {
					 duration : currentOpts.speedIn,
					 easing : currentOpts.easingIn,
					 step : _draw,
					 complete : _finish
				});

				return;
			}

			if (currentOpts.titlePosition == 'inside' && titleHeight > 0) {	
				title.show();	
			}

			content
				.css({
					'width' : final_pos.width - currentOpts.padding * 2,
					'height' : selectedOpts.autoDimensions ? 'auto' : final_pos.height - titleHeight - currentOpts.padding * 2
				})
				.html( tmp.contents() );

			wrap
				.css(final_pos)
				.fadeIn( currentOpts.transitionIn == 'none' ? 0 : currentOpts.speedIn, _finish );
		},

		_format_title = function(title) {
			if (title && title.length) {
				if (currentOpts.titlePosition == 'float') {
					return '<table id="fancybox-title-float-wrap" cellpadding="0" cellspacing="0"><tr><td id="fancybox-title-float-left"></td><td id="fancybox-title-float-main">' + title + '</td><td id="fancybox-title-float-right"></td></tr></table>';
				}

				return '<div id="fancybox-title-' + currentOpts.titlePosition + '">' + title + '</div>';
			}

			return false;
		},

		_process_title = function() {
			titleStr = currentOpts.title || '';
			titleHeight = 0;

			title
				.empty()
				.removeAttr('style')
				.removeClass();

			if (currentOpts.titleShow === false) {
				title.hide();
				return;
			}

			titleStr = $.isFunction(currentOpts.titleFormat) ? currentOpts.titleFormat(titleStr, currentArray, currentIndex, currentOpts) : _format_title(titleStr);

			if (!titleStr || titleStr === '') {
				title.hide();
				return;
			}

			title
				.addClass('fancybox-title-' + currentOpts.titlePosition)
				.html( titleStr )
				.appendTo( 'body' )
				.show();

			switch (currentOpts.titlePosition) {
				case 'inside':
					title
						.css({
							'width' : final_pos.width - (currentOpts.padding * 2),
							'marginLeft' : currentOpts.padding,
							'marginRight' : currentOpts.padding
						});

					titleHeight = title.outerHeight(true);

					title.appendTo( outer );

					final_pos.height += titleHeight;
				break;

				case 'over':
					title
						.css({
							'marginLeft' : currentOpts.padding,
							'width'	: final_pos.width - (currentOpts.padding * 2),
							'bottom' : currentOpts.padding
						})
						.appendTo( outer );
				break;

				case 'float':
					title
						.css('left', parseInt((title.width() - final_pos.width - 40)/ 2, 10) * -1)
						.appendTo( wrap );
				break;

				default:
					title
						.css({
							'width' : final_pos.width - (currentOpts.padding * 2),
							'paddingLeft' : currentOpts.padding,
							'paddingRight' : currentOpts.padding
						})
						.appendTo( wrap );
				break;
			}

			title.hide();
		},

		_set_navigation = function() {
			if (currentOpts.enableEscapeButton || currentOpts.enableKeyboardNav) {
				$(document).bind('keydown.fb', function(e) {
					if (e.keyCode == 27 && currentOpts.enableEscapeButton) {
						e.preventDefault();
						$.fancybox.close();

					} else if ((e.keyCode == 37 || e.keyCode == 39) && currentOpts.enableKeyboardNav && e.target.tagName !== 'INPUT' && e.target.tagName !== 'TEXTAREA' && e.target.tagName !== 'SELECT') {
						e.preventDefault();
						$.fancybox[ e.keyCode == 37 ? 'prev' : 'next']();
					}
				});
			}

			if (!currentOpts.showNavArrows) { 
				nav_left.hide();
				nav_right.hide();
				return;
			}

			if ((currentOpts.cyclic && currentArray.length > 1) || currentIndex !== 0) {
				nav_left.show();
			}

			if ((currentOpts.cyclic && currentArray.length > 1) || currentIndex != (currentArray.length -1)) {
				nav_right.show();
			}
		},

		_finish = function () {
			if (!$.support.opacity) {
				content.get(0).style.removeAttribute('filter');
				wrap.get(0).style.removeAttribute('filter');
			}

			if (selectedOpts.autoDimensions) {
				content.css('height', 'auto');
			}

			wrap.css('height', 'auto');

			if (titleStr && titleStr.length) {
				title.show();
			}

			if (currentOpts.showCloseButton) {
				close.show();
			}

			_set_navigation();
	
			if (currentOpts.hideOnContentClick)	{
				content.bind('click', $.fancybox.close);
			}

			if (currentOpts.hideOnOverlayClick)	{
				overlay.bind('click', $.fancybox.close);
			}

			$(window).bind("resize.fb", $.fancybox.resize);

			if (currentOpts.centerOnScroll) {
				$(window).bind("scroll.fb", $.fancybox.center);
			}

			if (currentOpts.type == 'iframe') {
				$('<iframe id="fancybox-frame" name="fancybox-frame' + new Date().getTime() + '" frameborder="0" hspace="0" ' + ($.browser.msie ? 'allowtransparency="true""' : '') + ' scrolling="' + selectedOpts.scrolling + '" src="' + currentOpts.href + '"></iframe>').appendTo(content);
			}

			wrap.show();

			busy = false;

			$.fancybox.center();

			currentOpts.onComplete(currentArray, currentIndex, currentOpts);

			_preload_images();
		},

		_preload_images = function() {
			var href, 
				objNext;

			if ((currentArray.length -1) > currentIndex) {
				href = currentArray[ currentIndex + 1 ].href;

				if (typeof href !== 'undefined' && href.match(imgRegExp)) {
					objNext = new Image();
					objNext.src = href;
				}
			}

			if (currentIndex > 0) {
				href = currentArray[ currentIndex - 1 ].href;

				if (typeof href !== 'undefined' && href.match(imgRegExp)) {
					objNext = new Image();
					objNext.src = href;
				}
			}
		},

		_draw = function(pos) {
			var dim = {
				width : parseInt(start_pos.width + (final_pos.width - start_pos.width) * pos, 10),
				height : parseInt(start_pos.height + (final_pos.height - start_pos.height) * pos, 10),

				top : parseInt(start_pos.top + (final_pos.top - start_pos.top) * pos, 10),
				left : parseInt(start_pos.left + (final_pos.left - start_pos.left) * pos, 10)
			};

			if (typeof final_pos.opacity !== 'undefined') {
				dim.opacity = pos < 0.5 ? 0.5 : pos;
			}

			wrap.css(dim);

			content.css({
				'width' : dim.width - currentOpts.padding * 2,
				'height' : dim.height - (titleHeight * pos) - currentOpts.padding * 2
			});
		},

		_get_viewport = function() {
			return [
				$(window).width() - (currentOpts.margin * 2),
				$(window).height() - (currentOpts.margin * 2),
				$(document).scrollLeft() + currentOpts.margin,
				$(document).scrollTop() + currentOpts.margin
			];
		},

		_get_zoom_to = function () {
			var view = _get_viewport(),
				to = {},
				resize = currentOpts.autoScale,
				double_padding = currentOpts.padding * 2,
				ratio;

			if (currentOpts.width.toString().indexOf('%') > -1) {
				to.width = parseInt((view[0] * parseFloat(currentOpts.width)) / 100, 10);
			} else {
				to.width = currentOpts.width + double_padding;
			}

			if (currentOpts.height.toString().indexOf('%') > -1) {
				to.height = parseInt((view[1] * parseFloat(currentOpts.height)) / 100, 10);
			} else {
				to.height = currentOpts.height + double_padding;
			}

			if (resize && (to.width > view[0] || to.height > view[1])) {
				if (selectedOpts.type == 'image' || selectedOpts.type == 'swf') {
					ratio = (currentOpts.width ) / (currentOpts.height );

					if ((to.width ) > view[0]) {
						to.width = view[0];
						to.height = parseInt(((to.width - double_padding) / ratio) + double_padding, 10);
					}

					if ((to.height) > view[1]) {
						to.height = view[1];
						to.width = parseInt(((to.height - double_padding) * ratio) + double_padding, 10);
					}

				} else {
					to.width = Math.min(to.width, view[0]);
					to.height = Math.min(to.height, view[1]);
				}
			}

			to.top = parseInt(Math.max(view[3] - 20, view[3] + ((view[1] - to.height - 40) * 0.5)), 10);
			to.left = parseInt(Math.max(view[2] - 20, view[2] + ((view[0] - to.width - 40) * 0.5)), 10);

			return to;
		},

		_get_obj_pos = function(obj) {
			var pos = obj.offset();

			pos.top += parseInt( obj.css('paddingTop'), 10 ) || 0;
			pos.left += parseInt( obj.css('paddingLeft'), 10 ) || 0;

			pos.top += parseInt( obj.css('border-top-width'), 10 ) || 0;
			pos.left += parseInt( obj.css('border-left-width'), 10 ) || 0;

			pos.width = obj.width();
			pos.height = obj.height();

			return pos;
		},

		_get_zoom_from = function() {
			var orig = selectedOpts.orig ? $(selectedOpts.orig) : false,
				from = {},
				pos,
				view;

			if (orig && orig.length) {
				pos = _get_obj_pos(orig);

				from = {
					width : pos.width + (currentOpts.padding * 2),
					height : pos.height + (currentOpts.padding * 2),
					top	: pos.top - currentOpts.padding - 20,
					left : pos.left - currentOpts.padding - 20
				};

			} else {
				view = _get_viewport();

				from = {
					width : currentOpts.padding * 2,
					height : currentOpts.padding * 2,
					top	: parseInt(view[3] + view[1] * 0.5, 10),
					left : parseInt(view[2] + view[0] * 0.5, 10)
				};
			}

			return from;
		},

		_animate_loading = function() {
			if (!loading.is(':visible')){
				clearInterval(loadingTimer);
				return;
			}

			$('div', loading).css('top', (loadingFrame * -40) + 'px');

			loadingFrame = (loadingFrame + 1) % 12;
		};

	/*
	 * Public methods 
	 */

	$.fn.fancybox = function(options) {
		if (!$(this).length) {
			return this;
		}

		$(this)
			.data('fancybox', $.extend({}, options, ($.metadata ? $(this).metadata() : {})))
			.unbind('click.fb')
			.bind('click.fb', function(e) {
				e.preventDefault();

				if (busy) {
					return;
				}

				busy = true;

				$(this).blur();

				selectedArray = [];
				selectedIndex = 0;

				var rel = $(this).attr('rel') || '';

				if (!rel || rel == '' || rel === 'nofollow') {
					selectedArray.push(this);

				} else {
					selectedArray = $("a[rel=" + rel + "], area[rel=" + rel + "]");
					selectedIndex = selectedArray.index( this );
				}

				_start();

				return;
			});

		return this;
	};

	$.fancybox = function(obj) {
		var opts;

		if (busy) {
			return;
		}

		busy = true;
		opts = typeof arguments[1] !== 'undefined' ? arguments[1] : {};

		selectedArray = [];
		selectedIndex = parseInt(opts.index, 10) || 0;

		if ($.isArray(obj)) {
			for (var i = 0, j = obj.length; i < j; i++) {
				if (typeof obj[i] == 'object') {
					$(obj[i]).data('fancybox', $.extend({}, opts, obj[i]));
				} else {
					obj[i] = $({}).data('fancybox', $.extend({content : obj[i]}, opts));
				}
			}

			selectedArray = jQuery.merge(selectedArray, obj);

		} else {
			if (typeof obj == 'object') {
				$(obj).data('fancybox', $.extend({}, opts, obj));
			} else {
				obj = $({}).data('fancybox', $.extend({content : obj}, opts));
			}

			selectedArray.push(obj);
		}

		if (selectedIndex > selectedArray.length || selectedIndex < 0) {
			selectedIndex = 0;
		}

		_start();
	};

	$.fancybox.showActivity = function() {
		clearInterval(loadingTimer);

		loading.show();
		loadingTimer = setInterval(_animate_loading, 66);
	};

	$.fancybox.hideActivity = function() {
		loading.hide();
	};

	$.fancybox.next = function() {
		return $.fancybox.pos( currentIndex + 1);
	};

	$.fancybox.prev = function() {
		return $.fancybox.pos( currentIndex - 1);
	};

	$.fancybox.pos = function(pos) {
		if (busy) {
			return;
		}

		pos = parseInt(pos);

		selectedArray = currentArray;

		if (pos > -1 && pos < currentArray.length) {
			selectedIndex = pos;
			_start();

		} else if (currentOpts.cyclic && currentArray.length > 1) {
			selectedIndex = pos >= currentArray.length ? 0 : currentArray.length - 1;
			_start();
		}

		return;
	};

	$.fancybox.cancel = function() {
		if (busy) {
			return;
		}

		busy = true;

		$.event.trigger('fancybox-cancel');

		_abort();

		selectedOpts.onCancel(selectedArray, selectedIndex, selectedOpts);

		busy = false;
	};

	// Note: within an iframe use - parent.$.fancybox.close();
	$.fancybox.close = function() {
		if (busy || wrap.is(':hidden')) {
			return;
		}

		busy = true;

		if (currentOpts && false === currentOpts.onCleanup(currentArray, currentIndex, currentOpts)) {
			busy = false;
			return;
		}

		_abort();

		$(close.add( nav_left ).add( nav_right )).hide();

		$(content.add( overlay )).unbind();

		$(window).unbind("resize.fb scroll.fb");
		$(document).unbind('keydown.fb');

		content.find('iframe').attr('src', isIE6 && /^https/i.test(window.location.href || '') ? 'javascript:void(false)' : 'about:blank');

		if (currentOpts.titlePosition !== 'inside') {
			title.empty();
		}

		wrap.stop();

		function _cleanup() {
			overlay.fadeOut('fast');

			title.empty().hide();
			wrap.hide();

			$.event.trigger('fancybox-cleanup');

			content.empty();

			currentOpts.onClosed(currentArray, currentIndex, currentOpts);

			currentArray = selectedOpts	= [];
			currentIndex = selectedIndex = 0;
			currentOpts = selectedOpts	= {};

			busy = false;
		}

		if (currentOpts.transitionOut == 'elastic') {
			start_pos = _get_zoom_from();

			var pos = wrap.position();

			final_pos = {
				top	 : pos.top ,
				left : pos.left,
				width :	wrap.width(),
				height : wrap.height()
			};

			if (currentOpts.opacity) {
				final_pos.opacity = 1;
			}

			title.empty().hide();

			fx.prop = 1;

			$(fx).animate({ prop: 0 }, {
				 duration : currentOpts.speedOut,
				 easing : currentOpts.easingOut,
				 step : _draw,
				 complete : _cleanup
			});

		} else {
			wrap.fadeOut( currentOpts.transitionOut == 'none' ? 0 : currentOpts.speedOut, _cleanup);
		}
	};

	$.fancybox.resize = function() {
		if (overlay.is(':visible')) {
			overlay.css('height', $(document).height());
		}

		$.fancybox.center(true);
	};

	$.fancybox.center = function() {
		var view, align;

		if (busy) {
			return;	
		}

		align = arguments[0] === true ? 1 : 0;
		view = _get_viewport();

		if (!align && (wrap.width() > view[0] || wrap.height() > view[1])) {
			return;	
		}

		wrap
			.stop()
			.animate({
				'top' : parseInt(Math.max(view[3] - 20, view[3] + ((view[1] - content.height() - 40) * 0.5) - currentOpts.padding)),
				'left' : parseInt(Math.max(view[2] - 20, view[2] + ((view[0] - content.width() - 40) * 0.5) - currentOpts.padding))
			}, typeof arguments[0] == 'number' ? arguments[0] : 200);
	};

	$.fancybox.init = function() {
		if ($("#fancybox-wrap").length) {
			return;
		}

		$('body').append(
			tmp	= $('<div id="fancybox-tmp"></div>'),
			loading	= $('<div id="fancybox-loading"><div></div></div>'),
			overlay	= $('<div id="fancybox-overlay"></div>'),
			wrap = $('<div id="fancybox-wrap"></div>')
		);

		outer = $('<div id="fancybox-outer"></div>')
			.append('<div class="fancybox-bg" id="fancybox-bg-n"></div><div class="fancybox-bg" id="fancybox-bg-ne"></div><div class="fancybox-bg" id="fancybox-bg-e"></div><div class="fancybox-bg" id="fancybox-bg-se"></div><div class="fancybox-bg" id="fancybox-bg-s"></div><div class="fancybox-bg" id="fancybox-bg-sw"></div><div class="fancybox-bg" id="fancybox-bg-w"></div><div class="fancybox-bg" id="fancybox-bg-nw"></div>')
			.appendTo( wrap );

		outer.append(
			content = $('<div id="fancybox-content"></div>'),
			close = $('<a id="fancybox-close"></a>'),
			title = $('<div id="fancybox-title"></div>'),

			nav_left = $('<a href="javascript:;" id="fancybox-left"><span class="fancy-ico" id="fancybox-left-ico"></span></a>'),
			nav_right = $('<a href="javascript:;" id="fancybox-right"><span class="fancy-ico" id="fancybox-right-ico"></span></a>')
		);

		close.click($.fancybox.close);
		loading.click($.fancybox.cancel);

		nav_left.click(function(e) {
			e.preventDefault();
			$.fancybox.prev();
		});

		nav_right.click(function(e) {
			e.preventDefault();
			$.fancybox.next();
		});

		if ($.fn.mousewheel) {
			wrap.bind('mousewheel.fb', function(e, delta) {
				if (busy) {
					e.preventDefault();

				} else if ($(e.target).get(0).clientHeight == 0 || $(e.target).get(0).scrollHeight === $(e.target).get(0).clientHeight) {
					e.preventDefault();
					$.fancybox[ delta > 0 ? 'prev' : 'next']();
				}
			});
		}

		if (!$.support.opacity) {
			wrap.addClass('fancybox-ie');
		}

		if (isIE6) {
			loading.addClass('fancybox-ie6');
			wrap.addClass('fancybox-ie6');

			$('<iframe id="fancybox-hide-sel-frame" src="' + (/^https/i.test(window.location.href || '') ? 'javascript:void(false)' : 'about:blank' ) + '" scrolling="no" border="0" frameborder="0" tabindex="-1"></iframe>').prependTo(outer);
		}
	};

	$.fn.fancybox.defaults = {
		padding : 10,
		margin : 40,
		opacity : false,
		modal : false,
		cyclic : false,
		scrolling : 'auto',	// 'auto', 'yes' or 'no'

		width : 560,
		height : 340,

		autoScale : true,
		autoDimensions : true,
		centerOnScroll : false,

		ajax : {},
		swf : { wmode: 'transparent' },

		hideOnOverlayClick : true,
		hideOnContentClick : false,

		overlayShow : true,
		overlayOpacity : 0.7,
		overlayColor : '#777',

		titleShow : true,
		titlePosition : 'float', // 'float', 'outside', 'inside' or 'over'
		titleFormat : null,
		titleFromAlt : false,

		transitionIn : 'fade', // 'elastic', 'fade' or 'none'
		transitionOut : 'fade', // 'elastic', 'fade' or 'none'

		speedIn : 300,
		speedOut : 300,

		changeSpeed : 300,
		changeFade : 'fast',

		easingIn : 'swing',
		easingOut : 'swing',

		showCloseButton	 : true,
		showNavArrows : true,
		enableEscapeButton : true,
		enableKeyboardNav : true,

		onStart : function(){},
		onCancel : function(){},
		onComplete : function(){},
		onCleanup : function(){},
		onClosed : function(){},
		onError : function(){}
	};

	$(document).ready(function() {
		$.fancybox.init();
	});

})(jQuery);
```
