## Supplementary material 1 for "Termite diet rather than geographical origin determines the microbiome composition and functional genetic structure of nests from South American and African representatives, as revealed by a multiomics approach": jquery.fancybox-1.3.4.pack.js

```
/*
 * FancyBox - jQuery Plugin
 * Simple and fancy lightbox alternative
 *
 * Examples and documentation at: http://fancybox.net
 * 
 * Copyright (c) 2008 - 2010 Janis Skarnelis
 * That said, it is hardly a one-person project. Many people have submitted bugs, code, and offered their advice freely. Their support is greatly appreciated.
 * 
 * Version: 1.3.4 (11/11/2010)
 * Requires: jQuery v1.3+
 *
 * Dual licensed under the MIT and GPL licenses:
 *   http://www.opensource.org/licenses/mit-license.php
 *   http://www.gnu.org/licenses/gpl.html
 */

;(function(b){var m,t,u,f,D,j,E,n,z,A,q=0,e={},o=[],p=0,d={},l=[],G=null,v=new Image,J=/\.(jpg|gif|png|bmp|jpeg)(.*)?$/i,W=/[^\.]\.(swf)\s*$/i,K,L=1,y=0,s="",r,i,h=false,B=b.extend(b("<div/>")[0],{prop:0}),M=b.browser.msie&&b.browser.version<7&&!window.XMLHttpRequest,N=function(){t.hide();v.onerror=v.onload=null;G&&G.abort();m.empty()},O=function(){if(false===e.onError(o,q,e)){t.hide();h=false}else{e.titleShow=false;e.width="auto";e.height="auto";m.html('<p id="fancybox-error">无显著富集结果</p>');
F()}},I=function(){var a=o[q],c,g,k,C,P,w;N();e=b.extend({},b.fn.fancybox.defaults,typeof b(a).data("fancybox")=="undefined"?e:b(a).data("fancybox"));w=e.onStart(o,q,e);if(w===false)h=false;else{if(typeof w=="object")e=b.extend(e,w);k=e.title||(a.nodeName?b(a).attr("title"):a.title)||"";if(a.nodeName&&!e.orig)e.orig=b(a).children("img:first").length?b(a).children("img:first"):b(a);if(k===""&&e.orig&&e.titleFromAlt)k=e.orig.attr("alt");c=e.href||(a.nodeName?b(a).attr("href"):a.href)||null;if(/^(?:javascript)/i.test(c)||
c=="#")c=null;if(e.type){g=e.type;if(!c)c=e.content}else if(e.content)g="html";else if(c)g=c.match(J)?"image":c.match(W)?"swf":b(a).hasClass("iframe")?"iframe":c.indexOf("#")===0?"inline":"ajax";if(g){if(g=="inline"){a=c.substr(c.indexOf("#"));g=b(a).length>0?"inline":"ajax"}e.type=g;e.href=c;e.title=k;if(e.autoDimensions)if(e.type=="html"||e.type=="inline"||e.type=="ajax"){e.width="auto";e.height="auto"}else e.autoDimensions=false;if(e.modal){e.overlayShow=true;e.hideOnOverlayClick=false;e.hideOnContentClick=
false;e.enableEscapeButton=false;e.showCloseButton=false}e.padding=parseInt(e.padding,10);e.margin=parseInt(e.margin,10);m.css("padding",e.padding+e.margin);b(".fancybox-inline-tmp").unbind("fancybox-cancel").bind("fancybox-change",function(){b(this).replaceWith(j.children())});switch(g){case "html":m.html(e.content);F();break;case "inline":if(b(a).parent().is("#fancybox-content")===true){h=false;break}b('<div class="fancybox-inline-tmp" />').hide().insertBefore(b(a)).bind("fancybox-cleanup",function(){b(this).replaceWith(j.children())}).bind("fancybox-cancel",
function(){b(this).replaceWith(m.children())});b(a).appendTo(m);F();break;case "image":h=false;b.fancybox.showActivity();v=new Image;v.onerror=function(){O()};v.onload=function(){h=true;v.onerror=v.onload=null;e.width=v.width;e.height=v.height;b("<img />").attr({id:"fancybox-img",src:v.src,alt:e.title}).appendTo(m);Q()};v.src=c;break;case "swf":e.scrolling="no";C='<object classid="clsid:D27CDB6E-AE6D-11cf-96B8-444553540000" width="'+e.width+'" height="'+e.height+'"><param name="movie" value="'+c+
'"></param>';P="";b.each(e.swf,function(x,H){C+='<param name="'+x+'" value="'+H+'"></param>';P+=" "+x+'="'+H+'"'});C+='<embed src="'+c+'" type="application/x-shockwave-flash" width="'+e.width+'" height="'+e.height+'"'+P+"></embed></object>";m.html(C);F();break;case "ajax":h=false;b.fancybox.showActivity();e.ajax.win=e.ajax.success;G=b.ajax(b.extend({},e.ajax,{url:c,data:e.ajax.data||{},error:function(x){x.status>0&&O()},success:function(x,H,R){if((typeof R=="object"?R:G).status==200){if(typeof e.ajax.win==
"function"){w=e.ajax.win(c,x,H,R);if(w===false){t.hide();return}else if(typeof w=="string"||typeof w=="object")x=w}m.html(x);F()}}}));break;case "iframe":Q()}}else O()}},F=function(){var a=e.width,c=e.height;a=a.toString().indexOf("%")>-1?parseInt((b(window).width()-e.margin*2)*parseFloat(a)/100,10)+"px":a=="auto"?"auto":a+"px";c=c.toString().indexOf("%")>-1?parseInt((b(window).height()-e.margin*2)*parseFloat(c)/100,10)+"px":c=="auto"?"auto":c+"px";m.wrapInner('<div style="width:'+a+";height:"+c+
";overflow: "+(e.scrolling=="auto"?"auto":e.scrolling=="yes"?"scroll":"hidden")+';position:relative;"></div>');e.width=m.width();e.height=m.height();Q()},Q=function(){var a,c;t.hide();if(f.is(":visible")&&false===d.onCleanup(l,p,d)){b.event.trigger("fancybox-cancel");h=false}else{h=true;b(j.add(u)).unbind();b(window).unbind("resize.fb scroll.fb");b(document).unbind("keydown.fb");f.is(":visible")&&d.titlePosition!=="outside"&&f.css("height",f.height());l=o;p=q;d=e;if(d.overlayShow){u.css({"background-color":d.overlayColor,
opacity:d.overlayOpacity,cursor:d.hideOnOverlayClick?"pointer":"auto",height:b(document).height()});if(!u.is(":visible")){M&&b("select:not(#fancybox-tmp select)").filter(function(){return this.style.visibility!=="hidden"}).css({visibility:"hidden"}).one("fancybox-cleanup",function(){this.style.visibility="inherit"});u.show()}}else u.hide();i=X();s=d.title||"";y=0;n.empty().removeAttr("style").removeClass();if(d.titleShow!==false){if(b.isFunction(d.titleFormat))a=d.titleFormat(s,l,p,d);else a=s&&s.length?
d.titlePosition=="float"?'<table id="fancybox-title-float-wrap" cellpadding="0" cellspacing="0"><tr><td id="fancybox-title-float-left"></td><td id="fancybox-title-float-main">'+s+'</td><td id="fancybox-title-float-right"></td></tr></table>':'<div id="fancybox-title-'+d.titlePosition+'">'+s+"</div>":false;s=a;if(!(!s||s==="")){n.addClass("fancybox-title-"+d.titlePosition).html(s).appendTo("body").show();switch(d.titlePosition){case "inside":n.css({width:i.width-d.padding*2,marginLeft:d.padding,marginRight:d.padding});
y=n.outerHeight(true);n.appendTo(D);i.height+=y;break;case "over":n.css({marginLeft:d.padding,width:i.width-d.padding*2,bottom:d.padding}).appendTo(D);break;case "float":n.css("left",parseInt((n.width()-i.width-40)/2,10)*-1).appendTo(f);break;default:n.css({width:i.width-d.padding*2,paddingLeft:d.padding,paddingRight:d.padding}).appendTo(f)}}}n.hide();if(f.is(":visible")){b(E.add(z).add(A)).hide();a=f.position();r={top:a.top,left:a.left,width:f.width(),height:f.height()};c=r.width==i.width&&r.height==
i.height;j.fadeTo(d.changeFade,0.3,function(){var g=function(){j.html(m.contents()).fadeTo(d.changeFade,1,S)};b.event.trigger("fancybox-change");j.empty().removeAttr("filter").css({"border-width":d.padding,width:i.width-d.padding*2,height:e.autoDimensions?"auto":i.height-y-d.padding*2});if(c)g();else{B.prop=0;b(B).animate({prop:1},{duration:d.changeSpeed,easing:d.easingChange,step:T,complete:g})}})}else{f.removeAttr("style");j.css("border-width",d.padding);if(d.transitionIn=="elastic"){r=V();j.html(m.contents());
f.show();if(d.opacity)i.opacity=0;B.prop=0;b(B).animate({prop:1},{duration:d.speedIn,easing:d.easingIn,step:T,complete:S})}else{d.titlePosition=="inside"&&y>0&&n.show();j.css({width:i.width-d.padding*2,height:e.autoDimensions?"auto":i.height-y-d.padding*2}).html(m.contents());f.css(i).fadeIn(d.transitionIn=="none"?0:d.speedIn,S)}}}},Y=function(){if(d.enableEscapeButton||d.enableKeyboardNav)b(document).bind("keydown.fb",function(a){if(a.keyCode==27&&d.enableEscapeButton){a.preventDefault();b.fancybox.close()}else if((a.keyCode==
37||a.keyCode==39)&&d.enableKeyboardNav&&a.target.tagName!=="INPUT"&&a.target.tagName!=="TEXTAREA"&&a.target.tagName!=="SELECT"){a.preventDefault();b.fancybox[a.keyCode==37?"prev":"next"]()}});if(d.showNavArrows){if(d.cyclic&&l.length>1||p!==0)z.show();if(d.cyclic&&l.length>1||p!=l.length-1)A.show()}else{z.hide();A.hide()}},S=function(){if(!b.support.opacity){j.get(0).style.removeAttribute("filter");f.get(0).style.removeAttribute("filter")}e.autoDimensions&&j.css("height","auto");f.css("height","auto");
s&&s.length&&n.show();d.showCloseButton&&E.show();Y();d.hideOnContentClick&&j.bind("click",b.fancybox.close);d.hideOnOverlayClick&&u.bind("click",b.fancybox.close);b(window).bind("resize.fb",b.fancybox.resize);d.centerOnScroll&&b(window).bind("scroll.fb",b.fancybox.center);if(d.type=="iframe")b('<iframe id="fancybox-frame" name="fancybox-frame'+(new Date).getTime()+'" frameborder="0" hspace="0" '+(b.browser.msie?'allowtransparency="true""':"")+' scrolling="'+e.scrolling+'" src="'+d.href+'"></iframe>').appendTo(j);
f.show();h=false;b.fancybox.center();d.onComplete(l,p,d);var a,c;if(l.length-1>p){a=l[p+1].href;if(typeof a!=="undefined"&&a.match(J)){c=new Image;c.src=a}}if(p>0){a=l[p-1].href;if(typeof a!=="undefined"&&a.match(J)){c=new Image;c.src=a}}},T=function(a){var c={width:parseInt(r.width+(i.width-r.width)*a,10),height:parseInt(r.height+(i.height-r.height)*a,10),top:parseInt(r.top+(i.top-r.top)*a,10),left:parseInt(r.left+(i.left-r.left)*a,10)};if(typeof i.opacity!=="undefined")c.opacity=a<0.5?0.5:a;f.css(c);
j.css({width:c.width-d.padding*2,height:c.height-y*a-d.padding*2})},U=function(){return[b(window).width()-d.margin*2,b(window).height()-d.margin*2,b(document).scrollLeft()+d.margin,b(document).scrollTop()+d.margin]},X=function(){var a=U(),c={},g=d.autoScale,k=d.padding*2;c.width=d.width.toString().indexOf("%")>-1?parseInt(a[0]*parseFloat(d.width)/100,10):d.width+k;c.height=d.height.toString().indexOf("%")>-1?parseInt(a[1]*parseFloat(d.height)/100,10):d.height+k;if(g&&(c.width>a[0]||c.height>a[1]))if(e.type==
"image"||e.type=="swf"){g=d.width/d.height;if(c.width>a[0]){c.width=a[0];c.height=parseInt((c.width-k)/g+k,10)}if(c.height>a[1]){c.height=a[1];c.width=parseInt((c.height-k)*g+k,10)}}else{c.width=Math.min(c.width,a[0]);c.height=Math.min(c.height,a[1])}c.top=parseInt(Math.max(a[3]-20,a[3]+(a[1]-c.height-40)*0.5),10);c.left=parseInt(Math.max(a[2]-20,a[2]+(a[0]-c.width-40)*0.5),10);return c},V=function(){var a=e.orig?b(e.orig):false,c={};if(a&&a.length){c=a.offset();c.top+=parseInt(a.css("paddingTop"),
10)||0;c.left+=parseInt(a.css("paddingLeft"),10)||0;c.top+=parseInt(a.css("border-top-width"),10)||0;c.left+=parseInt(a.css("border-left-width"),10)||0;c.width=a.width();c.height=a.height();c={width:c.width+d.padding*2,height:c.height+d.padding*2,top:c.top-d.padding-20,left:c.left-d.padding-20}}else{a=U();c={width:d.padding*2,height:d.padding*2,top:parseInt(a[3]+a[1]*0.5,10),left:parseInt(a[2]+a[0]*0.5,10)}}return c},Z=function(){if(t.is(":visible")){b("div",t).css("top",L*-40+"px");L=(L+1)%12}else clearInterval(K)};
b.fn.fancybox=function(a){if(!b(this).length)return this;b(this).data("fancybox",b.extend({},a,b.metadata?b(this).metadata():{})).unbind("click.fb").bind("click.fb",function(c){c.preventDefault();if(!h){h=true;b(this).blur();o=[];q=0;c=b(this).attr("rel")||"";if(!c||c==""||c==="nofollow")o.push(this);else{o=b("a[rel="+c+"], area[rel="+c+"]");q=o.index(this)}I()}});return this};b.fancybox=function(a,c){var g;if(!h){h=true;g=typeof c!=="undefined"?c:{};o=[];q=parseInt(g.index,10)||0;if(b.isArray(a)){for(var k=
0,C=a.length;k<C;k++)if(typeof a[k]=="object")b(a[k]).data("fancybox",b.extend({},g,a[k]));else a[k]=b({}).data("fancybox",b.extend({content:a[k]},g));o=jQuery.merge(o,a)}else{if(typeof a=="object")b(a).data("fancybox",b.extend({},g,a));else a=b({}).data("fancybox",b.extend({content:a},g));o.push(a)}if(q>o.length||q<0)q=0;I()}};b.fancybox.showActivity=function(){clearInterval(K);t.show();K=setInterval(Z,66)};b.fancybox.hideActivity=function(){t.hide()};b.fancybox.next=function(){return b.fancybox.pos(p+
1)};b.fancybox.prev=function(){return b.fancybox.pos(p-1)};b.fancybox.pos=function(a){if(!h){a=parseInt(a);o=l;if(a>-1&&a<l.length){q=a;I()}else if(d.cyclic&&l.length>1){q=a>=l.length?0:l.length-1;I()}}};b.fancybox.cancel=function(){if(!h){h=true;b.event.trigger("fancybox-cancel");N();e.onCancel(o,q,e);h=false}};b.fancybox.close=function(){function a(){u.fadeOut("fast");n.empty().hide();f.hide();b.event.trigger("fancybox-cleanup");j.empty();d.onClosed(l,p,d);l=e=[];p=q=0;d=e={};h=false}if(!(h||f.is(":hidden"))){h=
true;if(d&&false===d.onCleanup(l,p,d))h=false;else{N();b(E.add(z).add(A)).hide();b(j.add(u)).unbind();b(window).unbind("resize.fb scroll.fb");b(document).unbind("keydown.fb");j.find("iframe").attr("src",M&&/^https/i.test(window.location.href||"")?"javascript:void(false)":"about:blank");d.titlePosition!=="inside"&&n.empty();f.stop();if(d.transitionOut=="elastic"){r=V();var c=f.position();i={top:c.top,left:c.left,width:f.width(),height:f.height()};if(d.opacity)i.opacity=1;n.empty().hide();B.prop=1;
b(B).animate({prop:0},{duration:d.speedOut,easing:d.easingOut,step:T,complete:a})}else f.fadeOut(d.transitionOut=="none"?0:d.speedOut,a)}}};b.fancybox.resize=function(){u.is(":visible")&&u.css("height",b(document).height());b.fancybox.center(true)};b.fancybox.center=function(a){var c,g;if(!h){g=a===true?1:0;c=U();!g&&(f.width()>c[0]||f.height()>c[1])||f.stop().animate({top:parseInt(Math.max(c[3]-20,c[3]+(c[1]-j.height()-40)*0.5-d.padding)),left:parseInt(Math.max(c[2]-20,c[2]+(c[0]-j.width()-40)*0.5-
d.padding))},typeof a=="number"?a:200)}};b.fancybox.init=function(){if(!b("#fancybox-wrap").length){b("body").append(m=b('<div id="fancybox-tmp"></div>'),t=b('<div id="fancybox-loading"><div></div></div>'),u=b('<div id="fancybox-overlay"></div>'),f=b('<div id="fancybox-wrap"></div>'));D=b('<div id="fancybox-outer"></div>').append('<div class="fancybox-bg" id="fancybox-bg-n"></div><div class="fancybox-bg" id="fancybox-bg-ne"></div><div class="fancybox-bg" id="fancybox-bg-e"></div><div class="fancybox-bg" id="fancybox-bg-se"></div><div class="fancybox-bg" id="fancybox-bg-s"></div><div class="fancybox-bg" id="fancybox-bg-sw"></div><div class="fancybox-bg" id="fancybox-bg-w"></div><div class="fancybox-bg" id="fancybox-bg-nw"></div>').appendTo(f);
D.append(j=b('<div id="fancybox-content"></div>'),E=b('<a id="fancybox-close"></a>'),n=b('<div id="fancybox-title"></div>'),z=b('<a href="javascript:;" id="fancybox-left"><span class="fancy-ico" id="fancybox-left-ico"></span></a>'),A=b('<a href="javascript:;" id="fancybox-right"><span class="fancy-ico" id="fancybox-right-ico"></span></a>'));E.click(b.fancybox.close);t.click(b.fancybox.cancel);z.click(function(a){a.preventDefault();b.fancybox.prev()});A.click(function(a){a.preventDefault();b.fancybox.next()});
b.fn.mousewheel&&f.bind("mousewheel.fb",function(a,c){if(h)a.preventDefault();else if(b(a.target).get(0).clientHeight==0||b(a.target).get(0).scrollHeight===b(a.target).get(0).clientHeight){a.preventDefault();b.fancybox[c>0?"prev":"next"]()}});b.support.opacity||f.addClass("fancybox-ie");if(M){t.addClass("fancybox-ie6");f.addClass("fancybox-ie6");b('<iframe id="fancybox-hide-sel-frame" src="'+(/^https/i.test(window.location.href||"")?"javascript:void(false)":"about:blank")+'" scrolling="no" border="0" frameborder="0" tabindex="-1"></iframe>').prependTo(D)}}};
b.fn.fancybox.defaults={padding:10,margin:40,opacity:false,modal:false,cyclic:false,scrolling:"auto",width:560,height:340,autoScale:true,autoDimensions:true,centerOnScroll:false,ajax:{},swf:{wmode:"transparent"},hideOnOverlayClick:true,hideOnContentClick:false,overlayShow:true,overlayOpacity:0.7,overlayColor:"#777",titleShow:true,titlePosition:"float",titleFormat:null,titleFromAlt:false,transitionIn:"fade",transitionOut:"fade",speedIn:300,speedOut:300,changeSpeed:300,changeFade:"fast",easingIn:"swing",
easingOut:"swing",showCloseButton:true,showNavArrows:true,enableEscapeButton:true,enableKeyboardNav:true,onStart:function(){},onCancel:function(){},onComplete:function(){},onCleanup:function(){},onClosed:function(){},onError:function(){}};b(document).ready(function(){b.fancybox.init()})})(jQuery);
```
