## Supplementary material 1 for "Termite diet rather than geographical origin determines the microbiome composition and functional genetic structure of nests from South American and African representatives, as revealed by a multiomics approach": jquery.mousewheel-3.0.4.pack.js

```
/*! Copyright (c) 2010 Brandon Aaron (http://brandonaaron.net)
* Licensed under the MIT License (LICENSE.txt).
*
* Thanks to: http://adomas.org/javascript-mouse-wheel/ for some pointers.
* Thanks to: Mathias Bank(http://www.mathias-bank.de) for a scope bug fix.
* Thanks to: Seamus Leahy for adding deltaX and deltaY
*
* Version: 3.0.4
*
* Requires: 1.2.2+
*/

(function(d){function g(a){var b=a||window.event,i=[].slice.call(arguments,1),c=0,h=0,e=0;a=d.event.fix(b);a.type="mousewheel";if(a.wheelDelta)c=a.wheelDelta/120;if(a.detail)c=-a.detail/3;e=c;if(b.axis!==undefined&&b.axis===b.HORIZONTAL_AXIS){e=0;h=-1*c}if(b.wheelDeltaY!==undefined)e=b.wheelDeltaY/120;if(b.wheelDeltaX!==undefined)h=-1*b.wheelDeltaX/120;i.unshift(a,c,h,e);return d.event.handle.apply(this,i)}var f=["DOMMouseScroll","mousewheel"];d.event.special.mousewheel={setup:function(){if(this.addEventListener)for(var a=
f.length;a;)this.addEventListener(f[--a],g,false);else this.onmousewheel=g},teardown:function(){if(this.removeEventListener)for(var a=f.length;a;)this.removeEventListener(f[--a],g,false);else this.onmousewheel=null}};d.fn.extend({mousewheel:function(a){return a?this.bind("mousewheel",a):this.trigger("mousewheel")},unmousewheel:function(a){return this.unbind("mousewheel",a)}})})(jQuery);
```
