## Supplementary material 1 for "Termite diet rather than geographical origin determines the microbiome composition and functional genetic structure of nests from South American and African representatives, as revealed by a multiomics approach": jquery-1.4.2.js

```
/*!
 * jQuery JavaScript Library v1.4.2
 * http://jquery.com/
 *
 * Copyright 2010, John Resig
 * Dual licensed under the MIT or GPL Version 2 licenses.
 * http://jquery.org/license
 *
 * Includes Sizzle.js
 * http://sizzlejs.com/
 * Copyright 2010, The Dojo Foundation
 * Released under the MIT, BSD, and GPL Licenses.
 *
 * Date: Sat Feb 13 22:33:48 2010 -0500
 */
(function( window, undefined ) {

// Define a local copy of jQuery
var jQuery = function( selector, context ) {
		// The jQuery object is actually just the init constructor 'enhanced'
		return new jQuery.fn.init( selector, context );
	},

	// Map over jQuery in case of overwrite
	_jQuery = window.jQuery,

	// Map over the $ in case of overwrite
	_$ = window.$,

	// Use the correct document accordingly with window argument (sandbox)
	document = window.document,

	// A central reference to the root jQuery(document)
	rootjQuery,

	// A simple way to check for HTML strings or ID strings
	// (both of which we optimize for)
	quickExpr = /^[^<]*(<[\w\W]+>)[^>]*$|^#([\w-]+)$/,

	// Is it a simple selector
	isSimple = /^.[^:#\[\.,]*$/,

	// Check if a string has a non-whitespace character in it
	rnotwhite = /\S/,

	// Used for trimming whitespace
	rtrim = /^(\s|\u00A0)+|(\s|\u00A0)+$/g,

	// Match a standalone tag
	rsingleTag = /^<(\w+)\s*\/?>(?:<\/\1>)?$/,

	// Keep a UserAgent string for use with jQuery.browser
	userAgent = navigator.userAgent,

	// For matching the engine and version of the browser
	browserMatch,
	
	// Has the ready events already been bound?
	readyBound = false,
	
	// The functions to execute on DOM ready
	readyList = [],

	// The ready event handler
	DOMContentLoaded,

	// Save a reference to some core methods
	toString = Object.prototype.toString,
	hasOwnProperty = Object.prototype.hasOwnProperty,
	push = Array.prototype.push,
	slice = Array.prototype.slice,
	indexOf = Array.prototype.indexOf;

jQuery.fn = jQuery.prototype = {
	init: function( selector, context ) {
		var match, elem, ret, doc;

		// Handle $(""), $(null), or $(undefined)
		if ( !selector ) {
			return this;
		}

		// Handle $(DOMElement)
		if ( selector.nodeType ) {
			this.context = this[0] = selector;
			this.length = 1;
			return this;
		}
		
		// The body element only exists once, optimize finding it
		if ( selector === "body" && !context ) {
			this.context = document;
			this[0] = document.body;
			this.selector = "body";
			this.length = 1;
			return this;
		}

		// Handle HTML strings
		if ( typeof selector === "string" ) {
			// Are we dealing with HTML string or an ID?
			match = quickExpr.exec( selector );

			// Verify a match, and that no context was specified for #id
			if ( match && (match[1] || !context) ) {

				// HANDLE: $(html) -> $(array)
				if ( match[1] ) {
					doc = (context ? context.ownerDocument || context : document);

					// If a single string is passed in and it's a single tag
					// just do a createElement and skip the rest
					ret = rsingleTag.exec( selector );

					if ( ret ) {
						if ( jQuery.isPlainObject( context ) ) {
							selector = [ document.createElement( ret[1] ) ];
							jQuery.fn.attr.call( selector, context, true );

						} else {
							selector = [ doc.createElement( ret[1] ) ];
						}

					} else {
						ret = buildFragment( [ match[1] ], [ doc ] );
						selector = (ret.cacheable ? ret.fragment.cloneNode(true) : ret.fragment).childNodes;
					}
					
					return jQuery.merge( this, selector );
					
				// HANDLE: $("#id")
				} else {
					elem = document.getElementById( match[2] );

					if ( elem ) {
						// Handle the case where IE and Opera return items
						// by name instead of ID
						if ( elem.id !== match[2] ) {
							return rootjQuery.find( selector );
						}

						// Otherwise, we inject the element directly into the jQuery object
						this.length = 1;
						this[0] = elem;
					}

					this.context = document;
					this.selector = selector;
					return this;
				}

			// HANDLE: $("TAG")
			} else if ( !context && /^\w+$/.test( selector ) ) {
				this.selector = selector;
				this.context = document;
				selector = document.getElementsByTagName( selector );
				return jQuery.merge( this, selector );

			// HANDLE: $(expr, $(...))
			} else if ( !context || context.jquery ) {
				return (context || rootjQuery).find( selector );

			// HANDLE: $(expr, context)
			// (which is just equivalent to: $(context).find(expr)
			} else {
				return jQuery( context ).find( selector );
			}

		// HANDLE: $(function)
		// Shortcut for document ready
		} else if ( jQuery.isFunction( selector ) ) {
			return rootjQuery.ready( selector );
		}

		if (selector.selector !== undefined) {
			this.selector = selector.selector;
			this.context = selector.context;
		}

		return jQuery.makeArray( selector, this );
	},

	// Start with an empty selector
	selector: "",

	// The current version of jQuery being used
	jquery: "1.4.2",

	// The default length of a jQuery object is 0
	length: 0,

	// The number of elements contained in the matched element set
	size: function() {
		return this.length;
	},

	toArray: function() {
		return slice.call( this, 0 );
	},

	// Get the Nth element in the matched element set OR
	// Get the whole matched element set as a clean array
	get: function( num ) {
		return num == null ?

			// Return a 'clean' array
			this.toArray() :

			// Return just the object
			( num < 0 ? this.slice(num)[ 0 ] : this[ num ] );
	},

	// Take an array of elements and push it onto the stack
	// (returning the new matched element set)
	pushStack: function( elems, name, selector ) {
		// Build a new jQuery matched element set
		var ret = jQuery();

		if ( jQuery.isArray( elems ) ) {
			push.apply( ret, elems );
		
		} else {
			jQuery.merge( ret, elems );
		}

		// Add the old object onto the stack (as a reference)
		ret.prevObject = this;

		ret.context = this.context;

		if ( name === "find" ) {
			ret.selector = this.selector + (this.selector ? " " : "") + selector;
		} else if ( name ) {
			ret.selector = this.selector + "." + name + "(" + selector + ")";
		}

		// Return the newly-formed element set
		return ret;
	},

	// Execute a callback for every element in the matched set.
	// (You can seed the arguments with an array of args, but this is
	// only used internally.)
	each: function( callback, args ) {
		return jQuery.each( this, callback, args );
	},
	
	ready: function( fn ) {
		// Attach the listeners
		jQuery.bindReady();

		// If the DOM is already ready
		if ( jQuery.isReady ) {
			// Execute the function immediately
			fn.call( document, jQuery );

		// Otherwise, remember the function for later
		} else if ( readyList ) {
			// Add the function to the wait list
			readyList.push( fn );
		}

		return this;
	},
	
	eq: function( i ) {
		return i === -1 ?
			this.slice( i ) :
			this.slice( i, +i + 1 );
	},

	first: function() {
		return this.eq( 0 );
	},

	last: function() {
		return this.eq( -1 );
	},

	slice: function() {
		return this.pushStack( slice.apply( this, arguments ),
			"slice", slice.call(arguments).join(",") );
	},

	map: function( callback ) {
		return this.pushStack( jQuery.map(this, function( elem, i ) {
			return callback.call( elem, i, elem );
		}));
	},
	
	end: function() {
		return this.prevObject || jQuery(null);
	},

	// For internal use only.
	// Behaves like an Array's method, not like a jQuery method.
	push: push,
	sort: [].sort,
	splice: [].splice
};

// Give the init function the jQuery prototype for later instantiation
jQuery.fn.init.prototype = jQuery.fn;

jQuery.extend = jQuery.fn.extend = function() {
	// copy reference to target object
	var target = arguments[0] || {}, i = 1, length = arguments.length, deep = false, options, name, src, copy;

	// Handle a deep copy situation
	if ( typeof target === "boolean" ) {
		deep = target;
		target = arguments[1] || {};
		// skip the boolean and the target
		i = 2;
	}

	// Handle case when target is a string or something (possible in deep copy)
	if ( typeof target !== "object" && !jQuery.isFunction(target) ) {
		target = {};
	}

	// extend jQuery itself if only one argument is passed
	if ( length === i ) {
		target = this;
		--i;
	}

	for ( ; i < length; i++ ) {
		// Only deal with non-null/undefined values
		if ( (options = arguments[ i ]) != null ) {
			// Extend the base object
			for ( name in options ) {
				src = target[ name ];
				copy = options[ name ];

				// Prevent never-ending loop
				if ( target === copy ) {
					continue;
				}

				// Recurse if we're merging object literal values or arrays
				if ( deep && copy && ( jQuery.isPlainObject(copy) || jQuery.isArray(copy) ) ) {
					var clone = src && ( jQuery.isPlainObject(src) || jQuery.isArray(src) ) ? src
						: jQuery.isArray(copy) ? [] : {};

					// Never move original objects, clone them
					target[ name ] = jQuery.extend( deep, clone, copy );

				// Don't bring in undefined values
				} else if ( copy !== undefined ) {
					target[ name ] = copy;
				}
			}
		}
	}

	// Return the modified object
	return target;
};

jQuery.extend({
	noConflict: function( deep ) {
		window.$ = _$;

		if ( deep ) {
			window.jQuery = _jQuery;
		}

		return jQuery;
	},
	
	// Is the DOM ready to be used? Set to true once it occurs.
	isReady: false,
	
	// Handle when the DOM is ready
	ready: function() {
		// Make sure that the DOM is not already loaded
		if ( !jQuery.isReady ) {
			// Make sure body exists, at least, in case IE gets a little overzealous (ticket #5443).
			if ( !document.body ) {
				return setTimeout( jQuery.ready, 13 );
			}

			// Remember that the DOM is ready
			jQuery.isReady = true;

			// If there are functions bound, to execute
			if ( readyList ) {
				// Execute all of them
				var fn, i = 0;
				while ( (fn = readyList[ i++ ]) ) {
					fn.call( document, jQuery );
				}

				// Reset the list of functions
				readyList = null;
			}

			// Trigger any bound ready events
			if ( jQuery.fn.triggerHandler ) {
				jQuery( document ).triggerHandler( "ready" );
			}
		}
	},
	
	bindReady: function() {
		if ( readyBound ) {
			return;
		}

		readyBound = true;

		// Catch cases where $(document).ready() is called after the
		// browser event has already occurred.
		if ( document.readyState === "complete" ) {
			return jQuery.ready();
		}

		// Mozilla, Opera and webkit nightlies currently support this event
		if ( document.addEventListener ) {
			// Use the handy event callback
			document.addEventListener( "DOMContentLoaded", DOMContentLoaded, false );
			
			// A fallback to window.onload, that will always work
			window.addEventListener( "load", jQuery.ready, false );

		// If IE event model is used
		} else if ( document.attachEvent ) {
			// ensure firing before onload,
			// maybe late but safe also for iframes
			document.attachEvent("onreadystatechange", DOMContentLoaded);
			
			// A fallback to window.onload, that will always work
			window.attachEvent( "onload", jQuery.ready );

			// If IE and not a frame
			// continually check to see if the document is ready
			var toplevel = false;

			try {
				toplevel = window.frameElement == null;
			} catch(e) {}

			if ( document.documentElement.doScroll && toplevel ) {
				doScrollCheck();
			}
		}
	},

	// See test/unit/core.js for details concerning isFunction.
	// Since version 1.3, DOM methods and functions like alert
	// aren't supported. They return false on IE (#2968).
	isFunction: function( obj ) {
		return toString.call(obj) === "[object Function]";
	},

	isArray: function( obj ) {
		return toString.call(obj) === "[object Array]";
	},

	isPlainObject: function( obj ) {
		// Must be an Object.
		// Because of IE, we also have to check the presence of the constructor property.
		// Make sure that DOM nodes and window objects don't pass through, as well
		if ( !obj || toString.call(obj) !== "[object Object]" || obj.nodeType || obj.setInterval ) {
			return false;
		}
		
		// Not own constructor property must be Object
		if ( obj.constructor
			&& !hasOwnProperty.call(obj, "constructor")
			&& !hasOwnProperty.call(obj.constructor.prototype, "isPrototypeOf") ) {
			return false;
		}
		
		// Own properties are enumerated firstly, so to speed up,
		// if last one is own, then all properties are own.
	
		var key;
		for ( key in obj ) {}
		
		return key === undefined || hasOwnProperty.call( obj, key );
	},

	isEmptyObject: function( obj ) {
		for ( var name in obj ) {
			return false;
		}
		return true;
	},
	
	error: function( msg ) {
		throw msg;
	},
	
	parseJSON: function( data ) {
		if ( typeof data !== "string" || !data ) {
			return null;
		}

		// Make sure leading/trailing whitespace is removed (IE can't handle it)
		data = jQuery.trim( data );
		
		// Make sure the incoming data is actual JSON
		// Logic borrowed from http://json.org/json2.js
		if ( /^[\],:{}\s]*$/.test(data.replace(/\\(?:["\\\/bfnrt]|u[0-9a-fA-F]{4})/g, "@")
			.replace(/"[^"\\\n\r]*"|true|false|null|-?\d+(?:\.\d*)?(?:[eE][+\-]?\d+)?/g, "]")
			.replace(/(?:^|:|,)(?:\s*\[)+/g, "")) ) {

			// Try to use the native JSON parser first
			return window.JSON && window.JSON.parse ?
				window.JSON.parse( data ) :
				(new Function("return " + data))();

		} else {
			jQuery.error( "Invalid JSON: " + data );
		}
	},

	noop: function() {},

	// Evalulates a script in a global context
	globalEval: function( data ) {
		if ( data && rnotwhite.test(data) ) {
			// Inspired by code by Andrea Giammarchi
			// http://webreflection.blogspot.com/2007/08/global-scope-evaluation-and-dom.html
			var head = document.getElementsByTagName("head")[0] || document.documentElement,
				script = document.createElement("script");

			script.type = "text/javascript";

			if ( jQuery.support.scriptEval ) {
				script.appendChild( document.createTextNode( data ) );
			} else {
				script.text = data;
			}

			// Use insertBefore instead of appendChild to circumvent an IE6 bug.
			// This arises when a base node is used (#2709).
			head.insertBefore( script, head.firstChild );
			head.removeChild( script );
		}
	},

	nodeName: function( elem, name ) {
		return elem.nodeName && elem.nodeName.toUpperCase() === name.toUpperCase();
	},

	// args is for internal usage only
	each: function( object, callback, args ) {
		var name, i = 0,
			length = object.length,
			isObj = length === undefined || jQuery.isFunction(object);

		if ( args ) {
			if ( isObj ) {
				for ( name in object ) {
					if ( callback.apply( object[ name ], args ) === false ) {
						break;
					}
				}
			} else {
				for ( ; i < length; ) {
					if ( callback.apply( object[ i++ ], args ) === false ) {
						break;
					}
				}
			}

		// A special, fast, case for the most common use of each
		} else {
			if ( isObj ) {
				for ( name in object ) {
					if ( callback.call( object[ name ], name, object[ name ] ) === false ) {
						break;
					}
				}
			} else {
				for ( var value = object[0];
					i < length && callback.call( value, i, value ) !== false; value = object[++i] ) {}
			}
		}

		return object;
	},

	trim: function( text ) {
		return (text || "").replace( rtrim, "" );
	},

	// results is for internal usage only
	makeArray: function( array, results ) {
		var ret = results || [];

		if ( array != null ) {
			// The window, strings (and functions) also have 'length'
			// The extra typeof function check is to prevent crashes
			// in Safari 2 (See: #3039)
			if ( array.length == null || typeof array === "string" || jQuery.isFunction(array) || (typeof array !== "function" && array.setInterval) ) {
				push.call( ret, array );
			} else {
				jQuery.merge( ret, array );
			}
		}

		return ret;
	},

	inArray: function( elem, array ) {
		if ( array.indexOf ) {
			return array.indexOf( elem );
		}

		for ( var i = 0, length = array.length; i < length; i++ ) {
			if ( array[ i ] === elem ) {
				return i;
			}
		}

		return -1;
	},

	merge: function( first, second ) {
		var i = first.length, j = 0;

		if ( typeof second.length === "number" ) {
			for ( var l = second.length; j < l; j++ ) {
				first[ i++ ] = second[ j ];
			}
		
		} else {
			while ( second[j] !== undefined ) {
				first[ i++ ] = second[ j++ ];
			}
		}

		first.length = i;

		return first;
	},

	grep: function( elems, callback, inv ) {
		var ret = [];

		// Go through the array, only saving the items
		// that pass the validator function
		for ( var i = 0, length = elems.length; i < length; i++ ) {
			if ( !inv !== !callback( elems[ i ], i ) ) {
				ret.push( elems[ i ] );
			}
		}

		return ret;
	},

	// arg is for internal usage only
	map: function( elems, callback, arg ) {
		var ret = [], value;

		// Go through the array, translating each of the items to their
		// new value (or values).
		for ( var i = 0, length = elems.length; i < length; i++ ) {
			value = callback( elems[ i ], i, arg );

			if ( value != null ) {
				ret[ ret.length ] = value;
			}
		}

		return ret.concat.apply( [], ret );
	},

	// A global GUID counter for objects
	guid: 1,

	proxy: function( fn, proxy, thisObject ) {
		if ( arguments.length === 2 ) {
			if ( typeof proxy === "string" ) {
				thisObject = fn;
				fn = thisObject[ proxy ];
				proxy = undefined;

			} else if ( proxy && !jQuery.isFunction( proxy ) ) {
				thisObject = proxy;
				proxy = undefined;
			}
		}

		if ( !proxy && fn ) {
			proxy = function() {
				return fn.apply( thisObject || this, arguments );
			};
		}

		// Set the guid of unique handler to the same of original handler, so it can be removed
		if ( fn ) {
			proxy.guid = fn.guid = fn.guid || proxy.guid || jQuery.guid++;
		}

		// So proxy can be declared as an argument
		return proxy;
	},

	// Use of jQuery.browser is frowned upon.
	// More details: http://docs.jquery.com/Utilities/jQuery.browser
	uaMatch: function( ua ) {
		ua = ua.toLowerCase();

		var match = /(webkit)[ \/]([\w.]+)/.exec( ua ) ||
			/(opera)(?:.*version)?[ \/]([\w.]+)/.exec( ua ) ||
			/(msie) ([\w.]+)/.exec( ua ) ||
			!/compatible/.test( ua ) && /(mozilla)(?:.*? rv:([\w.]+))?/.exec( ua ) ||
		  	[];

		return { browser: match[1] || "", version: match[2] || "0" };
	},

	browser: {}
});

browserMatch = jQuery.uaMatch( userAgent );
if ( browserMatch.browser ) {
	jQuery.browser[ browserMatch.browser ] = true;
	jQuery.browser.version = browserMatch.version;
}

// Deprecated, use jQuery.browser.webkit instead
if ( jQuery.browser.webkit ) {
	jQuery.browser.safari = true;
}

if ( indexOf ) {
	jQuery.inArray = function( elem, array ) {
		return indexOf.call( array, elem );
	};
}

// All jQuery objects should point back to these
rootjQuery = jQuery(document);

// Cleanup functions for the document ready method
if ( document.addEventListener ) {
	DOMContentLoaded = function() {
		document.removeEventListener( "DOMContentLoaded", DOMContentLoaded, false );
		jQuery.ready();
	};

} else if ( document.attachEvent ) {
	DOMContentLoaded = function() {
		// Make sure body exists, at least, in case IE gets a little overzealous (ticket #5443).
		if ( document.readyState === "complete" ) {
			document.detachEvent( "onreadystatechange", DOMContentLoaded );
			jQuery.ready();
		}
	};
}

// The DOM ready check for Internet Explorer
function doScrollCheck() {
	if ( jQuery.isReady ) {
		return;
	}

	try {
		// If IE is used, use the trick by Diego Perini
		// http://javascript.nwbox.com/IEContentLoaded/
		document.documentElement.doScroll("left");
	} catch( error ) {
		setTimeout( doScrollCheck, 1 );
		return;
	}

	// and execute any waiting functions
	jQuery.ready();
}

function evalScript( i, elem ) {
	if ( elem.src ) {
		jQuery.ajax({
			url: elem.src,
			async: false,
			dataType: "script"
		});
	} else {
		jQuery.globalEval( elem.text || elem.textContent || elem.innerHTML || "" );
	}

	if ( elem.parentNode ) {
		elem.parentNode.removeChild( elem );
	}
}

// Mutifunctional method to get and set values to a collection
// The value/s can be optionally by executed if its a function
function access( elems, key, value, exec, fn, pass ) {
	var length = elems.length;
	
	// Setting many attributes
	if ( typeof key === "object" ) {
		for ( var k in key ) {
			access( elems, k, key[k], exec, fn, value );
		}
		return elems;
	}
	
	// Setting one attribute
	if ( value !== undefined ) {
		// Optionally, function values get executed if exec is true
		exec = !pass && exec && jQuery.isFunction(value);
		
		for ( var i = 0; i < length; i++ ) {
			fn( elems[i], key, exec ? value.call( elems[i], i, fn( elems[i], key ) ) : value, pass );
		}
		
		return elems;
	}
	
	// Getting an attribute
	return length ? fn( elems[0], key ) : undefined;
}

function now() {
	return (new Date).getTime();
}
(function() {

	jQuery.support = {};

	var root = document.documentElement,
		script = document.createElement("script"),
		div = document.createElement("div"),
		id = "script" + now();

	div.style.display = "none";
	div.innerHTML = "   <link/><table></table><a href='/a' style='color:red;float:left;opacity:.55;'>a</a><input type='checkbox'/>";

	var all = div.getElementsByTagName("*"),
		a = div.getElementsByTagName("a")[0];

	// Can't get basic test support
	if ( !all || !all.length || !a ) {
		return;
	}

	jQuery.support = {
		// IE strips leading whitespace when .innerHTML is used
		leadingWhitespace: div.firstChild.nodeType === 3,

		// Make sure that tbody elements aren't automatically inserted
		// IE will insert them into empty tables
		tbody: !div.getElementsByTagName("tbody").length,

		// Make sure that link elements get serialized correctly by innerHTML
		// This requires a wrapper element in IE
		htmlSerialize: !!div.getElementsByTagName("link").length,

		// Get the style information from getAttribute
		// (IE uses .cssText insted)
		style: /red/.test( a.getAttribute("style") ),

		// Make sure that URLs aren't manipulated
		// (IE normalizes it by default)
		hrefNormalized: a.getAttribute("href") === "/a",

		// Make sure that element opacity exists
		// (IE uses filter instead)
		// Use a regex to work around a WebKit issue. See #5145
		opacity: /^0.55$/.test( a.style.opacity ),

		// Verify style float existence
		// (IE uses styleFloat instead of cssFloat)
		cssFloat: !!a.style.cssFloat,

		// Make sure that if no value is specified for a checkbox
		// that it defaults to "on".
		// (WebKit defaults to "" instead)
		checkOn: div.getElementsByTagName("input")[0].value === "on",

		// Make sure that a selected-by-default option has a working selected property.
		// (WebKit defaults to false instead of true, IE too, if it's in an optgroup)
		optSelected: document.createElement("select").appendChild( document.createElement("option") ).selected,

		parentNode: div.removeChild( div.appendChild( document.createElement("div") ) ).parentNode === null,

		// Will be defined later
		deleteExpando: true,
		checkClone: false,
		scriptEval: false,
		noCloneEvent: true,
		boxModel: null
	};

	script.type = "text/javascript";
	try {
		script.appendChild( document.createTextNode( "window." + id + "=1;" ) );
	} catch(e) {}

	root.insertBefore( script, root.firstChild );

	// Make sure that the execution of code works by injecting a script
	// tag with appendChild/createTextNode
	// (IE doesn't support this, fails, and uses .text instead)
	if ( window[ id ] ) {
		jQuery.support.scriptEval = true;
		delete window[ id ];
	}

	// Test to see if it's possible to delete an expando from an element
	// Fails in Internet Explorer
	try {
		delete script.test;
	
	} catch(e) {
		jQuery.support.deleteExpando = false;
	}

	root.removeChild( script );

	if ( div.attachEvent && div.fireEvent ) {
		div.attachEvent("onclick", function click() {
			// Cloning a node shouldn't copy over any
			// bound event handlers (IE does this)
			jQuery.support.noCloneEvent = false;
			div.detachEvent("onclick", click);
		});
		div.cloneNode(true).fireEvent("onclick");
	}

	div = document.createElement("div");
	div.innerHTML = "<input type='radio' name='radiotest' checked='checked'/>";

	var fragment = document.createDocumentFragment();
	fragment.appendChild( div.firstChild );

	// WebKit doesn't clone checked state correctly in fragments
	jQuery.support.checkClone = fragment.cloneNode(true).cloneNode(true).lastChild.checked;

	// Figure out if the W3C box model works as expected
	// document.body must exist before we can do this
	jQuery(function() {
		var div = document.createElement("div");
		div.style.width = div.style.paddingLeft = "1px";

		document.body.appendChild( div );
		jQuery.boxModel = jQuery.support.boxModel = div.offsetWidth === 2;
		document.body.removeChild( div ).style.display = 'none';

		div = null;
	});

	// Technique from Juriy Zaytsev
	// http://thinkweb2.com/projects/prototype/detecting-event-support-without-browser-sniffing/
	var eventSupported = function( eventName ) { 
		var el = document.createElement("div"); 
		eventName = "on" + eventName; 

		var isSupported = (eventName in el); 
		if ( !isSupported ) { 
			el.setAttribute(eventName, "return;"); 
			isSupported = typeof el[eventName] === "function"; 
		} 
		el = null; 

		return isSupported; 
	};
	
	jQuery.support.submitBubbles = eventSupported("submit");
	jQuery.support.changeBubbles = eventSupported("change");

	// release memory in IE
	root = script = div = all = a = null;
})();

jQuery.props = {
	"for": "htmlFor",
	"class": "className",
	readonly: "readOnly",
	maxlength: "maxLength",
	cellspacing: "cellSpacing",
	rowspan: "rowSpan",
	colspan: "colSpan",
	tabindex: "tabIndex",
	usemap: "useMap",
	frameborder: "frameBorder"
};
var expando = "jQuery" + now(), uuid = 0, windowData = {};

jQuery.extend({
	cache: {},
	
	expando:expando,

	// The following elements throw uncatchable exceptions if you
	// attempt to add expando properties to them.
	noData: {
		"embed": true,
		"object": true,
		"applet": true
	},

	data: function( elem, name, data ) {
		if ( elem.nodeName && jQuery.noData[elem.nodeName.toLowerCase()] ) {
			return;
		}

		elem = elem == window ?
			windowData :
			elem;

		var id = elem[ expando ], cache = jQuery.cache, thisCache;

		if ( !id && typeof name === "string" && data === undefined ) {
			return null;
		}

		// Compute a unique ID for the element
		if ( !id ) { 
			id = ++uuid;
		}

		// Avoid generating a new cache unless none exists and we
		// want to manipulate it.
		if ( typeof name === "object" ) {
			elem[ expando ] = id;
			thisCache = cache[ id ] = jQuery.extend(true, {}, name);

		} else if ( !cache[ id ] ) {
			elem[ expando ] = id;
			cache[ id ] = {};
		}

		thisCache = cache[ id ];

		// Prevent overriding the named cache with undefined values
		if ( data !== undefined ) {
			thisCache[ name ] = data;
		}

		return typeof name === "string" ? thisCache[ name ] : thisCache;
	},

	removeData: function( elem, name ) {
		if ( elem.nodeName && jQuery.noData[elem.nodeName.toLowerCase()] ) {
			return;
		}

		elem = elem == window ?
			windowData :
			elem;

		var id = elem[ expando ], cache = jQuery.cache, thisCache = cache[ id ];

		// If we want to remove a specific section of the element's data
		if ( name ) {
			if ( thisCache ) {
				// Remove the section of cache data
				delete thisCache[ name ];

				// If we've removed all the data, remove the element's cache
				if ( jQuery.isEmptyObject(thisCache) ) {
					jQuery.removeData( elem );
				}
			}

		// Otherwise, we want to remove all of the element's data
		} else {
			if ( jQuery.support.deleteExpando ) {
				delete elem[ jQuery.expando ];

			} else if ( elem.removeAttribute ) {
				elem.removeAttribute( jQuery.expando );
			}

			// Completely remove the data cache
			delete cache[ id ];
		}
	}
});

jQuery.fn.extend({
	data: function( key, value ) {
		if ( typeof key === "undefined" && this.length ) {
			return jQuery.data( this[0] );

		} else if ( typeof key === "object" ) {
			return this.each(function() {
				jQuery.data( this, key );
			});
		}

		var parts = key.split(".");
		parts[1] = parts[1] ? "." + parts[1] : "";

		if ( value === undefined ) {
			var data = this.triggerHandler("getData" + parts[1] + "!", [parts[0]]);

			if ( data === undefined && this.length ) {
				data = jQuery.data( this[0], key );
			}
			return data === undefined && parts[1] ?
				this.data( parts[0] ) :
				data;
		} else {
			return this.trigger("setData" + parts[1] + "!", [parts[0], value]).each(function() {
				jQuery.data( this, key, value );
			});
		}
	},

	removeData: function( key ) {
		return this.each(function() {
			jQuery.removeData( this, key );
		});
	}
});
jQuery.extend({
	queue: function( elem, type, data ) {
		if ( !elem ) {
			return;
		}

		type = (type || "fx") + "queue";
		var q = jQuery.data( elem, type );

		// Speed up dequeue by getting out quickly if this is just a lookup
		if ( !data ) {
			return q || [];
		}

		if ( !q || jQuery.isArray(data) ) {
			q = jQuery.data( elem, type, jQuery.makeArray(data) );

		} else {
			q.push( data );
		}

		return q;
	},

	dequeue: function( elem, type ) {
		type = type || "fx";

		var queue = jQuery.queue( elem, type ), fn = queue.shift();

		// If the fx queue is dequeued, always remove the progress sentinel
		if ( fn === "inprogress" ) {
			fn = queue.shift();
		}

		if ( fn ) {
			// Add a progress sentinel to prevent the fx queue from being
			// automatically dequeued
			if ( type === "fx" ) {
				queue.unshift("inprogress");
			}

			fn.call(elem, function() {
				jQuery.dequeue(elem, type);
			});
		}
	}
});

jQuery.fn.extend({
	queue: function( type, data ) {
		if ( typeof type !== "string" ) {
			data = type;
			type = "fx";
		}

		if ( data === undefined ) {
			return jQuery.queue( this[0], type );
		}
		return this.each(function( i, elem ) {
			var queue = jQuery.queue( this, type, data );

			if ( type === "fx" && queue[0] !== "inprogress" ) {
				jQuery.dequeue( this, type );
			}
		});
	},
	dequeue: function( type ) {
		return this.each(function() {
			jQuery.dequeue( this, type );
		});
	},

	// Based off of the plugin by Clint Helfers, with permission.
	// http://blindsignals.com/index.php/2009/07/jquery-delay/
	delay: function( time, type ) {
		time = jQuery.fx ? jQuery.fx.speeds[time] || time : time;
		type = type || "fx";

		return this.queue( type, function() {
			var elem = this;
			setTimeout(function() {
				jQuery.dequeue( elem, type );
			}, time );
		});
	},

	clearQueue: function( type ) {
		return this.queue( type || "fx", [] );
	}
});
var rclass = /[\n\t]/g,
	rspace = /\s+/,
	rreturn = /\r/g,
	rspecialurl = /href|src|style/,
	rtype = /(button|input)/i,
	rfocusable = /(button|input|object|select|textarea)/i,
	rclickable = /^(a|area)$/i,
	rradiocheck = /radio|checkbox/;

jQuery.fn.extend({
	attr: function( name, value ) {
		return access( this, name, value, true, jQuery.attr );
	},

	removeAttr: function( name, fn ) {
		return this.each(function(){
			jQuery.attr( this, name, "" );
			if ( this.nodeType === 1 ) {
				this.removeAttribute( name );
			}
		});
	},

	addClass: function( value ) {
		if ( jQuery.isFunction(value) ) {
			return this.each(function(i) {
				var self = jQuery(this);
				self.addClass( value.call(this, i, self.attr("class")) );
			});
		}

		if ( value && typeof value === "string" ) {
			var classNames = (value || "").split( rspace );

			for ( var i = 0, l = this.length; i < l; i++ ) {
				var elem = this[i];

				if ( elem.nodeType === 1 ) {
					if ( !elem.className ) {
						elem.className = value;

					} else {
						var className = " " + elem.className + " ", setClass = elem.className;
						for ( var c = 0, cl = classNames.length; c < cl; c++ ) {
							if ( className.indexOf( " " + classNames[c] + " " ) < 0 ) {
								setClass += " " + classNames[c];
							}
						}
						elem.className = jQuery.trim( setClass );
					}
				}
			}
		}

		return this;
	},

	removeClass: function( value ) {
		if ( jQuery.isFunction(value) ) {
			return this.each(function(i) {
				var self = jQuery(this);
				self.removeClass( value.call(this, i, self.attr("class")) );
			});
		}

		if ( (value && typeof value === "string") || value === undefined ) {
			var classNames = (value || "").split(rspace);

			for ( var i = 0, l = this.length; i < l; i++ ) {
				var elem = this[i];

				if ( elem.nodeType === 1 && elem.className ) {
					if ( value ) {
						var className = (" " + elem.className + " ").replace(rclass, " ");
						for ( var c = 0, cl = classNames.length; c < cl; c++ ) {
							className = className.replace(" " + classNames[c] + " ", " ");
						}
						elem.className = jQuery.trim( className );

					} else {
						elem.className = "";
					}
				}
			}
		}

		return this;
	},

	toggleClass: function( value, stateVal ) {
		var type = typeof value, isBool = typeof stateVal === "boolean";

		if ( jQuery.isFunction( value ) ) {
			return this.each(function(i) {
				var self = jQuery(this);
				self.toggleClass( value.call(this, i, self.attr("class"), stateVal), stateVal );
			});
		}

		return this.each(function() {
			if ( type === "string" ) {
				// toggle individual class names
				var className, i = 0, self = jQuery(this),
					state = stateVal,
					classNames = value.split( rspace );

				while ( (className = classNames[ i++ ]) ) {
					// check each className given, space seperated list
					state = isBool ? state : !self.hasClass( className );
					self[ state ? "addClass" : "removeClass" ]( className );
				}

			} else if ( type === "undefined" || type === "boolean" ) {
				if ( this.className ) {
					// store className if set
					jQuery.data( this, "__className__", this.className );
				}

				// toggle whole className
				this.className = this.className || value === false ? "" : jQuery.data( this, "__className__" ) || "";
			}
		});
	},

	hasClass: function( selector ) {
		var className = " " + selector + " ";
		for ( var i = 0, l = this.length; i < l; i++ ) {
			if ( (" " + this[i].className + " ").replace(rclass, " ").indexOf( className ) > -1 ) {
				return true;
			}
		}

		return false;
	},

	val: function( value ) {
		if ( value === undefined ) {
			var elem = this[0];

			if ( elem ) {
				if ( jQuery.nodeName( elem, "option" ) ) {
					return (elem.attributes.value || {}).specified ? elem.value : elem.text;
				}

				// We need to handle select boxes special
				if ( jQuery.nodeName( elem, "select" ) ) {
					var index = elem.selectedIndex,
						values = [],
						options = elem.options,
						one = elem.type === "select-one";

					// Nothing was selected
					if ( index < 0 ) {
						return null;
					}

					// Loop through all the selected options
					for ( var i = one ? index : 0, max = one ? index + 1 : options.length; i < max; i++ ) {
						var option = options[ i ];

						if ( option.selected ) {
							// Get the specifc value for the option
							value = jQuery(option).val();

							// We don't need an array for one selects
							if ( one ) {
								return value;
							}

							// Multi-Selects return an array
							values.push( value );
						}
					}

					return values;
				}

				// Handle the case where in Webkit "" is returned instead of "on" if a value isn't specified
				if ( rradiocheck.test( elem.type ) && !jQuery.support.checkOn ) {
					return elem.getAttribute("value") === null ? "on" : elem.value;
				}
				

				// Everything else, we just grab the value
				return (elem.value || "").replace(rreturn, "");

			}

			return undefined;
		}

		var isFunction = jQuery.isFunction(value);

		return this.each(function(i) {
			var self = jQuery(this), val = value;

			if ( this.nodeType !== 1 ) {
				return;
			}

			if ( isFunction ) {
				val = value.call(this, i, self.val());
			}

			// Typecast each time if the value is a Function and the appended
			// value is therefore different each time.
			if ( typeof val === "number" ) {
				val += "";
			}

			if ( jQuery.isArray(val) && rradiocheck.test( this.type ) ) {
				this.checked = jQuery.inArray( self.val(), val ) >= 0;

			} else if ( jQuery.nodeName( this, "select" ) ) {
				var values = jQuery.makeArray(val);

				jQuery( "option", this ).each(function() {
					this.selected = jQuery.inArray( jQuery(this).val(), values ) >= 0;
				});

				if ( !values.length ) {
					this.selectedIndex = -1;
				}

			} else {
				this.value = val;
			}
		});
	}
});

jQuery.extend({
	attrFn: {
		val: true,
		css: true,
		html: true,
		text: true,
		data: true,
		width: true,
		height: true,
		offset: true
	},
		
	attr: function( elem, name, value, pass ) {
		// don't set attributes on text and comment nodes
		if ( !elem || elem.nodeType === 3 || elem.nodeType === 8 ) {
			return undefined;
		}

		if ( pass && name in jQuery.attrFn ) {
			return jQuery(elem)[name](value);
		}

		var notxml = elem.nodeType !== 1 || !jQuery.isXMLDoc( elem ),
			// Whether we are setting (or getting)
			set = value !== undefined;

		// Try to normalize/fix the name
		name = notxml && jQuery.props[ name ] || name;

		// Only do all the following if this is a node (faster for style)
		if ( elem.nodeType === 1 ) {
			// These attributes require special treatment
			var special = rspecialurl.test( name );

			// Safari mis-reports the default selected property of an option
			// Accessing the parent's selectedIndex property fixes it
			if ( name === "selected" && !jQuery.support.optSelected ) {
				var parent = elem.parentNode;
				if ( parent ) {
					parent.selectedIndex;
	
					// Make sure that it also works with optgroups, see #5701
					if ( parent.parentNode ) {
						parent.parentNode.selectedIndex;
					}
				}
			}

			// If applicable, access the attribute via the DOM 0 way
			if ( name in elem && notxml && !special ) {
				if ( set ) {
					// We can't allow the type property to be changed (since it causes problems in IE)
					if ( name === "type" && rtype.test( elem.nodeName ) && elem.parentNode ) {
						jQuery.error( "type property can't be changed" );
					}

					elem[ name ] = value;
				}

				// browsers index elements by id/name on forms, give priority to attributes.
				if ( jQuery.nodeName( elem, "form" ) && elem.getAttributeNode(name) ) {
					return elem.getAttributeNode( name ).nodeValue;
				}

				// elem.tabIndex doesn't always return the correct value when it hasn't been explicitly set
				// http://fluidproject.org/blog/2008/01/09/getting-setting-and-removing-tabindex-values-with-javascript/
				if ( name === "tabIndex" ) {
					var attributeNode = elem.getAttributeNode( "tabIndex" );

					return attributeNode && attributeNode.specified ?
						attributeNode.value :
						rfocusable.test( elem.nodeName ) || rclickable.test( elem.nodeName ) && elem.href ?
							0 :
							undefined;
				}

				return elem[ name ];
			}

			if ( !jQuery.support.style && notxml && name === "style" ) {
				if ( set ) {
					elem.style.cssText = "" + value;
				}

				return elem.style.cssText;
			}

			if ( set ) {
				// convert the value to a string (all browsers do this but IE) see #1070
				elem.setAttribute( name, "" + value );
			}

			var attr = !jQuery.support.hrefNormalized && notxml && special ?
					// Some attributes require a special call on IE
					elem.getAttribute( name, 2 ) :
					elem.getAttribute( name );

			// Non-existent attributes return null, we normalize to undefined
			return attr === null ? undefined : attr;
		}

		// elem is actually elem.style ... set the style
		// Using attr for specific style information is now deprecated. Use style instead.
		return jQuery.style( elem, name, value );
	}
});
var rnamespaces = /\.(.*)$/,
	fcleanup = function( nm ) {
		return nm.replace(/[^\w\s\.\|`]/g, function( ch ) {
			return "\\" + ch;
		});
	};

/*
 * A number of helper functions used for managing events.
 * Many of the ideas behind this code originated from
 * Dean Edwards' addEvent library.
 */
jQuery.event = {

	// Bind an event to an element
	// Original by Dean Edwards
	add: function( elem, types, handler, data ) {
		if ( elem.nodeType === 3 || elem.nodeType === 8 ) {
			return;
		}

		// For whatever reason, IE has trouble passing the window object
		// around, causing it to be cloned in the process
		if ( elem.setInterval && ( elem !== window && !elem.frameElement ) ) {
			elem = window;
		}

		var handleObjIn, handleObj;

		if ( handler.handler ) {
			handleObjIn = handler;
			handler = handleObjIn.handler;
		}

		// Make sure that the function being executed has a unique ID
		if ( !handler.guid ) {
			handler.guid = jQuery.guid++;
		}

		// Init the element's event structure
		var elemData = jQuery.data( elem );

		// If no elemData is found then we must be trying to bind to one of the
		// banned noData elements
		if ( !elemData ) {
			return;
		}

		var events = elemData.events = elemData.events || {},
			eventHandle = elemData.handle, eventHandle;

		if ( !eventHandle ) {
			elemData.handle = eventHandle = function() {
				// Handle the second event of a trigger and when
				// an event is called after a page has unloaded
				return typeof jQuery !== "undefined" && !jQuery.event.triggered ?
					jQuery.event.handle.apply( eventHandle.elem, arguments ) :
					undefined;
			};
		}

		// Add elem as a property of the handle function
		// This is to prevent a memory leak with non-native events in IE.
		eventHandle.elem = elem;

		// Handle multiple events separated by a space
		// jQuery(...).bind("mouseover mouseout", fn);
		types = types.split(" ");

		var type, i = 0, namespaces;

		while ( (type = types[ i++ ]) ) {
			handleObj = handleObjIn ?
				jQuery.extend({}, handleObjIn) :
				{ handler: handler, data: data };

			// Namespaced event handlers
			if ( type.indexOf(".") > -1 ) {
				namespaces = type.split(".");
				type = namespaces.shift();
				handleObj.namespace = namespaces.slice(0).sort().join(".");

			} else {
				namespaces = [];
				handleObj.namespace = "";
			}

			handleObj.type = type;
			handleObj.guid = handler.guid;

			// Get the current list of functions bound to this event
			var handlers = events[ type ],
				special = jQuery.event.special[ type ] || {};

			// Init the event handler queue
			if ( !handlers ) {
				handlers = events[ type ] = [];

				// Check for a special event handler
				// Only use addEventListener/attachEvent if the special
				// events handler returns false
				if ( !special.setup || special.setup.call( elem, data, namespaces, eventHandle ) === false ) {
					// Bind the global event handler to the element
					if ( elem.addEventListener ) {
						elem.addEventListener( type, eventHandle, false );

					} else if ( elem.attachEvent ) {
						elem.attachEvent( "on" + type, eventHandle );
					}
				}
			}
			
			if ( special.add ) { 
				special.add.call( elem, handleObj ); 

				if ( !handleObj.handler.guid ) {
					handleObj.handler.guid = handler.guid;
				}
			}

			// Add the function to the element's handler list
			handlers.push( handleObj );

			// Keep track of which events have been used, for global triggering
			jQuery.event.global[ type ] = true;
		}

		// Nullify elem to prevent memory leaks in IE
		elem = null;
	},

	global: {},

	// Detach an event or set of events from an element
	remove: function( elem, types, handler, pos ) {
		// don't do events on text and comment nodes
		if ( elem.nodeType === 3 || elem.nodeType === 8 ) {
			return;
		}

		var ret, type, fn, i = 0, all, namespaces, namespace, special, eventType, handleObj, origType,
			elemData = jQuery.data( elem ),
			events = elemData && elemData.events;

		if ( !elemData || !events ) {
			return;
		}

		// types is actually an event object here
		if ( types && types.type ) {
			handler = types.handler;
			types = types.type;
		}

		// Unbind all events for the element
		if ( !types || typeof types === "string" && types.charAt(0) === "." ) {
			types = types || "";

			for ( type in events ) {
				jQuery.event.remove( elem, type + types );
			}

			return;
		}

		// Handle multiple events separated by a space
		// jQuery(...).unbind("mouseover mouseout", fn);
		types = types.split(" ");

		while ( (type = types[ i++ ]) ) {
			origType = type;
			handleObj = null;
			all = type.indexOf(".") < 0;
			namespaces = [];

			if ( !all ) {
				// Namespaced event handlers
				namespaces = type.split(".");
				type = namespaces.shift();

				namespace = new RegExp("(^|\\.)" + 
					jQuery.map( namespaces.slice(0).sort(), fcleanup ).join("\\.(?:.*\\.)?") + "(\\.|$)")
			}

			eventType = events[ type ];

			if ( !eventType ) {
				continue;
			}

			if ( !handler ) {
				for ( var j = 0; j < eventType.length; j++ ) {
					handleObj = eventType[ j ];

					if ( all || namespace.test( handleObj.namespace ) ) {
						jQuery.event.remove( elem, origType, handleObj.handler, j );
						eventType.splice( j--, 1 );
					}
				}

				continue;
			}

			special = jQuery.event.special[ type ] || {};

			for ( var j = pos || 0; j < eventType.length; j++ ) {
				handleObj = eventType[ j ];

				if ( handler.guid === handleObj.guid ) {
					// remove the given handler for the given type
					if ( all || namespace.test( handleObj.namespace ) ) {
						if ( pos == null ) {
							eventType.splice( j--, 1 );
						}

						if ( special.remove ) {
							special.remove.call( elem, handleObj );
						}
					}

					if ( pos != null ) {
						break;
					}
				}
			}

			// remove generic event handler if no more handlers exist
			if ( eventType.length === 0 || pos != null && eventType.length === 1 ) {
				if ( !special.teardown || special.teardown.call( elem, namespaces ) === false ) {
					removeEvent( elem, type, elemData.handle );
				}

				ret = null;
				delete events[ type ];
			}
		}

		// Remove the expando if it's no longer used
		if ( jQuery.isEmptyObject( events ) ) {
			var handle = elemData.handle;
			if ( handle ) {
				handle.elem = null;
			}

			delete elemData.events;
			delete elemData.handle;

			if ( jQuery.isEmptyObject( elemData ) ) {
				jQuery.removeData( elem );
			}
		}
	},

	// bubbling is internal
	trigger: function( event, data, elem /*, bubbling */ ) {
		// Event object or event type
		var type = event.type || event,
			bubbling = arguments[3];

		if ( !bubbling ) {
			event = typeof event === "object" ?
				// jQuery.Event object
				event[expando] ? event :
				// Object literal
				jQuery.extend( jQuery.Event(type), event ) :
				// Just the event type (string)
				jQuery.Event(type);

			if ( type.indexOf("!") >= 0 ) {
				event.type = type = type.slice(0, -1);
				event.exclusive = true;
			}

			// Handle a global trigger
			if ( !elem ) {
				// Don't bubble custom events when global (to avoid too much overhead)
				event.stopPropagation();

				// Only trigger if we've ever bound an event for it
				if ( jQuery.event.global[ type ] ) {
					jQuery.each( jQuery.cache, function() {
						if ( this.events && this.events[type] ) {
							jQuery.event.trigger( event, data, this.handle.elem );
						}
					});
				}
			}

			// Handle triggering a single element

			// don't do events on text and comment nodes
			if ( !elem || elem.nodeType === 3 || elem.nodeType === 8 ) {
				return undefined;
			}

			// Clean up in case it is reused
			event.result = undefined;
			event.target = elem;

			// Clone the incoming data, if any
			data = jQuery.makeArray( data );
			data.unshift( event );
		}

		event.currentTarget = elem;

		// Trigger the event, it is assumed that "handle" is a function
		var handle = jQuery.data( elem, "handle" );
		if ( handle ) {
			handle.apply( elem, data );
		}

		var parent = elem.parentNode || elem.ownerDocument;

		// Trigger an inline bound script
		try {
			if ( !(elem && elem.nodeName && jQuery.noData[elem.nodeName.toLowerCase()]) ) {
				if ( elem[ "on" + type ] && elem[ "on" + type ].apply( elem, data ) === false ) {
					event.result = false;
				}
			}

		// prevent IE from throwing an error for some elements with some event types, see #3533
		} catch (e) {}

		if ( !event.isPropagationStopped() && parent ) {
			jQuery.event.trigger( event, data, parent, true );

		} else if ( !event.isDefaultPrevented() ) {
			var target = event.target, old,
				isClick = jQuery.nodeName(target, "a") && type === "click",
				special = jQuery.event.special[ type ] || {};

			if ( (!special._default || special._default.call( elem, event ) === false) && 
				!isClick && !(target && target.nodeName && jQuery.noData[target.nodeName.toLowerCase()]) ) {

				try {
					if ( target[ type ] ) {
						// Make sure that we don't accidentally re-trigger the onFOO events
						old = target[ "on" + type ];

						if ( old ) {
							target[ "on" + type ] = null;
						}

						jQuery.event.triggered = true;
						target[ type ]();
					}

				// prevent IE from throwing an error for some elements with some event types, see #3533
				} catch (e) {}

				if ( old ) {
					target[ "on" + type ] = old;
				}

				jQuery.event.triggered = false;
			}
		}
	},

	handle: function( event ) {
		var all, handlers, namespaces, namespace, events;

		event = arguments[0] = jQuery.event.fix( event || window.event );
		event.currentTarget = this;

		// Namespaced event handlers
		all = event.type.indexOf(".") < 0 && !event.exclusive;

		if ( !all ) {
			namespaces = event.type.split(".");
			event.type = namespaces.shift();
			namespace = new RegExp("(^|\\.)" + namespaces.slice(0).sort().join("\\.(?:.*\\.)?") + "(\\.|$)");
		}

		var events = jQuery.data(this, "events"), handlers = events[ event.type ];

		if ( events && handlers ) {
			// Clone the handlers to prevent manipulation
			handlers = handlers.slice(0);

			for ( var j = 0, l = handlers.length; j < l; j++ ) {
				var handleObj = handlers[ j ];

				// Filter the functions by class
				if ( all || namespace.test( handleObj.namespace ) ) {
					// Pass in a reference to the handler function itself
					// So that we can later remove it
					event.handler = handleObj.handler;
					event.data = handleObj.data;
					event.handleObj = handleObj;
	
					var ret = handleObj.handler.apply( this, arguments );

					if ( ret !== undefined ) {
						event.result = ret;
						if ( ret === false ) {
							event.preventDefault();
							event.stopPropagation();
						}
					}

					if ( event.isImmediatePropagationStopped() ) {
						break;
					}
				}
			}
		}

		return event.result;
	},

	props: "altKey attrChange attrName bubbles button cancelable charCode clientX clientY ctrlKey currentTarget data detail eventPhase fromElement handler keyCode layerX layerY metaKey newValue offsetX offsetY originalTarget pageX pageY prevValue relatedNode relatedTarget screenX screenY shiftKey srcElement target toElement view wheelDelta which".split(" "),

	fix: function( event ) {
		if ( event[ expando ] ) {
			return event;
		}

		// store a copy of the original event object
		// and "clone" to set read-only properties
		var originalEvent = event;
		event = jQuery.Event( originalEvent );

		for ( var i = this.props.length, prop; i; ) {
			prop = this.props[ --i ];
			event[ prop ] = originalEvent[ prop ];
		}

		// Fix target property, if necessary
		if ( !event.target ) {
			event.target = event.srcElement || document; // Fixes #1925 where srcElement might not be defined either
		}

		// check if target is a textnode (safari)
		if ( event.target.nodeType === 3 ) {
			event.target = event.target.parentNode;
		}

		// Add relatedTarget, if necessary
		if ( !event.relatedTarget && event.fromElement ) {
			event.relatedTarget = event.fromElement === event.target ? event.toElement : event.fromElement;
		}

		// Calculate pageX/Y if missing and clientX/Y available
		if ( event.pageX == null && event.clientX != null ) {
			var doc = document.documentElement, body = document.body;
			event.pageX = event.clientX + (doc && doc.scrollLeft || body && body.scrollLeft || 0) - (doc && doc.clientLeft || body && body.clientLeft || 0);
			event.pageY = event.clientY + (doc && doc.scrollTop  || body && body.scrollTop  || 0) - (doc && doc.clientTop  || body && body.clientTop  || 0);
		}

		// Add which for key events
		if ( !event.which && ((event.charCode || event.charCode === 0) ? event.charCode : event.keyCode) ) {
			event.which = event.charCode || event.keyCode;
		}

		// Add metaKey to non-Mac browsers (use ctrl for PC's and Meta for Macs)
		if ( !event.metaKey && event.ctrlKey ) {
			event.metaKey = event.ctrlKey;
		}

		// Add which for click: 1 === left; 2 === middle; 3 === right
		// Note: button is not normalized, so don't use it
		if ( !event.which && event.button !== undefined ) {
			event.which = (event.button & 1 ? 1 : ( event.button & 2 ? 3 : ( event.button & 4 ? 2 : 0 ) ));
		}

		return event;
	},

	// Deprecated, use jQuery.guid instead
	guid: 1E8,

	// Deprecated, use jQuery.proxy instead
	proxy: jQuery.proxy,

	special: {
		ready: {
			// Make sure the ready event is setup
			setup: jQuery.bindReady,
			teardown: jQuery.noop
		},

		live: {
			add: function( handleObj ) {
				jQuery.event.add( this, handleObj.origType, jQuery.extend({}, handleObj, {handler: liveHandler}) ); 
			},

			remove: function( handleObj ) {
				var remove = true,
					type = handleObj.origType.replace(rnamespaces, "");
				
				jQuery.each( jQuery.data(this, "events").live || [], function() {
					if ( type === this.origType.replace(rnamespaces, "") ) {
						remove = false;
						return false;
					}
				});

				if ( remove ) {
					jQuery.event.remove( this, handleObj.origType, liveHandler );
				}
			}

		},

		beforeunload: {
			setup: function( data, namespaces, eventHandle ) {
				// We only want to do this special case on windows
				if ( this.setInterval ) {
					this.onbeforeunload = eventHandle;
				}

				return false;
			},
			teardown: function( namespaces, eventHandle ) {
				if ( this.onbeforeunload === eventHandle ) {
					this.onbeforeunload = null;
				}
			}
		}
	}
};

var removeEvent = document.removeEventListener ?
	function( elem, type, handle ) {
		elem.removeEventListener( type, handle, false );
	} : 
	function( elem, type, handle ) {
		elem.detachEvent( "on" + type, handle );
	};

jQuery.Event = function( src ) {
	// Allow instantiation without the 'new' keyword
	if ( !this.preventDefault ) {
		return new jQuery.Event( src );
	}

	// Event object
	if ( src && src.type ) {
		this.originalEvent = src;
		this.type = src.type;
	// Event type
	} else {
		this.type = src;
	}

	// timeStamp is buggy for some events on Firefox(#3843)
	// So we won't rely on the native value
	this.timeStamp = now();

	// Mark it as fixed
	this[ expando ] = true;
};

function returnFalse() {
	return false;
}
function returnTrue() {
	return true;
}

// jQuery.Event is based on DOM3 Events as specified by the ECMAScript Language Binding
// http://www.w3.org/TR/2003/WD-DOM-Level-3-Events-20030331/ecma-script-binding.html
jQuery.Event.prototype = {
	preventDefault: function() {
		this.isDefaultPrevented = returnTrue;

		var e = this.originalEvent;
		if ( !e ) {
			return;
		}
		
		// if preventDefault exists run it on the original event
		if ( e.preventDefault ) {
			e.preventDefault();
		}
		// otherwise set the returnValue property of the original event to false (IE)
		e.returnValue = false;
	},
	stopPropagation: function() {
		this.isPropagationStopped = returnTrue;

		var e = this.originalEvent;
		if ( !e ) {
			return;
		}
		// if stopPropagation exists run it on the original event
		if ( e.stopPropagation ) {
			e.stopPropagation();
		}
		// otherwise set the cancelBubble property of the original event to true (IE)
		e.cancelBubble = true;
	},
	stopImmediatePropagation: function() {
		this.isImmediatePropagationStopped = returnTrue;
		this.stopPropagation();
	},
	isDefaultPrevented: returnFalse,
	isPropagationStopped: returnFalse,
	isImmediatePropagationStopped: returnFalse
};

// Checks if an event happened on an element within another element
// Used in jQuery.event.special.mouseenter and mouseleave handlers
var withinElement = function( event ) {
	// Check if mouse(over|out) are still within the same parent element
	var parent = event.relatedTarget;

	// Firefox sometimes assigns relatedTarget a XUL element
	// which we cannot access the parentNode property of
	try {
		// Traverse up the tree
		while ( parent && parent !== this ) {
			parent = parent.parentNode;
		}

		if ( parent !== this ) {
			// set the correct event type
			event.type = event.data;

			// handle event if we actually just moused on to a non sub-element
			jQuery.event.handle.apply( this, arguments );
		}

	// assuming we've left the element since we most likely mousedover a xul element
	} catch(e) { }
},

// In case of event delegation, we only need to rename the event.type,
// liveHandler will take care of the rest.
delegate = function( event ) {
	event.type = event.data;
	jQuery.event.handle.apply( this, arguments );
};

// Create mouseenter and mouseleave events
jQuery.each({
	mouseenter: "mouseover",
	mouseleave: "mouseout"
}, function( orig, fix ) {
	jQuery.event.special[ orig ] = {
		setup: function( data ) {
			jQuery.event.add( this, fix, data && data.selector ? delegate : withinElement, orig );
		},
		teardown: function( data ) {
			jQuery.event.remove( this, fix, data && data.selector ? delegate : withinElement );
		}
	};
});

// submit delegation
if ( !jQuery.support.submitBubbles ) {

	jQuery.event.special.submit = {
		setup: function( data, namespaces ) {
			if ( this.nodeName.toLowerCase() !== "form" ) {
				jQuery.event.add(this, "click.specialSubmit", function( e ) {
					var elem = e.target, type = elem.type;

					if ( (type === "submit" || type === "image") && jQuery( elem ).closest("form").length ) {
						return trigger( "submit", this, arguments );
					}
				});
	 
				jQuery.event.add(this, "keypress.specialSubmit", function( e ) {
					var elem = e.target, type = elem.type;

					if ( (type === "text" || type === "password") && jQuery( elem ).closest("form").length && e.keyCode === 13 ) {
						return trigger( "submit", this, arguments );
					}
				});

			} else {
				return false;
			}
		},

		teardown: function( namespaces ) {
			jQuery.event.remove( this, ".specialSubmit" );
		}
	};

}

// change delegation, happens here so we have bind.
if ( !jQuery.support.changeBubbles ) {

	var formElems = /textarea|input|select/i,

	changeFilters,

	getVal = function( elem ) {
		var type = elem.type, val = elem.value;

		if ( type === "radio" || type === "checkbox" ) {
			val = elem.checked;

		} else if ( type === "select-multiple" ) {
			val = elem.selectedIndex > -1 ?
				jQuery.map( elem.options, function( elem ) {
					return elem.selected;
				}).join("-") :
				"";

		} else if ( elem.nodeName.toLowerCase() === "select" ) {
			val = elem.selectedIndex;
		}

		return val;
	},

	testChange = function testChange( e ) {
		var elem = e.target, data, val;

		if ( !formElems.test( elem.nodeName ) || elem.readOnly ) {
			return;
		}

		data = jQuery.data( elem, "_change_data" );
		val = getVal(elem);

		// the current data will be also retrieved by beforeactivate
		if ( e.type !== "focusout" || elem.type !== "radio" ) {
			jQuery.data( elem, "_change_data", val );
		}
		
		if ( data === undefined || val === data ) {
			return;
		}

		if ( data != null || val ) {
			e.type = "change";
			return jQuery.event.trigger( e, arguments[1], elem );
		}
	};

	jQuery.event.special.change = {
		filters: {
			focusout: testChange, 

			click: function( e ) {
				var elem = e.target, type = elem.type;

				if ( type === "radio" || type === "checkbox" || elem.nodeName.toLowerCase() === "select" ) {
					return testChange.call( this, e );
				}
			},

			// Change has to be called before submit
			// Keydown will be called before keypress, which is used in submit-event delegation
			keydown: function( e ) {
				var elem = e.target, type = elem.type;

				if ( (e.keyCode === 13 && elem.nodeName.toLowerCase() !== "textarea") ||
					(e.keyCode === 32 && (type === "checkbox" || type === "radio")) ||
					type === "select-multiple" ) {
					return testChange.call( this, e );
				}
			},

			// Beforeactivate happens also before the previous element is blurred
			// with this event you can't trigger a change event, but you can store
			// information/focus[in] is not needed anymore
			beforeactivate: function( e ) {
				var elem = e.target;
				jQuery.data( elem, "_change_data", getVal(elem) );
			}
		},

		setup: function( data, namespaces ) {
			if ( this.type === "file" ) {
				return false;
			}

			for ( var type in changeFilters ) {
				jQuery.event.add( this, type + ".specialChange", changeFilters[type] );
			}

			return formElems.test( this.nodeName );
		},

		teardown: function( namespaces ) {
			jQuery.event.remove( this, ".specialChange" );

			return formElems.test( this.nodeName );
		}
	};

	changeFilters = jQuery.event.special.change.filters;
}

function trigger( type, elem, args ) {
	args[0].type = type;
	return jQuery.event.handle.apply( elem, args );
}

// Create "bubbling" focus and blur events
if ( document.addEventListener ) {
	jQuery.each({ focus: "focusin", blur: "focusout" }, function( orig, fix ) {
		jQuery.event.special[ fix ] = {
			setup: function() {
				this.addEventListener( orig, handler, true );
			}, 
			teardown: function() { 
				this.removeEventListener( orig, handler, true );
			}
		};

		function handler( e ) { 
			e = jQuery.event.fix( e );
			e.type = fix;
			return jQuery.event.handle.call( this, e );
		}
	});
}

jQuery.each(["bind", "one"], function( i, name ) {
	jQuery.fn[ name ] = function( type, data, fn ) {
		// Handle object literals
		if ( typeof type === "object" ) {
			for ( var key in type ) {
				this[ name ](key, data, type[key], fn);
			}
			return this;
		}
		
		if ( jQuery.isFunction( data ) ) {
			fn = data;
			data = undefined;
		}

		var handler = name === "one" ? jQuery.proxy( fn, function( event ) {
			jQuery( this ).unbind( event, handler );
			return fn.apply( this, arguments );
		}) : fn;

		if ( type === "unload" && name !== "one" ) {
			this.one( type, data, fn );

		} else {
			for ( var i = 0, l = this.length; i < l; i++ ) {
				jQuery.event.add( this[i], type, handler, data );
			}
		}

		return this;
	};
});

jQuery.fn.extend({
	unbind: function( type, fn ) {
		// Handle object literals
		if ( typeof type === "object" && !type.preventDefault ) {
			for ( var key in type ) {
				this.unbind(key, type[key]);
			}

		} else {
			for ( var i = 0, l = this.length; i < l; i++ ) {
				jQuery.event.remove( this[i], type, fn );
			}
		}

		return this;
	},
	
	delegate: function( selector, types, data, fn ) {
		return this.live( types, data, fn, selector );
	},
	
	undelegate: function( selector, types, fn ) {
		if ( arguments.length === 0 ) {
				return this.unbind( "live" );
		
		} else {
			return this.die( types, null, fn, selector );
		}
	},
	
	trigger: function( type, data ) {
		return this.each(function() {
			jQuery.event.trigger( type, data, this );
		});
	},

	triggerHandler: function( type, data ) {
		if ( this[0] ) {
			var event = jQuery.Event( type );
			event.preventDefault();
			event.stopPropagation();
			jQuery.event.trigger( event, data, this[0] );
			return event.result;
		}
	},

	toggle: function( fn ) {
		// Save reference to arguments for access in closure
		var args = arguments, i = 1;

		// link all the functions, so any of them can unbind this click handler
		while ( i < args.length ) {
			jQuery.proxy( fn, args[ i++ ] );
		}

		return this.click( jQuery.proxy( fn, function( event ) {
			// Figure out which function to execute
			var lastToggle = ( jQuery.data( this, "lastToggle" + fn.guid ) || 0 ) % i;
			jQuery.data( this, "lastToggle" + fn.guid, lastToggle + 1 );

			// Make sure that clicks stop
			event.preventDefault();

			// and execute the function
			return args[ lastToggle ].apply( this, arguments ) || false;
		}));
	},

	hover: function( fnOver, fnOut ) {
		return this.mouseenter( fnOver ).mouseleave( fnOut || fnOver );
	}
});

var liveMap = {
	focus: "focusin",
	blur: "focusout",
	mouseenter: "mouseover",
	mouseleave: "mouseout"
};

jQuery.each(["live", "die"], function( i, name ) {
	jQuery.fn[ name ] = function( types, data, fn, origSelector /* Internal Use Only */ ) {
		var type, i = 0, match, namespaces, preType,
			selector = origSelector || this.selector,
			context = origSelector ? this : jQuery( this.context );

		if ( jQuery.isFunction( data ) ) {
			fn = data;
			data = undefined;
		}

		types = (types || "").split(" ");

		while ( (type = types[ i++ ]) != null ) {
			match = rnamespaces.exec( type );
			namespaces = "";

			if ( match )  {
				namespaces = match[0];
				type = type.replace( rnamespaces, "" );
			}

			if ( type === "hover" ) {
				types.push( "mouseenter" + namespaces, "mouseleave" + namespaces );
				continue;
			}

			preType = type;

			if ( type === "focus" || type === "blur" ) {
				types.push( liveMap[ type ] + namespaces );
				type = type + namespaces;

			} else {
				type = (liveMap[ type ] || type) + namespaces;
			}

			if ( name === "live" ) {
				// bind live handler
				context.each(function(){
					jQuery.event.add( this, liveConvert( type, selector ),
						{ data: data, selector: selector, handler: fn, origType: type, origHandler: fn, preType: preType } );
				});

			} else {
				// unbind live handler
				context.unbind( liveConvert( type, selector ), fn );
			}
		}
		
		return this;
	}
});

function liveHandler( event ) {
	var stop, elems = [], selectors = [], args = arguments,
		related, match, handleObj, elem, j, i, l, data,
		events = jQuery.data( this, "events" );

	// Make sure we avoid non-left-click bubbling in Firefox (#3861)
	if ( event.liveFired === this || !events || !events.live || event.button && event.type === "click" ) {
		return;
	}

	event.liveFired = this;

	var live = events.live.slice(0);

	for ( j = 0; j < live.length; j++ ) {
		handleObj = live[j];

		if ( handleObj.origType.replace( rnamespaces, "" ) === event.type ) {
			selectors.push( handleObj.selector );

		} else {
			live.splice( j--, 1 );
		}
	}

	match = jQuery( event.target ).closest( selectors, event.currentTarget );

	for ( i = 0, l = match.length; i < l; i++ ) {
		for ( j = 0; j < live.length; j++ ) {
			handleObj = live[j];

			if ( match[i].selector === handleObj.selector ) {
				elem = match[i].elem;
				related = null;

				// Those two events require additional checking
				if ( handleObj.preType === "mouseenter" || handleObj.preType === "mouseleave" ) {
					related = jQuery( event.relatedTarget ).closest( handleObj.selector )[0];
				}

				if ( !related || related !== elem ) {
					elems.push({ elem: elem, handleObj: handleObj });
				}
			}
		}
	}

	for ( i = 0, l = elems.length; i < l; i++ ) {
		match = elems[i];
		event.currentTarget = match.elem;
		event.data = match.handleObj.data;
		event.handleObj = match.handleObj;

		if ( match.handleObj.origHandler.apply( match.elem, args ) === false ) {
			stop = false;
			break;
		}
	}

	return stop;
}

function liveConvert( type, selector ) {
	return "live." + (type && type !== "*" ? type + "." : "") + selector.replace(/\./g, "`").replace(/ /g, "&");
}

jQuery.each( ("blur focus focusin focusout load resize scroll unload click dblclick " +
	"mousedown mouseup mousemove mouseover mouseout mouseenter mouseleave " +
	"change select submit keydown keypress keyup error").split(" "), function( i, name ) {

	// Handle event binding
	jQuery.fn[ name ] = function( fn ) {
		return fn ? this.bind( name, fn ) : this.trigger( name );
	};

	if ( jQuery.attrFn ) {
		jQuery.attrFn[ name ] = true;
	}
});

// Prevent memory leaks in IE
// Window isn't included so as not to unbind existing unload events
// More info:
//  - http://isaacschlueter.com/2006/10/msie-memory-leaks/
if ( window.attachEvent && !window.addEventListener ) {
	window.attachEvent("onunload", function() {
		for ( var id in jQuery.cache ) {
			if ( jQuery.cache[ id ].handle ) {
				// Try/Catch is to handle iframes being unloaded, see #4280
				try {
					jQuery.event.remove( jQuery.cache[ id ].handle.elem );
				} catch(e) {}
			}
		}
	});
}
/*!
 * Sizzle CSS Selector Engine - v1.0
 *  Copyright 2009, The Dojo Foundation
 *  Released under the MIT, BSD, and GPL Licenses.
 *  More information: http://sizzlejs.com/
 */
(function(){

var chunker = /((?:\((?:\([^()]+\)|[^()]+)+\)|\[(?:\[[^[\]]*\]|['"][^'"]*['"]|[^[\]'"]+)+\]|\\.|[^ >+~,(\[\\]+)+|[>+~])(\s*,\s*)?((?:.|\r|\n)*)/g,
	done = 0,
	toString = Object.prototype.toString,
	hasDuplicate = false,
	baseHasDuplicate = true;

// Here we check if the JavaScript engine is using some sort of
// optimization where it does not always call our comparision
// function. If that is the case, discard the hasDuplicate value.
//   Thus far that includes Google Chrome.
[0, 0].sort(function(){
	baseHasDuplicate = false;
	return 0;
});

var Sizzle = function(selector, context, results, seed) {
	results = results || [];
	var origContext = context = context || document;

	if ( context.nodeType !== 1 && context.nodeType !== 9 ) {
		return [];
	}
	
	if ( !selector || typeof selector !== "string" ) {
		return results;
	}

	var parts = [], m, set, checkSet, extra, prune = true, contextXML = isXML(context),
		soFar = selector;
	
	// Reset the position of the chunker regexp (start from head)
	while ( (chunker.exec(""), m = chunker.exec(soFar)) !== null ) {
		soFar = m[3];
		
		parts.push( m[1] );
		
		if ( m[2] ) {
			extra = m[3];
			break;
		}
	}

	if ( parts.length > 1 && origPOS.exec( selector ) ) {
		if ( parts.length === 2 && Expr.relative[ parts[0] ] ) {
			set = posProcess( parts[0] + parts[1], context );
		} else {
			set = Expr.relative[ parts[0] ] ?
				[ context ] :
				Sizzle( parts.shift(), context );

			while ( parts.length ) {
				selector = parts.shift();

				if ( Expr.relative[ selector ] ) {
					selector += parts.shift();
				}
				
				set = posProcess( selector, set );
			}
		}
	} else {
		// Take a shortcut and set the context if the root selector is an ID
		// (but not if it'll be faster if the inner selector is an ID)
		if ( !seed && parts.length > 1 && context.nodeType === 9 && !contextXML &&
				Expr.match.ID.test(parts[0]) && !Expr.match.ID.test(parts[parts.length - 1]) ) {
			var ret = Sizzle.find( parts.shift(), context, contextXML );
			context = ret.expr ? Sizzle.filter( ret.expr, ret.set )[0] : ret.set[0];
		}

		if ( context ) {
			var ret = seed ?
				{ expr: parts.pop(), set: makeArray(seed) } :
				Sizzle.find( parts.pop(), parts.length === 1 && (parts[0] === "~" || parts[0] === "+") && context.parentNode ? context.parentNode : context, contextXML );
			set = ret.expr ? Sizzle.filter( ret.expr, ret.set ) : ret.set;

			if ( parts.length > 0 ) {
				checkSet = makeArray(set);
			} else {
				prune = false;
			}

			while ( parts.length ) {
				var cur = parts.pop(), pop = cur;

				if ( !Expr.relative[ cur ] ) {
					cur = "";
				} else {
					pop = parts.pop();
				}

				if ( pop == null ) {
					pop = context;
				}

				Expr.relative[ cur ]( checkSet, pop, contextXML );
			}
		} else {
			checkSet = parts = [];
		}
	}

	if ( !checkSet ) {
		checkSet = set;
	}

	if ( !checkSet ) {
		Sizzle.error( cur || selector );
	}

	if ( toString.call(checkSet) === "[object Array]" ) {
		if ( !prune ) {
			results.push.apply( results, checkSet );
		} else if ( context && context.nodeType === 1 ) {
			for ( var i = 0; checkSet[i] != null; i++ ) {
				if ( checkSet[i] && (checkSet[i] === true || checkSet[i].nodeType === 1 && contains(context, checkSet[i])) ) {
					results.push( set[i] );
				}
			}
		} else {
			for ( var i = 0; checkSet[i] != null; i++ ) {
				if ( checkSet[i] && checkSet[i].nodeType === 1 ) {
					results.push( set[i] );
				}
			}
		}
	} else {
		makeArray( checkSet, results );
	}

	if ( extra ) {
		Sizzle( extra, origContext, results, seed );
		Sizzle.uniqueSort( results );
	}

	return results;
};

Sizzle.uniqueSort = function(results){
	if ( sortOrder ) {
		hasDuplicate = baseHasDuplicate;
		results.sort(sortOrder);

		if ( hasDuplicate ) {
			for ( var i = 1; i < results.length; i++ ) {
				if ( results[i] === results[i-1] ) {
					results.splice(i--, 1);
				}
			}
		}
	}

	return results;
};

Sizzle.matches = function(expr, set){
	return Sizzle(expr, null, null, set);
};

Sizzle.find = function(expr, context, isXML){
	var set, match;

	if ( !expr ) {
		return [];
	}

	for ( var i = 0, l = Expr.order.length; i < l; i++ ) {
		var type = Expr.order[i], match;
		
		if ( (match = Expr.leftMatch[ type ].exec( expr )) ) {
			var left = match[1];
			match.splice(1,1);

			if ( left.substr( left.length - 1 ) !== "\\" ) {
				match[1] = (match[1] || "").replace(/\\/g, "");
				set = Expr.find[ type ]( match, context, isXML );
				if ( set != null ) {
					expr = expr.replace( Expr.match[ type ], "" );
					break;
				}
			}
		}
	}

	if ( !set ) {
		set = context.getElementsByTagName("*");
	}

	return {set: set, expr: expr};
};

Sizzle.filter = function(expr, set, inplace, not){
	var old = expr, result = [], curLoop = set, match, anyFound,
		isXMLFilter = set && set[0] && isXML(set[0]);

	while ( expr && set.length ) {
		for ( var type in Expr.filter ) {
			if ( (match = Expr.leftMatch[ type ].exec( expr )) != null && match[2] ) {
				var filter = Expr.filter[ type ], found, item, left = match[1];
				anyFound = false;

				match.splice(1,1);

				if ( left.substr( left.length - 1 ) === "\\" ) {
					continue;
				}

				if ( curLoop === result ) {
					result = [];
				}

				if ( Expr.preFilter[ type ] ) {
					match = Expr.preFilter[ type ]( match, curLoop, inplace, result, not, isXMLFilter );

					if ( !match ) {
						anyFound = found = true;
					} else if ( match === true ) {
						continue;
					}
				}

				if ( match ) {
					for ( var i = 0; (item = curLoop[i]) != null; i++ ) {
						if ( item ) {
							found = filter( item, match, i, curLoop );
							var pass = not ^ !!found;

							if ( inplace && found != null ) {
								if ( pass ) {
									anyFound = true;
								} else {
									curLoop[i] = false;
								}
							} else if ( pass ) {
								result.push( item );
								anyFound = true;
							}
						}
					}
				}

				if ( found !== undefined ) {
					if ( !inplace ) {
						curLoop = result;
					}

					expr = expr.replace( Expr.match[ type ], "" );

					if ( !anyFound ) {
						return [];
					}

					break;
				}
			}
		}

		// Improper expression
		if ( expr === old ) {
			if ( anyFound == null ) {
				Sizzle.error( expr );
			} else {
				break;
			}
		}

		old = expr;
	}

	return curLoop;
};

Sizzle.error = function( msg ) {
	throw "Syntax error, unrecognized expression: " + msg;
};

var Expr = Sizzle.selectors = {
	order: [ "ID", "NAME", "TAG" ],
	match: {
		ID: /#((?:[\w\u00c0-\uFFFF-]|\\.)+)/,
		CLASS: /\.((?:[\w\u00c0-\uFFFF-]|\\.)+)/,
		NAME: /\[name=['"]*((?:[\w\u00c0-\uFFFF-]|\\.)+)['"]*\]/,
		ATTR: /\[\s*((?:[\w\u00c0-\uFFFF-]|\\.)+)\s*(?:(\S?=)\s*(['"]*)(.*?)\3|)\s*\]/,
		TAG: /^((?:[\w\u00c0-\uFFFF\*-]|\\.)+)/,
		CHILD: /:(only|nth|last|first)-child(?:\((even|odd|[\dn+-]*)\))?/,
		POS: /:(nth|eq|gt|lt|first|last|even|odd)(?:\((\d*)\))?(?=[^-]|$)/,
		PSEUDO: /:((?:[\w\u00c0-\uFFFF-]|\\.)+)(?:\((['"]?)((?:\([^\)]+\)|[^\(\)]*)+)\2\))?/
	},
	leftMatch: {},
	attrMap: {
		"class": "className",
		"for": "htmlFor"
	},
	attrHandle: {
		href: function(elem){
			return elem.getAttribute("href");
		}
	},
	relative: {
		"+": function(checkSet, part){
			var isPartStr = typeof part === "string",
				isTag = isPartStr && !/\W/.test(part),
				isPartStrNotTag = isPartStr && !isTag;

			if ( isTag ) {
				part = part.toLowerCase();
			}

			for ( var i = 0, l = checkSet.length, elem; i < l; i++ ) {
				if ( (elem = checkSet[i]) ) {
					while ( (elem = elem.previousSibling) && elem.nodeType !== 1 ) {}

					checkSet[i] = isPartStrNotTag || elem && elem.nodeName.toLowerCase() === part ?
						elem || false :
						elem === part;
				}
			}

			if ( isPartStrNotTag ) {
				Sizzle.filter( part, checkSet, true );
			}
		},
		">": function(checkSet, part){
			var isPartStr = typeof part === "string";

			if ( isPartStr && !/\W/.test(part) ) {
				part = part.toLowerCase();

				for ( var i = 0, l = checkSet.length; i < l; i++ ) {
					var elem = checkSet[i];
					if ( elem ) {
						var parent = elem.parentNode;
						checkSet[i] = parent.nodeName.toLowerCase() === part ? parent : false;
					}
				}
			} else {
				for ( var i = 0, l = checkSet.length; i < l; i++ ) {
					var elem = checkSet[i];
					if ( elem ) {
						checkSet[i] = isPartStr ?
							elem.parentNode :
							elem.parentNode === part;
					}
				}

				if ( isPartStr ) {
					Sizzle.filter( part, checkSet, true );
				}
			}
		},
		"": function(checkSet, part, isXML){
			var doneName = done++, checkFn = dirCheck;

			if ( typeof part === "string" && !/\W/.test(part) ) {
				var nodeCheck = part = part.toLowerCase();
				checkFn = dirNodeCheck;
			}

			checkFn("parentNode", part, doneName, checkSet, nodeCheck, isXML);
		},
		"~": function(checkSet, part, isXML){
			var doneName = done++, checkFn = dirCheck;

			if ( typeof part === "string" && !/\W/.test(part) ) {
				var nodeCheck = part = part.toLowerCase();
				checkFn = dirNodeCheck;
			}

			checkFn("previousSibling", part, doneName, checkSet, nodeCheck, isXML);
		}
	},
	find: {
		ID: function(match, context, isXML){
			if ( typeof context.getElementById !== "undefined" && !isXML ) {
				var m = context.getElementById(match[1]);
				return m ? [m] : [];
			}
		},
		NAME: function(match, context){
			if ( typeof context.getElementsByName !== "undefined" ) {
				var ret = [], results = context.getElementsByName(match[1]);

				for ( var i = 0, l = results.length; i < l; i++ ) {
					if ( results[i].getAttribute("name") === match[1] ) {
						ret.push( results[i] );
					}
				}

				return ret.length === 0 ? null : ret;
			}
		},
		TAG: function(match, context){
			return context.getElementsByTagName(match[1]);
		}
	},
	preFilter: {
		CLASS: function(match, curLoop, inplace, result, not, isXML){
			match = " " + match[1].replace(/\\/g, "") + " ";

			if ( isXML ) {
				return match;
			}

			for ( var i = 0, elem; (elem = curLoop[i]) != null; i++ ) {
				if ( elem ) {
					if ( not ^ (elem.className && (" " + elem.className + " ").replace(/[\t\n]/g, " ").indexOf(match) >= 0) ) {
						if ( !inplace ) {
							result.push( elem );
						}
					} else if ( inplace ) {
						curLoop[i] = false;
					}
				}
			}

			return false;
		},
		ID: function(match){
			return match[1].replace(/\\/g, "");
		},
		TAG: function(match, curLoop){
			return match[1].toLowerCase();
		},
		CHILD: function(match){
			if ( match[1] === "nth" ) {
				// parse equations like 'even', 'odd', '5', '2n', '3n+2', '4n-1', '-n+6'
				var test = /(-?)(\d*)n((?:\+|-)?\d*)/.exec(
					match[2] === "even" && "2n" || match[2] === "odd" && "2n+1" ||
					!/\D/.test( match[2] ) && "0n+" + match[2] || match[2]);

				// calculate the numbers (first)n+(last) including if they are negative
				match[2] = (test[1] + (test[2] || 1)) - 0;
				match[3] = test[3] - 0;
			}

			// TODO: Move to normal caching system
			match[0] = done++;

			return match;
		},
		ATTR: function(match, curLoop, inplace, result, not, isXML){
			var name = match[1].replace(/\\/g, "");
			
			if ( !isXML && Expr.attrMap[name] ) {
				match[1] = Expr.attrMap[name];
			}

			if ( match[2] === "~=" ) {
				match[4] = " " + match[4] + " ";
			}

			return match;
		},
		PSEUDO: function(match, curLoop, inplace, result, not){
			if ( match[1] === "not" ) {
				// If we're dealing with a complex expression, or a simple one
				if ( ( chunker.exec(match[3]) || "" ).length > 1 || /^\w/.test(match[3]) ) {
					match[3] = Sizzle(match[3], null, null, curLoop);
				} else {
					var ret = Sizzle.filter(match[3], curLoop, inplace, true ^ not);
					if ( !inplace ) {
						result.push.apply( result, ret );
					}
					return false;
				}
			} else if ( Expr.match.POS.test( match[0] ) || Expr.match.CHILD.test( match[0] ) ) {
				return true;
			}
			
			return match;
		},
		POS: function(match){
			match.unshift( true );
			return match;
		}
	},
	filters: {
		enabled: function(elem){
			return elem.disabled === false && elem.type !== "hidden";
		},
		disabled: function(elem){
			return elem.disabled === true;
		},
		checked: function(elem){
			return elem.checked === true;
		},
		selected: function(elem){
			// Accessing this property makes selected-by-default
			// options in Safari work properly
			elem.parentNode.selectedIndex;
			return elem.selected === true;
		},
		parent: function(elem){
			return !!elem.firstChild;
		},
		empty: function(elem){
			return !elem.firstChild;
		},
		has: function(elem, i, match){
			return !!Sizzle( match[3], elem ).length;
		},
		header: function(elem){
			return /h\d/i.test( elem.nodeName );
		},
		text: function(elem){
			return "text" === elem.type;
		},
		radio: function(elem){
			return "radio" === elem.type;
		},
		checkbox: function(elem){
			return "checkbox" === elem.type;
		},
		file: function(elem){
			return "file" === elem.type;
		},
		password: function(elem){
			return "password" === elem.type;
		},
		submit: function(elem){
			return "submit" === elem.type;
		},
		image: function(elem){
			return "image" === elem.type;
		},
		reset: function(elem){
			return "reset" === elem.type;
		},
		button: function(elem){
			return "button" === elem.type || elem.nodeName.toLowerCase() === "button";
		},
		input: function(elem){
			return /input|select|textarea|button/i.test(elem.nodeName);
		}
	},
	setFilters: {
		first: function(elem, i){
			return i === 0;
		},
		last: function(elem, i, match, array){
			return i === array.length - 1;
		},
		even: function(elem, i){
			return i % 2 === 0;
		},
		odd: function(elem, i){
			return i % 2 === 1;
		},
		lt: function(elem, i, match){
			return i < match[3] - 0;
		},
		gt: function(elem, i, match){
			return i > match[3] - 0;
		},
		nth: function(elem, i, match){
			return match[3] - 0 === i;
		},
		eq: function(elem, i, match){
			return match[3] - 0 === i;
		}
	},
	filter: {
		PSEUDO: function(elem, match, i, array){
			var name = match[1], filter = Expr.filters[ name ];

			if ( filter ) {
				return filter( elem, i, match, array );
			} else if ( name === "contains" ) {
				return (elem.textContent || elem.innerText || getText([ elem ]) || "").indexOf(match[3]) >= 0;
			} else if ( name === "not" ) {
				var not = match[3];

				for ( var i = 0, l = not.length; i < l; i++ ) {
					if ( not[i] === elem ) {
						return false;
					}
				}

				return true;
			} else {
				Sizzle.error( "Syntax error, unrecognized expression: " + name );
			}
		},
		CHILD: function(elem, match){
			var type = match[1], node = elem;
			switch (type) {
				case 'only':
				case 'first':
					while ( (node = node.previousSibling) )	 {
						if ( node.nodeType === 1 ) { 
							return false; 
						}
					}
					if ( type === "first" ) { 
						return true; 
					}
					node = elem;
				case 'last':
					while ( (node = node.nextSibling) )	 {
						if ( node.nodeType === 1 ) { 
							return false; 
						}
					}
					return true;
				case 'nth':
					var first = match[2], last = match[3];

					if ( first === 1 && last === 0 ) {
						return true;
					}
					
					var doneName = match[0],
						parent = elem.parentNode;
	
					if ( parent && (parent.sizcache !== doneName || !elem.nodeIndex) ) {
						var count = 0;
						for ( node = parent.firstChild; node; node = node.nextSibling ) {
							if ( node.nodeType === 1 ) {
								node.nodeIndex = ++count;
							}
						} 
						parent.sizcache = doneName;
					}
					
					var diff = elem.nodeIndex - last;
					if ( first === 0 ) {
						return diff === 0;
					} else {
						return ( diff % first === 0 && diff / first >= 0 );
					}
			}
		},
		ID: function(elem, match){
			return elem.nodeType === 1 && elem.getAttribute("id") === match;
		},
		TAG: function(elem, match){
			return (match === "*" && elem.nodeType === 1) || elem.nodeName.toLowerCase() === match;
		},
		CLASS: function(elem, match){
			return (" " + (elem.className || elem.getAttribute("class")) + " ")
				.indexOf( match ) > -1;
		},
		ATTR: function(elem, match){
			var name = match[1],
				result = Expr.attrHandle[ name ] ?
					Expr.attrHandle[ name ]( elem ) :
					elem[ name ] != null ?
						elem[ name ] :
						elem.getAttribute( name ),
				value = result + "",
				type = match[2],
				check = match[4];

			return result == null ?
				type === "!=" :
				type === "=" ?
				value === check :
				type === "*=" ?
				value.indexOf(check) >= 0 :
				type === "~=" ?
				(" " + value + " ").indexOf(check) >= 0 :
				!check ?
				value && result !== false :
				type === "!=" ?
				value !== check :
				type === "^=" ?
				value.indexOf(check) === 0 :
				type === "$=" ?
				value.substr(value.length - check.length) === check :
				type === "|=" ?
				value === check || value.substr(0, check.length + 1) === check + "-" :
				false;
		},
		POS: function(elem, match, i, array){
			var name = match[2], filter = Expr.setFilters[ name ];

			if ( filter ) {
				return filter( elem, i, match, array );
			}
		}
	}
};

var origPOS = Expr.match.POS;

for ( var type in Expr.match ) {
	Expr.match[ type ] = new RegExp( Expr.match[ type ].source + /(?![^\[]*\])(?![^\(]*\))/.source );
	Expr.leftMatch[ type ] = new RegExp( /(^(?:.|\r|\n)*?)/.source + Expr.match[ type ].source.replace(/\\(\d+)/g, function(all, num){
		return "\\" + (num - 0 + 1);
	}));
}

var makeArray = function(array, results) {
	array = Array.prototype.slice.call( array, 0 );

	if ( results ) {
		results.push.apply( results, array );
		return results;
	}
	
	return array;
};

// Perform a simple check to determine if the browser is capable of
// converting a NodeList to an array using builtin methods.
// Also verifies that the returned array holds DOM nodes
// (which is not the case in the Blackberry browser)
try {
	Array.prototype.slice.call( document.documentElement.childNodes, 0 )[0].nodeType;

// Provide a fallback method if it does not work
} catch(e){
	makeArray = function(array, results) {
		var ret = results || [];

		if ( toString.call(array) === "[object Array]" ) {
			Array.prototype.push.apply( ret, array );
		} else {
			if ( typeof array.length === "number" ) {
				for ( var i = 0, l = array.length; i < l; i++ ) {
					ret.push( array[i] );
				}
			} else {
				for ( var i = 0; array[i]; i++ ) {
					ret.push( array[i] );
				}
			}
		}

		return ret;
	};
}

var sortOrder;

if ( document.documentElement.compareDocumentPosition ) {
	sortOrder = function( a, b ) {
		if ( !a.compareDocumentPosition || !b.compareDocumentPosition ) {
			if ( a == b ) {
				hasDuplicate = true;
			}
			return a.compareDocumentPosition ? -1 : 1;
		}

		var ret = a.compareDocumentPosition(b) & 4 ? -1 : a === b ? 0 : 1;
		if ( ret === 0 ) {
			hasDuplicate = true;
		}
		return ret;
	};
} else if ( "sourceIndex" in document.documentElement ) {
	sortOrder = function( a, b ) {
		if ( !a.sourceIndex || !b.sourceIndex ) {
			if ( a == b ) {
				hasDuplicate = true;
			}
			return a.sourceIndex ? -1 : 1;
		}

		var ret = a.sourceIndex - b.sourceIndex;
		if ( ret === 0 ) {
			hasDuplicate = true;
		}
		return ret;
	};
} else if ( document.createRange ) {
	sortOrder = function( a, b ) {
		if ( !a.ownerDocument || !b.ownerDocument ) {
			if ( a == b ) {
				hasDuplicate = true;
			}
			return a.ownerDocument ? -1 : 1;
		}

		var aRange = a.ownerDocument.createRange(), bRange = b.ownerDocument.createRange();
		aRange.setStart(a, 0);
		aRange.setEnd(a, 0);
		bRange.setStart(b, 0);
		bRange.setEnd(b, 0);
		var ret = aRange.compareBoundaryPoints(Range.START_TO_END, bRange);
		if ( ret === 0 ) {
			hasDuplicate = true;
		}
		return ret;
	};
}

// Utility function for retreiving the text value of an array of DOM nodes
function getText( elems ) {
	var ret = "", elem;

	for ( var i = 0; elems[i]; i++ ) {
		elem = elems[i];

		// Get the text from text nodes and CDATA nodes
		if ( elem.nodeType === 3 || elem.nodeType === 4 ) {
			ret += elem.nodeValue;

		// Traverse everything else, except comment nodes
		} else if ( elem.nodeType !== 8 ) {
			ret += getText( elem.childNodes );
		}
	}

	return ret;
}

// Check to see if the browser returns elements by name when
// querying by getElementById (and provide a workaround)
(function(){
	// We're going to inject a fake input element with a specified name
	var form = document.createElement("div"),
		id = "script" + (new Date).getTime();
	form.innerHTML = "<a name='" + id + "'/>";

	// Inject it into the root element, check its status, and remove it quickly
	var root = document.documentElement;
	root.insertBefore( form, root.firstChild );

	// The workaround has to do additional checks after a getElementById
	// Which slows things down for other browsers (hence the branching)
	if ( document.getElementById( id ) ) {
		Expr.find.ID = function(match, context, isXML){
			if ( typeof context.getElementById !== "undefined" && !isXML ) {
				var m = context.getElementById(match[1]);
				return m ? m.id === match[1] || typeof m.getAttributeNode !== "undefined" && m.getAttributeNode("id").nodeValue === match[1] ? [m] : undefined : [];
			}
		};

		Expr.filter.ID = function(elem, match){
			var node = typeof elem.getAttributeNode !== "undefined" && elem.getAttributeNode("id");
			return elem.nodeType === 1 && node && node.nodeValue === match;
		};
	}

	root.removeChild( form );
	root = form = null; // release memory in IE
})();

(function(){
	// Check to see if the browser returns only elements
	// when doing getElementsByTagName("*")

	// Create a fake element
	var div = document.createElement("div");
	div.appendChild( document.createComment("") );

	// Make sure no comments are found
	if ( div.getElementsByTagName("*").length > 0 ) {
		Expr.find.TAG = function(match, context){
			var results = context.getElementsByTagName(match[1]);

			// Filter out possible comments
			if ( match[1] === "*" ) {
				var tmp = [];

				for ( var i = 0; results[i]; i++ ) {
					if ( results[i].nodeType === 1 ) {
						tmp.push( results[i] );
					}
				}

				results = tmp;
			}

			return results;
		};
	}

	// Check to see if an attribute returns normalized href attributes
	div.innerHTML = "<a href='#'></a>";
	if ( div.firstChild && typeof div.firstChild.getAttribute !== "undefined" &&
			div.firstChild.getAttribute("href") !== "#" ) {
		Expr.attrHandle.href = function(elem){
			return elem.getAttribute("href", 2);
		};
	}

	div = null; // release memory in IE
})();

if ( document.querySelectorAll ) {
	(function(){
		var oldSizzle = Sizzle, div = document.createElement("div");
		div.innerHTML = "<p class='TEST'></p>";

		// Safari can't handle uppercase or unicode characters when
		// in quirks mode.
		if ( div.querySelectorAll && div.querySelectorAll(".TEST").length === 0 ) {
			return;
		}
	
		Sizzle = function(query, context, extra, seed){
			context = context || document;

			// Only use querySelectorAll on non-XML documents
			// (ID selectors don't work in non-HTML documents)
			if ( !seed && context.nodeType === 9 && !isXML(context) ) {
				try {
					return makeArray( context.querySelectorAll(query), extra );
				} catch(e){}
			}
		
			return oldSizzle(query, context, extra, seed);
		};

		for ( var prop in oldSizzle ) {
			Sizzle[ prop ] = oldSizzle[ prop ];
		}

		div = null; // release memory in IE
	})();
}

(function(){
	var div = document.createElement("div");

	div.innerHTML = "<div class='test e'></div><div class='test'></div>";

	// Opera can't find a second classname (in 9.6)
	// Also, make sure that getElementsByClassName actually exists
	if ( !div.getElementsByClassName || div.getElementsByClassName("e").length === 0 ) {
		return;
	}

	// Safari caches class attributes, doesn't catch changes (in 3.2)
	div.lastChild.className = "e";

	if ( div.getElementsByClassName("e").length === 1 ) {
		return;
	}
	
	Expr.order.splice(1, 0, "CLASS");
	Expr.find.CLASS = function(match, context, isXML) {
		if ( typeof context.getElementsByClassName !== "undefined" && !isXML ) {
			return context.getElementsByClassName(match[1]);
		}
	};

	div = null; // release memory in IE
})();

function dirNodeCheck( dir, cur, doneName, checkSet, nodeCheck, isXML ) {
	for ( var i = 0, l = checkSet.length; i < l; i++ ) {
		var elem = checkSet[i];
		if ( elem ) {
			elem = elem[dir];
			var match = false;

			while ( elem ) {
				if ( elem.sizcache === doneName ) {
					match = checkSet[elem.sizset];
					break;
				}

				if ( elem.nodeType === 1 && !isXML ){
					elem.sizcache = doneName;
					elem.sizset = i;
				}

				if ( elem.nodeName.toLowerCase() === cur ) {
					match = elem;
					break;
				}

				elem = elem[dir];
			}

			checkSet[i] = match;
		}
	}
}

function dirCheck( dir, cur, doneName, checkSet, nodeCheck, isXML ) {
	for ( var i = 0, l = checkSet.length; i < l; i++ ) {
		var elem = checkSet[i];
		if ( elem ) {
			elem = elem[dir];
			var match = false;

			while ( elem ) {
				if ( elem.sizcache === doneName ) {
					match = checkSet[elem.sizset];
					break;
				}

				if ( elem.nodeType === 1 ) {
					if ( !isXML ) {
						elem.sizcache = doneName;
						elem.sizset = i;
					}
					if ( typeof cur !== "string" ) {
						if ( elem === cur ) {
							match = true;
							break;
						}

					} else if ( Sizzle.filter( cur, [elem] ).length > 0 ) {
						match = elem;
						break;
					}
				}

				elem = elem[dir];
			}

			checkSet[i] = match;
		}
	}
}

var contains = document.compareDocumentPosition ? function(a, b){
	return !!(a.compareDocumentPosition(b) & 16);
} : function(a, b){
	return a !== b && (a.contains ? a.contains(b) : true);
};

var isXML = function(elem){
	// documentElement is verified for cases where it doesn't yet exist
	// (such as loading iframes in IE - #4833) 
	var documentElement = (elem ? elem.ownerDocument || elem : 0).documentElement;
	return documentElement ? documentElement.nodeName !== "HTML" : false;
};

var posProcess = function(selector, context){
	var tmpSet = [], later = "", match,
		root = context.nodeType ? [context] : context;

	// Position selectors must be done after the filter
	// And so must :not(positional) so we move all PSEUDOs to the end
	while ( (match = Expr.match.PSEUDO.exec( selector )) ) {
		later += match[0];
		selector = selector.replace( Expr.match.PSEUDO, "" );
	}

	selector = Expr.relative[selector] ? selector + "*" : selector;

	for ( var i = 0, l = root.length; i < l; i++ ) {
		Sizzle( selector, root[i], tmpSet );
	}

	return Sizzle.filter( later, tmpSet );
};

// EXPOSE
jQuery.find = Sizzle;
jQuery.expr = Sizzle.selectors;
jQuery.expr[":"] = jQuery.expr.filters;
jQuery.unique = Sizzle.uniqueSort;
jQuery.text = getText;
jQuery.isXMLDoc = isXML;
jQuery.contains = contains;

return;

window.Sizzle = Sizzle;

})();
var runtil = /Until$/,
	rparentsprev = /^(?:parents|prevUntil|prevAll)/,
	// Note: This RegExp should be improved, or likely pulled from Sizzle
	rmultiselector = /,/,
	slice = Array.prototype.slice;

// Implement the identical functionality for filter and not
var winnow = function( elements, qualifier, keep ) {
	if ( jQuery.isFunction( qualifier ) ) {
		return jQuery.grep(elements, function( elem, i ) {
			return !!qualifier.call( elem, i, elem ) === keep;
		});

	} else if ( qualifier.nodeType ) {
		return jQuery.grep(elements, function( elem, i ) {
			return (elem === qualifier) === keep;
		});

	} else if ( typeof qualifier === "string" ) {
		var filtered = jQuery.grep(elements, function( elem ) {
			return elem.nodeType === 1;
		});

		if ( isSimple.test( qualifier ) ) {
			return jQuery.filter(qualifier, filtered, !keep);
		} else {
			qualifier = jQuery.filter( qualifier, filtered );
		}
	}

	return jQuery.grep(elements, function( elem, i ) {
		return (jQuery.inArray( elem, qualifier ) >= 0) === keep;
	});
};

jQuery.fn.extend({
	find: function( selector ) {
		var ret = this.pushStack( "", "find", selector ), length = 0;

		for ( var i = 0, l = this.length; i < l; i++ ) {
			length = ret.length;
			jQuery.find( selector, this[i], ret );

			if ( i > 0 ) {
				// Make sure that the results are unique
				for ( var n = length; n < ret.length; n++ ) {
					for ( var r = 0; r < length; r++ ) {
						if ( ret[r] === ret[n] ) {
							ret.splice(n--, 1);
							break;
						}
					}
				}
			}
		}

		return ret;
	},

	has: function( target ) {
		var targets = jQuery( target );
		return this.filter(function() {
			for ( var i = 0, l = targets.length; i < l; i++ ) {
				if ( jQuery.contains( this, targets[i] ) ) {
					return true;
				}
			}
		});
	},

	not: function( selector ) {
		return this.pushStack( winnow(this, selector, false), "not", selector);
	},

	filter: function( selector ) {
		return this.pushStack( winnow(this, selector, true), "filter", selector );
	},
	
	is: function( selector ) {
		return !!selector && jQuery.filter( selector, this ).length > 0;
	},

	closest: function( selectors, context ) {
		if ( jQuery.isArray( selectors ) ) {
			var ret = [], cur = this[0], match, matches = {}, selector;

			if ( cur && selectors.length ) {
				for ( var i = 0, l = selectors.length; i < l; i++ ) {
					selector = selectors[i];

					if ( !matches[selector] ) {
						matches[selector] = jQuery.expr.match.POS.test( selector ) ? 
							jQuery( selector, context || this.context ) :
							selector;
					}
				}

				while ( cur && cur.ownerDocument && cur !== context ) {
					for ( selector in matches ) {
						match = matches[selector];

						if ( match.jquery ? match.index(cur) > -1 : jQuery(cur).is(match) ) {
							ret.push({ selector: selector, elem: cur });
							delete matches[selector];
						}
					}
					cur = cur.parentNode;
				}
			}

			return ret;
		}

		var pos = jQuery.expr.match.POS.test( selectors ) ? 
			jQuery( selectors, context || this.context ) : null;

		return this.map(function( i, cur ) {
			while ( cur && cur.ownerDocument && cur !== context ) {
				if ( pos ? pos.index(cur) > -1 : jQuery(cur).is(selectors) ) {
					return cur;
				}
				cur = cur.parentNode;
			}
			return null;
		});
	},
	
	// Determine the position of an element within
	// the matched set of elements
	index: function( elem ) {
		if ( !elem || typeof elem === "string" ) {
			return jQuery.inArray( this[0],
				// If it receives a string, the selector is used
				// If it receives nothing, the siblings are used
				elem ? jQuery( elem ) : this.parent().children() );
		}
		// Locate the position of the desired element
		return jQuery.inArray(
			// If it receives a jQuery object, the first element is used
			elem.jquery ? elem[0] : elem, this );
	},

	add: function( selector, context ) {
		var set = typeof selector === "string" ?
				jQuery( selector, context || this.context ) :
				jQuery.makeArray( selector ),
			all = jQuery.merge( this.get(), set );

		return this.pushStack( isDisconnected( set[0] ) || isDisconnected( all[0] ) ?
			all :
			jQuery.unique( all ) );
	},

	andSelf: function() {
		return this.add( this.prevObject );
	}
});

// A painfully simple check to see if an element is disconnected
// from a document (should be improved, where feasible).
function isDisconnected( node ) {
	return !node || !node.parentNode || node.parentNode.nodeType === 11;
}

jQuery.each({
	parent: function( elem ) {
		var parent = elem.parentNode;
		return parent && parent.nodeType !== 11 ? parent : null;
	},
	parents: function( elem ) {
		return jQuery.dir( elem, "parentNode" );
	},
	parentsUntil: function( elem, i, until ) {
		return jQuery.dir( elem, "parentNode", until );
	},
	next: function( elem ) {
		return jQuery.nth( elem, 2, "nextSibling" );
	},
	prev: function( elem ) {
		return jQuery.nth( elem, 2, "previousSibling" );
	},
	nextAll: function( elem ) {
		return jQuery.dir( elem, "nextSibling" );
	},
	prevAll: function( elem ) {
		return jQuery.dir( elem, "previousSibling" );
	},
	nextUntil: function( elem, i, until ) {
		return jQuery.dir( elem, "nextSibling", until );
	},
	prevUntil: function( elem, i, until ) {
		return jQuery.dir( elem, "previousSibling", until );
	},
	siblings: function( elem ) {
		return jQuery.sibling( elem.parentNode.firstChild, elem );
	},
	children: function( elem ) {
		return jQuery.sibling( elem.firstChild );
	},
	contents: function( elem ) {
		return jQuery.nodeName( elem, "iframe" ) ?
			elem.contentDocument || elem.contentWindow.document :
			jQuery.makeArray( elem.childNodes );
	}
}, function( name, fn ) {
	jQuery.fn[ name ] = function( until, selector ) {
		var ret = jQuery.map( this, fn, until );
		
		if ( !runtil.test( name ) ) {
			selector = until;
		}

		if ( selector && typeof selector === "string" ) {
			ret = jQuery.filter( selector, ret );
		}

		ret = this.length > 1 ? jQuery.unique( ret ) : ret;

		if ( (this.length > 1 || rmultiselector.test( selector )) && rparentsprev.test( name ) ) {
			ret = ret.reverse();
		}

		return this.pushStack( ret, name, slice.call(arguments).join(",") );
	};
});

jQuery.extend({
	filter: function( expr, elems, not ) {
		if ( not ) {
			expr = ":not(" + expr + ")";
		}

		return jQuery.find.matches(expr, elems);
	},
	
	dir: function( elem, dir, until ) {
		var matched = [], cur = elem[dir];
		while ( cur && cur.nodeType !== 9 && (until === undefined || cur.nodeType !== 1 || !jQuery( cur ).is( until )) ) {
			if ( cur.nodeType === 1 ) {
				matched.push( cur );
			}
			cur = cur[dir];
		}
		return matched;
	},

	nth: function( cur, result, dir, elem ) {
		result = result || 1;
		var num = 0;

		for ( ; cur; cur = cur[dir] ) {
			if ( cur.nodeType === 1 && ++num === result ) {
				break;
			}
		}

		return cur;
	},

	sibling: function( n, elem ) {
		var r = [];

		for ( ; n; n = n.nextSibling ) {
			if ( n.nodeType === 1 && n !== elem ) {
				r.push( n );
			}
		}

		return r;
	}
});
var rinlinejQuery = / jQuery\d+="(?:\d+|null)"/g,
	rleadingWhitespace = /^\s+/,
	rxhtmlTag = /(<([\w:]+)[^>]*?)\/>/g,
	rselfClosing = /^(?:area|br|col|embed|hr|img|input|link|meta|param)$/i,
	rtagName = /<([\w:]+)/,
	rtbody = /<tbody/i,
	rhtml = /<|&#?\w+;/,
	rnocache = /<script|<object|<embed|<option|<style/i,
	rchecked = /checked\s*(?:[^=]|=\s*.checked.)/i,  // checked="checked" or checked (html5)
	fcloseTag = function( all, front, tag ) {
		return rselfClosing.test( tag ) ?
			all :
			front + "></" + tag + ">";
	},
	wrapMap = {
		option: [ 1, "<select multiple='multiple'>", "</select>" ],
		legend: [ 1, "<fieldset>", "</fieldset>" ],
		thead: [ 1, "<table>", "</table>" ],
		tr: [ 2, "<table><tbody>", "</tbody></table>" ],
		td: [ 3, "<table><tbody><tr>", "</tr></tbody></table>" ],
		col: [ 2, "<table><tbody></tbody><colgroup>", "</colgroup></table>" ],
		area: [ 1, "<map>", "</map>" ],
		_default: [ 0, "", "" ]
	};

wrapMap.optgroup = wrapMap.option;
wrapMap.tbody = wrapMap.tfoot = wrapMap.colgroup = wrapMap.caption = wrapMap.thead;
wrapMap.th = wrapMap.td;

// IE can't serialize <link> and <script> tags normally
if ( !jQuery.support.htmlSerialize ) {
	wrapMap._default = [ 1, "div<div>", "</div>" ];
}

jQuery.fn.extend({
	text: function( text ) {
		if ( jQuery.isFunction(text) ) {
			return this.each(function(i) {
				var self = jQuery(this);
				self.text( text.call(this, i, self.text()) );
			});
		}

		if ( typeof text !== "object" && text !== undefined ) {
			return this.empty().append( (this[0] && this[0].ownerDocument || document).createTextNode( text ) );
		}

		return jQuery.text( this );
	},

	wrapAll: function( html ) {
		if ( jQuery.isFunction( html ) ) {
			return this.each(function(i) {
				jQuery(this).wrapAll( html.call(this, i) );
			});
		}

		if ( this[0] ) {
			// The elements to wrap the target around
			var wrap = jQuery( html, this[0].ownerDocument ).eq(0).clone(true);

			if ( this[0].parentNode ) {
				wrap.insertBefore( this[0] );
			}

			wrap.map(function() {
				var elem = this;

				while ( elem.firstChild && elem.firstChild.nodeType === 1 ) {
					elem = elem.firstChild;
				}

				return elem;
			}).append(this);
		}

		return this;
	},

	wrapInner: function( html ) {
		if ( jQuery.isFunction( html ) ) {
			return this.each(function(i) {
				jQuery(this).wrapInner( html.call(this, i) );
			});
		}

		return this.each(function() {
			var self = jQuery( this ), contents = self.contents();

			if ( contents.length ) {
				contents.wrapAll( html );

			} else {
				self.append( html );
			}
		});
	},

	wrap: function( html ) {
		return this.each(function() {
			jQuery( this ).wrapAll( html );
		});
	},

	unwrap: function() {
		return this.parent().each(function() {
			if ( !jQuery.nodeName( this, "body" ) ) {
				jQuery( this ).replaceWith( this.childNodes );
			}
		}).end();
	},

	append: function() {
		return this.domManip(arguments, true, function( elem ) {
			if ( this.nodeType === 1 ) {
				this.appendChild( elem );
			}
		});
	},

	prepend: function() {
		return this.domManip(arguments, true, function( elem ) {
			if ( this.nodeType === 1 ) {
				this.insertBefore( elem, this.firstChild );
			}
		});
	},

	before: function() {
		if ( this[0] && this[0].parentNode ) {
			return this.domManip(arguments, false, function( elem ) {
				this.parentNode.insertBefore( elem, this );
			});
		} else if ( arguments.length ) {
			var set = jQuery(arguments[0]);
			set.push.apply( set, this.toArray() );
			return this.pushStack( set, "before", arguments );
		}
	},

	after: function() {
		if ( this[0] && this[0].parentNode ) {
			return this.domManip(arguments, false, function( elem ) {
				this.parentNode.insertBefore( elem, this.nextSibling );
			});
		} else if ( arguments.length ) {
			var set = this.pushStack( this, "after", arguments );
			set.push.apply( set, jQuery(arguments[0]).toArray() );
			return set;
		}
	},
	
	// keepData is for internal use only--do not document
	remove: function( selector, keepData ) {
		for ( var i = 0, elem; (elem = this[i]) != null; i++ ) {
			if ( !selector || jQuery.filter( selector, [ elem ] ).length ) {
				if ( !keepData && elem.nodeType === 1 ) {
					jQuery.cleanData( elem.getElementsByTagName("*") );
					jQuery.cleanData( [ elem ] );
				}

				if ( elem.parentNode ) {
					 elem.parentNode.removeChild( elem );
				}
			}
		}
		
		return this;
	},

	empty: function() {
		for ( var i = 0, elem; (elem = this[i]) != null; i++ ) {
			// Remove element nodes and prevent memory leaks
			if ( elem.nodeType === 1 ) {
				jQuery.cleanData( elem.getElementsByTagName("*") );
			}

			// Remove any remaining nodes
			while ( elem.firstChild ) {
				elem.removeChild( elem.firstChild );
			}
		}
		
		return this;
	},

	clone: function( events ) {
		// Do the clone
		var ret = this.map(function() {
			if ( !jQuery.support.noCloneEvent && !jQuery.isXMLDoc(this) ) {
				// IE copies events bound via attachEvent when
				// using cloneNode. Calling detachEvent on the
				// clone will also remove the events from the orignal
				// In order to get around this, we use innerHTML.
				// Unfortunately, this means some modifications to
				// attributes in IE that are actually only stored
				// as properties will not be copied (such as the
				// the name attribute on an input).
				var html = this.outerHTML, ownerDocument = this.ownerDocument;
				if ( !html ) {
					var div = ownerDocument.createElement("div");
					div.appendChild( this.cloneNode(true) );
					html = div.innerHTML;
				}

				return jQuery.clean([html.replace(rinlinejQuery, "")
					// Handle the case in IE 8 where action=/test/> self-closes a tag
					.replace(/=([^="'>\s]+\/)>/g, '="$1">')
					.replace(rleadingWhitespace, "")], ownerDocument)[0];
			} else {
				return this.cloneNode(true);
			}
		});

		// Copy the events from the original to the clone
		if ( events === true ) {
			cloneCopyEvent( this, ret );
			cloneCopyEvent( this.find("*"), ret.find("*") );
		}

		// Return the cloned set
		return ret;
	},

	html: function( value ) {
		if ( value === undefined ) {
			return this[0] && this[0].nodeType === 1 ?
				this[0].innerHTML.replace(rinlinejQuery, "") :
				null;

		// See if we can take a shortcut and just use innerHTML
		} else if ( typeof value === "string" && !rnocache.test( value ) &&
			(jQuery.support.leadingWhitespace || !rleadingWhitespace.test( value )) &&
			!wrapMap[ (rtagName.exec( value ) || ["", ""])[1].toLowerCase() ] ) {

			value = value.replace(rxhtmlTag, fcloseTag);

			try {
				for ( var i = 0, l = this.length; i < l; i++ ) {
					// Remove element nodes and prevent memory leaks
					if ( this[i].nodeType === 1 ) {
						jQuery.cleanData( this[i].getElementsByTagName("*") );
						this[i].innerHTML = value;
					}
				}

			// If using innerHTML throws an exception, use the fallback method
			} catch(e) {
				this.empty().append( value );
			}

		} else if ( jQuery.isFunction( value ) ) {
			this.each(function(i){
				var self = jQuery(this), old = self.html();
				self.empty().append(function(){
					return value.call( this, i, old );
				});
			});

		} else {
			this.empty().append( value );
		}

		return this;
	},

	replaceWith: function( value ) {
		if ( this[0] && this[0].parentNode ) {
			// Make sure that the elements are removed from the DOM before they are inserted
			// this can help fix replacing a parent with child elements
			if ( jQuery.isFunction( value ) ) {
				return this.each(function(i) {
					var self = jQuery(this), old = self.html();
					self.replaceWith( value.call( this, i, old ) );
				});
			}

			if ( typeof value !== "string" ) {
				value = jQuery(value).detach();
			}

			return this.each(function() {
				var next = this.nextSibling, parent = this.parentNode;

				jQuery(this).remove();

				if ( next ) {
					jQuery(next).before( value );
				} else {
					jQuery(parent).append( value );
				}
			});
		} else {
			return this.pushStack( jQuery(jQuery.isFunction(value) ? value() : value), "replaceWith", value );
		}
	},

	detach: function( selector ) {
		return this.remove( selector, true );
	},

	domManip: function( args, table, callback ) {
		var results, first, value = args[0], scripts = [], fragment, parent;

		// We can't cloneNode fragments that contain checked, in WebKit
		if ( !jQuery.support.checkClone && arguments.length === 3 && typeof value === "string" && rchecked.test( value ) ) {
			return this.each(function() {
				jQuery(this).domManip( args, table, callback, true );
			});
		}

		if ( jQuery.isFunction(value) ) {
			return this.each(function(i) {
				var self = jQuery(this);
				args[0] = value.call(this, i, table ? self.html() : undefined);
				self.domManip( args, table, callback );
			});
		}

		if ( this[0] ) {
			parent = value && value.parentNode;

			// If we're in a fragment, just use that instead of building a new one
			if ( jQuery.support.parentNode && parent && parent.nodeType === 11 && parent.childNodes.length === this.length ) {
				results = { fragment: parent };

			} else {
				results = buildFragment( args, this, scripts );
			}
			
			fragment = results.fragment;
			
			if ( fragment.childNodes.length === 1 ) {
				first = fragment = fragment.firstChild;
			} else {
				first = fragment.firstChild;
			}

			if ( first ) {
				table = table && jQuery.nodeName( first, "tr" );

				for ( var i = 0, l = this.length; i < l; i++ ) {
					callback.call(
						table ?
							root(this[i], first) :
							this[i],
						i > 0 || results.cacheable || this.length > 1  ?
							fragment.cloneNode(true) :
							fragment
					);
				}
			}

			if ( scripts.length ) {
				jQuery.each( scripts, evalScript );
			}
		}

		return this;

		function root( elem, cur ) {
			return jQuery.nodeName(elem, "table") ?
				(elem.getElementsByTagName("tbody")[0] ||
				elem.appendChild(elem.ownerDocument.createElement("tbody"))) :
				elem;
		}
	}
});

function cloneCopyEvent(orig, ret) {
	var i = 0;

	ret.each(function() {
		if ( this.nodeName !== (orig[i] && orig[i].nodeName) ) {
			return;
		}

		var oldData = jQuery.data( orig[i++] ), curData = jQuery.data( this, oldData ), events = oldData && oldData.events;

		if ( events ) {
			delete curData.handle;
			curData.events = {};

			for ( var type in events ) {
				for ( var handler in events[ type ] ) {
					jQuery.event.add( this, type, events[ type ][ handler ], events[ type ][ handler ].data );
				}
			}
		}
	});
}

function buildFragment( args, nodes, scripts ) {
	var fragment, cacheable, cacheresults,
		doc = (nodes && nodes[0] ? nodes[0].ownerDocument || nodes[0] : document);

	// Only cache "small" (1/2 KB) strings that are associated with the main document
	// Cloning options loses the selected state, so don't cache them
	// IE 6 doesn't like it when you put <object> or <embed> elements in a fragment
	// Also, WebKit does not clone 'checked' attributes on cloneNode, so don't cache
	if ( args.length === 1 && typeof args[0] === "string" && args[0].length < 512 && doc === document &&
		!rnocache.test( args[0] ) && (jQuery.support.checkClone || !rchecked.test( args[0] )) ) {

		cacheable = true;
		cacheresults = jQuery.fragments[ args[0] ];
		if ( cacheresults ) {
			if ( cacheresults !== 1 ) {
				fragment = cacheresults;
			}
		}
	}

	if ( !fragment ) {
		fragment = doc.createDocumentFragment();
		jQuery.clean( args, doc, fragment, scripts );
	}

	if ( cacheable ) {
		jQuery.fragments[ args[0] ] = cacheresults ? fragment : 1;
	}

	return { fragment: fragment, cacheable: cacheable };
}

jQuery.fragments = {};

jQuery.each({
	appendTo: "append",
	prependTo: "prepend",
	insertBefore: "before",
	insertAfter: "after",
	replaceAll: "replaceWith"
}, function( name, original ) {
	jQuery.fn[ name ] = function( selector ) {
		var ret = [], insert = jQuery( selector ),
			parent = this.length === 1 && this[0].parentNode;
		
		if ( parent && parent.nodeType === 11 && parent.childNodes.length === 1 && insert.length === 1 ) {
			insert[ original ]( this[0] );
			return this;
			
		} else {
			for ( var i = 0, l = insert.length; i < l; i++ ) {
				var elems = (i > 0 ? this.clone(true) : this).get();
				jQuery.fn[ original ].apply( jQuery(insert[i]), elems );
				ret = ret.concat( elems );
			}
		
			return this.pushStack( ret, name, insert.selector );
		}
	};
});

jQuery.extend({
	clean: function( elems, context, fragment, scripts ) {
		context = context || document;

		// !context.createElement fails in IE with an error but returns typeof 'object'
		if ( typeof context.createElement === "undefined" ) {
			context = context.ownerDocument || context[0] && context[0].ownerDocument || document;
		}

		var ret = [];

		for ( var i = 0, elem; (elem = elems[i]) != null; i++ ) {
			if ( typeof elem === "number" ) {
				elem += "";
			}

			if ( !elem ) {
				continue;
			}

			// Convert html string into DOM nodes
			if ( typeof elem === "string" && !rhtml.test( elem ) ) {
				elem = context.createTextNode( elem );

			} else if ( typeof elem === "string" ) {
				// Fix "XHTML"-style tags in all browsers
				elem = elem.replace(rxhtmlTag, fcloseTag);

				// Trim whitespace, otherwise indexOf won't work as expected
				var tag = (rtagName.exec( elem ) || ["", ""])[1].toLowerCase(),
					wrap = wrapMap[ tag ] || wrapMap._default,
					depth = wrap[0],
					div = context.createElement("div");

				// Go to html and back, then peel off extra wrappers
				div.innerHTML = wrap[1] + elem + wrap[2];

				// Move to the right depth
				while ( depth-- ) {
					div = div.lastChild;
				}

				// Remove IE's autoinserted <tbody> from table fragments
				if ( !jQuery.support.tbody ) {

					// String was a <table>, *may* have spurious <tbody>
					var hasBody = rtbody.test(elem),
						tbody = tag === "table" && !hasBody ?
							div.firstChild && div.firstChild.childNodes :

							// String was a bare <thead> or <tfoot>
							wrap[1] === "<table>" && !hasBody ?
								div.childNodes :
								[];

					for ( var j = tbody.length - 1; j >= 0 ; --j ) {
						if ( jQuery.nodeName( tbody[ j ], "tbody" ) && !tbody[ j ].childNodes.length ) {
							tbody[ j ].parentNode.removeChild( tbody[ j ] );
						}
					}

				}

				// IE completely kills leading whitespace when innerHTML is used
				if ( !jQuery.support.leadingWhitespace && rleadingWhitespace.test( elem ) ) {
					div.insertBefore( context.createTextNode( rleadingWhitespace.exec(elem)[0] ), div.firstChild );
				}

				elem = div.childNodes;
			}

			if ( elem.nodeType ) {
				ret.push( elem );
			} else {
				ret = jQuery.merge( ret, elem );
			}
		}

		if ( fragment ) {
			for ( var i = 0; ret[i]; i++ ) {
				if ( scripts && jQuery.nodeName( ret[i], "script" ) && (!ret[i].type || ret[i].type.toLowerCase() === "text/javascript") ) {
					scripts.push( ret[i].parentNode ? ret[i].parentNode.removeChild( ret[i] ) : ret[i] );
				
				} else {
					if ( ret[i].nodeType === 1 ) {
						ret.splice.apply( ret, [i + 1, 0].concat(jQuery.makeArray(ret[i].getElementsByTagName("script"))) );
					}
					fragment.appendChild( ret[i] );
				}
			}
		}

		return ret;
	},
	
	cleanData: function( elems ) {
		var data, id, cache = jQuery.cache,
			special = jQuery.event.special,
			deleteExpando = jQuery.support.deleteExpando;
		
		for ( var i = 0, elem; (elem = elems[i]) != null; i++ ) {
			id = elem[ jQuery.expando ];
			
			if ( id ) {
				data = cache[ id ];
				
				if ( data.events ) {
					for ( var type in data.events ) {
						if ( special[ type ] ) {
							jQuery.event.remove( elem, type );

						} else {
							removeEvent( elem, type, data.handle );
						}
					}
				}
				
				if ( deleteExpando ) {
					delete elem[ jQuery.expando ];

				} else if ( elem.removeAttribute ) {
					elem.removeAttribute( jQuery.expando );
				}
				
				delete cache[ id ];
			}
		}
	}
});
// exclude the following css properties to add px
var rexclude = /z-?index|font-?weight|opacity|zoom|line-?height/i,
	ralpha = /alpha\([^)]*\)/,
	ropacity = /opacity=([^)]*)/,
	rfloat = /float/i,
	rdashAlpha = /-([a-z])/ig,
	rupper = /([A-Z])/g,
	rnumpx = /^-?\d+(?:px)?$/i,
	rnum = /^-?\d/,

	cssShow = { position: "absolute", visibility: "hidden", display:"block" },
	cssWidth = [ "Left", "Right" ],
	cssHeight = [ "Top", "Bottom" ],

	// cache check for defaultView.getComputedStyle
	getComputedStyle = document.defaultView && document.defaultView.getComputedStyle,
	// normalize float css property
	styleFloat = jQuery.support.cssFloat ? "cssFloat" : "styleFloat",
	fcamelCase = function( all, letter ) {
		return letter.toUpperCase();
	};

jQuery.fn.css = function( name, value ) {
	return access( this, name, value, true, function( elem, name, value ) {
		if ( value === undefined ) {
			return jQuery.curCSS( elem, name );
		}
		
		if ( typeof value === "number" && !rexclude.test(name) ) {
			value += "px";
		}

		jQuery.style( elem, name, value );
	});
};

jQuery.extend({
	style: function( elem, name, value ) {
		// don't set styles on text and comment nodes
		if ( !elem || elem.nodeType === 3 || elem.nodeType === 8 ) {
			return undefined;
		}

		// ignore negative width and height values #1599
		if ( (name === "width" || name === "height") && parseFloat(value) < 0 ) {
			value = undefined;
		}

		var style = elem.style || elem, set = value !== undefined;

		// IE uses filters for opacity
		if ( !jQuery.support.opacity && name === "opacity" ) {
			if ( set ) {
				// IE has trouble with opacity if it does not have layout
				// Force it by setting the zoom level
				style.zoom = 1;

				// Set the alpha filter to set the opacity
				var opacity = parseInt( value, 10 ) + "" === "NaN" ? "" : "alpha(opacity=" + value * 100 + ")";
				var filter = style.filter || jQuery.curCSS( elem, "filter" ) || "";
				style.filter = ralpha.test(filter) ? filter.replace(ralpha, opacity) : opacity;
			}

			return style.filter && style.filter.indexOf("opacity=") >= 0 ?
				(parseFloat( ropacity.exec(style.filter)[1] ) / 100) + "":
				"";
		}

		// Make sure we're using the right name for getting the float value
		if ( rfloat.test( name ) ) {
			name = styleFloat;
		}

		name = name.replace(rdashAlpha, fcamelCase);

		if ( set ) {
			style[ name ] = value;
		}

		return style[ name ];
	},

	css: function( elem, name, force, extra ) {
		if ( name === "width" || name === "height" ) {
			var val, props = cssShow, which = name === "width" ? cssWidth : cssHeight;

			function getWH() {
				val = name === "width" ? elem.offsetWidth : elem.offsetHeight;

				if ( extra === "border" ) {
					return;
				}

				jQuery.each( which, function() {
					if ( !extra ) {
						val -= parseFloat(jQuery.curCSS( elem, "padding" + this, true)) || 0;
					}

					if ( extra === "margin" ) {
						val += parseFloat(jQuery.curCSS( elem, "margin" + this, true)) || 0;
					} else {
						val -= parseFloat(jQuery.curCSS( elem, "border" + this + "Width", true)) || 0;
					}
				});
			}

			if ( elem.offsetWidth !== 0 ) {
				getWH();
			} else {
				jQuery.swap( elem, props, getWH );
			}

			return Math.max(0, Math.round(val));
		}

		return jQuery.curCSS( elem, name, force );
	},

	curCSS: function( elem, name, force ) {
		var ret, style = elem.style, filter;

		// IE uses filters for opacity
		if ( !jQuery.support.opacity && name === "opacity" && elem.currentStyle ) {
			ret = ropacity.test(elem.currentStyle.filter || "") ?
				(parseFloat(RegExp.$1) / 100) + "" :
				"";

			return ret === "" ?
				"1" :
				ret;
		}

		// Make sure we're using the right name for getting the float value
		if ( rfloat.test( name ) ) {
			name = styleFloat;
		}

		if ( !force && style && style[ name ] ) {
			ret = style[ name ];

		} else if ( getComputedStyle ) {

			// Only "float" is needed here
			if ( rfloat.test( name ) ) {
				name = "float";
			}

			name = name.replace( rupper, "-$1" ).toLowerCase();

			var defaultView = elem.ownerDocument.defaultView;

			if ( !defaultView ) {
				return null;
			}

			var computedStyle = defaultView.getComputedStyle( elem, null );

			if ( computedStyle ) {
				ret = computedStyle.getPropertyValue( name );
			}

			// We should always get a number back from opacity
			if ( name === "opacity" && ret === "" ) {
				ret = "1";
			}

		} else if ( elem.currentStyle ) {
			var camelCase = name.replace(rdashAlpha, fcamelCase);

			ret = elem.currentStyle[ name ] || elem.currentStyle[ camelCase ];

			// From the awesome hack by Dean Edwards
			// http://erik.eae.net/archives/2007/07/27/18.54.15/#comment-102291

			// If we're not dealing with a regular pixel number
			// but a number that has a weird ending, we need to convert it to pixels
			if ( !rnumpx.test( ret ) && rnum.test( ret ) ) {
				// Remember the original values
				var left = style.left, rsLeft = elem.runtimeStyle.left;

				// Put in the new values to get a computed value out
				elem.runtimeStyle.left = elem.currentStyle.left;
				style.left = camelCase === "fontSize" ? "1em" : (ret || 0);
				ret = style.pixelLeft + "px";

				// Revert the changed values
				style.left = left;
				elem.runtimeStyle.left = rsLeft;
			}
		}

		return ret;
	},

	// A method for quickly swapping in/out CSS properties to get correct calculations
	swap: function( elem, options, callback ) {
		var old = {};

		// Remember the old values, and insert the new ones
		for ( var name in options ) {
			old[ name ] = elem.style[ name ];
			elem.style[ name ] = options[ name ];
		}

		callback.call( elem );

		// Revert the old values
		for ( var name in options ) {
			elem.style[ name ] = old[ name ];
		}
	}
});

if ( jQuery.expr && jQuery.expr.filters ) {
	jQuery.expr.filters.hidden = function( elem ) {
		var width = elem.offsetWidth, height = elem.offsetHeight,
			skip = elem.nodeName.toLowerCase() === "tr";

		return width === 0 && height === 0 && !skip ?
			true :
			width > 0 && height > 0 && !skip ?
				false :
				jQuery.curCSS(elem, "display") === "none";
	};

	jQuery.expr.filters.visible = function( elem ) {
		return !jQuery.expr.filters.hidden( elem );
	};
}
var jsc = now(),
	rscript = /<script(.|\s)*?\/script>/gi,
	rselectTextarea = /select|textarea/i,
	rinput = /color|date|datetime|email|hidden|month|number|password|range|search|tel|text|time|url|week/i,
	jsre = /=\?(&|$)/,
	rquery = /\?/,
	rts = /(\?|&)_=.*?(&|$)/,
	rurl = /^(\w+:)?\/\/([^\/?#]+)/,
	r20 = /%20/g,

	// Keep a copy of the old load method
	_load = jQuery.fn.load;

jQuery.fn.extend({
	load: function( url, params, callback ) {
		if ( typeof url !== "string" ) {
			return _load.call( this, url );

		// Don't do a request if no elements are being requested
		} else if ( !this.length ) {
			return this;
		}

		var off = url.indexOf(" ");
		if ( off >= 0 ) {
			var selector = url.slice(off, url.length);
			url = url.slice(0, off);
		}

		// Default to a GET request
		var type = "GET";

		// If the second parameter was provided
		if ( params ) {
			// If it's a function
			if ( jQuery.isFunction( params ) ) {
				// We assume that it's the callback
				callback = params;
				params = null;

			// Otherwise, build a param string
			} else if ( typeof params === "object" ) {
				params = jQuery.param( params, jQuery.ajaxSettings.traditional );
				type = "POST";
			}
		}

		var self = this;

		// Request the remote document
		jQuery.ajax({
			url: url,
			type: type,
			dataType: "html",
			data: params,
			complete: function( res, status ) {
				// If successful, inject the HTML into all the matched elements
				if ( status === "success" || status === "notmodified" ) {
					// See if a selector was specified
					self.html( selector ?
						// Create a dummy div to hold the results
						jQuery("<div />")
							// inject the contents of the document in, removing the scripts
							// to avoid any 'Permission Denied' errors in IE
							.append(res.responseText.replace(rscript, ""))

							// Locate the specified elements
							.find(selector) :

						// If not, just inject the full result
						res.responseText );
				}

				if ( callback ) {
					self.each( callback, [res.responseText, status, res] );
				}
			}
		});

		return this;
	},

	serialize: function() {
		return jQuery.param(this.serializeArray());
	},
	serializeArray: function() {
		return this.map(function() {
			return this.elements ? jQuery.makeArray(this.elements) : this;
		})
		.filter(function() {
			return this.name && !this.disabled &&
				(this.checked || rselectTextarea.test(this.nodeName) ||
					rinput.test(this.type));
		})
		.map(function( i, elem ) {
			var val = jQuery(this).val();

			return val == null ?
				null :
				jQuery.isArray(val) ?
					jQuery.map( val, function( val, i ) {
						return { name: elem.name, value: val };
					}) :
					{ name: elem.name, value: val };
		}).get();
	}
});

// Attach a bunch of functions for handling common AJAX events
jQuery.each( "ajaxStart ajaxStop ajaxComplete ajaxError ajaxSuccess ajaxSend".split(" "), function( i, o ) {
	jQuery.fn[o] = function( f ) {
		return this.bind(o, f);
	};
});

jQuery.extend({

	get: function( url, data, callback, type ) {
		// shift arguments if data argument was omited
		if ( jQuery.isFunction( data ) ) {
			type = type || callback;
			callback = data;
			data = null;
		}

		return jQuery.ajax({
			type: "GET",
			url: url,
			data: data,
			success: callback,
			dataType: type
		});
	},

	getScript: function( url, callback ) {
		return jQuery.get(url, null, callback, "script");
	},

	getJSON: function( url, data, callback ) {
		return jQuery.get(url, data, callback, "json");
	},

	post: function( url, data, callback, type ) {
		// shift arguments if data argument was omited
		if ( jQuery.isFunction( data ) ) {
			type = type || callback;
			callback = data;
			data = {};
		}

		return jQuery.ajax({
			type: "POST",
			url: url,
			data: data,
			success: callback,
			dataType: type
		});
	},

	ajaxSetup: function( settings ) {
		jQuery.extend( jQuery.ajaxSettings, settings );
	},

	ajaxSettings: {
		url: location.href,
		global: true,
		type: "GET",
		contentType: "application/x-www-form-urlencoded",
		processData: true,
		async: true,
		/*
		timeout: 0,
		data: null,
		username: null,
		password: null,
		traditional: false,
		*/
		// Create the request object; Microsoft failed to properly
		// implement the XMLHttpRequest in IE7 (can't request local files),
		// so we use the ActiveXObject when it is available
		// This function can be overriden by calling jQuery.ajaxSetup
		xhr: window.XMLHttpRequest && (window.location.protocol !== "file:" || !window.ActiveXObject) ?
			function() {
				return new window.XMLHttpRequest();
			} :
			function() {
				try {
					return new window.ActiveXObject("Microsoft.XMLHTTP");
				} catch(e) {}
			},
		accepts: {
			xml: "application/xml, text/xml",
			html: "text/html",
			script: "text/javascript, application/javascript",
			json: "application/json, text/javascript",
			text: "text/plain",
			_default: "*/*"
		}
	},

	// Last-Modified header cache for next request
	lastModified: {},
	etag: {},

	ajax: function( origSettings ) {
		var s = jQuery.extend(true, {}, jQuery.ajaxSettings, origSettings);
		
		var jsonp, status, data,
			callbackContext = origSettings && origSettings.context || s,
			type = s.type.toUpperCase();

		// convert data if not already a string
		if ( s.data && s.processData && typeof s.data !== "string" ) {
			s.data = jQuery.param( s.data, s.traditional );
		}

		// Handle JSONP Parameter Callbacks
		if ( s.dataType === "jsonp" ) {
			if ( type === "GET" ) {
				if ( !jsre.test( s.url ) ) {
					s.url += (rquery.test( s.url ) ? "&" : "?") + (s.jsonp || "callback") + "=?";
				}
			} else if ( !s.data || !jsre.test(s.data) ) {
				s.data = (s.data ? s.data + "&" : "") + (s.jsonp || "callback") + "=?";
			}
			s.dataType = "json";
		}

		// Build temporary JSONP function
		if ( s.dataType === "json" && (s.data && jsre.test(s.data) || jsre.test(s.url)) ) {
			jsonp = s.jsonpCallback || ("jsonp" + jsc++);

			// Replace the =? sequence both in the query string and the data
			if ( s.data ) {
				s.data = (s.data + "").replace(jsre, "=" + jsonp + "$1");
			}

			s.url = s.url.replace(jsre, "=" + jsonp + "$1");

			// We need to make sure
			// that a JSONP style response is executed properly
			s.dataType = "script";

			// Handle JSONP-style loading
			window[ jsonp ] = window[ jsonp ] || function( tmp ) {
				data = tmp;
				success();
				complete();
				// Garbage collect
				window[ jsonp ] = undefined;

				try {
					delete window[ jsonp ];
				} catch(e) {}

				if ( head ) {
					head.removeChild( script );
				}
			};
		}

		if ( s.dataType === "script" && s.cache === null ) {
			s.cache = false;
		}

		if ( s.cache === false && type === "GET" ) {
			var ts = now();

			// try replacing _= if it is there
			var ret = s.url.replace(rts, "$1_=" + ts + "$2");

			// if nothing was replaced, add timestamp to the end
			s.url = ret + ((ret === s.url) ? (rquery.test(s.url) ? "&" : "?") + "_=" + ts : "");
		}

		// If data is available, append data to url for get requests
		if ( s.data && type === "GET" ) {
			s.url += (rquery.test(s.url) ? "&" : "?") + s.data;
		}

		// Watch for a new set of requests
		if ( s.global && ! jQuery.active++ ) {
			jQuery.event.trigger( "ajaxStart" );
		}

		// Matches an absolute URL, and saves the domain
		var parts = rurl.exec( s.url ),
			remote = parts && (parts[1] && parts[1] !== location.protocol || parts[2] !== location.host);

		// If we're requesting a remote document
		// and trying to load JSON or Script with a GET
		if ( s.dataType === "script" && type === "GET" && remote ) {
			var head = document.getElementsByTagName("head")[0] || document.documentElement;
			var script = document.createElement("script");
			script.src = s.url;
			if ( s.scriptCharset ) {
				script.charset = s.scriptCharset;
			}

			// Handle Script loading
			if ( !jsonp ) {
				var done = false;

				// Attach handlers for all browsers
				script.onload = script.onreadystatechange = function() {
					if ( !done && (!this.readyState ||
							this.readyState === "loaded" || this.readyState === "complete") ) {
						done = true;
						success();
						complete();

						// Handle memory leak in IE
						script.onload = script.onreadystatechange = null;
						if ( head && script.parentNode ) {
							head.removeChild( script );
						}
					}
				};
			}

			// Use insertBefore instead of appendChild  to circumvent an IE6 bug.
			// This arises when a base node is used (#2709 and #4378).
			head.insertBefore( script, head.firstChild );

			// We handle everything using the script element injection
			return undefined;
		}

		var requestDone = false;

		// Create the request object
		var xhr = s.xhr();

		if ( !xhr ) {
			return;
		}

		// Open the socket
		// Passing null username, generates a login popup on Opera (#2865)
		if ( s.username ) {
			xhr.open(type, s.url, s.async, s.username, s.password);
		} else {
			xhr.open(type, s.url, s.async);
		}

		// Need an extra try/catch for cross domain requests in Firefox 3
		try {
			// Set the correct header, if data is being sent
			if ( s.data || origSettings && origSettings.contentType ) {
				xhr.setRequestHeader("Content-Type", s.contentType);
			}

			// Set the If-Modified-Since and/or If-None-Match header, if in ifModified mode.
			if ( s.ifModified ) {
				if ( jQuery.lastModified[s.url] ) {
					xhr.setRequestHeader("If-Modified-Since", jQuery.lastModified[s.url]);
				}

				if ( jQuery.etag[s.url] ) {
					xhr.setRequestHeader("If-None-Match", jQuery.etag[s.url]);
				}
			}

			// Set header so the called script knows that it's an XMLHttpRequest
			// Only send the header if it's not a remote XHR
			if ( !remote ) {
				xhr.setRequestHeader("X-Requested-With", "XMLHttpRequest");
			}

			// Set the Accepts header for the server, depending on the dataType
			xhr.setRequestHeader("Accept", s.dataType && s.accepts[ s.dataType ] ?
				s.accepts[ s.dataType ] + ", */*" :
				s.accepts._default );
		} catch(e) {}

		// Allow custom headers/mimetypes and early abort
		if ( s.beforeSend && s.beforeSend.call(callbackContext, xhr, s) === false ) {
			// Handle the global AJAX counter
			if ( s.global && ! --jQuery.active ) {
				jQuery.event.trigger( "ajaxStop" );
			}

			// close opended socket
			xhr.abort();
			return false;
		}

		if ( s.global ) {
			trigger("ajaxSend", [xhr, s]);
		}

		// Wait for a response to come back
		var onreadystatechange = xhr.onreadystatechange = function( isTimeout ) {
			// The request was aborted
			if ( !xhr || xhr.readyState === 0 || isTimeout === "abort" ) {
				// Opera doesn't call onreadystatechange before this point
				// so we simulate the call
				if ( !requestDone ) {
					complete();
				}

				requestDone = true;
				if ( xhr ) {
					xhr.onreadystatechange = jQuery.noop;
				}

			// The transfer is complete and the data is available, or the request timed out
			} else if ( !requestDone && xhr && (xhr.readyState === 4 || isTimeout === "timeout") ) {
				requestDone = true;
				xhr.onreadystatechange = jQuery.noop;

				status = isTimeout === "timeout" ?
					"timeout" :
					!jQuery.httpSuccess( xhr ) ?
						"error" :
						s.ifModified && jQuery.httpNotModified( xhr, s.url ) ?
							"notmodified" :
							"success";

				var errMsg;

				if ( status === "success" ) {
					// Watch for, and catch, XML document parse errors
					try {
						// process the data (runs the xml through httpData regardless of callback)
						data = jQuery.httpData( xhr, s.dataType, s );
					} catch(err) {
						status = "parsererror";
						errMsg = err;
					}
				}

				// Make sure that the request was successful or notmodified
				if ( status === "success" || status === "notmodified" ) {
					// JSONP handles its own success callback
					if ( !jsonp ) {
						success();
					}
				} else {
					jQuery.handleError(s, xhr, status, errMsg);
				}

				// Fire the complete handlers
				complete();

				if ( isTimeout === "timeout" ) {
					xhr.abort();
				}

				// Stop memory leaks
				if ( s.async ) {
					xhr = null;
				}
			}
		};

		// Override the abort handler, if we can (IE doesn't allow it, but that's OK)
		// Opera doesn't fire onreadystatechange at all on abort
		try {
			var oldAbort = xhr.abort;
			xhr.abort = function() {
				if ( xhr ) {
					oldAbort.call( xhr );
				}

				onreadystatechange( "abort" );
			};
		} catch(e) { }

		// Timeout checker
		if ( s.async && s.timeout > 0 ) {
			setTimeout(function() {
				// Check to see if the request is still happening
				if ( xhr && !requestDone ) {
					onreadystatechange( "timeout" );
				}
			}, s.timeout);
		}

		// Send the data
		try {
			xhr.send( type === "POST" || type === "PUT" || type === "DELETE" ? s.data : null );
		} catch(e) {
			jQuery.handleError(s, xhr, null, e);
			// Fire the complete handlers
			complete();
		}

		// firefox 1.5 doesn't fire statechange for sync requests
		if ( !s.async ) {
			onreadystatechange();
		}

		function success() {
			// If a local callback was specified, fire it and pass it the data
			if ( s.success ) {
				s.success.call( callbackContext, data, status, xhr );
			}

			// Fire the global callback
			if ( s.global ) {
				trigger( "ajaxSuccess", [xhr, s] );
			}
		}

		function complete() {
			// Process result
			if ( s.complete ) {
				s.complete.call( callbackContext, xhr, status);
			}

			// The request was completed
			if ( s.global ) {
				trigger( "ajaxComplete", [xhr, s] );
			}

			// Handle the global AJAX counter
			if ( s.global && ! --jQuery.active ) {
				jQuery.event.trigger( "ajaxStop" );
			}
		}
		
		function trigger(type, args) {
			(s.context ? jQuery(s.context) : jQuery.event).trigger(type, args);
		}

		// return XMLHttpRequest to allow aborting the request etc.
		return xhr;
	},

	handleError: function( s, xhr, status, e ) {
		// If a local callback was specified, fire it
		if ( s.error ) {
			s.error.call( s.context || s, xhr, status, e );
		}

		// Fire the global callback
		if ( s.global ) {
			(s.context ? jQuery(s.context) : jQuery.event).trigger( "ajaxError", [xhr, s, e] );
		}
	},

	// Counter for holding the number of active queries
	active: 0,

	// Determines if an XMLHttpRequest was successful or not
	httpSuccess: function( xhr ) {
		try {
			// IE error sometimes returns 1223 when it should be 204 so treat it as success, see #1450
			return !xhr.status && location.protocol === "file:" ||
				// Opera returns 0 when status is 304
				( xhr.status >= 200 && xhr.status < 300 ) ||
				xhr.status === 304 || xhr.status === 1223 || xhr.status === 0;
		} catch(e) {}

		return false;
	},

	// Determines if an XMLHttpRequest returns NotModified
	httpNotModified: function( xhr, url ) {
		var lastModified = xhr.getResponseHeader("Last-Modified"),
			etag = xhr.getResponseHeader("Etag");

		if ( lastModified ) {
			jQuery.lastModified[url] = lastModified;
		}

		if ( etag ) {
			jQuery.etag[url] = etag;
		}

		// Opera returns 0 when status is 304
		return xhr.status === 304 || xhr.status === 0;
	},

	httpData: function( xhr, type, s ) {
		var ct = xhr.getResponseHeader("content-type") || "",
			xml = type === "xml" || !type && ct.indexOf("xml") >= 0,
			data = xml ? xhr.responseXML : xhr.responseText;

		if ( xml && data.documentElement.nodeName === "parsererror" ) {
			jQuery.error( "parsererror" );
		}

		// Allow a pre-filtering function to sanitize the response
		// s is checked to keep backwards compatibility
		if ( s && s.dataFilter ) {
			data = s.dataFilter( data, type );
		}

		// The filter can actually parse the response
		if ( typeof data === "string" ) {
			// Get the JavaScript object, if JSON is used.
			if ( type === "json" || !type && ct.indexOf("json") >= 0 ) {
				data = jQuery.parseJSON( data );

			// If the type is "script", eval it in global context
			} else if ( type === "script" || !type && ct.indexOf("javascript") >= 0 ) {
				jQuery.globalEval( data );
			}
		}

		return data;
	},

	// Serialize an array of form elements or a set of
	// key/values into a query string
	param: function( a, traditional ) {
		var s = [];
		
		// Set traditional to true for jQuery <= 1.3.2 behavior.
		if ( traditional === undefined ) {
			traditional = jQuery.ajaxSettings.traditional;
		}
		
		// If an array was passed in, assume that it is an array of form elements.
		if ( jQuery.isArray(a) || a.jquery ) {
			// Serialize the form elements
			jQuery.each( a, function() {
				add( this.name, this.value );
			});
			
		} else {
			// If traditional, encode the "old" way (the way 1.3.2 or older
			// did it), otherwise encode params recursively.
			for ( var prefix in a ) {
				buildParams( prefix, a[prefix] );
			}
		}

		// Return the resulting serialization
		return s.join("&").replace(r20, "+");

		function buildParams( prefix, obj ) {
			if ( jQuery.isArray(obj) ) {
				// Serialize array item.
				jQuery.each( obj, function( i, v ) {
					if ( traditional || /\[\]$/.test( prefix ) ) {
						// Treat each array item as a scalar.
						add( prefix, v );
					} else {
						// If array item is non-scalar (array or object), encode its
						// numeric index to resolve deserialization ambiguity issues.
						// Note that rack (as of 1.0.0) can't currently deserialize
						// nested arrays properly, and attempting to do so may cause
						// a server error. Possible fixes are to modify rack's
						// deserialization algorithm or to provide an option or flag
						// to force array serialization to be shallow.
						buildParams( prefix + "[" + ( typeof v === "object" || jQuery.isArray(v) ? i : "" ) + "]", v );
					}
				});
					
			} else if ( !traditional && obj != null && typeof obj === "object" ) {
				// Serialize object item.
				jQuery.each( obj, function( k, v ) {
					buildParams( prefix + "[" + k + "]", v );
				});
					
			} else {
				// Serialize scalar item.
				add( prefix, obj );
			}
		}

		function add( key, value ) {
			// If value is a function, invoke it and return its value
			value = jQuery.isFunction(value) ? value() : value;
			s[ s.length ] = encodeURIComponent(key) + "=" + encodeURIComponent(value);
		}
	}
});
var elemdisplay = {},
	rfxtypes = /toggle|show|hide/,
	rfxnum = /^([+-]=)?([\d+-.]+)(.*)$/,
	timerId,
	fxAttrs = [
		// height animations
		[ "height", "marginTop", "marginBottom", "paddingTop", "paddingBottom" ],
		// width animations
		[ "width", "marginLeft", "marginRight", "paddingLeft", "paddingRight" ],
		// opacity animations
		[ "opacity" ]
	];

jQuery.fn.extend({
	show: function( speed, callback ) {
		if ( speed || speed === 0) {
			return this.animate( genFx("show", 3), speed, callback);

		} else {
			for ( var i = 0, l = this.length; i < l; i++ ) {
				var old = jQuery.data(this[i], "olddisplay");

				this[i].style.display = old || "";

				if ( jQuery.css(this[i], "display") === "none" ) {
					var nodeName = this[i].nodeName, display;

					if ( elemdisplay[ nodeName ] ) {
						display = elemdisplay[ nodeName ];

					} else {
						var elem = jQuery("<" + nodeName + " />").appendTo("body");

						display = elem.css("display");

						if ( display === "none" ) {
							display = "block";
						}

						elem.remove();

						elemdisplay[ nodeName ] = display;
					}

					jQuery.data(this[i], "olddisplay", display);
				}
			}

			// Set the display of the elements in a second loop
			// to avoid the constant reflow
			for ( var j = 0, k = this.length; j < k; j++ ) {
				this[j].style.display = jQuery.data(this[j], "olddisplay") || "";
			}

			return this;
		}
	},

	hide: function( speed, callback ) {
		if ( speed || speed === 0 ) {
			return this.animate( genFx("hide", 3), speed, callback);

		} else {
			for ( var i = 0, l = this.length; i < l; i++ ) {
				var old = jQuery.data(this[i], "olddisplay");
				if ( !old && old !== "none" ) {
					jQuery.data(this[i], "olddisplay", jQuery.css(this[i], "display"));
				}
			}

			// Set the display of the elements in a second loop
			// to avoid the constant reflow
			for ( var j = 0, k = this.length; j < k; j++ ) {
				this[j].style.display = "none";
			}

			return this;
		}
	},

	// Save the old toggle function
	_toggle: jQuery.fn.toggle,

	toggle: function( fn, fn2 ) {
		var bool = typeof fn === "boolean";

		if ( jQuery.isFunction(fn) && jQuery.isFunction(fn2) ) {
			this._toggle.apply( this, arguments );

		} else if ( fn == null || bool ) {
			this.each(function() {
				var state = bool ? fn : jQuery(this).is(":hidden");
				jQuery(this)[ state ? "show" : "hide" ]();
			});

		} else {
			this.animate(genFx("toggle", 3), fn, fn2);
		}

		return this;
	},

	fadeTo: function( speed, to, callback ) {
		return this.filter(":hidden").css("opacity", 0).show().end()
					.animate({opacity: to}, speed, callback);
	},

	animate: function( prop, speed, easing, callback ) {
		var optall = jQuery.speed(speed, easing, callback);

		if ( jQuery.isEmptyObject( prop ) ) {
			return this.each( optall.complete );
		}

		return this[ optall.queue === false ? "each" : "queue" ](function() {
			var opt = jQuery.extend({}, optall), p,
				hidden = this.nodeType === 1 && jQuery(this).is(":hidden"),
				self = this;

			for ( p in prop ) {
				var name = p.replace(rdashAlpha, fcamelCase);

				if ( p !== name ) {
					prop[ name ] = prop[ p ];
					delete prop[ p ];
					p = name;
				}

				if ( prop[p] === "hide" && hidden || prop[p] === "show" && !hidden ) {
					return opt.complete.call(this);
				}

				if ( ( p === "height" || p === "width" ) && this.style ) {
					// Store display property
					opt.display = jQuery.css(this, "display");

					// Make sure that nothing sneaks out
					opt.overflow = this.style.overflow;
				}

				if ( jQuery.isArray( prop[p] ) ) {
					// Create (if needed) and add to specialEasing
					(opt.specialEasing = opt.specialEasing || {})[p] = prop[p][1];
					prop[p] = prop[p][0];
				}
			}

			if ( opt.overflow != null ) {
				this.style.overflow = "hidden";
			}

			opt.curAnim = jQuery.extend({}, prop);

			jQuery.each( prop, function( name, val ) {
				var e = new jQuery.fx( self, opt, name );

				if ( rfxtypes.test(val) ) {
					e[ val === "toggle" ? hidden ? "show" : "hide" : val ]( prop );

				} else {
					var parts = rfxnum.exec(val),
						start = e.cur(true) || 0;

					if ( parts ) {
						var end = parseFloat( parts[2] ),
							unit = parts[3] || "px";

						// We need to compute starting value
						if ( unit !== "px" ) {
							self.style[ name ] = (end || 1) + unit;
							start = ((end || 1) / e.cur(true)) * start;
							self.style[ name ] = start + unit;
						}

						// If a +=/-= token was provided, we're doing a relative animation
						if ( parts[1] ) {
							end = ((parts[1] === "-=" ? -1 : 1) * end) + start;
						}

						e.custom( start, end, unit );

					} else {
						e.custom( start, val, "" );
					}
				}
			});

			// For JS strict compliance
			return true;
		});
	},

	stop: function( clearQueue, gotoEnd ) {
		var timers = jQuery.timers;

		if ( clearQueue ) {
			this.queue([]);
		}

		this.each(function() {
			// go in reverse order so anything added to the queue during the loop is ignored
			for ( var i = timers.length - 1; i >= 0; i-- ) {
				if ( timers[i].elem === this ) {
					if (gotoEnd) {
						// force the next step to be the last
						timers[i](true);
					}

					timers.splice(i, 1);
				}
			}
		});

		// start the next in the queue if the last step wasn't forced
		if ( !gotoEnd ) {
			this.dequeue();
		}

		return this;
	}

});

// Generate shortcuts for custom animations
jQuery.each({
	slideDown: genFx("show", 1),
	slideUp: genFx("hide", 1),
	slideToggle: genFx("toggle", 1),
	fadeIn: { opacity: "show" },
	fadeOut: { opacity: "hide" }
}, function( name, props ) {
	jQuery.fn[ name ] = function( speed, callback ) {
		return this.animate( props, speed, callback );
	};
});

jQuery.extend({
	speed: function( speed, easing, fn ) {
		var opt = speed && typeof speed === "object" ? speed : {
			complete: fn || !fn && easing ||
				jQuery.isFunction( speed ) && speed,
			duration: speed,
			easing: fn && easing || easing && !jQuery.isFunction(easing) && easing
		};

		opt.duration = jQuery.fx.off ? 0 : typeof opt.duration === "number" ? opt.duration :
			jQuery.fx.speeds[opt.duration] || jQuery.fx.speeds._default;

		// Queueing
		opt.old = opt.complete;
		opt.complete = function() {
			if ( opt.queue !== false ) {
				jQuery(this).dequeue();
			}
			if ( jQuery.isFunction( opt.old ) ) {
				opt.old.call( this );
			}
		};

		return opt;
	},

	easing: {
		linear: function( p, n, firstNum, diff ) {
			return firstNum + diff * p;
		},
		swing: function( p, n, firstNum, diff ) {
			return ((-Math.cos(p*Math.PI)/2) + 0.5) * diff + firstNum;
		}
	},

	timers: [],

	fx: function( elem, options, prop ) {
		this.options = options;
		this.elem = elem;
		this.prop = prop;

		if ( !options.orig ) {
			options.orig = {};
		}
	}

});

jQuery.fx.prototype = {
	// Simple function for setting a style value
	update: function() {
		if ( this.options.step ) {
			this.options.step.call( this.elem, this.now, this );
		}

		(jQuery.fx.step[this.prop] || jQuery.fx.step._default)( this );

		// Set display property to block for height/width animations
		if ( ( this.prop === "height" || this.prop === "width" ) && this.elem.style ) {
			this.elem.style.display = "block";
		}
	},

	// Get the current size
	cur: function( force ) {
		if ( this.elem[this.prop] != null && (!this.elem.style || this.elem.style[this.prop] == null) ) {
			return this.elem[ this.prop ];
		}

		var r = parseFloat(jQuery.css(this.elem, this.prop, force));
		return r && r > -10000 ? r : parseFloat(jQuery.curCSS(this.elem, this.prop)) || 0;
	},

	// Start an animation from one number to another
	custom: function( from, to, unit ) {
		this.startTime = now();
		this.start = from;
		this.end = to;
		this.unit = unit || this.unit || "px";
		this.now = this.start;
		this.pos = this.state = 0;

		var self = this;
		function t( gotoEnd ) {
			return self.step(gotoEnd);
		}

		t.elem = this.elem;

		if ( t() && jQuery.timers.push(t) && !timerId ) {
			timerId = setInterval(jQuery.fx.tick, 13);
		}
	},

	// Simple 'show' function
	show: function() {
		// Remember where we started, so that we can go back to it later
		this.options.orig[this.prop] = jQuery.style( this.elem, this.prop );
		this.options.show = true;

		// Begin the animation
		// Make sure that we start at a small width/height to avoid any
		// flash of content
		this.custom(this.prop === "width" || this.prop === "height" ? 1 : 0, this.cur());

		// Start by showing the element
		jQuery( this.elem ).show();
	},

	// Simple 'hide' function
	hide: function() {
		// Remember where we started, so that we can go back to it later
		this.options.orig[this.prop] = jQuery.style( this.elem, this.prop );
		this.options.hide = true;

		// Begin the animation
		this.custom(this.cur(), 0);
	},

	// Each step of an animation
	step: function( gotoEnd ) {
		var t = now(), done = true;

		if ( gotoEnd || t >= this.options.duration + this.startTime ) {
			this.now = this.end;
			this.pos = this.state = 1;
			this.update();

			this.options.curAnim[ this.prop ] = true;

			for ( var i in this.options.curAnim ) {
				if ( this.options.curAnim[i] !== true ) {
					done = false;
				}
			}

			if ( done ) {
				if ( this.options.display != null ) {
					// Reset the overflow
					this.elem.style.overflow = this.options.overflow;

					// Reset the display
					var old = jQuery.data(this.elem, "olddisplay");
					this.elem.style.display = old ? old : this.options.display;

					if ( jQuery.css(this.elem, "display") === "none" ) {
						this.elem.style.display = "block";
					}
				}

				// Hide the element if the "hide" operation was done
				if ( this.options.hide ) {
					jQuery(this.elem).hide();
				}

				// Reset the properties, if the item has been hidden or shown
				if ( this.options.hide || this.options.show ) {
					for ( var p in this.options.curAnim ) {
						jQuery.style(this.elem, p, this.options.orig[p]);
					}
				}

				// Execute the complete function
				this.options.complete.call( this.elem );
			}

			return false;

		} else {
			var n = t - this.startTime;
			this.state = n / this.options.duration;

			// Perform the easing function, defaults to swing
			var specialEasing = this.options.specialEasing && this.options.specialEasing[this.prop];
			var defaultEasing = this.options.easing || (jQuery.easing.swing ? "swing" : "linear");
			this.pos = jQuery.easing[specialEasing || defaultEasing](this.state, n, 0, 1, this.options.duration);
			this.now = this.start + ((this.end - this.start) * this.pos);

			// Perform the next step of the animation
			this.update();
		}

		return true;
	}
};

jQuery.extend( jQuery.fx, {
	tick: function() {
		var timers = jQuery.timers;

		for ( var i = 0; i < timers.length; i++ ) {
			if ( !timers[i]() ) {
				timers.splice(i--, 1);
			}
		}

		if ( !timers.length ) {
			jQuery.fx.stop();
		}
	},
		
	stop: function() {
		clearInterval( timerId );
		timerId = null;
	},
	
	speeds: {
		slow: 600,
 		fast: 200,
 		// Default speed
 		_default: 400
	},

	step: {
		opacity: function( fx ) {
			jQuery.style(fx.elem, "opacity", fx.now);
		},

		_default: function( fx ) {
			if ( fx.elem.style && fx.elem.style[ fx.prop ] != null ) {
				fx.elem.style[ fx.prop ] = (fx.prop === "width" || fx.prop === "height" ? Math.max(0, fx.now) : fx.now) + fx.unit;
			} else {
				fx.elem[ fx.prop ] = fx.now;
			}
		}
	}
});

if ( jQuery.expr && jQuery.expr.filters ) {
	jQuery.expr.filters.animated = function( elem ) {
		return jQuery.grep(jQuery.timers, function( fn ) {
			return elem === fn.elem;
		}).length;
	};
}

function genFx( type, num ) {
	var obj = {};

	jQuery.each( fxAttrs.concat.apply([], fxAttrs.slice(0,num)), function() {
		obj[ this ] = type;
	});

	return obj;
}
if ( "getBoundingClientRect" in document.documentElement ) {
	jQuery.fn.offset = function( options ) {
		var elem = this[0];

		if ( options ) { 
			return this.each(function( i ) {
				jQuery.offset.setOffset( this, options, i );
			});
		}

		if ( !elem || !elem.ownerDocument ) {
			return null;
		}

		if ( elem === elem.ownerDocument.body ) {
			return jQuery.offset.bodyOffset( elem );
		}

		var box = elem.getBoundingClientRect(), doc = elem.ownerDocument, body = doc.body, docElem = doc.documentElement,
			clientTop = docElem.clientTop || body.clientTop || 0, clientLeft = docElem.clientLeft || body.clientLeft || 0,
			top  = box.top  + (self.pageYOffset || jQuery.support.boxModel && docElem.scrollTop  || body.scrollTop ) - clientTop,
			left = box.left + (self.pageXOffset || jQuery.support.boxModel && docElem.scrollLeft || body.scrollLeft) - clientLeft;

		return { top: top, left: left };
	};

} else {
	jQuery.fn.offset = function( options ) {
		var elem = this[0];

		if ( options ) { 
			return this.each(function( i ) {
				jQuery.offset.setOffset( this, options, i );
			});
		}

		if ( !elem || !elem.ownerDocument ) {
			return null;
		}

		if ( elem === elem.ownerDocument.body ) {
			return jQuery.offset.bodyOffset( elem );
		}

		jQuery.offset.initialize();

		var offsetParent = elem.offsetParent, prevOffsetParent = elem,
			doc = elem.ownerDocument, computedStyle, docElem = doc.documentElement,
			body = doc.body, defaultView = doc.defaultView,
			prevComputedStyle = defaultView ? defaultView.getComputedStyle( elem, null ) : elem.currentStyle,
			top = elem.offsetTop, left = elem.offsetLeft;

		while ( (elem = elem.parentNode) && elem !== body && elem !== docElem ) {
			if ( jQuery.offset.supportsFixedPosition && prevComputedStyle.position === "fixed" ) {
				break;
			}

			computedStyle = defaultView ? defaultView.getComputedStyle(elem, null) : elem.currentStyle;
			top  -= elem.scrollTop;
			left -= elem.scrollLeft;

			if ( elem === offsetParent ) {
				top  += elem.offsetTop;
				left += elem.offsetLeft;

				if ( jQuery.offset.doesNotAddBorder && !(jQuery.offset.doesAddBorderForTableAndCells && /^t(able|d|h)$/i.test(elem.nodeName)) ) {
					top  += parseFloat( computedStyle.borderTopWidth  ) || 0;
					left += parseFloat( computedStyle.borderLeftWidth ) || 0;
				}

				prevOffsetParent = offsetParent, offsetParent = elem.offsetParent;
			}

			if ( jQuery.offset.subtractsBorderForOverflowNotVisible && computedStyle.overflow !== "visible" ) {
				top  += parseFloat( computedStyle.borderTopWidth  ) || 0;
				left += parseFloat( computedStyle.borderLeftWidth ) || 0;
			}

			prevComputedStyle = computedStyle;
		}

		if ( prevComputedStyle.position === "relative" || prevComputedStyle.position === "static" ) {
			top  += body.offsetTop;
			left += body.offsetLeft;
		}

		if ( jQuery.offset.supportsFixedPosition && prevComputedStyle.position === "fixed" ) {
			top  += Math.max( docElem.scrollTop, body.scrollTop );
			left += Math.max( docElem.scrollLeft, body.scrollLeft );
		}

		return { top: top, left: left };
	};
}

jQuery.offset = {
	initialize: function() {
		var body = document.body, container = document.createElement("div"), innerDiv, checkDiv, table, td, bodyMarginTop = parseFloat( jQuery.curCSS(body, "marginTop", true) ) || 0,
			html = "<div style='position:absolute;top:0;left:0;margin:0;border:5px solid #000;padding:0;width:1px;height:1px;'><div></div></div><table style='position:absolute;top:0;left:0;margin:0;border:5px solid #000;padding:0;width:1px;height:1px;' cellpadding='0' cellspacing='0'><tr><td></td></tr></table>";

		jQuery.extend( container.style, { position: "absolute", top: 0, left: 0, margin: 0, border: 0, width: "1px", height: "1px", visibility: "hidden" } );

		container.innerHTML = html;
		body.insertBefore( container, body.firstChild );
		innerDiv = container.firstChild;
		checkDiv = innerDiv.firstChild;
		td = innerDiv.nextSibling.firstChild.firstChild;

		this.doesNotAddBorder = (checkDiv.offsetTop !== 5);
		this.doesAddBorderForTableAndCells = (td.offsetTop === 5);

		checkDiv.style.position = "fixed", checkDiv.style.top = "20px";
		// safari subtracts parent border width here which is 5px
		this.supportsFixedPosition = (checkDiv.offsetTop === 20 || checkDiv.offsetTop === 15);
		checkDiv.style.position = checkDiv.style.top = "";

		innerDiv.style.overflow = "hidden", innerDiv.style.position = "relative";
		this.subtractsBorderForOverflowNotVisible = (checkDiv.offsetTop === -5);

		this.doesNotIncludeMarginInBodyOffset = (body.offsetTop !== bodyMarginTop);

		body.removeChild( container );
		body = container = innerDiv = checkDiv = table = td = null;
		jQuery.offset.initialize = jQuery.noop;
	},

	bodyOffset: function( body ) {
		var top = body.offsetTop, left = body.offsetLeft;

		jQuery.offset.initialize();

		if ( jQuery.offset.doesNotIncludeMarginInBodyOffset ) {
			top  += parseFloat( jQuery.curCSS(body, "marginTop",  true) ) || 0;
			left += parseFloat( jQuery.curCSS(body, "marginLeft", true) ) || 0;
		}

		return { top: top, left: left };
	},
	
	setOffset: function( elem, options, i ) {
		// set position first, in-case top/left are set even on static elem
		if ( /static/.test( jQuery.curCSS( elem, "position" ) ) ) {
			elem.style.position = "relative";
		}
		var curElem   = jQuery( elem ),
			curOffset = curElem.offset(),
			curTop    = parseInt( jQuery.curCSS( elem, "top",  true ), 10 ) || 0,
			curLeft   = parseInt( jQuery.curCSS( elem, "left", true ), 10 ) || 0;

		if ( jQuery.isFunction( options ) ) {
			options = options.call( elem, i, curOffset );
		}

		var props = {
			top:  (options.top  - curOffset.top)  + curTop,
			left: (options.left - curOffset.left) + curLeft
		};
		
		if ( "using" in options ) {
			options.using.call( elem, props );
		} else {
			curElem.css( props );
		}
	}
};


jQuery.fn.extend({
	position: function() {
		if ( !this[0] ) {
			return null;
		}

		var elem = this[0],

		// Get *real* offsetParent
		offsetParent = this.offsetParent(),

		// Get correct offsets
		offset       = this.offset(),
		parentOffset = /^body|html$/i.test(offsetParent[0].nodeName) ? { top: 0, left: 0 } : offsetParent.offset();

		// Subtract element margins
		// note: when an element has margin: auto the offsetLeft and marginLeft
		// are the same in Safari causing offset.left to incorrectly be 0
		offset.top  -= parseFloat( jQuery.curCSS(elem, "marginTop",  true) ) || 0;
		offset.left -= parseFloat( jQuery.curCSS(elem, "marginLeft", true) ) || 0;

		// Add offsetParent borders
		parentOffset.top  += parseFloat( jQuery.curCSS(offsetParent[0], "borderTopWidth",  true) ) || 0;
		parentOffset.left += parseFloat( jQuery.curCSS(offsetParent[0], "borderLeftWidth", true) ) || 0;

		// Subtract the two offsets
		return {
			top:  offset.top  - parentOffset.top,
			left: offset.left - parentOffset.left
		};
	},

	offsetParent: function() {
		return this.map(function() {
			var offsetParent = this.offsetParent || document.body;
			while ( offsetParent && (!/^body|html$/i.test(offsetParent.nodeName) && jQuery.css(offsetParent, "position") === "static") ) {
				offsetParent = offsetParent.offsetParent;
			}
			return offsetParent;
		});
	}
});


// Create scrollLeft and scrollTop methods
jQuery.each( ["Left", "Top"], function( i, name ) {
	var method = "scroll" + name;

	jQuery.fn[ method ] = function(val) {
		var elem = this[0], win;
		
		if ( !elem ) {
			return null;
		}

		if ( val !== undefined ) {
			// Set the scroll offset
			return this.each(function() {
				win = getWindow( this );

				if ( win ) {
					win.scrollTo(
						!i ? val : jQuery(win).scrollLeft(),
						 i ? val : jQuery(win).scrollTop()
					);

				} else {
					this[ method ] = val;
				}
			});
		} else {
			win = getWindow( elem );

			// Return the scroll offset
			return win ? ("pageXOffset" in win) ? win[ i ? "pageYOffset" : "pageXOffset" ] :
				jQuery.support.boxModel && win.document.documentElement[ method ] ||
					win.document.body[ method ] :
				elem[ method ];
		}
	};
});

function getWindow( elem ) {
	return ("scrollTo" in elem && elem.document) ?
		elem :
		elem.nodeType === 9 ?
			elem.defaultView || elem.parentWindow :
			false;
}
// Create innerHeight, innerWidth, outerHeight and outerWidth methods
jQuery.each([ "Height", "Width" ], function( i, name ) {

	var type = name.toLowerCase();

	// innerHeight and innerWidth
	jQuery.fn["inner" + name] = function() {
		return this[0] ?
			jQuery.css( this[0], type, false, "padding" ) :
			null;
	};

	// outerHeight and outerWidth
	jQuery.fn["outer" + name] = function( margin ) {
		return this[0] ?
			jQuery.css( this[0], type, false, margin ? "margin" : "border" ) :
			null;
	};

	jQuery.fn[ type ] = function( size ) {
		// Get window width or height
		var elem = this[0];
		if ( !elem ) {
			return size == null ? null : this;
		}
		
		if ( jQuery.isFunction( size ) ) {
			return this.each(function( i ) {
				var self = jQuery( this );
				self[ type ]( size.call( this, i, self[ type ]() ) );
			});
		}

		return ("scrollTo" in elem && elem.document) ? // does it walk and quack like a window?
			// Everyone else use document.documentElement or document.body depending on Quirks vs Standards mode
			elem.document.compatMode === "CSS1Compat" && elem.document.documentElement[ "client" + name ] ||
			elem.document.body[ "client" + name ] :

			// Get document width or height
			(elem.nodeType === 9) ? // is it a document
				// Either scroll[Width/Height] or offset[Width/Height], whichever is greater
				Math.max(
					elem.documentElement["client" + name],
					elem.body["scroll" + name], elem.documentElement["scroll" + name],
					elem.body["offset" + name], elem.documentElement["offset" + name]
				) :

				// Get or set width or height on the element
				size === undefined ?
					// Get width or height on the element
					jQuery.css( elem, type ) :

					// Set the width or height on the element (default to pixels if value is unitless)
					this.css( type, typeof size === "string" ? size : size + "px" );
	};

});
// Expose jQuery to the global object
window.jQuery = window.$ = jQuery;

})(window);
```
