## Supplementary material 1 for "Termite diet rather than geographical origin determines the microbiome composition and functional genetic structure of nests from South American and African representatives, as revealed by a multiomics approach": jquery-1.4.2.min.js

```
/*
* jQuery JavaScript Library v1.4.2
* Copyright 2010, John Resig, http://jquery.com/
* Distributed in whole under the terms of the MIT license
* http://www.opensource.org/licenses/mit-license.php 
* 
* Includes Sizzle.js
* http://sizzlejs.com/
* Copyright 2010, The Dojo Foundation
* Released under the MIT, BSD, and GPL Licenses.
*/
(function(A, w) {
    function ma() { if (!c.isReady) { try { s.documentElement.doScroll("left") } catch (a) { setTimeout(ma, 1); return } c.ready() } } function Qa(a, b) { b.src ? c.ajax({ url: b.src, async: false, dataType: "script" }) : c.globalEval(b.text || b.textContent || b.innerHTML || ""); b.parentNode && b.parentNode.removeChild(b) } function X(a, b, d, f, e, j) {
        var i = a.length; if (typeof b === "object") { for (var o in b) X(a, o, b[o], f, e, d); return a } if (d !== w) { f = !j && f && c.isFunction(d); for (o = 0; o < i; o++) e(a[o], b, f ? d.call(a[o], o, e(a[o], b)) : d, j); return a } return i ?
e(a[0],b):w}function J(){return(new Date).getTime()}function Y(){return false}function Z(){return true}function na(a,b,d){d[0].type=a;return c.event.handle.apply(b,d)}function oa(a){var b,d=[],f=[],e=arguments,j,i,o,k,n,r;i=c.data(this,"events");if(!(a.liveFired===this||!i||!i.live||a.button&&a.type==="click")){a.liveFired=this;var u=i.live.slice(0);for(k=0;k<u.length;k++){i=u[k];i.origType.replace(O,"")===a.type?f.push(i.selector):u.splice(k--,1)}j=c(a.target).closest(f,a.currentTarget);n=0;for(r=
j.length;n<r;n++)for(k=0;k<u.length;k++){i=u[k];if(j[n].selector===i.selector){o=j[n].elem;f=null;if(i.preType==="mouseenter"||i.preType==="mouseleave")f=c(a.relatedTarget).closest(i.selector)[0];if(!f||f!==o)d.push({elem:o,handleObj:i})}}n=0;for(r=d.length;n<r;n++){j=d[n];a.currentTarget=j.elem;a.data=j.handleObj.data;a.handleObj=j.handleObj;if(j.handleObj.origHandler.apply(j.elem,e)===false){b=false;break}}return b}}function pa(a,b){return"live."+(a&&a!=="*"?a+".":"")+b.replace(/\./g,"`").replace(/ /g,
"&")}function qa(a){return!a||!a.parentNode||a.parentNode.nodeType===11}function ra(a,b){var d=0;b.each(function(){if(this.nodeName===(a[d]&&a[d].nodeName)){var f=c.data(a[d++]),e=c.data(this,f);if(f=f&&f.events){delete e.handle;e.events={};for(var j in f)for(var i in f[j])c.event.add(this,j,f[j][i],f[j][i].data)}}})}function sa(a,b,d){var f,e,j;b=b&&b[0]?b[0].ownerDocument||b[0]:s;if(a.length===1&&typeof a[0]==="string"&&a[0].length<512&&b===s&&!ta.test(a[0])&&(c.support.checkClone||!ua.test(a[0]))){e=
true;if(j=c.fragments[a[0]])if(j!==1)f=j}if(!f){f=b.createDocumentFragment();c.clean(a,b,f,d)}if(e)c.fragments[a[0]]=j?f:1;return{fragment:f,cacheable:e}}function K(a,b){var d={};c.each(va.concat.apply([],va.slice(0,b)),function(){d[this]=a});return d}function wa(a){return"scrollTo"in a&&a.document?a:a.nodeType===9?a.defaultView||a.parentWindow:false}var c=function(a,b){return new c.fn.init(a,b)},Ra=A.jQuery,Sa=A.$,s=A.document,T,Ta=/^[^<]*(<[\w\W]+>)[^>]*$|^#([\w-]+)$/,Ua=/^.[^:#\[\.,]*$/,Va=/\S/,
Wa=/^(\s|\u00A0)+|(\s|\u00A0)+$/g,Xa=/^<(\w+)\s*\/?>(?:<\/\1>)?$/,P=navigator.userAgent,xa=false,Q=[],L,$=Object.prototype.toString,aa=Object.prototype.hasOwnProperty,ba=Array.prototype.push,R=Array.prototype.slice,ya=Array.prototype.indexOf;c.fn=c.prototype={init:function(a,b){var d,f;if(!a)return this;if(a.nodeType){this.context=this[0]=a;this.length=1;return this}if(a==="body"&&!b){this.context=s;this[0]=s.body;this.selector="body";this.length=1;return this}if(typeof a==="string")if((d=Ta.exec(a))&&
(d[1]||!b))if(d[1]){f=b?b.ownerDocument||b:s;if(a=Xa.exec(a))if(c.isPlainObject(b)){a=[s.createElement(a[1])];c.fn.attr.call(a,b,true)}else a=[f.createElement(a[1])];else{a=sa([d[1]],[f]);a=(a.cacheable?a.fragment.cloneNode(true):a.fragment).childNodes}return c.merge(this,a)}else{if(b=s.getElementById(d[2])){if(b.id!==d[2])return T.find(a);this.length=1;this[0]=b}this.context=s;this.selector=a;return this}else if(!b&&/^\w+$/.test(a)){this.selector=a;this.context=s;a=s.getElementsByTagName(a);return c.merge(this,
a)}else return!b||b.jquery?(b||T).find(a):c(b).find(a);else if(c.isFunction(a))return T.ready(a);if(a.selector!==w){this.selector=a.selector;this.context=a.context}return c.makeArray(a,this)},selector:"",jquery:"1.4.2",length:0,size:function(){return this.length},toArray:function(){return R.call(this,0)},get:function(a){return a==null?this.toArray():a<0?this.slice(a)[0]:this[a]},pushStack:function(a,b,d){var f=c();c.isArray(a)?ba.apply(f,a):c.merge(f,a);f.prevObject=this;f.context=this.context;if(b===
"find")f.selector=this.selector+(this.selector?" ":"")+d;else if(b)f.selector=this.selector+"."+b+"("+d+")";return f},each:function(a,b){return c.each(this,a,b)},ready:function(a){c.bindReady();if(c.isReady)a.call(s,c);else Q&&Q.push(a);return this},eq:function(a){return a===-1?this.slice(a):this.slice(a,+a+1)},first:function(){return this.eq(0)},last:function(){return this.eq(-1)},slice:function(){return this.pushStack(R.apply(this,arguments),"slice",R.call(arguments).join(","))},map:function(a){return this.pushStack(c.map(this,
function(b,d){return a.call(b,d,b)}))},end:function(){return this.prevObject||c(null)},push:ba,sort:[].sort,splice:[].splice};c.fn.init.prototype=c.fn;c.extend=c.fn.extend=function(){var a=arguments[0]||{},b=1,d=arguments.length,f=false,e,j,i,o;if(typeof a==="boolean"){f=a;a=arguments[1]||{};b=2}if(typeof a!=="object"&&!c.isFunction(a))a={};if(d===b){a=this;--b}for(;b<d;b++)if((e=arguments[b])!=null)for(j in e){i=a[j];o=e[j];if(a!==o)if(f&&o&&(c.isPlainObject(o)||c.isArray(o))){i=i&&(c.isPlainObject(i)||
c.isArray(i))?i:c.isArray(o)?[]:{};a[j]=c.extend(f,i,o)}else if(o!==w)a[j]=o}return a};c.extend({noConflict:function(a){A.$=Sa;if(a)A.jQuery=Ra;return c},isReady:false,ready:function(){if(!c.isReady){if(!s.body)return setTimeout(c.ready,13);c.isReady=true;if(Q){for(var a,b=0;a=Q[b++];)a.call(s,c);Q=null}c.fn.triggerHandler&&c(s).triggerHandler("ready")}},bindReady:function(){if(!xa){xa=true;if(s.readyState==="complete")return c.ready();if(s.addEventListener){s.addEventListener("DOMContentLoaded",
L,false);A.addEventListener("load",c.ready,false)}else if(s.attachEvent){s.attachEvent("onreadystatechange",L);A.attachEvent("onload",c.ready);var a=false;try{a=A.frameElement==null}catch(b){}s.documentElement.doScroll&&a&&ma()}}},isFunction:function(a){return $.call(a)==="[object Function]"},isArray:function(a){return $.call(a)==="[object Array]"},isPlainObject:function(a){if(!a||$.call(a)!=="[object Object]"||a.nodeType||a.setInterval)return false;if(a.constructor&&!aa.call(a,"constructor")&&!aa.call(a.constructor.prototype,
"isPrototypeOf"))return false;var b;for(b in a);return b===w||aa.call(a,b)},isEmptyObject:function(a){for(var b in a)return false;return true},error:function(a){throw a;},parseJSON:function(a){if(typeof a!=="string"||!a)return null;a=c.trim(a);if(/^[\],:{}\s]*$/.test(a.replace(/\\(?:["\\\/bfnrt]|u[0-9a-fA-F]{4})/g,"@").replace(/"[^"\\\n\r]*"|true|false|null|-?\d+(?:\.\d*)?(?:[eE][+\-]?\d+)?/g,"]").replace(/(?:^|:|,)(?:\s*\[)+/g,"")))return A.JSON&&A.JSON.parse?A.JSON.parse(a):(new Function("return "+
a))();else c.error("Invalid JSON: "+a)},noop:function(){},globalEval:function(a){if(a&&Va.test(a)){var b=s.getElementsByTagName("head")[0]||s.documentElement,d=s.createElement("script");d.type="text/javascript";if(c.support.scriptEval)d.appendChild(s.createTextNode(a));else d.text=a;b.insertBefore(d,b.firstChild);b.removeChild(d)}},nodeName:function(a,b){return a.nodeName&&a.nodeName.toUpperCase()===b.toUpperCase()},each:function(a,b,d){var f,e=0,j=a.length,i=j===w||c.isFunction(a);if(d)if(i)for(f in a){if(b.apply(a[f],
d)===false)break}else for(;e<j;){if(b.apply(a[e++],d)===false)break}else if(i)for(f in a){if(b.call(a[f],f,a[f])===false)break}else for(d=a[0];e<j&&b.call(d,e,d)!==false;d=a[++e]);return a},trim:function(a){return(a||"").replace(Wa,"")},makeArray:function(a,b){b=b||[];if(a!=null)a.length==null||typeof a==="string"||c.isFunction(a)||typeof a!=="function"&&a.setInterval?ba.call(b,a):c.merge(b,a);return b},inArray:function(a,b){if(b.indexOf)return b.indexOf(a);for(var d=0,f=b.length;d<f;d++)if(b[d]===
a)return d;return-1},merge:function(a,b){var d=a.length,f=0;if(typeof b.length==="number")for(var e=b.length;f<e;f++)a[d++]=b[f];else for(;b[f]!==w;)a[d++]=b[f++];a.length=d;return a},grep:function(a,b,d){for(var f=[],e=0,j=a.length;e<j;e++)!d!==!b(a[e],e)&&f.push(a[e]);return f},map:function(a,b,d){for(var f=[],e,j=0,i=a.length;j<i;j++){e=b(a[j],j,d);if(e!=null)f[f.length]=e}return f.concat.apply([],f)},guid:1,proxy:function(a,b,d){if(arguments.length===2)if(typeof b==="string"){d=a;a=d[b];b=w}else if(b&&
!c.isFunction(b)){d=b;b=w}if(!b&&a)b=function(){return a.apply(d||this,arguments)};if(a)b.guid=a.guid=a.guid||b.guid||c.guid++;return b},uaMatch:function(a){a=a.toLowerCase();a=/(webkit)[ \/]([\w.]+)/.exec(a)||/(opera)(?:.*version)?[ \/]([\w.]+)/.exec(a)||/(msie) ([\w.]+)/.exec(a)||!/compatible/.test(a)&&/(mozilla)(?:.*? rv:([\w.]+))?/.exec(a)||[];return{browser:a[1]||"",version:a[2]||"0"}},browser:{}});P=c.uaMatch(P);if(P.browser){c.browser[P.browser]=true;c.browser.version=P.version}if(c.browser.webkit)c.browser.safari=
true;if(ya)c.inArray=function(a,b){return ya.call(b,a)};T=c(s);if(s.addEventListener)L=function(){s.removeEventListener("DOMContentLoaded",L,false);c.ready()};else if(s.attachEvent)L=function(){if(s.readyState==="complete"){s.detachEvent("onreadystatechange",L);c.ready()}};(function(){c.support={};var a=s.documentElement,b=s.createElement("script"),d=s.createElement("div"),f="script"+J();d.style.display="none";d.innerHTML="   <link/><table></table><a href='/a' style='color:red;float:left;opacity:.55;'>a</a><input type='checkbox'/>";
var e=d.getElementsByTagName("*"),j=d.getElementsByTagName("a")[0];if(!(!e||!e.length||!j)){c.support={leadingWhitespace:d.firstChild.nodeType===3,tbody:!d.getElementsByTagName("tbody").length,htmlSerialize:!!d.getElementsByTagName("link").length,style:/red/.test(j.getAttribute("style")),hrefNormalized:j.getAttribute("href")==="/a",opacity:/^0.55$/.test(j.style.opacity),cssFloat:!!j.style.cssFloat,checkOn:d.getElementsByTagName("input")[0].value==="on",optSelected:s.createElement("select").appendChild(s.createElement("option")).selected,
parentNode:d.removeChild(d.appendChild(s.createElement("div"))).parentNode===null,deleteExpando:true,checkClone:false,scriptEval:false,noCloneEvent:true,boxModel:null};b.type="text/javascript";try{b.appendChild(s.createTextNode("window."+f+"=1;"))}catch(i){}a.insertBefore(b,a.firstChild);if(A[f]){c.support.scriptEval=true;delete A[f]}try{delete b.test}catch(o){c.support.deleteExpando=false}a.removeChild(b);if(d.attachEvent&&d.fireEvent){d.attachEvent("onclick",function k(){c.support.noCloneEvent=
false;d.detachEvent("onclick",k)});d.cloneNode(true).fireEvent("onclick")}d=s.createElement("div");d.innerHTML="<input type='radio' name='radiotest' checked='checked'/>";a=s.createDocumentFragment();a.appendChild(d.firstChild);c.support.checkClone=a.cloneNode(true).cloneNode(true).lastChild.checked;c(function(){var k=s.createElement("div");k.style.width=k.style.paddingLeft="1px";s.body.appendChild(k);c.boxModel=c.support.boxModel=k.offsetWidth===2;s.body.removeChild(k).style.display="none"});a=function(k){var n=
s.createElement("div");k="on"+k;var r=k in n;if(!r){n.setAttribute(k,"return;");r=typeof n[k]==="function"}return r};c.support.submitBubbles=a("submit");c.support.changeBubbles=a("change");a=b=d=e=j=null}})();c.props={"for":"htmlFor","class":"className",readonly:"readOnly",maxlength:"maxLength",cellspacing:"cellSpacing",rowspan:"rowSpan",colspan:"colSpan",tabindex:"tabIndex",usemap:"useMap",frameborder:"frameBorder"};var G="jQuery"+J(),Ya=0,za={};c.extend({cache:{},expando:G,noData:{embed:true,object:true,
applet:true},data:function(a,b,d){if(!(a.nodeName&&c.noData[a.nodeName.toLowerCase()])){a=a==A?za:a;var f=a[G],e=c.cache;if(!f&&typeof b==="string"&&d===w)return null;f||(f=++Ya);if(typeof b==="object"){a[G]=f;e[f]=c.extend(true,{},b)}else if(!e[f]){a[G]=f;e[f]={}}a=e[f];if(d!==w)a[b]=d;return typeof b==="string"?a[b]:a}},removeData:function(a,b){if(!(a.nodeName&&c.noData[a.nodeName.toLowerCase()])){a=a==A?za:a;var d=a[G],f=c.cache,e=f[d];if(b){if(e){delete e[b];c.isEmptyObject(e)&&c.removeData(a)}}else{if(c.support.deleteExpando)delete a[c.expando];
else a.removeAttribute&&a.removeAttribute(c.expando);delete f[d]}}}});c.fn.extend({data:function(a,b){if(typeof a==="undefined"&&this.length)return c.data(this[0]);else if(typeof a==="object")return this.each(function(){c.data(this,a)});var d=a.split(".");d[1]=d[1]?"."+d[1]:"";if(b===w){var f=this.triggerHandler("getData"+d[1]+"!",[d[0]]);if(f===w&&this.length)f=c.data(this[0],a);return f===w&&d[1]?this.data(d[0]):f}else return this.trigger("setData"+d[1]+"!",[d[0],b]).each(function(){c.data(this,
a,b)})},removeData:function(a){return this.each(function(){c.removeData(this,a)})}});c.extend({queue:function(a,b,d){if(a){b=(b||"fx")+"queue";var f=c.data(a,b);if(!d)return f||[];if(!f||c.isArray(d))f=c.data(a,b,c.makeArray(d));else f.push(d);return f}},dequeue:function(a,b){b=b||"fx";var d=c.queue(a,b),f=d.shift();if(f==="inprogress")f=d.shift();if(f){b==="fx"&&d.unshift("inprogress");f.call(a,function(){c.dequeue(a,b)})}}});c.fn.extend({queue:function(a,b){if(typeof a!=="string"){b=a;a="fx"}if(b===
w)return c.queue(this[0],a);return this.each(function(){var d=c.queue(this,a,b);a==="fx"&&d[0]!=="inprogress"&&c.dequeue(this,a)})},dequeue:function(a){return this.each(function(){c.dequeue(this,a)})},delay:function(a,b){a=c.fx?c.fx.speeds[a]||a:a;b=b||"fx";return this.queue(b,function(){var d=this;setTimeout(function(){c.dequeue(d,b)},a)})},clearQueue:function(a){return this.queue(a||"fx",[])}});var Aa=/[\n\t]/g,ca=/\s+/,Za=/\r/g,$a=/href|src|style/,ab=/(button|input)/i,bb=/(button|input|object|select|textarea)/i,
cb=/^(a|area)$/i,Ba=/radio|checkbox/;c.fn.extend({attr:function(a,b){return X(this,a,b,true,c.attr)},removeAttr:function(a){return this.each(function(){c.attr(this,a,"");this.nodeType===1&&this.removeAttribute(a)})},addClass:function(a){if(c.isFunction(a))return this.each(function(n){var r=c(this);r.addClass(a.call(this,n,r.attr("class")))});if(a&&typeof a==="string")for(var b=(a||"").split(ca),d=0,f=this.length;d<f;d++){var e=this[d];if(e.nodeType===1)if(e.className){for(var j=" "+e.className+" ",
i=e.className,o=0,k=b.length;o<k;o++)if(j.indexOf(" "+b[o]+" ")<0)i+=" "+b[o];e.className=c.trim(i)}else e.className=a}return this},removeClass:function(a){if(c.isFunction(a))return this.each(function(k){var n=c(this);n.removeClass(a.call(this,k,n.attr("class")))});if(a&&typeof a==="string"||a===w)for(var b=(a||"").split(ca),d=0,f=this.length;d<f;d++){var e=this[d];if(e.nodeType===1&&e.className)if(a){for(var j=(" "+e.className+" ").replace(Aa," "),i=0,o=b.length;i<o;i++)j=j.replace(" "+b[i]+" ",
" ");e.className=c.trim(j)}else e.className=""}return this},toggleClass:function(a,b){var d=typeof a,f=typeof b==="boolean";if(c.isFunction(a))return this.each(function(e){var j=c(this);j.toggleClass(a.call(this,e,j.attr("class"),b),b)});return this.each(function(){if(d==="string")for(var e,j=0,i=c(this),o=b,k=a.split(ca);e=k[j++];){o=f?o:!i.hasClass(e);i[o?"addClass":"removeClass"](e)}else if(d==="undefined"||d==="boolean"){this.className&&c.data(this,"__className__",this.className);this.className=
this.className||a===false?"":c.data(this,"__className__")||""}})},hasClass:function(a){a=" "+a+" ";for(var b=0,d=this.length;b<d;b++)if((" "+this[b].className+" ").replace(Aa," ").indexOf(a)>-1)return true;return false},val:function(a){if(a===w){var b=this[0];if(b){if(c.nodeName(b,"option"))return(b.attributes.value||{}).specified?b.value:b.text;if(c.nodeName(b,"select")){var d=b.selectedIndex,f=[],e=b.options;b=b.type==="select-one";if(d<0)return null;var j=b?d:0;for(d=b?d+1:e.length;j<d;j++){var i=
e[j];if(i.selected){a=c(i).val();if(b)return a;f.push(a)}}return f}if(Ba.test(b.type)&&!c.support.checkOn)return b.getAttribute("value")===null?"on":b.value;return(b.value||"").replace(Za,"")}return w}var o=c.isFunction(a);return this.each(function(k){var n=c(this),r=a;if(this.nodeType===1){if(o)r=a.call(this,k,n.val());if(typeof r==="number")r+="";if(c.isArray(r)&&Ba.test(this.type))this.checked=c.inArray(n.val(),r)>=0;else if(c.nodeName(this,"select")){var u=c.makeArray(r);c("option",this).each(function(){this.selected=
c.inArray(c(this).val(),u)>=0});if(!u.length)this.selectedIndex=-1}else this.value=r}})}});c.extend({attrFn:{val:true,css:true,html:true,text:true,data:true,width:true,height:true,offset:true},attr:function(a,b,d,f){if(!a||a.nodeType===3||a.nodeType===8)return w;if(f&&b in c.attrFn)return c(a)[b](d);f=a.nodeType!==1||!c.isXMLDoc(a);var e=d!==w;b=f&&c.props[b]||b;if(a.nodeType===1){var j=$a.test(b);if(b in a&&f&&!j){if(e){b==="type"&&ab.test(a.nodeName)&&a.parentNode&&c.error("type property can't be changed");
a[b]=d}if(c.nodeName(a,"form")&&a.getAttributeNode(b))return a.getAttributeNode(b).nodeValue;if(b==="tabIndex")return(b=a.getAttributeNode("tabIndex"))&&b.specified?b.value:bb.test(a.nodeName)||cb.test(a.nodeName)&&a.href?0:w;return a[b]}if(!c.support.style&&f&&b==="style"){if(e)a.style.cssText=""+d;return a.style.cssText}e&&a.setAttribute(b,""+d);a=!c.support.hrefNormalized&&f&&j?a.getAttribute(b,2):a.getAttribute(b);return a===null?w:a}return c.style(a,b,d)}});var O=/\.(.*)$/,db=function(a){return a.replace(/[^\w\s\.\|`]/g,
function(b){return"\\"+b})};c.event={add:function(a,b,d,f){if(!(a.nodeType===3||a.nodeType===8)){if(a.setInterval&&a!==A&&!a.frameElement)a=A;var e,j;if(d.handler){e=d;d=e.handler}if(!d.guid)d.guid=c.guid++;if(j=c.data(a)){var i=j.events=j.events||{},o=j.handle;if(!o)j.handle=o=function(){return typeof c!=="undefined"&&!c.event.triggered?c.event.handle.apply(o.elem,arguments):w};o.elem=a;b=b.split(" ");for(var k,n=0,r;k=b[n++];){j=e?c.extend({},e):{handler:d,data:f};if(k.indexOf(".")>-1){r=k.split(".");
k=r.shift();j.namespace=r.slice(0).sort().join(".")}else{r=[];j.namespace=""}j.type=k;j.guid=d.guid;var u=i[k],z=c.event.special[k]||{};if(!u){u=i[k]=[];if(!z.setup||z.setup.call(a,f,r,o)===false)if(a.addEventListener)a.addEventListener(k,o,false);else a.attachEvent&&a.attachEvent("on"+k,o)}if(z.add){z.add.call(a,j);if(!j.handler.guid)j.handler.guid=d.guid}u.push(j);c.event.global[k]=true}a=null}}},global:{},remove:function(a,b,d,f){if(!(a.nodeType===3||a.nodeType===8)){var e,j=0,i,o,k,n,r,u,z=c.data(a),
C=z&&z.events;if(z&&C){if(b&&b.type){d=b.handler;b=b.type}if(!b||typeof b==="string"&&b.charAt(0)==="."){b=b||"";for(e in C)c.event.remove(a,e+b)}else{for(b=b.split(" ");e=b[j++];){n=e;i=e.indexOf(".")<0;o=[];if(!i){o=e.split(".");e=o.shift();k=new RegExp("(^|\\.)"+c.map(o.slice(0).sort(),db).join("\\.(?:.*\\.)?")+"(\\.|$)")}if(r=C[e])if(d){n=c.event.special[e]||{};for(B=f||0;B<r.length;B++){u=r[B];if(d.guid===u.guid){if(i||k.test(u.namespace)){f==null&&r.splice(B--,1);n.remove&&n.remove.call(a,u)}if(f!=
null)break}}if(r.length===0||f!=null&&r.length===1){if(!n.teardown||n.teardown.call(a,o)===false)Ca(a,e,z.handle);delete C[e]}}else for(var B=0;B<r.length;B++){u=r[B];if(i||k.test(u.namespace)){c.event.remove(a,n,u.handler,B);r.splice(B--,1)}}}if(c.isEmptyObject(C)){if(b=z.handle)b.elem=null;delete z.events;delete z.handle;c.isEmptyObject(z)&&c.removeData(a)}}}}},trigger:function(a,b,d,f){var e=a.type||a;if(!f){a=typeof a==="object"?a[G]?a:c.extend(c.Event(e),a):c.Event(e);if(e.indexOf("!")>=0){a.type=
e=e.slice(0,-1);a.exclusive=true}if(!d){a.stopPropagation();c.event.global[e]&&c.each(c.cache,function(){this.events&&this.events[e]&&c.event.trigger(a,b,this.handle.elem)})}if(!d||d.nodeType===3||d.nodeType===8)return w;a.result=w;a.target=d;b=c.makeArray(b);b.unshift(a)}a.currentTarget=d;(f=c.data(d,"handle"))&&f.apply(d,b);f=d.parentNode||d.ownerDocument;try{if(!(d&&d.nodeName&&c.noData[d.nodeName.toLowerCase()]))if(d["on"+e]&&d["on"+e].apply(d,b)===false)a.result=false}catch(j){}if(!a.isPropagationStopped()&&
f)c.event.trigger(a,b,f,true);else if(!a.isDefaultPrevented()){f=a.target;var i,o=c.nodeName(f,"a")&&e==="click",k=c.event.special[e]||{};if((!k._default||k._default.call(d,a)===false)&&!o&&!(f&&f.nodeName&&c.noData[f.nodeName.toLowerCase()])){try{if(f[e]){if(i=f["on"+e])f["on"+e]=null;c.event.triggered=true;f[e]()}}catch(n){}if(i)f["on"+e]=i;c.event.triggered=false}}},handle:function(a){var b,d,f,e;a=arguments[0]=c.event.fix(a||A.event);a.currentTarget=this;b=a.type.indexOf(".")<0&&!a.exclusive;
if(!b){d=a.type.split(".");a.type=d.shift();f=new RegExp("(^|\\.)"+d.slice(0).sort().join("\\.(?:.*\\.)?")+"(\\.|$)")}e=c.data(this,"events");d=e[a.type];if(e&&d){d=d.slice(0);e=0;for(var j=d.length;e<j;e++){var i=d[e];if(b||f.test(i.namespace)){a.handler=i.handler;a.data=i.data;a.handleObj=i;i=i.handler.apply(this,arguments);if(i!==w){a.result=i;if(i===false){a.preventDefault();a.stopPropagation()}}if(a.isImmediatePropagationStopped())break}}}return a.result},props:"altKey attrChange attrName bubbles button cancelable charCode clientX clientY ctrlKey currentTarget data detail eventPhase fromElement handler keyCode layerX layerY metaKey newValue offsetX offsetY originalTarget pageX pageY prevValue relatedNode relatedTarget screenX screenY shiftKey srcElement target toElement view wheelDelta which".split(" "),
fix:function(a){if(a[G])return a;var b=a;a=c.Event(b);for(var d=this.props.length,f;d;){f=this.props[--d];a[f]=b[f]}if(!a.target)a.target=a.srcElement||s;if(a.target.nodeType===3)a.target=a.target.parentNode;if(!a.relatedTarget&&a.fromElement)a.relatedTarget=a.fromElement===a.target?a.toElement:a.fromElement;if(a.pageX==null&&a.clientX!=null){b=s.documentElement;d=s.body;a.pageX=a.clientX+(b&&b.scrollLeft||d&&d.scrollLeft||0)-(b&&b.clientLeft||d&&d.clientLeft||0);a.pageY=a.clientY+(b&&b.scrollTop||
d&&d.scrollTop||0)-(b&&b.clientTop||d&&d.clientTop||0)}if(!a.which&&(a.charCode||a.charCode===0?a.charCode:a.keyCode))a.which=a.charCode||a.keyCode;if(!a.metaKey&&a.ctrlKey)a.metaKey=a.ctrlKey;if(!a.which&&a.button!==w)a.which=a.button&1?1:a.button&2?3:a.button&4?2:0;return a},guid:1E8,proxy:c.proxy,special:{ready:{setup:c.bindReady,teardown:c.noop},live:{add:function(a){c.event.add(this,a.origType,c.extend({},a,{handler:oa}))},remove:function(a){var b=true,d=a.origType.replace(O,"");c.each(c.data(this,
"events").live||[],function(){if(d===this.origType.replace(O,""))return b=false});b&&c.event.remove(this,a.origType,oa)}},beforeunload:{setup:function(a,b,d){if(this.setInterval)this.onbeforeunload=d;return false},teardown:function(a,b){if(this.onbeforeunload===b)this.onbeforeunload=null}}}};var Ca=s.removeEventListener?function(a,b,d){a.removeEventListener(b,d,false)}:function(a,b,d){a.detachEvent("on"+b,d)};c.Event=function(a){if(!this.preventDefault)return new c.Event(a);if(a&&a.type){this.originalEvent=
a;this.type=a.type}else this.type=a;this.timeStamp=J();this[G]=true};c.Event.prototype={preventDefault:function(){this.isDefaultPrevented=Z;var a=this.originalEvent;if(a){a.preventDefault&&a.preventDefault();a.returnValue=false}},stopPropagation:function(){this.isPropagationStopped=Z;var a=this.originalEvent;if(a){a.stopPropagation&&a.stopPropagation();a.cancelBubble=true}},stopImmediatePropagation:function(){this.isImmediatePropagationStopped=Z;this.stopPropagation()},isDefaultPrevented:Y,isPropagationStopped:Y,
isImmediatePropagationStopped:Y};var Da=function(a){var b=a.relatedTarget;try{for(;b&&b!==this;)b=b.parentNode;if(b!==this){a.type=a.data;c.event.handle.apply(this,arguments)}}catch(d){}},Ea=function(a){a.type=a.data;c.event.handle.apply(this,arguments)};c.each({mouseenter:"mouseover",mouseleave:"mouseout"},function(a,b){c.event.special[a]={setup:function(d){c.event.add(this,b,d&&d.selector?Ea:Da,a)},teardown:function(d){c.event.remove(this,b,d&&d.selector?Ea:Da)}}});if(!c.support.submitBubbles)c.event.special.submit=
{setup:function(){if(this.nodeName.toLowerCase()!=="form"){c.event.add(this,"click.specialSubmit",function(a){var b=a.target,d=b.type;if((d==="submit"||d==="image")&&c(b).closest("form").length)return na("submit",this,arguments)});c.event.add(this,"keypress.specialSubmit",function(a){var b=a.target,d=b.type;if((d==="text"||d==="password")&&c(b).closest("form").length&&a.keyCode===13)return na("submit",this,arguments)})}else return false},teardown:function(){c.event.remove(this,".specialSubmit")}};
if(!c.support.changeBubbles){var da=/textarea|input|select/i,ea,Fa=function(a){var b=a.type,d=a.value;if(b==="radio"||b==="checkbox")d=a.checked;else if(b==="select-multiple")d=a.selectedIndex>-1?c.map(a.options,function(f){return f.selected}).join("-"):"";else if(a.nodeName.toLowerCase()==="select")d=a.selectedIndex;return d},fa=function(a,b){var d=a.target,f,e;if(!(!da.test(d.nodeName)||d.readOnly)){f=c.data(d,"_change_data");e=Fa(d);if(a.type!=="focusout"||d.type!=="radio")c.data(d,"_change_data",
e);if(!(f===w||e===f))if(f!=null||e){a.type="change";return c.event.trigger(a,b,d)}}};c.event.special.change={filters:{focusout:fa,click:function(a){var b=a.target,d=b.type;if(d==="radio"||d==="checkbox"||b.nodeName.toLowerCase()==="select")return fa.call(this,a)},keydown:function(a){var b=a.target,d=b.type;if(a.keyCode===13&&b.nodeName.toLowerCase()!=="textarea"||a.keyCode===32&&(d==="checkbox"||d==="radio")||d==="select-multiple")return fa.call(this,a)},beforeactivate:function(a){a=a.target;c.data(a,
"_change_data",Fa(a))}},setup:function(){if(this.type==="file")return false;for(var a in ea)c.event.add(this,a+".specialChange",ea[a]);return da.test(this.nodeName)},teardown:function(){c.event.remove(this,".specialChange");return da.test(this.nodeName)}};ea=c.event.special.change.filters}s.addEventListener&&c.each({focus:"focusin",blur:"focusout"},function(a,b){function d(f){f=c.event.fix(f);f.type=b;return c.event.handle.call(this,f)}c.event.special[b]={setup:function(){this.addEventListener(a,
d,true)},teardown:function(){this.removeEventListener(a,d,true)}}});c.each(["bind","one"],function(a,b){c.fn[b]=function(d,f,e){if(typeof d==="object"){for(var j in d)this[b](j,f,d[j],e);return this}if(c.isFunction(f)){e=f;f=w}var i=b==="one"?c.proxy(e,function(k){c(this).unbind(k,i);return e.apply(this,arguments)}):e;if(d==="unload"&&b!=="one")this.one(d,f,e);else{j=0;for(var o=this.length;j<o;j++)c.event.add(this[j],d,i,f)}return this}});c.fn.extend({unbind:function(a,b){if(typeof a==="object"&&
!a.preventDefault)for(var d in a)this.unbind(d,a[d]);else{d=0;for(var f=this.length;d<f;d++)c.event.remove(this[d],a,b)}return this},delegate:function(a,b,d,f){return this.live(b,d,f,a)},undelegate:function(a,b,d){return arguments.length===0?this.unbind("live"):this.die(b,null,d,a)},trigger:function(a,b){return this.each(function(){c.event.trigger(a,b,this)})},triggerHandler:function(a,b){if(this[0]){a=c.Event(a);a.preventDefault();a.stopPropagation();c.event.trigger(a,b,this[0]);return a.result}},
toggle:function(a){for(var b=arguments,d=1;d<b.length;)c.proxy(a,b[d++]);return this.click(c.proxy(a,function(f){var e=(c.data(this,"lastToggle"+a.guid)||0)%d;c.data(this,"lastToggle"+a.guid,e+1);f.preventDefault();return b[e].apply(this,arguments)||false}))},hover:function(a,b){return this.mouseenter(a).mouseleave(b||a)}});var Ga={focus:"focusin",blur:"focusout",mouseenter:"mouseover",mouseleave:"mouseout"};c.each(["live","die"],function(a,b){c.fn[b]=function(d,f,e,j){var i,o=0,k,n,r=j||this.selector,
u=j?this:c(this.context);if(c.isFunction(f)){e=f;f=w}for(d=(d||"").split(" ");(i=d[o++])!=null;){j=O.exec(i);k="";if(j){k=j[0];i=i.replace(O,"")}if(i==="hover")d.push("mouseenter"+k,"mouseleave"+k);else{n=i;if(i==="focus"||i==="blur"){d.push(Ga[i]+k);i+=k}else i=(Ga[i]||i)+k;b==="live"?u.each(function(){c.event.add(this,pa(i,r),{data:f,selector:r,handler:e,origType:i,origHandler:e,preType:n})}):u.unbind(pa(i,r),e)}}return this}});c.each("blur focus focusin focusout load resize scroll unload click dblclick mousedown mouseup mousemove mouseover mouseout mouseenter mouseleave change select submit keydown keypress keyup error".split(" "),
function(a,b){c.fn[b]=function(d){return d?this.bind(b,d):this.trigger(b)};if(c.attrFn)c.attrFn[b]=true});A.attachEvent&&!A.addEventListener&&A.attachEvent("onunload",function(){for(var a in c.cache)if(c.cache[a].handle)try{c.event.remove(c.cache[a].handle.elem)}catch(b){}});(function(){function a(g){for(var h="",l,m=0;g[m];m++){l=g[m];if(l.nodeType===3||l.nodeType===4)h+=l.nodeValue;else if(l.nodeType!==8)h+=a(l.childNodes)}return h}function b(g,h,l,m,q,p){q=0;for(var v=m.length;q<v;q++){var t=m[q];
if(t){t=t[g];for(var y=false;t;){if(t.sizcache===l){y=m[t.sizset];break}if(t.nodeType===1&&!p){t.sizcache=l;t.sizset=q}if(t.nodeName.toLowerCase()===h){y=t;break}t=t[g]}m[q]=y}}}function d(g,h,l,m,q,p){q=0;for(var v=m.length;q<v;q++){var t=m[q];if(t){t=t[g];for(var y=false;t;){if(t.sizcache===l){y=m[t.sizset];break}if(t.nodeType===1){if(!p){t.sizcache=l;t.sizset=q}if(typeof h!=="string"){if(t===h){y=true;break}}else if(k.filter(h,[t]).length>0){y=t;break}}t=t[g]}m[q]=y}}}var f=/((?:\((?:\([^()]+\)|[^()]+)+\)|\[(?:\[[^[\]]*\]|['"][^'"]*['"]|[^[\]'"]+)+\]|\\.|[^ >+~,(\[\\]+)+|[>+~])(\s*,\s*)?((?:.|\r|\n)*)/g,
e=0,j=Object.prototype.toString,i=false,o=true;[0,0].sort(function(){o=false;return 0});var k=function(g,h,l,m){l=l||[];var q=h=h||s;if(h.nodeType!==1&&h.nodeType!==9)return[];if(!g||typeof g!=="string")return l;for(var p=[],v,t,y,S,H=true,M=x(h),I=g;(f.exec(""),v=f.exec(I))!==null;){I=v[3];p.push(v[1]);if(v[2]){S=v[3];break}}if(p.length>1&&r.exec(g))if(p.length===2&&n.relative[p[0]])t=ga(p[0]+p[1],h);else for(t=n.relative[p[0]]?[h]:k(p.shift(),h);p.length;){g=p.shift();if(n.relative[g])g+=p.shift();
t=ga(g,t)}else{if(!m&&p.length>1&&h.nodeType===9&&!M&&n.match.ID.test(p[0])&&!n.match.ID.test(p[p.length-1])){v=k.find(p.shift(),h,M);h=v.expr?k.filter(v.expr,v.set)[0]:v.set[0]}if(h){v=m?{expr:p.pop(),set:z(m)}:k.find(p.pop(),p.length===1&&(p[0]==="~"||p[0]==="+")&&h.parentNode?h.parentNode:h,M);t=v.expr?k.filter(v.expr,v.set):v.set;if(p.length>0)y=z(t);else H=false;for(;p.length;){var D=p.pop();v=D;if(n.relative[D])v=p.pop();else D="";if(v==null)v=h;n.relative[D](y,v,M)}}else y=[]}y||(y=t);y||k.error(D||
g);if(j.call(y)==="[object Array]")if(H)if(h&&h.nodeType===1)for(g=0;y[g]!=null;g++){if(y[g]&&(y[g]===true||y[g].nodeType===1&&E(h,y[g])))l.push(t[g])}else for(g=0;y[g]!=null;g++)y[g]&&y[g].nodeType===1&&l.push(t[g]);else l.push.apply(l,y);else z(y,l);if(S){k(S,q,l,m);k.uniqueSort(l)}return l};k.uniqueSort=function(g){if(B){i=o;g.sort(B);if(i)for(var h=1;h<g.length;h++)g[h]===g[h-1]&&g.splice(h--,1)}return g};k.matches=function(g,h){return k(g,null,null,h)};k.find=function(g,h,l){var m,q;if(!g)return[];
for(var p=0,v=n.order.length;p<v;p++){var t=n.order[p];if(q=n.leftMatch[t].exec(g)){var y=q[1];q.splice(1,1);if(y.substr(y.length-1)!=="\\"){q[1]=(q[1]||"").replace(/\\/g,"");m=n.find[t](q,h,l);if(m!=null){g=g.replace(n.match[t],"");break}}}}m||(m=h.getElementsByTagName("*"));return{set:m,expr:g}};k.filter=function(g,h,l,m){for(var q=g,p=[],v=h,t,y,S=h&&h[0]&&x(h[0]);g&&h.length;){for(var H in n.filter)if((t=n.leftMatch[H].exec(g))!=null&&t[2]){var M=n.filter[H],I,D;D=t[1];y=false;t.splice(1,1);if(D.substr(D.length-
1)!=="\\"){if(v===p)p=[];if(n.preFilter[H])if(t=n.preFilter[H](t,v,l,p,m,S)){if(t===true)continue}else y=I=true;if(t)for(var U=0;(D=v[U])!=null;U++)if(D){I=M(D,t,U,v);var Ha=m^!!I;if(l&&I!=null)if(Ha)y=true;else v[U]=false;else if(Ha){p.push(D);y=true}}if(I!==w){l||(v=p);g=g.replace(n.match[H],"");if(!y)return[];break}}}if(g===q)if(y==null)k.error(g);else break;q=g}return v};k.error=function(g){throw"Syntax error, unrecognized expression: "+g;};var n=k.selectors={order:["ID","NAME","TAG"],match:{ID:/#((?:[\w\u00c0-\uFFFF-]|\\.)+)/,
CLASS:/\.((?:[\w\u00c0-\uFFFF-]|\\.)+)/,NAME:/\[name=['"]*((?:[\w\u00c0-\uFFFF-]|\\.)+)['"]*\]/,ATTR:/\[\s*((?:[\w\u00c0-\uFFFF-]|\\.)+)\s*(?:(\S?=)\s*(['"]*)(.*?)\3|)\s*\]/,TAG:/^((?:[\w\u00c0-\uFFFF\*-]|\\.)+)/,CHILD:/:(only|nth|last|first)-child(?:\((even|odd|[\dn+-]*)\))?/,POS:/:(nth|eq|gt|lt|first|last|even|odd)(?:\((\d*)\))?(?=[^-]|$)/,PSEUDO:/:((?:[\w\u00c0-\uFFFF-]|\\.)+)(?:\((['"]?)((?:\([^\)]+\)|[^\(\)]*)+)\2\))?/},leftMatch:{},attrMap:{"class":"className","for":"htmlFor"},attrHandle:{href:function(g){return g.getAttribute("href")}},
relative:{"+":function(g,h){var l=typeof h==="string",m=l&&!/\W/.test(h);l=l&&!m;if(m)h=h.toLowerCase();m=0;for(var q=g.length,p;m<q;m++)if(p=g[m]){for(;(p=p.previousSibling)&&p.nodeType!==1;);g[m]=l||p&&p.nodeName.toLowerCase()===h?p||false:p===h}l&&k.filter(h,g,true)},">":function(g,h){var l=typeof h==="string";if(l&&!/\W/.test(h)){h=h.toLowerCase();for(var m=0,q=g.length;m<q;m++){var p=g[m];if(p){l=p.parentNode;g[m]=l.nodeName.toLowerCase()===h?l:false}}}else{m=0;for(q=g.length;m<q;m++)if(p=g[m])g[m]=
l?p.parentNode:p.parentNode===h;l&&k.filter(h,g,true)}},"":function(g,h,l){var m=e++,q=d;if(typeof h==="string"&&!/\W/.test(h)){var p=h=h.toLowerCase();q=b}q("parentNode",h,m,g,p,l)},"~":function(g,h,l){var m=e++,q=d;if(typeof h==="string"&&!/\W/.test(h)){var p=h=h.toLowerCase();q=b}q("previousSibling",h,m,g,p,l)}},find:{ID:function(g,h,l){if(typeof h.getElementById!=="undefined"&&!l)return(g=h.getElementById(g[1]))?[g]:[]},NAME:function(g,h){if(typeof h.getElementsByName!=="undefined"){var l=[];
h=h.getElementsByName(g[1]);for(var m=0,q=h.length;m<q;m++)h[m].getAttribute("name")===g[1]&&l.push(h[m]);return l.length===0?null:l}},TAG:function(g,h){return h.getElementsByTagName(g[1])}},preFilter:{CLASS:function(g,h,l,m,q,p){g=" "+g[1].replace(/\\/g,"")+" ";if(p)return g;p=0;for(var v;(v=h[p])!=null;p++)if(v)if(q^(v.className&&(" "+v.className+" ").replace(/[\t\n]/g," ").indexOf(g)>=0))l||m.push(v);else if(l)h[p]=false;return false},ID:function(g){return g[1].replace(/\\/g,"")},TAG:function(g){return g[1].toLowerCase()},
CHILD:function(g){if(g[1]==="nth"){var h=/(-?)(\d*)n((?:\+|-)?\d*)/.exec(g[2]==="even"&&"2n"||g[2]==="odd"&&"2n+1"||!/\D/.test(g[2])&&"0n+"+g[2]||g[2]);g[2]=h[1]+(h[2]||1)-0;g[3]=h[3]-0}g[0]=e++;return g},ATTR:function(g,h,l,m,q,p){h=g[1].replace(/\\/g,"");if(!p&&n.attrMap[h])g[1]=n.attrMap[h];if(g[2]==="~=")g[4]=" "+g[4]+" ";return g},PSEUDO:function(g,h,l,m,q){if(g[1]==="not")if((f.exec(g[3])||"").length>1||/^\w/.test(g[3]))g[3]=k(g[3],null,null,h);else{g=k.filter(g[3],h,l,true^q);l||m.push.apply(m,
g);return false}else if(n.match.POS.test(g[0])||n.match.CHILD.test(g[0]))return true;return g},POS:function(g){g.unshift(true);return g}},filters:{enabled:function(g){return g.disabled===false&&g.type!=="hidden"},disabled:function(g){return g.disabled===true},checked:function(g){return g.checked===true},selected:function(g){return g.selected===true},parent:function(g){return!!g.firstChild},empty:function(g){return!g.firstChild},has:function(g,h,l){return!!k(l[3],g).length},header:function(g){return/h\d/i.test(g.nodeName)},
text:function(g){return"text"===g.type},radio:function(g){return"radio"===g.type},checkbox:function(g){return"checkbox"===g.type},file:function(g){return"file"===g.type},password:function(g){return"password"===g.type},submit:function(g){return"submit"===g.type},image:function(g){return"image"===g.type},reset:function(g){return"reset"===g.type},button:function(g){return"button"===g.type||g.nodeName.toLowerCase()==="button"},input:function(g){return/input|select|textarea|button/i.test(g.nodeName)}},
setFilters:{first:function(g,h){return h===0},last:function(g,h,l,m){return h===m.length-1},even:function(g,h){return h%2===0},odd:function(g,h){return h%2===1},lt:function(g,h,l){return h<l[3]-0},gt:function(g,h,l){return h>l[3]-0},nth:function(g,h,l){return l[3]-0===h},eq:function(g,h,l){return l[3]-0===h}},filter:{PSEUDO:function(g,h,l,m){var q=h[1],p=n.filters[q];if(p)return p(g,l,h,m);else if(q==="contains")return(g.textContent||g.innerText||a([g])||"").indexOf(h[3])>=0;else if(q==="not"){h=
h[3];l=0;for(m=h.length;l<m;l++)if(h[l]===g)return false;return true}else k.error("Syntax error, unrecognized expression: "+q)},CHILD:function(g,h){var l=h[1],m=g;switch(l){case "only":case "first":for(;m=m.previousSibling;)if(m.nodeType===1)return false;if(l==="first")return true;m=g;case "last":for(;m=m.nextSibling;)if(m.nodeType===1)return false;return true;case "nth":l=h[2];var q=h[3];if(l===1&&q===0)return true;h=h[0];var p=g.parentNode;if(p&&(p.sizcache!==h||!g.nodeIndex)){var v=0;for(m=p.firstChild;m;m=
m.nextSibling)if(m.nodeType===1)m.nodeIndex=++v;p.sizcache=h}g=g.nodeIndex-q;return l===0?g===0:g%l===0&&g/l>=0}},ID:function(g,h){return g.nodeType===1&&g.getAttribute("id")===h},TAG:function(g,h){return h==="*"&&g.nodeType===1||g.nodeName.toLowerCase()===h},CLASS:function(g,h){return(" "+(g.className||g.getAttribute("class"))+" ").indexOf(h)>-1},ATTR:function(g,h){var l=h[1];g=n.attrHandle[l]?n.attrHandle[l](g):g[l]!=null?g[l]:g.getAttribute(l);l=g+"";var m=h[2];h=h[4];return g==null?m==="!=":m===
"="?l===h:m==="*="?l.indexOf(h)>=0:m==="~="?(" "+l+" ").indexOf(h)>=0:!h?l&&g!==false:m==="!="?l!==h:m==="^="?l.indexOf(h)===0:m==="$="?l.substr(l.length-h.length)===h:m==="|="?l===h||l.substr(0,h.length+1)===h+"-":false},POS:function(g,h,l,m){var q=n.setFilters[h[2]];if(q)return q(g,l,h,m)}}},r=n.match.POS;for(var u in n.match){n.match[u]=new RegExp(n.match[u].source+/(?![^\[]*\])(?![^\(]*\))/.source);n.leftMatch[u]=new RegExp(/(^(?:.|\r|\n)*?)/.source+n.match[u].source.replace(/\\(\d+)/g,function(g,
h){return"\\"+(h-0+1)}))}var z=function(g,h){g=Array.prototype.slice.call(g,0);if(h){h.push.apply(h,g);return h}return g};try{Array.prototype.slice.call(s.documentElement.childNodes,0)}catch(C){z=function(g,h){h=h||[];if(j.call(g)==="[object Array]")Array.prototype.push.apply(h,g);else if(typeof g.length==="number")for(var l=0,m=g.length;l<m;l++)h.push(g[l]);else for(l=0;g[l];l++)h.push(g[l]);return h}}var B;if(s.documentElement.compareDocumentPosition)B=function(g,h){if(!g.compareDocumentPosition||
!h.compareDocumentPosition){if(g==h)i=true;return g.compareDocumentPosition?-1:1}g=g.compareDocumentPosition(h)&4?-1:g===h?0:1;if(g===0)i=true;return g};else if("sourceIndex"in s.documentElement)B=function(g,h){if(!g.sourceIndex||!h.sourceIndex){if(g==h)i=true;return g.sourceIndex?-1:1}g=g.sourceIndex-h.sourceIndex;if(g===0)i=true;return g};else if(s.createRange)B=function(g,h){if(!g.ownerDocument||!h.ownerDocument){if(g==h)i=true;return g.ownerDocument?-1:1}var l=g.ownerDocument.createRange(),m=
h.ownerDocument.createRange();l.setStart(g,0);l.setEnd(g,0);m.setStart(h,0);m.setEnd(h,0);g=l.compareBoundaryPoints(Range.START_TO_END,m);if(g===0)i=true;return g};(function(){var g=s.createElement("div"),h="script"+(new Date).getTime();g.innerHTML="<a name='"+h+"'/>";var l=s.documentElement;l.insertBefore(g,l.firstChild);if(s.getElementById(h)){n.find.ID=function(m,q,p){if(typeof q.getElementById!=="undefined"&&!p)return(q=q.getElementById(m[1]))?q.id===m[1]||typeof q.getAttributeNode!=="undefined"&&
q.getAttributeNode("id").nodeValue===m[1]?[q]:w:[]};n.filter.ID=function(m,q){var p=typeof m.getAttributeNode!=="undefined"&&m.getAttributeNode("id");return m.nodeType===1&&p&&p.nodeValue===q}}l.removeChild(g);l=g=null})();(function(){var g=s.createElement("div");g.appendChild(s.createComment(""));if(g.getElementsByTagName("*").length>0)n.find.TAG=function(h,l){l=l.getElementsByTagName(h[1]);if(h[1]==="*"){h=[];for(var m=0;l[m];m++)l[m].nodeType===1&&h.push(l[m]);l=h}return l};g.innerHTML="<a href='#'></a>";
if(g.firstChild&&typeof g.firstChild.getAttribute!=="undefined"&&g.firstChild.getAttribute("href")!=="#")n.attrHandle.href=function(h){return h.getAttribute("href",2)};g=null})();s.querySelectorAll&&function(){var g=k,h=s.createElement("div");h.innerHTML="<p class='TEST'></p>";if(!(h.querySelectorAll&&h.querySelectorAll(".TEST").length===0)){k=function(m,q,p,v){q=q||s;if(!v&&q.nodeType===9&&!x(q))try{return z(q.querySelectorAll(m),p)}catch(t){}return g(m,q,p,v)};for(var l in g)k[l]=g[l];h=null}}();
(function(){var g=s.createElement("div");g.innerHTML="<div class='test e'></div><div class='test'></div>";if(!(!g.getElementsByClassName||g.getElementsByClassName("e").length===0)){g.lastChild.className="e";if(g.getElementsByClassName("e").length!==1){n.order.splice(1,0,"CLASS");n.find.CLASS=function(h,l,m){if(typeof l.getElementsByClassName!=="undefined"&&!m)return l.getElementsByClassName(h[1])};g=null}}})();var E=s.compareDocumentPosition?function(g,h){return!!(g.compareDocumentPosition(h)&16)}:
function(g,h){return g!==h&&(g.contains?g.contains(h):true)},x=function(g){return(g=(g?g.ownerDocument||g:0).documentElement)?g.nodeName!=="HTML":false},ga=function(g,h){var l=[],m="",q;for(h=h.nodeType?[h]:h;q=n.match.PSEUDO.exec(g);){m+=q[0];g=g.replace(n.match.PSEUDO,"")}g=n.relative[g]?g+"*":g;q=0;for(var p=h.length;q<p;q++)k(g,h[q],l);return k.filter(m,l)};c.find=k;c.expr=k.selectors;c.expr[":"]=c.expr.filters;c.unique=k.uniqueSort;c.text=a;c.isXMLDoc=x;c.contains=E})();var eb=/Until$/,fb=/^(?:parents|prevUntil|prevAll)/,
gb=/,/;R=Array.prototype.slice;var Ia=function(a,b,d){if(c.isFunction(b))return c.grep(a,function(e,j){return!!b.call(e,j,e)===d});else if(b.nodeType)return c.grep(a,function(e){return e===b===d});else if(typeof b==="string"){var f=c.grep(a,function(e){return e.nodeType===1});if(Ua.test(b))return c.filter(b,f,!d);else b=c.filter(b,f)}return c.grep(a,function(e){return c.inArray(e,b)>=0===d})};c.fn.extend({find:function(a){for(var b=this.pushStack("","find",a),d=0,f=0,e=this.length;f<e;f++){d=b.length;
c.find(a,this[f],b);if(f>0)for(var j=d;j<b.length;j++)for(var i=0;i<d;i++)if(b[i]===b[j]){b.splice(j--,1);break}}return b},has:function(a){var b=c(a);return this.filter(function(){for(var d=0,f=b.length;d<f;d++)if(c.contains(this,b[d]))return true})},not:function(a){return this.pushStack(Ia(this,a,false),"not",a)},filter:function(a){return this.pushStack(Ia(this,a,true),"filter",a)},is:function(a){return!!a&&c.filter(a,this).length>0},closest:function(a,b){if(c.isArray(a)){var d=[],f=this[0],e,j=
{},i;if(f&&a.length){e=0;for(var o=a.length;e<o;e++){i=a[e];j[i]||(j[i]=c.expr.match.POS.test(i)?c(i,b||this.context):i)}for(;f&&f.ownerDocument&&f!==b;){for(i in j){e=j[i];if(e.jquery?e.index(f)>-1:c(f).is(e)){d.push({selector:i,elem:f});delete j[i]}}f=f.parentNode}}return d}var k=c.expr.match.POS.test(a)?c(a,b||this.context):null;return this.map(function(n,r){for(;r&&r.ownerDocument&&r!==b;){if(k?k.index(r)>-1:c(r).is(a))return r;r=r.parentNode}return null})},index:function(a){if(!a||typeof a===
"string")return c.inArray(this[0],a?c(a):this.parent().children());return c.inArray(a.jquery?a[0]:a,this)},add:function(a,b){a=typeof a==="string"?c(a,b||this.context):c.makeArray(a);b=c.merge(this.get(),a);return this.pushStack(qa(a[0])||qa(b[0])?b:c.unique(b))},andSelf:function(){return this.add(this.prevObject)}});c.each({parent:function(a){return(a=a.parentNode)&&a.nodeType!==11?a:null},parents:function(a){return c.dir(a,"parentNode")},parentsUntil:function(a,b,d){return c.dir(a,"parentNode",
d)},next:function(a){return c.nth(a,2,"nextSibling")},prev:function(a){return c.nth(a,2,"previousSibling")},nextAll:function(a){return c.dir(a,"nextSibling")},prevAll:function(a){return c.dir(a,"previousSibling")},nextUntil:function(a,b,d){return c.dir(a,"nextSibling",d)},prevUntil:function(a,b,d){return c.dir(a,"previousSibling",d)},siblings:function(a){return c.sibling(a.parentNode.firstChild,a)},children:function(a){return c.sibling(a.firstChild)},contents:function(a){return c.nodeName(a,"iframe")?
a.contentDocument||a.contentWindow.document:c.makeArray(a.childNodes)}},function(a,b){c.fn[a]=function(d,f){var e=c.map(this,b,d);eb.test(a)||(f=d);if(f&&typeof f==="string")e=c.filter(f,e);e=this.length>1?c.unique(e):e;if((this.length>1||gb.test(f))&&fb.test(a))e=e.reverse();return this.pushStack(e,a,R.call(arguments).join(","))}});c.extend({filter:function(a,b,d){if(d)a=":not("+a+")";return c.find.matches(a,b)},dir:function(a,b,d){var f=[];for(a=a[b];a&&a.nodeType!==9&&(d===w||a.nodeType!==1||!c(a).is(d));){a.nodeType===
1&&f.push(a);a=a[b]}return f},nth:function(a,b,d){b=b||1;for(var f=0;a;a=a[d])if(a.nodeType===1&&++f===b)break;return a},sibling:function(a,b){for(var d=[];a;a=a.nextSibling)a.nodeType===1&&a!==b&&d.push(a);return d}});var Ja=/ jQuery\d+="(?:\d+|null)"/g,V=/^\s+/,Ka=/(<([\w:]+)[^>]*?)\/>/g,hb=/^(?:area|br|col|embed|hr|img|input|link|meta|param)$/i,La=/<([\w:]+)/,ib=/<tbody/i,jb=/<|&#?\w+;/,ta=/<script|<object|<embed|<option|<style/i,ua=/checked\s*(?:[^=]|=\s*.checked.)/i,Ma=function(a,b,d){return hb.test(d)?
a:b+"></"+d+">"},F={option:[1,"<select multiple='multiple'>","</select>"],legend:[1,"<fieldset>","</fieldset>"],thead:[1,"<table>","</table>"],tr:[2,"<table><tbody>","</tbody></table>"],td:[3,"<table><tbody><tr>","</tr></tbody></table>"],col:[2,"<table><tbody></tbody><colgroup>","</colgroup></table>"],area:[1,"<map>","</map>"],_default:[0,"",""]};F.optgroup=F.option;F.tbody=F.tfoot=F.colgroup=F.caption=F.thead;F.th=F.td;if(!c.support.htmlSerialize)F._default=[1,"div<div>","</div>"];c.fn.extend({text:function(a){if(c.isFunction(a))return this.each(function(b){var d=
c(this);d.text(a.call(this,b,d.text()))});if(typeof a!=="object"&&a!==w)return this.empty().append((this[0]&&this[0].ownerDocument||s).createTextNode(a));return c.text(this)},wrapAll:function(a){if(c.isFunction(a))return this.each(function(d){c(this).wrapAll(a.call(this,d))});if(this[0]){var b=c(a,this[0].ownerDocument).eq(0).clone(true);this[0].parentNode&&b.insertBefore(this[0]);b.map(function(){for(var d=this;d.firstChild&&d.firstChild.nodeType===1;)d=d.firstChild;return d}).append(this)}return this},
wrapInner:function(a){if(c.isFunction(a))return this.each(function(b){c(this).wrapInner(a.call(this,b))});return this.each(function(){var b=c(this),d=b.contents();d.length?d.wrapAll(a):b.append(a)})},wrap:function(a){return this.each(function(){c(this).wrapAll(a)})},unwrap:function(){return this.parent().each(function(){c.nodeName(this,"body")||c(this).replaceWith(this.childNodes)}).end()},append:function(){return this.domManip(arguments,true,function(a){this.nodeType===1&&this.appendChild(a)})},
prepend:function(){return this.domManip(arguments,true,function(a){this.nodeType===1&&this.insertBefore(a,this.firstChild)})},before:function(){if(this[0]&&this[0].parentNode)return this.domManip(arguments,false,function(b){this.parentNode.insertBefore(b,this)});else if(arguments.length){var a=c(arguments[0]);a.push.apply(a,this.toArray());return this.pushStack(a,"before",arguments)}},after:function(){if(this[0]&&this[0].parentNode)return this.domManip(arguments,false,function(b){this.parentNode.insertBefore(b,
this.nextSibling)});else if(arguments.length){var a=this.pushStack(this,"after",arguments);a.push.apply(a,c(arguments[0]).toArray());return a}},remove:function(a,b){for(var d=0,f;(f=this[d])!=null;d++)if(!a||c.filter(a,[f]).length){if(!b&&f.nodeType===1){c.cleanData(f.getElementsByTagName("*"));c.cleanData([f])}f.parentNode&&f.parentNode.removeChild(f)}return this},empty:function(){for(var a=0,b;(b=this[a])!=null;a++)for(b.nodeType===1&&c.cleanData(b.getElementsByTagName("*"));b.firstChild;)b.removeChild(b.firstChild);
return this},clone:function(a){var b=this.map(function(){if(!c.support.noCloneEvent&&!c.isXMLDoc(this)){var d=this.outerHTML,f=this.ownerDocument;if(!d){d=f.createElement("div");d.appendChild(this.cloneNode(true));d=d.innerHTML}return c.clean([d.replace(Ja,"").replace(/=([^="'>\s]+\/)>/g,'="$1">').replace(V,"")],f)[0]}else return this.cloneNode(true)});if(a===true){ra(this,b);ra(this.find("*"),b.find("*"))}return b},html:function(a){if(a===w)return this[0]&&this[0].nodeType===1?this[0].innerHTML.replace(Ja,
""):null;else if(typeof a==="string"&&!ta.test(a)&&(c.support.leadingWhitespace||!V.test(a))&&!F[(La.exec(a)||["",""])[1].toLowerCase()]){a=a.replace(Ka,Ma);try{for(var b=0,d=this.length;b<d;b++)if(this[b].nodeType===1){c.cleanData(this[b].getElementsByTagName("*"));this[b].innerHTML=a}}catch(f){this.empty().append(a)}}else c.isFunction(a)?this.each(function(e){var j=c(this),i=j.html();j.empty().append(function(){return a.call(this,e,i)})}):this.empty().append(a);return this},replaceWith:function(a){if(this[0]&&
this[0].parentNode){if(c.isFunction(a))return this.each(function(b){var d=c(this),f=d.html();d.replaceWith(a.call(this,b,f))});if(typeof a!=="string")a=c(a).detach();return this.each(function(){var b=this.nextSibling,d=this.parentNode;c(this).remove();b?c(b).before(a):c(d).append(a)})}else return this.pushStack(c(c.isFunction(a)?a():a),"replaceWith",a)},detach:function(a){return this.remove(a,true)},domManip:function(a,b,d){function f(u){return c.nodeName(u,"table")?u.getElementsByTagName("tbody")[0]||
u.appendChild(u.ownerDocument.createElement("tbody")):u}var e,j,i=a[0],o=[],k;if(!c.support.checkClone&&arguments.length===3&&typeof i==="string"&&ua.test(i))return this.each(function(){c(this).domManip(a,b,d,true)});if(c.isFunction(i))return this.each(function(u){var z=c(this);a[0]=i.call(this,u,b?z.html():w);z.domManip(a,b,d)});if(this[0]){e=i&&i.parentNode;e=c.support.parentNode&&e&&e.nodeType===11&&e.childNodes.length===this.length?{fragment:e}:sa(a,this,o);k=e.fragment;if(j=k.childNodes.length===
1?(k=k.firstChild):k.firstChild){b=b&&c.nodeName(j,"tr");for(var n=0,r=this.length;n<r;n++)d.call(b?f(this[n],j):this[n],n>0||e.cacheable||this.length>1?k.cloneNode(true):k)}o.length&&c.each(o,Qa)}return this}});c.fragments={};c.each({appendTo:"append",prependTo:"prepend",insertBefore:"before",insertAfter:"after",replaceAll:"replaceWith"},function(a,b){c.fn[a]=function(d){var f=[];d=c(d);var e=this.length===1&&this[0].parentNode;if(e&&e.nodeType===11&&e.childNodes.length===1&&d.length===1){d[b](this[0]);
return this}else{e=0;for(var j=d.length;e<j;e++){var i=(e>0?this.clone(true):this).get();c.fn[b].apply(c(d[e]),i);f=f.concat(i)}return this.pushStack(f,a,d.selector)}}});c.extend({clean:function(a,b,d,f){b=b||s;if(typeof b.createElement==="undefined")b=b.ownerDocument||b[0]&&b[0].ownerDocument||s;for(var e=[],j=0,i;(i=a[j])!=null;j++){if(typeof i==="number")i+="";if(i){if(typeof i==="string"&&!jb.test(i))i=b.createTextNode(i);else if(typeof i==="string"){i=i.replace(Ka,Ma);var o=(La.exec(i)||["",
""])[1].toLowerCase(),k=F[o]||F._default,n=k[0],r=b.createElement("div");for(r.innerHTML=k[1]+i+k[2];n--;)r=r.lastChild;if(!c.support.tbody){n=ib.test(i);o=o==="table"&&!n?r.firstChild&&r.firstChild.childNodes:k[1]==="<table>"&&!n?r.childNodes:[];for(k=o.length-1;k>=0;--k)c.nodeName(o[k],"tbody")&&!o[k].childNodes.length&&o[k].parentNode.removeChild(o[k])}!c.support.leadingWhitespace&&V.test(i)&&r.insertBefore(b.createTextNode(V.exec(i)[0]),r.firstChild);i=r.childNodes}if(i.nodeType)e.push(i);else e=
c.merge(e,i)}}if(d)for(j=0;e[j];j++)if(f&&c.nodeName(e[j],"script")&&(!e[j].type||e[j].type.toLowerCase()==="text/javascript"))f.push(e[j].parentNode?e[j].parentNode.removeChild(e[j]):e[j]);else{e[j].nodeType===1&&e.splice.apply(e,[j+1,0].concat(c.makeArray(e[j].getElementsByTagName("script"))));d.appendChild(e[j])}return e},cleanData:function(a){for(var b,d,f=c.cache,e=c.event.special,j=c.support.deleteExpando,i=0,o;(o=a[i])!=null;i++)if(d=o[c.expando]){b=f[d];if(b.events)for(var k in b.events)e[k]?
c.event.remove(o,k):Ca(o,k,b.handle);if(j)delete o[c.expando];else o.removeAttribute&&o.removeAttribute(c.expando);delete f[d]}}});var kb=/z-?index|font-?weight|opacity|zoom|line-?height/i,Na=/alpha\([^)]*\)/,Oa=/opacity=([^)]*)/,ha=/float/i,ia=/-([a-z])/ig,lb=/([A-Z])/g,mb=/^-?\d+(?:px)?$/i,nb=/^-?\d/,ob={position:"absolute",visibility:"hidden",display:"block"},pb=["Left","Right"],qb=["Top","Bottom"],rb=s.defaultView&&s.defaultView.getComputedStyle,Pa=c.support.cssFloat?"cssFloat":"styleFloat",ja=
function(a,b){return b.toUpperCase()};c.fn.css=function(a,b){return X(this,a,b,true,function(d,f,e){if(e===w)return c.curCSS(d,f);if(typeof e==="number"&&!kb.test(f))e+="px";c.style(d,f,e)})};c.extend({style:function(a,b,d){if(!a||a.nodeType===3||a.nodeType===8)return w;if((b==="width"||b==="height")&&parseFloat(d)<0)d=w;var f=a.style||a,e=d!==w;if(!c.support.opacity&&b==="opacity"){if(e){f.zoom=1;b=parseInt(d,10)+""==="NaN"?"":"alpha(opacity="+d*100+")";a=f.filter||c.curCSS(a,"filter")||"";f.filter=
Na.test(a)?a.replace(Na,b):b}return f.filter&&f.filter.indexOf("opacity=")>=0?parseFloat(Oa.exec(f.filter)[1])/100+"":""}if(ha.test(b))b=Pa;b=b.replace(ia,ja);if(e)f[b]=d;return f[b]},css:function(a,b,d,f){if(b==="width"||b==="height"){var e,j=b==="width"?pb:qb;function i(){e=b==="width"?a.offsetWidth:a.offsetHeight;f!=="border"&&c.each(j,function(){f||(e-=parseFloat(c.curCSS(a,"padding"+this,true))||0);if(f==="margin")e+=parseFloat(c.curCSS(a,"margin"+this,true))||0;else e-=parseFloat(c.curCSS(a,
"border"+this+"Width",true))||0})}a.offsetWidth!==0?i():c.swap(a,ob,i);return Math.max(0,Math.round(e))}return c.curCSS(a,b,d)},curCSS:function(a,b,d){var f,e=a.style;if(!c.support.opacity&&b==="opacity"&&a.currentStyle){f=Oa.test(a.currentStyle.filter||"")?parseFloat(RegExp.$1)/100+"":"";return f===""?"1":f}if(ha.test(b))b=Pa;if(!d&&e&&e[b])f=e[b];else if(rb){if(ha.test(b))b="float";b=b.replace(lb,"-$1").toLowerCase();e=a.ownerDocument.defaultView;if(!e)return null;if(a=e.getComputedStyle(a,null))f=
a.getPropertyValue(b);if(b==="opacity"&&f==="")f="1"}else if(a.currentStyle){d=b.replace(ia,ja);f=a.currentStyle[b]||a.currentStyle[d];if(!mb.test(f)&&nb.test(f)){b=e.left;var j=a.runtimeStyle.left;a.runtimeStyle.left=a.currentStyle.left;e.left=d==="fontSize"?"1em":f||0;f=e.pixelLeft+"px";e.left=b;a.runtimeStyle.left=j}}return f},swap:function(a,b,d){var f={};for(var e in b){f[e]=a.style[e];a.style[e]=b[e]}d.call(a);for(e in b)a.style[e]=f[e]}});if(c.expr&&c.expr.filters){c.expr.filters.hidden=function(a){var b=
a.offsetWidth,d=a.offsetHeight,f=a.nodeName.toLowerCase()==="tr";return b===0&&d===0&&!f?true:b>0&&d>0&&!f?false:c.curCSS(a,"display")==="none"};c.expr.filters.visible=function(a){return!c.expr.filters.hidden(a)}}var sb=J(),tb=/<script(.|\s)*?\/script>/gi,ub=/select|textarea/i,vb=/color|date|datetime|email|hidden|month|number|password|range|search|tel|text|time|url|week/i,N=/=\?(&|$)/,ka=/\?/,wb=/(\?|&)_=.*?(&|$)/,xb=/^(\w+:)?\/\/([^\/?#]+)/,yb=/%20/g,zb=c.fn.load;c.fn.extend({load:function(a,b,d){if(typeof a!==
"string")return zb.call(this,a);else if(!this.length)return this;var f=a.indexOf(" ");if(f>=0){var e=a.slice(f,a.length);a=a.slice(0,f)}f="GET";if(b)if(c.isFunction(b)){d=b;b=null}else if(typeof b==="object"){b=c.param(b,c.ajaxSettings.traditional);f="POST"}var j=this;c.ajax({url:a,type:f,dataType:"html",data:b,complete:function(i,o){if(o==="success"||o==="notmodified")j.html(e?c("<div />").append(i.responseText.replace(tb,"")).find(e):i.responseText);d&&j.each(d,[i.responseText,o,i])}});return this},
serialize:function(){return c.param(this.serializeArray())},serializeArray:function(){return this.map(function(){return this.elements?c.makeArray(this.elements):this}).filter(function(){return this.name&&!this.disabled&&(this.checked||ub.test(this.nodeName)||vb.test(this.type))}).map(function(a,b){a=c(this).val();return a==null?null:c.isArray(a)?c.map(a,function(d){return{name:b.name,value:d}}):{name:b.name,value:a}}).get()}});c.each("ajaxStart ajaxStop ajaxComplete ajaxError ajaxSuccess ajaxSend".split(" "),
function(a,b){c.fn[b]=function(d){return this.bind(b,d)}});c.extend({get:function(a,b,d,f){if(c.isFunction(b)){f=f||d;d=b;b=null}return c.ajax({type:"GET",url:a,data:b,success:d,dataType:f})},getScript:function(a,b){return c.get(a,null,b,"script")},getJSON:function(a,b,d){return c.get(a,b,d,"json")},post:function(a,b,d,f){if(c.isFunction(b)){f=f||d;d=b;b={}}return c.ajax({type:"POST",url:a,data:b,success:d,dataType:f})},ajaxSetup:function(a){c.extend(c.ajaxSettings,a)},ajaxSettings:{url:location.href,
global:true,type:"GET",contentType:"application/x-www-form-urlencoded",processData:true,async:true,xhr:A.XMLHttpRequest&&(A.location.protocol!=="file:"||!A.ActiveXObject)?function(){return new A.XMLHttpRequest}:function(){try{return new A.ActiveXObject("Microsoft.XMLHTTP")}catch(a){}},accepts:{xml:"application/xml, text/xml",html:"text/html",script:"text/javascript, application/javascript",json:"application/json, text/javascript",text:"text/plain",_default:"*/*"}},lastModified:{},etag:{},ajax:function(a){function b(){e.success&&
e.success.call(k,o,i,x);e.global&&f("ajaxSuccess",[x,e])}function d(){e.complete&&e.complete.call(k,x,i);e.global&&f("ajaxComplete",[x,e]);e.global&&!--c.active&&c.event.trigger("ajaxStop")}function f(q,p){(e.context?c(e.context):c.event).trigger(q,p)}var e=c.extend(true,{},c.ajaxSettings,a),j,i,o,k=a&&a.context||e,n=e.type.toUpperCase();if(e.data&&e.processData&&typeof e.data!=="string")e.data=c.param(e.data,e.traditional);if(e.dataType==="jsonp"){if(n==="GET")N.test(e.url)||(e.url+=(ka.test(e.url)?
"&":"?")+(e.jsonp||"callback")+"=?");else if(!e.data||!N.test(e.data))e.data=(e.data?e.data+"&":"")+(e.jsonp||"callback")+"=?";e.dataType="json"}if(e.dataType==="json"&&(e.data&&N.test(e.data)||N.test(e.url))){j=e.jsonpCallback||"jsonp"+sb++;if(e.data)e.data=(e.data+"").replace(N,"="+j+"$1");e.url=e.url.replace(N,"="+j+"$1");e.dataType="script";A[j]=A[j]||function(q){o=q;b();d();A[j]=w;try{delete A[j]}catch(p){}z&&z.removeChild(C)}}if(e.dataType==="script"&&e.cache===null)e.cache=false;if(e.cache===
false&&n==="GET"){var r=J(),u=e.url.replace(wb,"$1_="+r+"$2");e.url=u+(u===e.url?(ka.test(e.url)?"&":"?")+"_="+r:"")}if(e.data&&n==="GET")e.url+=(ka.test(e.url)?"&":"?")+e.data;e.global&&!c.active++&&c.event.trigger("ajaxStart");r=(r=xb.exec(e.url))&&(r[1]&&r[1]!==location.protocol||r[2]!==location.host);if(e.dataType==="script"&&n==="GET"&&r){var z=s.getElementsByTagName("head")[0]||s.documentElement,C=s.createElement("script");C.src=e.url;if(e.scriptCharset)C.charset=e.scriptCharset;if(!j){var B=
false;C.onload=C.onreadystatechange=function(){if(!B&&(!this.readyState||this.readyState==="loaded"||this.readyState==="complete")){B=true;b();d();C.onload=C.onreadystatechange=null;z&&C.parentNode&&z.removeChild(C)}}}z.insertBefore(C,z.firstChild);return w}var E=false,x=e.xhr();if(x){e.username?x.open(n,e.url,e.async,e.username,e.password):x.open(n,e.url,e.async);try{if(e.data||a&&a.contentType)x.setRequestHeader("Content-Type",e.contentType);if(e.ifModified){c.lastModified[e.url]&&x.setRequestHeader("If-Modified-Since",
c.lastModified[e.url]);c.etag[e.url]&&x.setRequestHeader("If-None-Match",c.etag[e.url])}r||x.setRequestHeader("X-Requested-With","XMLHttpRequest");x.setRequestHeader("Accept",e.dataType&&e.accepts[e.dataType]?e.accepts[e.dataType]+", */*":e.accepts._default)}catch(ga){}if(e.beforeSend&&e.beforeSend.call(k,x,e)===false){e.global&&!--c.active&&c.event.trigger("ajaxStop");x.abort();return false}e.global&&f("ajaxSend",[x,e]);var g=x.onreadystatechange=function(q){if(!x||x.readyState===0||q==="abort"){E||
d();E=true;if(x)x.onreadystatechange=c.noop}else if(!E&&x&&(x.readyState===4||q==="timeout")){E=true;x.onreadystatechange=c.noop;i=q==="timeout"?"timeout":!c.httpSuccess(x)?"error":e.ifModified&&c.httpNotModified(x,e.url)?"notmodified":"success";var p;if(i==="success")try{o=c.httpData(x,e.dataType,e)}catch(v){i="parsererror";p=v}if(i==="success"||i==="notmodified")j||b();else c.handleError(e,x,i,p);d();q==="timeout"&&x.abort();if(e.async)x=null}};try{var h=x.abort;x.abort=function(){x&&h.call(x);
g("abort")}}catch(l){}e.async&&e.timeout>0&&setTimeout(function(){x&&!E&&g("timeout")},e.timeout);try{x.send(n==="POST"||n==="PUT"||n==="DELETE"?e.data:null)}catch(m){c.handleError(e,x,null,m);d()}e.async||g();return x}},handleError:function(a,b,d,f){if(a.error)a.error.call(a.context||a,b,d,f);if(a.global)(a.context?c(a.context):c.event).trigger("ajaxError",[b,a,f])},active:0,httpSuccess:function(a){try{return!a.status&&location.protocol==="file:"||a.status>=200&&a.status<300||a.status===304||a.status===
1223||a.status===0}catch(b){}return false},httpNotModified:function(a,b){var d=a.getResponseHeader("Last-Modified"),f=a.getResponseHeader("Etag");if(d)c.lastModified[b]=d;if(f)c.etag[b]=f;return a.status===304||a.status===0},httpData:function(a,b,d){var f=a.getResponseHeader("content-type")||"",e=b==="xml"||!b&&f.indexOf("xml")>=0;a=e?a.responseXML:a.responseText;e&&a.documentElement.nodeName==="parsererror"&&c.error("parsererror");if(d&&d.dataFilter)a=d.dataFilter(a,b);if(typeof a==="string")if(b===
"json"||!b&&f.indexOf("json")>=0)a=c.parseJSON(a);else if(b==="script"||!b&&f.indexOf("javascript")>=0)c.globalEval(a);return a},param:function(a,b){function d(i,o){if(c.isArray(o))c.each(o,function(k,n){b||/\[\]$/.test(i)?f(i,n):d(i+"["+(typeof n==="object"||c.isArray(n)?k:"")+"]",n)});else!b&&o!=null&&typeof o==="object"?c.each(o,function(k,n){d(i+"["+k+"]",n)}):f(i,o)}function f(i,o){o=c.isFunction(o)?o():o;e[e.length]=encodeURIComponent(i)+"="+encodeURIComponent(o)}var e=[];if(b===w)b=c.ajaxSettings.traditional;
if(c.isArray(a)||a.jquery)c.each(a,function(){f(this.name,this.value)});else for(var j in a)d(j,a[j]);return e.join("&").replace(yb,"+")}});var la={},Ab=/toggle|show|hide/,Bb=/^([+-]=)?([\d+-.]+)(.*)$/,W,va=[["height","marginTop","marginBottom","paddingTop","paddingBottom"],["width","marginLeft","marginRight","paddingLeft","paddingRight"],["opacity"]];c.fn.extend({show:function(a,b){if(a||a===0)return this.animate(K("show",3),a,b);else{a=0;for(b=this.length;a<b;a++){var d=c.data(this[a],"olddisplay");
this[a].style.display=d||"";if(c.css(this[a],"display")==="none"){d=this[a].nodeName;var f;if(la[d])f=la[d];else{var e=c("<"+d+" />").appendTo("body");f=e.css("display");if(f==="none")f="block";e.remove();la[d]=f}c.data(this[a],"olddisplay",f)}}a=0;for(b=this.length;a<b;a++)this[a].style.display=c.data(this[a],"olddisplay")||"";return this}},hide:function(a,b){if(a||a===0)return this.animate(K("hide",3),a,b);else{a=0;for(b=this.length;a<b;a++){var d=c.data(this[a],"olddisplay");!d&&d!=="none"&&c.data(this[a],
"olddisplay",c.css(this[a],"display"))}a=0;for(b=this.length;a<b;a++)this[a].style.display="none";return this}},_toggle:c.fn.toggle,toggle:function(a,b){var d=typeof a==="boolean";if(c.isFunction(a)&&c.isFunction(b))this._toggle.apply(this,arguments);else a==null||d?this.each(function(){var f=d?a:c(this).is(":hidden");c(this)[f?"show":"hide"]()}):this.animate(K("toggle",3),a,b);return this},fadeTo:function(a,b,d){return this.filter(":hidden").css("opacity",0).show().end().animate({opacity:b},a,d)},
animate:function(a,b,d,f){var e=c.speed(b,d,f);if(c.isEmptyObject(a))return this.each(e.complete);return this[e.queue===false?"each":"queue"](function(){var j=c.extend({},e),i,o=this.nodeType===1&&c(this).is(":hidden"),k=this;for(i in a){var n=i.replace(ia,ja);if(i!==n){a[n]=a[i];delete a[i];i=n}if(a[i]==="hide"&&o||a[i]==="show"&&!o)return j.complete.call(this);if((i==="height"||i==="width")&&this.style){j.display=c.css(this,"display");j.overflow=this.style.overflow}if(c.isArray(a[i])){(j.specialEasing=
j.specialEasing||{})[i]=a[i][1];a[i]=a[i][0]}}if(j.overflow!=null)this.style.overflow="hidden";j.curAnim=c.extend({},a);c.each(a,function(r,u){var z=new c.fx(k,j,r);if(Ab.test(u))z[u==="toggle"?o?"show":"hide":u](a);else{var C=Bb.exec(u),B=z.cur(true)||0;if(C){u=parseFloat(C[2]);var E=C[3]||"px";if(E!=="px"){k.style[r]=(u||1)+E;B=(u||1)/z.cur(true)*B;k.style[r]=B+E}if(C[1])u=(C[1]==="-="?-1:1)*u+B;z.custom(B,u,E)}else z.custom(B,u,"")}});return true})},stop:function(a,b){var d=c.timers;a&&this.queue([]);
this.each(function(){for(var f=d.length-1;f>=0;f--)if(d[f].elem===this){b&&d[f](true);d.splice(f,1)}});b||this.dequeue();return this}});c.each({slideDown:K("show",1),slideUp:K("hide",1),slideToggle:K("toggle",1),fadeIn:{opacity:"show"},fadeOut:{opacity:"hide"}},function(a,b){c.fn[a]=function(d,f){return this.animate(b,d,f)}});c.extend({speed:function(a,b,d){var f=a&&typeof a==="object"?a:{complete:d||!d&&b||c.isFunction(a)&&a,duration:a,easing:d&&b||b&&!c.isFunction(b)&&b};f.duration=c.fx.off?0:typeof f.duration===
"number"?f.duration:c.fx.speeds[f.duration]||c.fx.speeds._default;f.old=f.complete;f.complete=function(){f.queue!==false&&c(this).dequeue();c.isFunction(f.old)&&f.old.call(this)};return f},easing:{linear:function(a,b,d,f){return d+f*a},swing:function(a,b,d,f){return(-Math.cos(a*Math.PI)/2+0.5)*f+d}},timers:[],fx:function(a,b,d){this.options=b;this.elem=a;this.prop=d;if(!b.orig)b.orig={}}});c.fx.prototype={update:function(){this.options.step&&this.options.step.call(this.elem,this.now,this);(c.fx.step[this.prop]||
c.fx.step._default)(this);if((this.prop==="height"||this.prop==="width")&&this.elem.style)this.elem.style.display="block"},cur:function(a){if(this.elem[this.prop]!=null&&(!this.elem.style||this.elem.style[this.prop]==null))return this.elem[this.prop];return(a=parseFloat(c.css(this.elem,this.prop,a)))&&a>-10000?a:parseFloat(c.curCSS(this.elem,this.prop))||0},custom:function(a,b,d){function f(j){return e.step(j)}this.startTime=J();this.start=a;this.end=b;this.unit=d||this.unit||"px";this.now=this.start;
this.pos=this.state=0;var e=this;f.elem=this.elem;if(f()&&c.timers.push(f)&&!W)W=setInterval(c.fx.tick,13)},show:function(){this.options.orig[this.prop]=c.style(this.elem,this.prop);this.options.show=true;this.custom(this.prop==="width"||this.prop==="height"?1:0,this.cur());c(this.elem).show()},hide:function(){this.options.orig[this.prop]=c.style(this.elem,this.prop);this.options.hide=true;this.custom(this.cur(),0)},step:function(a){var b=J(),d=true;if(a||b>=this.options.duration+this.startTime){this.now=
this.end;this.pos=this.state=1;this.update();this.options.curAnim[this.prop]=true;for(var f in this.options.curAnim)if(this.options.curAnim[f]!==true)d=false;if(d){if(this.options.display!=null){this.elem.style.overflow=this.options.overflow;a=c.data(this.elem,"olddisplay");this.elem.style.display=a?a:this.options.display;if(c.css(this.elem,"display")==="none")this.elem.style.display="block"}this.options.hide&&c(this.elem).hide();if(this.options.hide||this.options.show)for(var e in this.options.curAnim)c.style(this.elem,
e,this.options.orig[e]);this.options.complete.call(this.elem)}return false}else{e=b-this.startTime;this.state=e/this.options.duration;a=this.options.easing||(c.easing.swing?"swing":"linear");this.pos=c.easing[this.options.specialEasing&&this.options.specialEasing[this.prop]||a](this.state,e,0,1,this.options.duration);this.now=this.start+(this.end-this.start)*this.pos;this.update()}return true}};c.extend(c.fx,{tick:function(){for(var a=c.timers,b=0;b<a.length;b++)a[b]()||a.splice(b--,1);a.length||
c.fx.stop()},stop:function(){clearInterval(W);W=null},speeds:{slow:600,fast:200,_default:400},step:{opacity:function(a){c.style(a.elem,"opacity",a.now)},_default:function(a){if(a.elem.style&&a.elem.style[a.prop]!=null)a.elem.style[a.prop]=(a.prop==="width"||a.prop==="height"?Math.max(0,a.now):a.now)+a.unit;else a.elem[a.prop]=a.now}}});if(c.expr&&c.expr.filters)c.expr.filters.animated=function(a){return c.grep(c.timers,function(b){return a===b.elem}).length};c.fn.offset="getBoundingClientRect"in s.documentElement?
function(a){var b=this[0];if(a)return this.each(function(e){c.offset.setOffset(this,a,e)});if(!b||!b.ownerDocument)return null;if(b===b.ownerDocument.body)return c.offset.bodyOffset(b);var d=b.getBoundingClientRect(),f=b.ownerDocument;b=f.body;f=f.documentElement;return{top:d.top+(self.pageYOffset||c.support.boxModel&&f.scrollTop||b.scrollTop)-(f.clientTop||b.clientTop||0),left:d.left+(self.pageXOffset||c.support.boxModel&&f.scrollLeft||b.scrollLeft)-(f.clientLeft||b.clientLeft||0)}}:function(a){var b=
this[0];if(a)return this.each(function(r){c.offset.setOffset(this,a,r)});if(!b||!b.ownerDocument)return null;if(b===b.ownerDocument.body)return c.offset.bodyOffset(b);c.offset.initialize();var d=b.offsetParent,f=b,e=b.ownerDocument,j,i=e.documentElement,o=e.body;f=(e=e.defaultView)?e.getComputedStyle(b,null):b.currentStyle;for(var k=b.offsetTop,n=b.offsetLeft;(b=b.parentNode)&&b!==o&&b!==i;){if(c.offset.supportsFixedPosition&&f.position==="fixed")break;j=e?e.getComputedStyle(b,null):b.currentStyle;
k-=b.scrollTop;n-=b.scrollLeft;if(b===d){k+=b.offsetTop;n+=b.offsetLeft;if(c.offset.doesNotAddBorder&&!(c.offset.doesAddBorderForTableAndCells&&/^t(able|d|h)$/i.test(b.nodeName))){k+=parseFloat(j.borderTopWidth)||0;n+=parseFloat(j.borderLeftWidth)||0}f=d;d=b.offsetParent}if(c.offset.subtractsBorderForOverflowNotVisible&&j.overflow!=="visible"){k+=parseFloat(j.borderTopWidth)||0;n+=parseFloat(j.borderLeftWidth)||0}f=j}if(f.position==="relative"||f.position==="static"){k+=o.offsetTop;n+=o.offsetLeft}if(c.offset.supportsFixedPosition&&
f.position==="fixed"){k+=Math.max(i.scrollTop,o.scrollTop);n+=Math.max(i.scrollLeft,o.scrollLeft)}return{top:k,left:n}};c.offset={initialize:function(){var a=s.body,b=s.createElement("div"),d,f,e,j=parseFloat(c.curCSS(a,"marginTop",true))||0;c.extend(b.style,{position:"absolute",top:0,left:0,margin:0,border:0,width:"1px",height:"1px",visibility:"hidden"});b.innerHTML="<div style='position:absolute;top:0;left:0;margin:0;border:5px solid #000;padding:0;width:1px;height:1px;'><div></div></div><table style='position:absolute;top:0;left:0;margin:0;border:5px solid #000;padding:0;width:1px;height:1px;' cellpadding='0' cellspacing='0'><tr><td></td></tr></table>";
a.insertBefore(b,a.firstChild);d=b.firstChild;f=d.firstChild;e=d.nextSibling.firstChild.firstChild;this.doesNotAddBorder=f.offsetTop!==5;this.doesAddBorderForTableAndCells=e.offsetTop===5;f.style.position="fixed";f.style.top="20px";this.supportsFixedPosition=f.offsetTop===20||f.offsetTop===15;f.style.position=f.style.top="";d.style.overflow="hidden";d.style.position="relative";this.subtractsBorderForOverflowNotVisible=f.offsetTop===-5;this.doesNotIncludeMarginInBodyOffset=a.offsetTop!==j;a.removeChild(b);
c.offset.initialize=c.noop},bodyOffset:function(a){var b=a.offsetTop,d=a.offsetLeft;c.offset.initialize();if(c.offset.doesNotIncludeMarginInBodyOffset){b+=parseFloat(c.curCSS(a,"marginTop",true))||0;d+=parseFloat(c.curCSS(a,"marginLeft",true))||0}return{top:b,left:d}},setOffset:function(a,b,d){if(/static/.test(c.curCSS(a,"position")))a.style.position="relative";var f=c(a),e=f.offset(),j=parseInt(c.curCSS(a,"top",true),10)||0,i=parseInt(c.curCSS(a,"left",true),10)||0;if(c.isFunction(b))b=b.call(a,
d,e);d={top:b.top-e.top+j,left:b.left-e.left+i};"using"in b?b.using.call(a,d):f.css(d)}};c.fn.extend({position:function(){if(!this[0])return null;var a=this[0],b=this.offsetParent(),d=this.offset(),f=/^body|html$/i.test(b[0].nodeName)?{top:0,left:0}:b.offset();d.top-=parseFloat(c.curCSS(a,"marginTop",true))||0;d.left-=parseFloat(c.curCSS(a,"marginLeft",true))||0;f.top+=parseFloat(c.curCSS(b[0],"borderTopWidth",true))||0;f.left+=parseFloat(c.curCSS(b[0],"borderLeftWidth",true))||0;return{top:d.top-
f.top,left:d.left-f.left}},offsetParent:function(){return this.map(function(){for(var a=this.offsetParent||s.body;a&&!/^body|html$/i.test(a.nodeName)&&c.css(a,"position")==="static";)a=a.offsetParent;return a})}});c.each(["Left","Top"],function(a,b){var d="scroll"+b;c.fn[d]=function(f){var e=this[0],j;if(!e)return null;if(f!==w)return this.each(function(){if(j=wa(this))j.scrollTo(!a?f:c(j).scrollLeft(),a?f:c(j).scrollTop());else this[d]=f});else return(j=wa(e))?"pageXOffset"in j?j[a?"pageYOffset":
"pageXOffset"]:c.support.boxModel&&j.document.documentElement[d]||j.document.body[d]:e[d]}});c.each(["Height","Width"],function(a,b){var d=b.toLowerCase();c.fn["inner"+b]=function(){return this[0]?c.css(this[0],d,false,"padding"):null};c.fn["outer"+b]=function(f){return this[0]?c.css(this[0],d,false,f?"margin":"border"):null};c.fn[d]=function(f){var e=this[0];if(!e)return f==null?null:this;if(c.isFunction(f))return this.each(function(j){var i=c(this);i[d](f.call(this,j,i[d]()))});return"scrollTo"in
e&&e.document?e.document.compatMode==="CSS1Compat"&&e.document.documentElement["client"+b]||e.document.body["client"+b]:e.nodeType===9?Math.max(e.documentElement["client"+b],e.body["scroll"+b],e.documentElement["scroll"+b],e.body["offset"+b],e.documentElement["offset"+b]):f===w?c.css(e,d):this.css(d,typeof f==="string"?f:f+"px")}});A.jQuery=A.$=c})(window);
```
