## Supplementary material 1 for "Termite diet rather than geographical origin determines the microbiome composition and functional genetic structure of nests from South American and African representatives, as revealed by a multiomics approach": jquery.albumSlider.min.js

```
(function($){$.fn.albumSlider=function(j){return this.each(function(){var b=$.extend({step:2,imgContainer:'div.fullview',listContainer:'ul.imglist',event:'mouseover',direction:'v'},j||{});var c=$(b.imgContainer,this),$list=$(b.listContainer,this),currId=0,currPage=0,size=$list.children().length-1,pageSize=Math.floor(size/b.step);var f=b.direction=='v';var g=f?'top':'left';var h=(size>=b.step)?$('li',$list).eq(b.step).offset()[g]-$('li',$list).eq(0).offset()[g]:0;var i=function(){var a=$(this);if(a.is('.current')){return false}$('img',c).fadeOut(800,function(){$(this).remove()});$('<img>').hide().attr('src',$('a',a).attr('href')).appendTo(c).fadeIn(800);a.addClass('current').siblings().removeClass('current');return false};$.proxy(i,$('li',$list).eq(0))();$list.delegate('li',b.event,$.proxy(i)).bind('moveforward movebackward',function(e){var a=e.type=='moveforward';if(a){currId+=b.step;if(currId>size){currId=size}if(++currPage>pageSize){currPage=pageSize;return false}}else{currId-=b.step;if(currId<0){currId=0}if(--currPage<0){currPage=0;return false}};var d=(a?'-=':'+=')+h;$(this).stop(true,true).animate(f?{top:d}:{left:d},500,function(){$.proxy(i,$('li',$list).eq(currId))()})});$('div.button',this).click(function(){$list.trigger($(this).is('.moveforward')?'moveforward':'movebackward')})})}})(jQuery);
```
