## Supplementary material 1 for "Termite diet rather than geographical origin determines the microbiome composition and functional genetic structure of nests from South American and African representatives, as revealed by a multiomics approach": scrolltop.js

```
$(document).ready(function(){
	//1.Ê×ÏÈ½«#goTopBtnÒþ²Ø
	$("#goTopBtn").hide();

	//2.¸ø´°¿ÚÌí¼ÓÊÂ¼þ£º¹ö¶¯Ê±´¥·¢
	$(window).scroll(function(){
		//µ±¹ö¶¯ÌõµÄÎ»ÖÃ´¦ÓÚ¾à¶¥²¿0ÏñËØÒÔÉÏÊ±£¬·µ»Ø¶¥²¿°´Å¥³öÏÖ£¬·ñÔòÏûÊ§
		if ($(window).scrollTop()>0){
			$("#goTopBtn").fadeIn(1500);
			}
		else{
			$("#goTopBtn").fadeOut(1500);
			}
	});
	
	//3.µ±µã»÷·µ»Ø¶¥²¿°´Å¥ºó£¬»Øµ½Ò³Ãæ¶¥²¿Î»ÖÃ
	$("#goTopBtn").click(function(){
		$('body,html').animate({scrollTop:0},1000);
	});
});
```
