## Supplementary material 1 for "Termite diet rather than geographical origin determines the microbiome composition and functional genetic structure of nests from South American and African representatives, as revealed by a multiomics approach": X201SC20094093-Z01-F001_sequence_Report.html

 X201SC20094093-Z01-F001 Analysis Report 


- Workflow
  - Experiment process and sequencing
  - Analysis process
- Results
  - Sequencing Quality
  - Summary of Sequencing
- Appendix
  - Introduction of Data Format
  - Paired-end reads assembly and quality control
  - References


### X201SC20094093-Z01-F001 QC Analysis Report

16-November-2020

- Workflow

- Experiment process and sequencing
- Analysis process

- Results and Instructions

- Data Quality Control

- Distribution of Sequencing Quality
- Distribution of Sequencing Error Rate
- Distribution of Tags Length

- Summary of Data Information
- Appendix

- Introduction of Sequenced Data Format
- Paired-end reads assembly and quality control
- References

Novogene Co., Ltd


---

  

#### A. Workflow

From the DNA samples to the final data, each step, including sample test, PCR, library preparation, and sequencing, influences the quality of the data, and data quality directly impacts the analysis results. To guarantee the reliability of the data, quality control (QC) is performed at each step of the procedure.
The workflow is as follows:

##### 1 Sample Quality Control

There are three main methods of QC for DNA samples:

(1) Nanodrop: tests DNA purity (OD260/OD280).

(2) Agarose Gel Electrophoresis: tests DNA degradation and potential contamination.

(3) Qubit 2.0: quantifies the DNA concentration precisely.


##### 2 Library Construction and Sequencing

PCR were performed using specific primers with the barcodes. The construction of the DNA libraries is through the processes of end repairing, adding A to tails, purification and etc. Libraries was performed on a paired-end Illumina platform to generate 250bp paired-end raw reads. The principle of library construction is as follows:

##### 3 Analysis process

Paired-end reads was assigned to samples based on their unique barcode and truncated by cutting off the barcode and primer sequence. Paired-end reads were merged using FLASH based on the reads overlap. Quality filtering on the raw tags were performed under specific filtering conditions to obtain the high-quality clean tags.

Novogene Co., Ltd


---

  

#### B. Results and Instructions

##### 1 Data Quality Control

###### 1.1 Distribution of Sequencing Quality

The “e” represents the sequence error rate and Q_phred_ represents the base quality value,Q_phred_=-10log_10_(e).
The relationship between sequencing error rate (e) and sequencing base quality value (Q_phred_) is as below:

| Phred score | error base | right base | Q-score |
| --- | --- | --- | --- |
| 10 | 1/10 | 90% | Q10 |
| 20 | 1/100 | 99% | Q20 |
| 30 | 1/1000 | 99.9% | Q30 |
| 40 | 1/10000 | 99.99% | Q40 |


The distribution of sequencing quality of the effective tags can be seen in **Fig.1**:

**Fig.1 Distribution of Sequencing Quality**

The base position is on the horizontal axis and the sequencing quality is on the vertical axis

Novogene Co., Ltd


---

  


###### 1.2 Distribution of Sequencing Error Rate

For Illumina SBS technology, the distribution of sequencing error rate has two features:

(1) Error rate grows with sequenced reads extension because of the consumption of sequencing reagent. The phenomenon is common in the Illumina high-throughput sequencing platform (Erlich Y. et al. 2008; Jiang et al. 2011).

(2) The reason for the high error rate of the first six bases is that the random hex-primers and RNA template bind incompletely in the process of cDNA synthesis (Jiang et al.2011).

The error rate of the effective tags is shown in **Fig.2**:

  

**Fig.2 Error Rate Distribution**

The base position is on the horizontal axis and the single base error rate is on the vertical axis

The first half part of the distribution is about for reads1 and the latter half part is about for reversed reads2

Novogene Co., Ltd


---

  


###### 1.3 Distribution of Tags Length

Before assembly, Paired-end reads were truncated by cutting off the barcode and primer sequence. The overlap of paired-end reads fluctuate reasonably according to the diversity of PCR products and cutted reads.

The distribution of length for effective tags is shown in **Fig.3**:

**Fig.3 Effective Tags Length Distribution**

The tags length is on the horizontal axis and the tags number is on the vertical axis

Novogene Co., Ltd


---

  


##### 2 Summary of Sequencing Data Information

Amplicon was performed on a paired-end Illumina platform to generate 250bp paired-end raw reads (Raw PE), and then assembled and pretreated to obtain Clean Tags. The chimeric sequences on Clean Tags are detected and removed to obtain the Effective Tags finally.

The data output of the above steps are shown in**Table 1**.

**Table 1 Data Quality Summary**

| Sample | Novo ID | Raw PE(#) | Combined(#) | Qualified(#) | Nochime(#) | Effective Rate(%) | AvgLen(nt) | Q20(%) | Q30(%) | GC Content(%) |
| --- | --- | --- | --- | --- | --- | --- | --- | --- | --- | --- |
| NasutsimFG | FKDN202550963-1A | 152,162 | 147,055 | 146,752 | 126,638 | 83.23 | 252 | 99.11 | 97.64 | 57.23 |
| CoptoBR | FKDN202550953-1A | 151,040 | 146,119 | 145,846 | 121,959 | 80.75 | 253 | 99.15 | 97.73 | 56.78 |
| NasutBR | FKDN202550969-1A | 164,768 | 159,572 | 158,976 | 122,132 | 74.12 | 253 | 98.74 | 96.57 | 56.36 |
| Cubiter2ML | FKDN202550956-1A | 153,640 | 149,729 | 149,426 | 129,358 | 84.20 | 252 | 99.21 | 97.73 | 57.70 |
| LabiotFG | FKDN202550961-1A | 151,111 | 147,202 | 146,926 | 126,636 | 83.80 | 253 | 99.26 | 97.88 | 57.94 |
| Anba1FG | FKDN202550947-1A | 160,689 | 156,780 | 156,527 | 135,201 | 84.14 | 253 | 99.32 | 98.04 | 58.23 |
| NasutspFG | FKDN202550968-1A | 156,367 | 152,463 | 152,180 | 130,681 | 83.57 | 252 | 99.27 | 97.89 | 57.19 |
| Nasutluj1CM | FKDN202550965-1A | 155,070 | 150,315 | 150,071 | 128,373 | 82.78 | 253 | 99.16 | 97.81 | 56.91 |
| NasutspCM | FKDN202550966-1A | 156,990 | 152,923 | 152,490 | 132,162 | 84.18 | 253 | 99.12 | 97.40 | 56.46 |
| Nasutluj2CM | FKDN202550967-1A | 155,759 | 151,476 | 151,210 | 128,273 | 82.35 | 253 | 99.23 | 97.86 | 57.20 |
| EmbiraFG | FKDN202550960-1A | 152,255 | 147,987 | 147,723 | 128,092 | 84.13 | 253 | 99.23 | 97.87 | 57.39 |
| Cubiter1CM | FKDN202550954-1A | 152,871 | 148,590 | 148,274 | 127,011 | 83.08 | 253 | 99.20 | 97.76 | 57.53 |
| CephCM | FKDN202550951-1A | 157,185 | 152,745 | 152,512 | 130,702 | 83.15 | 252 | 99.16 | 97.80 | 57.83 |
| Anba3FG | FKDN202550948-1A | 152,185 | 148,346 | 148,149 | 128,244 | 84.27 | 253 | 99.30 | 98.02 | 57.83 |
| MicroceroCM | FKDN202550962-1A | 156,539 | 152,071 | 151,797 | 133,657 | 85.38 | 253 | 99.22 | 97.84 | 57.03 |
| Cubiter2CM | FKDN202550959-1A | 153,116 | 149,230 | 148,948 | 128,695 | 84.05 | 253 | 99.24 | 97.85 | 58.26 |
| TaracuaFG20 | FKDN202550970-1A | 168,359 | 164,190 | 163,939 | 137,877 | 81.89 | 253 | 99.30 | 98.01 | 57.39 |
| Cubiter1ML | FKDN202550955-1A | 161,686 | 157,166 | 156,883 | 134,717 | 83.32 | 253 | 99.23 | 97.81 | 57.47 |
| Cubiter3ML | FKDN202550957-1A | 165,879 | 161,344 | 161,042 | 134,642 | 81.17 | 253 | 99.19 | 97.70 | 57.27 |
| Nasutluj3CM | FKDN202550964-1A | 168,354 | 163,269 | 162,978 | 142,192 | 84.46 | 253 | 99.13 | 97.72 | 56.93 |
| Taracua0FG19 | FKDN202550972-1A | 151,241 | 147,146 | 146,903 | 126,168 | 83.42 | 253 | 99.22 | 97.89 | 57.40 |
| Anba2FG | FKDN202550950-1A | 155,814 | 151,564 | 151,338 | 133,823 | 85.89 | 252 | 99.30 | 98.06 | 57.77 |
| Negative | FKDN202550975-1A | 49,427 | 44,414 | 44,273 | 36,406 | 73.66 | 253 | 98.89 | 97.10 | 55.15 |
| ConstrCM | FKDN202550952-1A | 161,528 | 156,899 | 156,688 | 134,317 | 83.15 | 251 | 99.18 | 97.86 | 56.82 |
| Cubiter4ML | FKDN202550958-1A | 161,687 | 157,529 | 157,232 | 133,520 | 82.58 | 253 | 99.28 | 97.95 | 57.85 |
| TermesFG | FKDN202550974-1A | 158,893 | 154,425 | 154,174 | 134,938 | 84.92 | 253 | 99.17 | 97.78 | 57.34 |
| SilvestFG | FKDN202550973-1A | 164,298 | 160,417 | 160,137 | 136,654 | 83.17 | 253 | 99.29 | 97.99 | 57.80 |
| Taracua4FG19 | FKDN202550971-1A | 167,842 | 163,525 | 163,239 | 138,987 | 82.81 | 253 | 99.27 | 97.94 | 57.86 |
| Anba4FG | FKDN202550949-1A | 148,098 | 144,608 | 144,389 | 126,669 | 85.53 | 252 | 99.33 | 98.07 | 57.83 |

Raw PE(#): Four rows are taken as a unit to calculate the total amount of read1 and read2 in raw data files split by barcode  
Combined(#): Raw tags generated from PE reads based on overlap  
Qualified(#): Clean tags after qualified from raw tags under specific filtering conditions  
Nochime(#): Effective tags after removing the chimera sequences  
Effective Rate(%): (Effective tags)/(Raw PE) \* 100%  
AvgLen(nt): Average length of effecitve tags  
Q20, Q30: (Base count of Phred value > 20 or 30) / (Total base count)  
GC content: (G & C base count) / (Total base count)


Novogene Co., Ltd


---

  


#### C. Appendix

##### 1. Introduction of Sequencing Data Format

The original data obtained from the high throughput sequencing platforms are transformed to sequenced reads by base calling. Raw data are recorded in a FASTQ file which contains sequenced reads and corresponding sequencing quality information. Every read in FASTQ format is stored in four lines as follows (Cock P.J.A. et al. 2010):

@HWI-ST1276:71:C1162ACXX:1:1101:1208:2458 1:N:0:CGATGT     
NAAGAACACGTTCGGTCACCTCAGCACACTTGTGAATGTCATGGGATCCAT  
+  
#55???BBBBB?BA@DEEFFCFFHHFFCFFHHHHHHHFAE0ECFFD/AEHH

Line 1 begins with a '@' character and is followed by a sequence identifier and an optional description (such as a FASTA title line).

Line 2 is the sequence of the read.

Line 3 begins with a '+' character and is optionally followed by the same sequence identifier (and any description) again.

Line 4 encodes the quality values for the bases in Line 2.

The details of Illumina sequence identifier are as follows:

|  |  |
| --- | --- |
| HWI-ST1276 | Instrument – unique identifier of the sequencer |
| 71 | run number – Run number on instrument |
| C1162ACXX | FlowCell ID – ID of flowcell |
| 1 | LaneNumber – positive integer |
| 1101 | TileNumber – positive integer |
| 1208 | X – x coordinate of the spot. Integer which can be negative |
| 2458 | Y – y coordinate of the spot. Integer which can be negative |
| 1 | ReadNumber - 1 for single reads; 1 or 2 for paired ends |
| N | whether it is filtered - NB: Y if the read is filtered out, not in the delivered fastq file, N otherwise |
| 0 | control number - 0 when none of the control bits are on, otherwise it is an even number |
| CGATGT | Illumina index sequences |

Novogene Co., Ltd


---

  


##### 2. Paired-end reads assembly and quality control

2.1 Data split: Paired-end reads was assigned to samples based on their unique barcode and truncated by cutting off the barcode and primer sequence.

2.2 Sequence assembly: Paired-end reads were merged using FLASH (V1.2.7, http://ccb.jhu.edu/software/FLASH/), a very fast and accurate analysis tool, which was designed to merge paired-end reads when at least some of the reads overlap the read generated from the opposite end of the same DNA fragment, and the splicing sequences were called raw tags.

2.3 Data Filtration:
Quality filtering on the raw tags were performed under specific filtering conditions to obtain the high-quality clean tags according to the Qiime(V1.7.0，http://qiime.org/scripts/split\_libraries\_fastq.html) quality controlled process.

2.4 Chimera removal:
The tags were compared with the reference database(Gold database，http://drive5.com/uchime/uchime\_download.html) using UCHIME algorithm(UCHIME Algorithm，http://www.drive5.com/usearch/manual/uchime\_algo.html) to detect chimera sequences, and then the chimera sequences were removed. Then the Effective Tags finally obtained.

Novogene Co., Ltd


---

  
