## Supplementary figures and images for "Termite diet rather than geographical origin determines the microbiome composition and functional genetic structure of nests from South American and African representatives, as revealed by a multiomics approach"

### 16s_pipeline_en.png

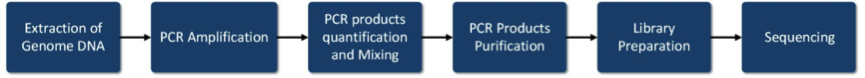

### album-slider-arrow_box.png

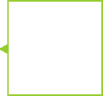

### Anba1FG.Error.png

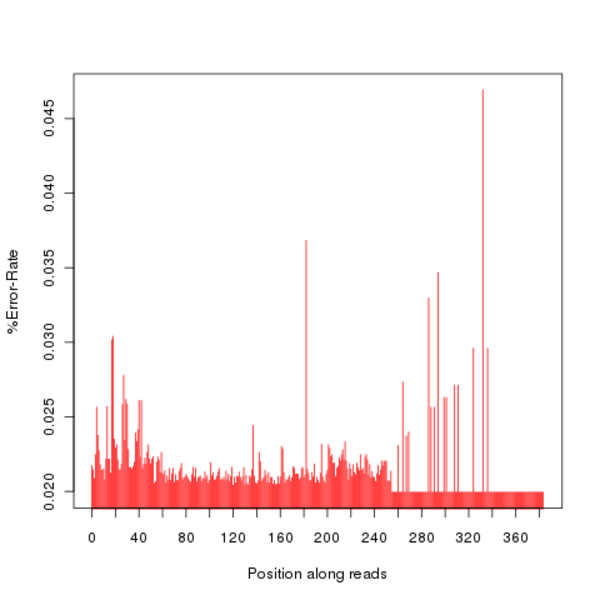

### Anba1FG.GC.png

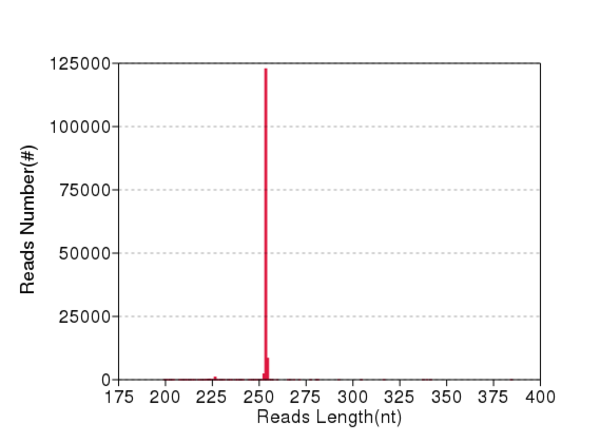

### Anba2FG.Error.JPEG

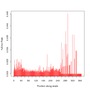

### Anba2FG.GC.png

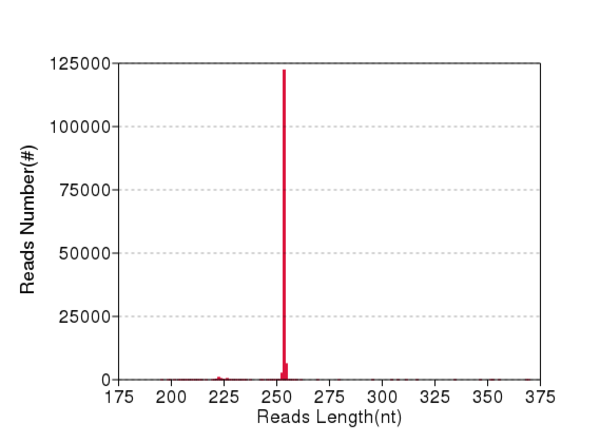

### Anba2FG.QD.png

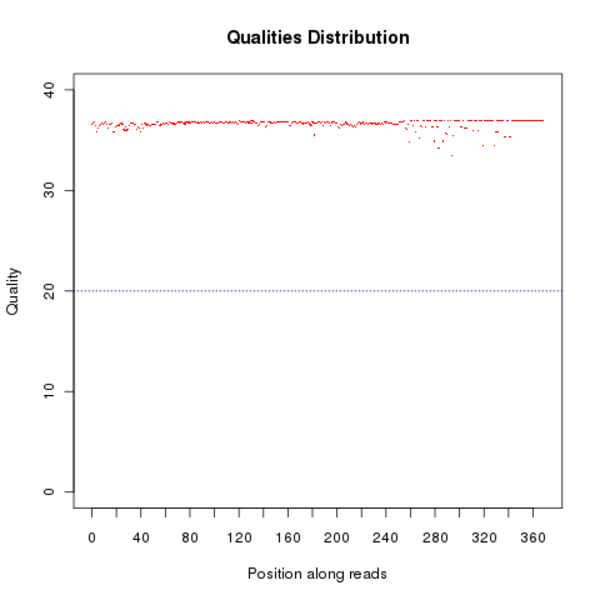

### Anba3FG.GC.png

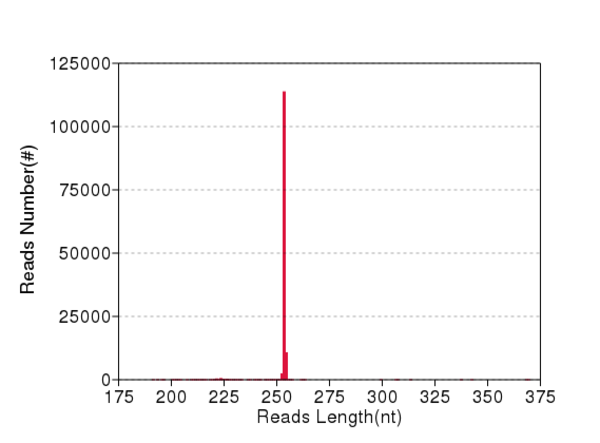

### Anba3FG.QD.png

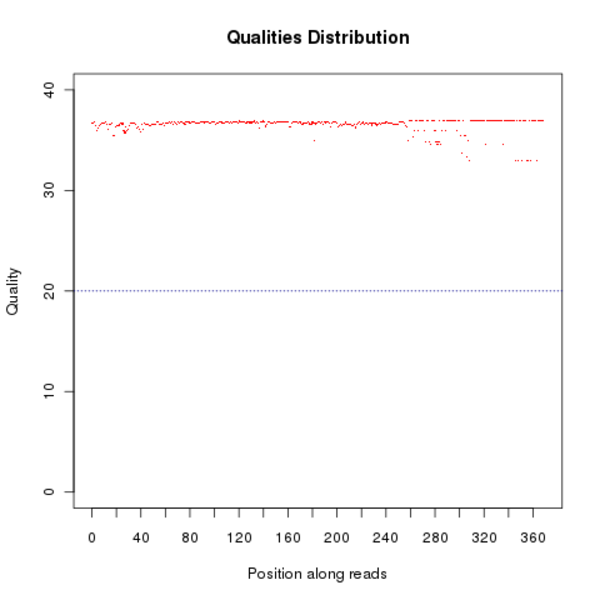

### Anba4FG.Error.JPEG

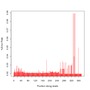

### Anba4FG.Error.png

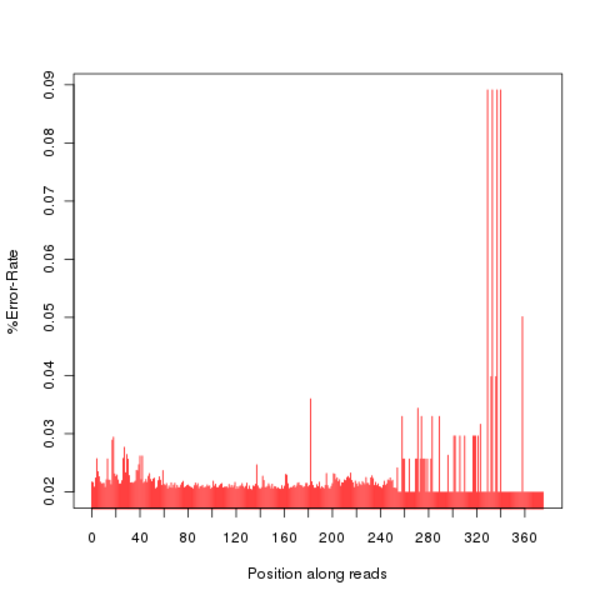

### Anba4FG.GC.JPEG

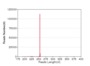

### Anba4FG.QD.JPEG

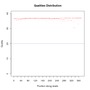

### CephCM.Error.JPEG

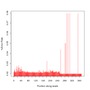

### CephCM.Error.png

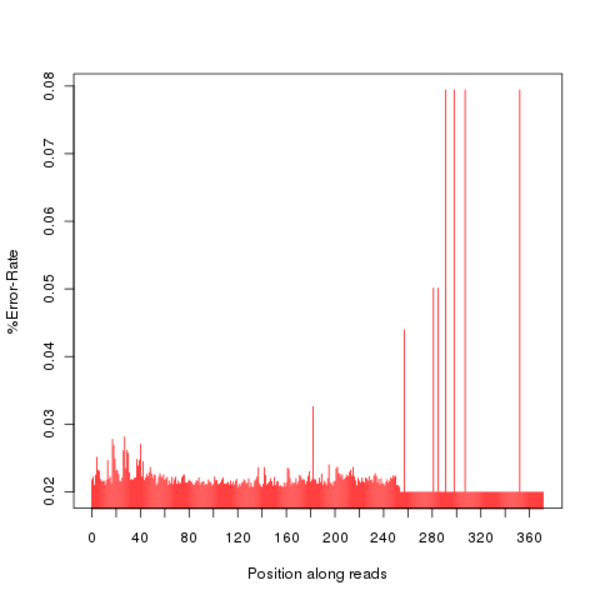

### CephCM.GC.JPEG

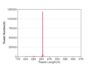

### CephCM.QD.JPEG

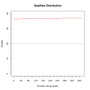

### close.gif

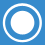

### ConstrCM.GC.png

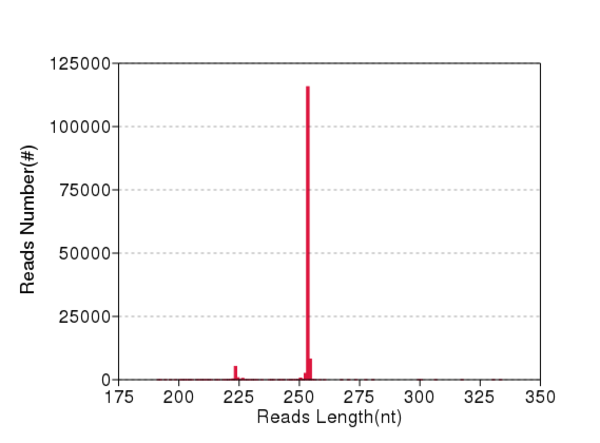

### CoptoBR.Error.png

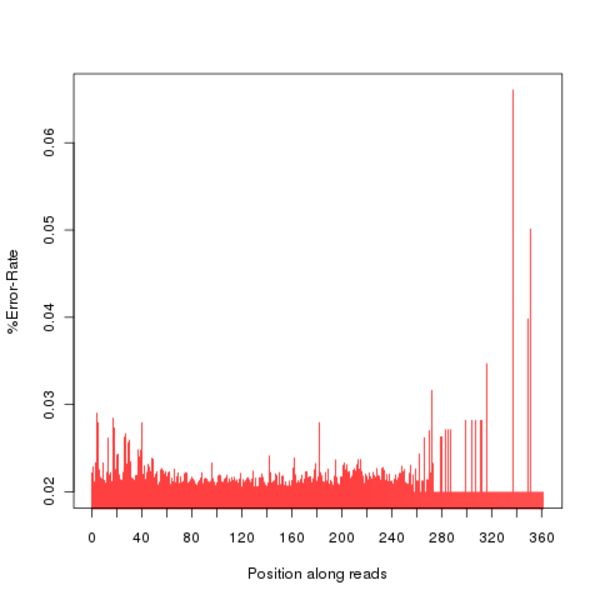

### CoptoBR.GC.png

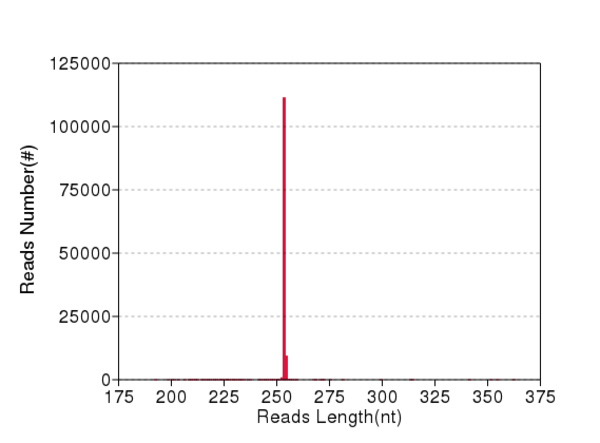

### CoptoBR.QD.JPEG

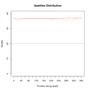

### Cubiter1CM.Error.JPEG

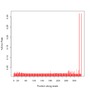

### Cubiter1CM.GC.JPEG

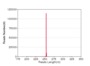

### Cubiter1CM.QD.JPEG

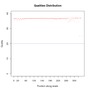

### Cubiter1CM.QD.png

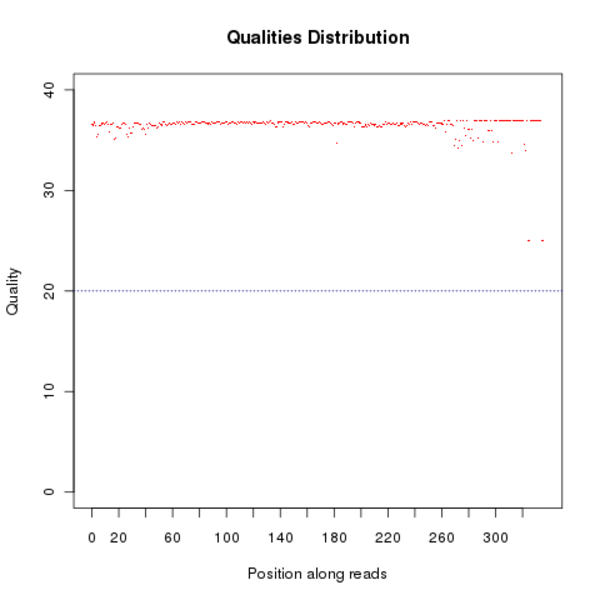

### Cubiter1ML.Error.png

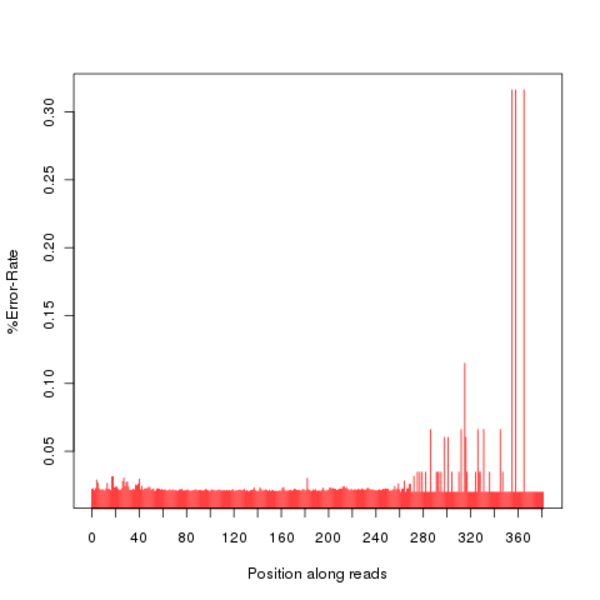

### Cubiter1ML.GC.JPEG

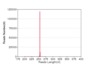

### Cubiter1ML.QD.JPEG

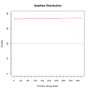

### Cubiter2CM.Error.png

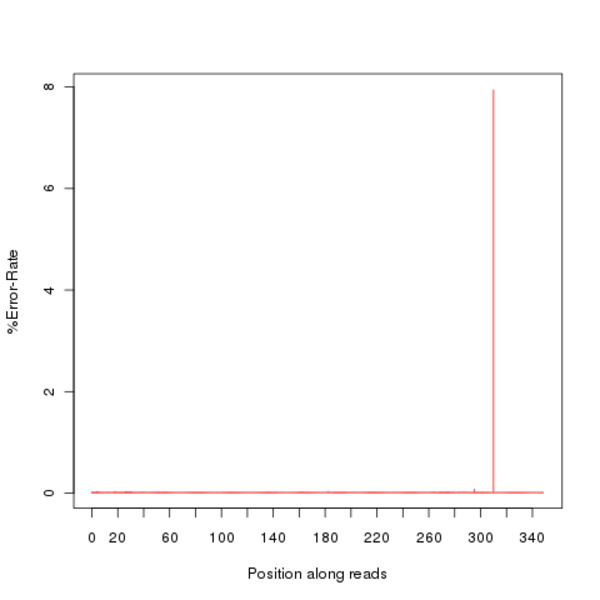

### Cubiter2ML.Error.JPEG

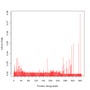

### Cubiter2ML.GC.png

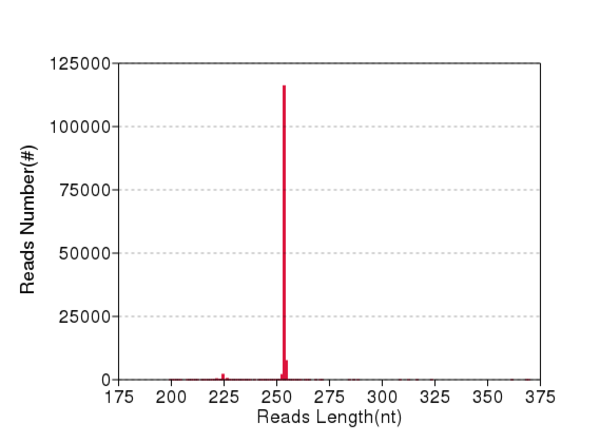

### Cubiter2ML.QD.JPEG

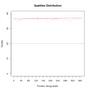

### Cubiter3ML.Error.JPEG

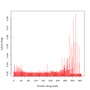

### Cubiter3ML.Error.png

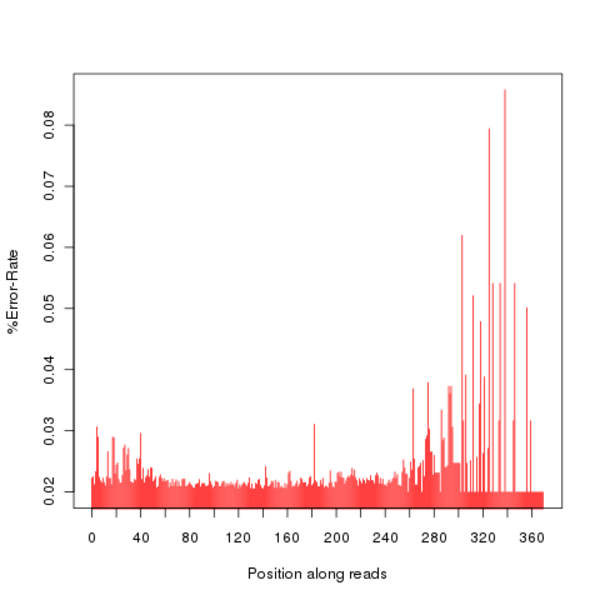

### Cubiter3ML.GC.JPEG

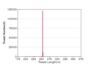

### Cubiter3ML.QD.JPEG

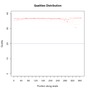

### Cubiter4ML.Error.png

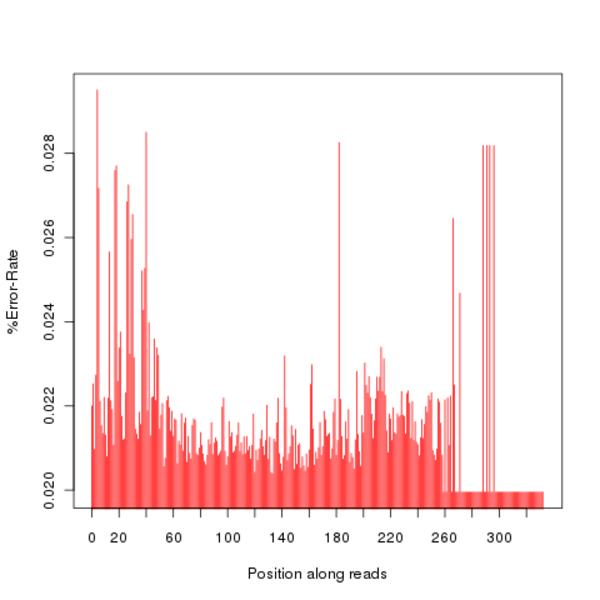

### Cubiter4ML.GC.JPEG

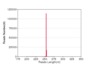

### Cubiter4ML.QD.JPEG

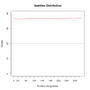

### EmbiraFG.Error.JPEG

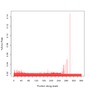

### EmbiraFG.GC.png

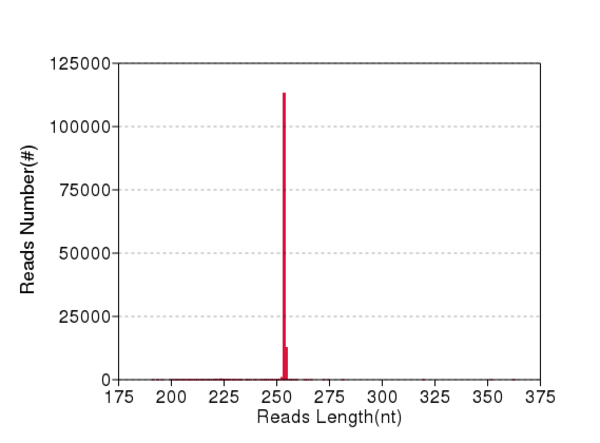

### exp_pipeline.png

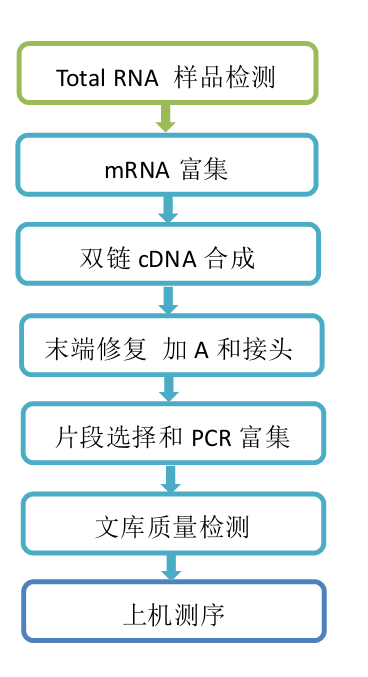

### LabiotFG.GC.JPEG

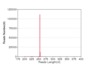

### LabiotFG.GC.png

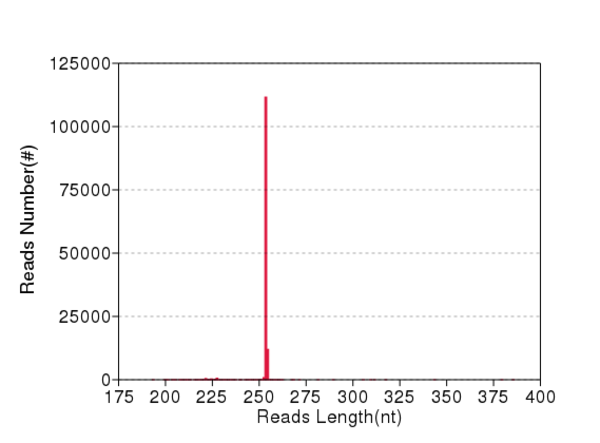

### LabiotFG.QD.JPEG

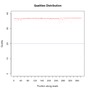

### LabiotFG.QD.png

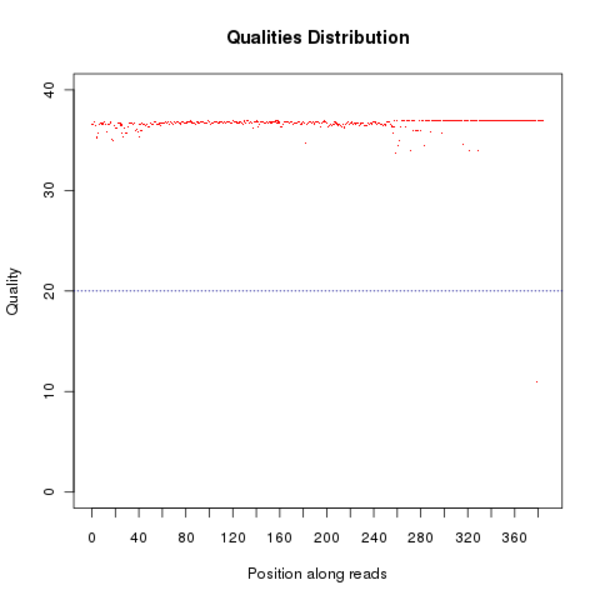

### MicroceroCM.Error.JPEG

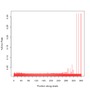

### MicroceroCM.GC.JPEG

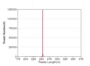

### MicroceroCM.GC.png

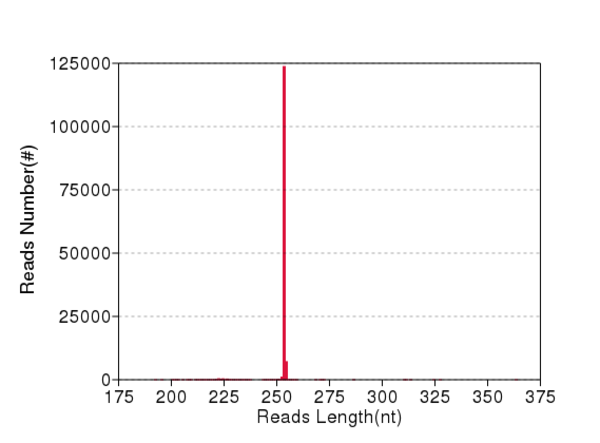

### MicroceroCM.QD.JPEG

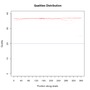

### NasutBR.Error.png

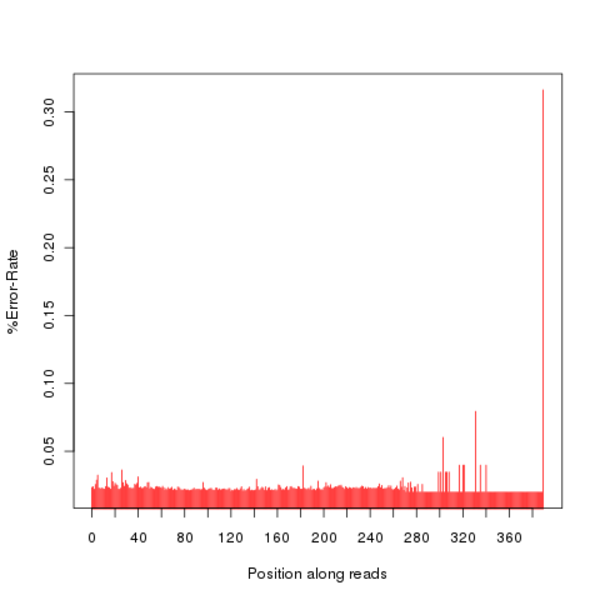

### NasutBR.GC.JPEG

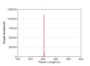

### Nasutluj1CM.Error.JPEG

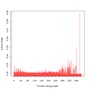

### Nasutluj1CM.Error.png

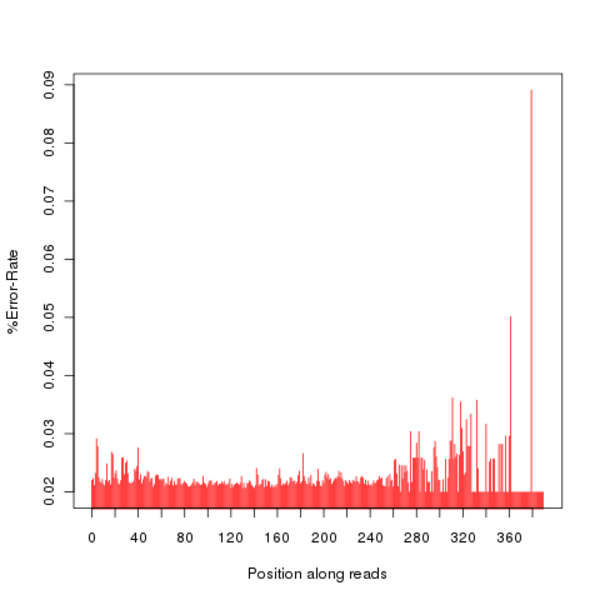

### Nasutluj1CM.GC.JPEG

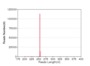

### Nasutluj2CM.Error.JPEG

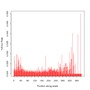

### Nasutluj2CM.GC.png

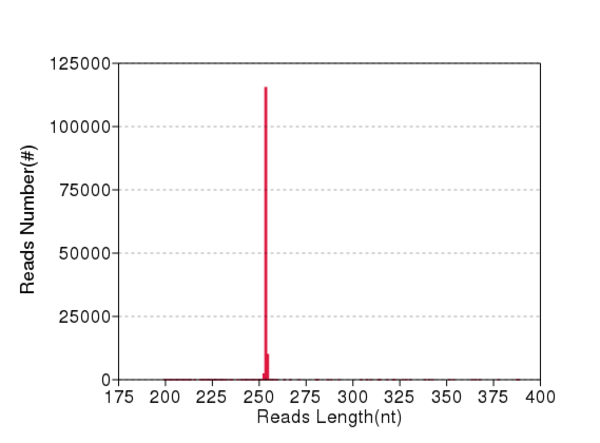

### Nasutluj2CM.QD.JPEG

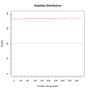

### Nasutluj2CM.QD.png

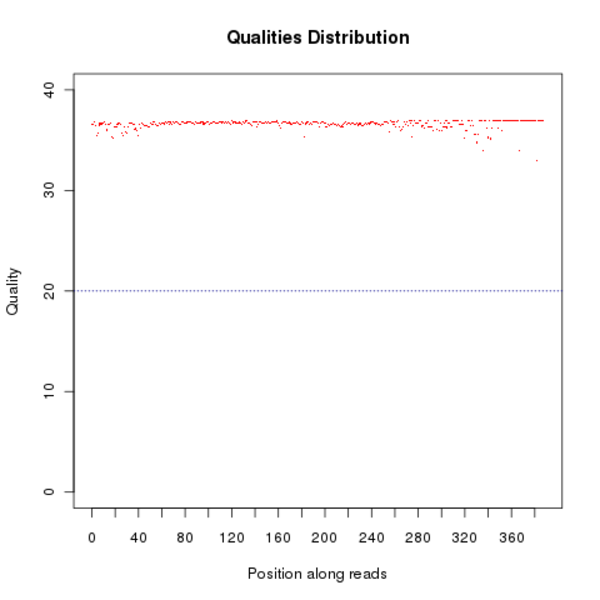

### Nasutluj3CM.GC.JPEG

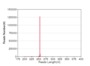

### Nasutluj3CM.GC.png

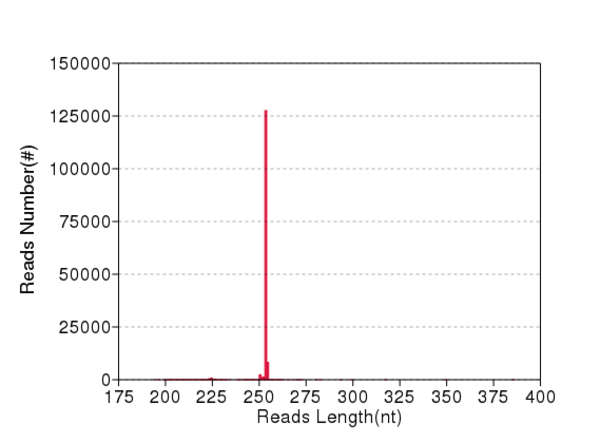

### NasutsimFG.Error.png

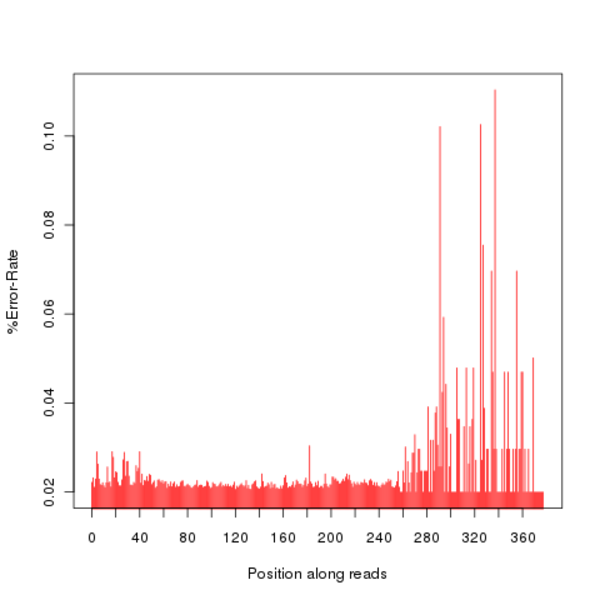

### NasutsimFG.QD.JPEG

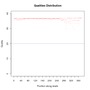

### NasutspCM.Error.JPEG

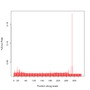

### NasutspCM.GC.png

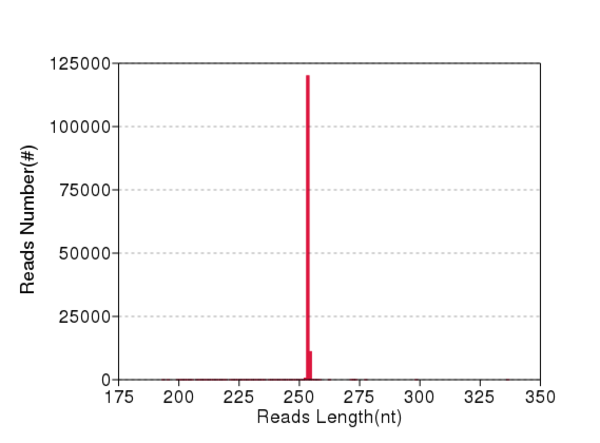

### NasutspFG.Error.JPEG

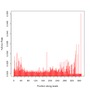

### NasutspFG.QD.JPEG

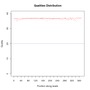

### NasutspFG.QD.png

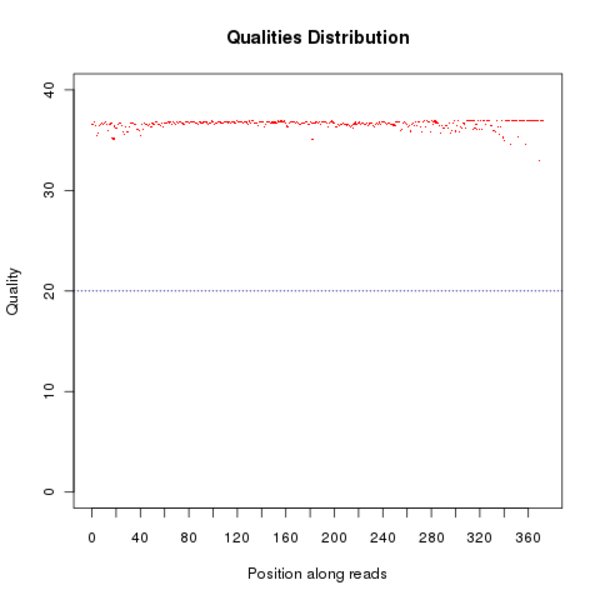

### Negative.Error.JPEG

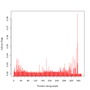

### Negative.Error.png

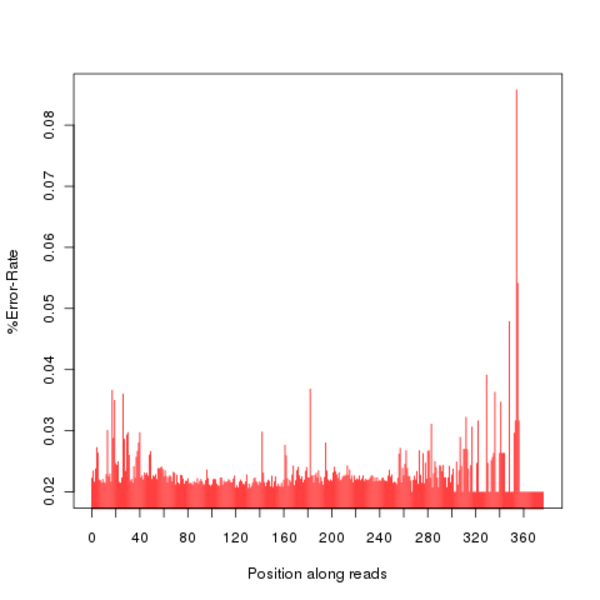

### Negative.QD.JPEG

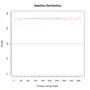

### open.gif

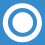

### sequence_pipeline.png

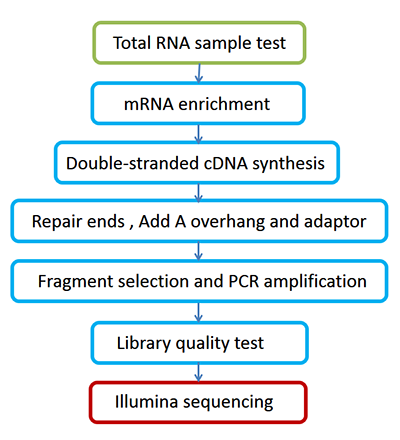

### sequence_pipeline_strand.png

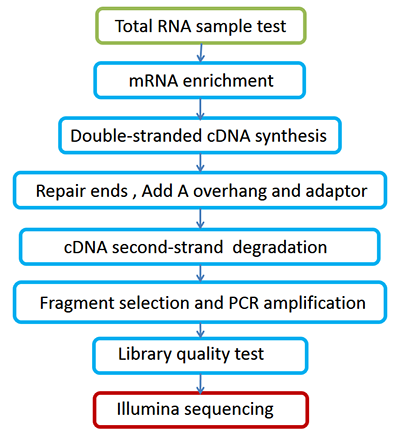

### SilvestFG.Error.png

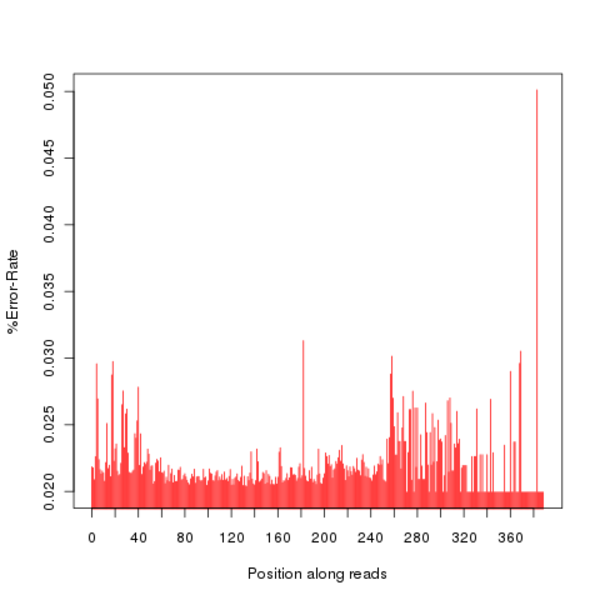

### Taracua0FG19.GC.png

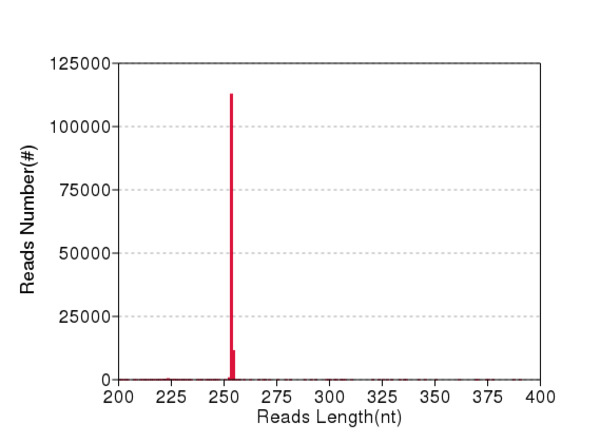

### Taracua0FG19.QD.JPEG

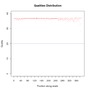

### Taracua4FG19.Error.JPEG

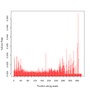

### Taracua4FG19.GC.JPEG

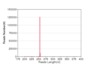

### Taracua4FG19.QD.png

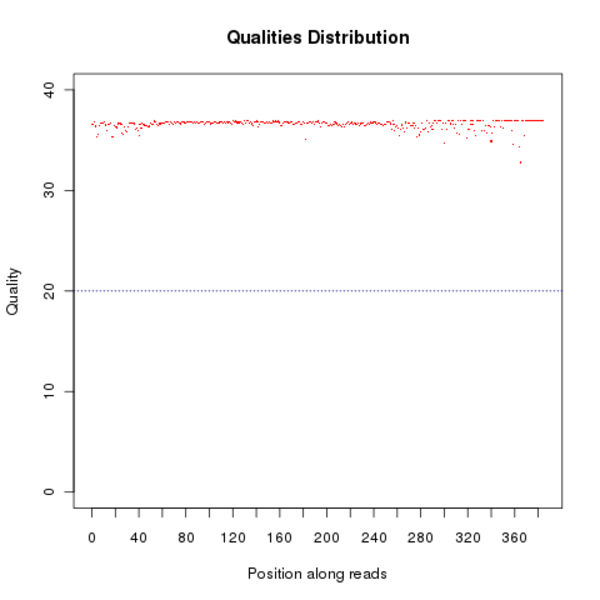

### TaracuaFG20.GC.JPEG

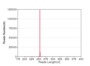

### TermesFG.GC.JPEG

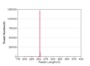

### TermesFG.QD.png

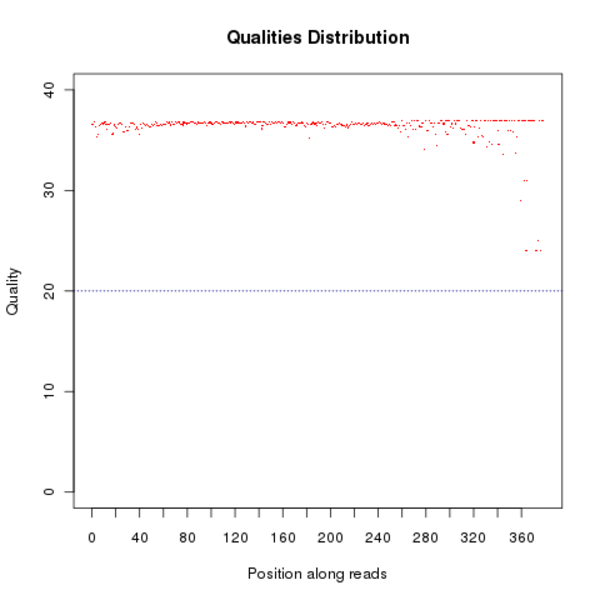
