## Supplementary figures and images for "Termite diet rather than geographical origin determines the microbiome composition and functional genetic structure of nests from South American and African representatives, as revealed by a multiomics approach"

### 16s_lib_en.png

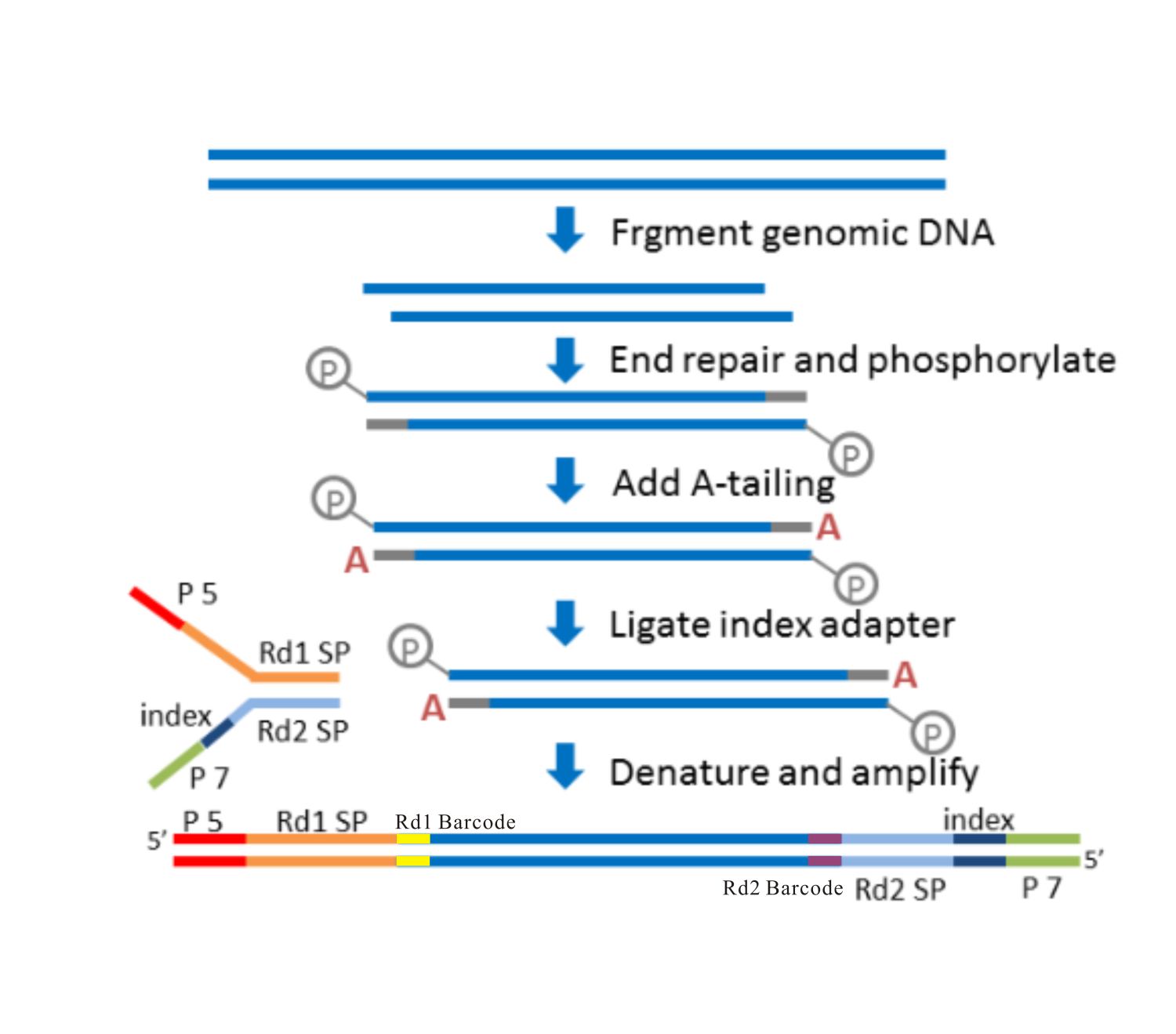

### Anba1FG.GC.JPEG

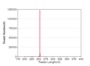

### Anba1FG.QD.JPEG

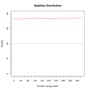

### Anba2FG.Error.png

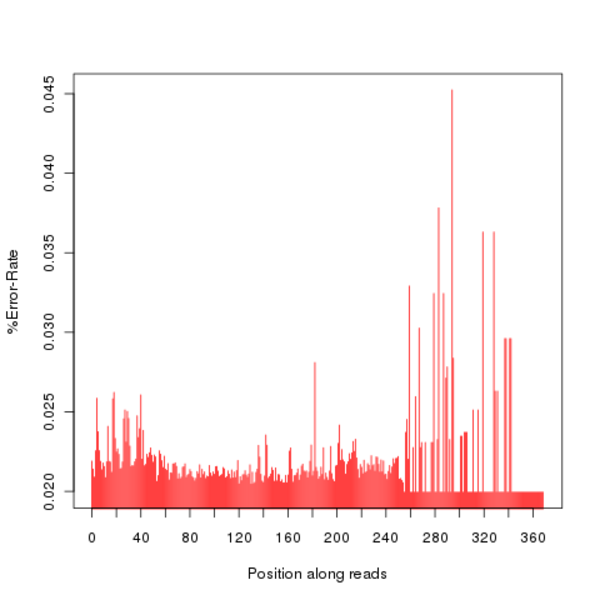

### Anba2FG.GC.JPEG

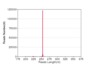

### Anba3FG.Error.JPEG

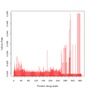

### Anba3FG.Error.png

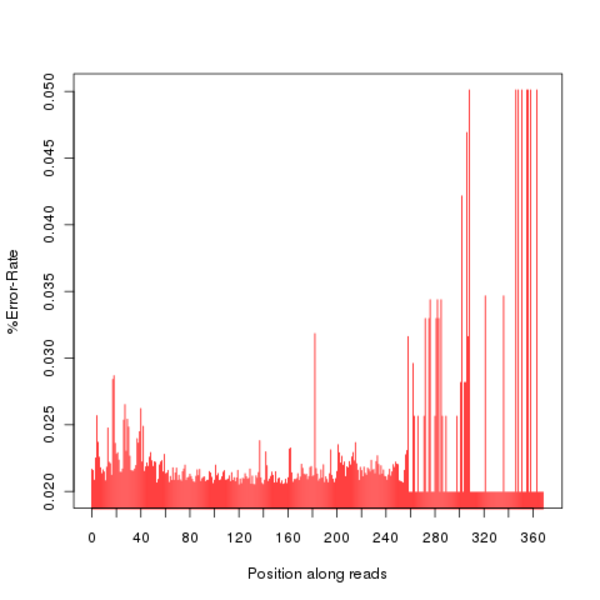

### Anba3FG.GC.JPEG

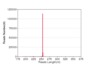

### Anba4FG.GC.png

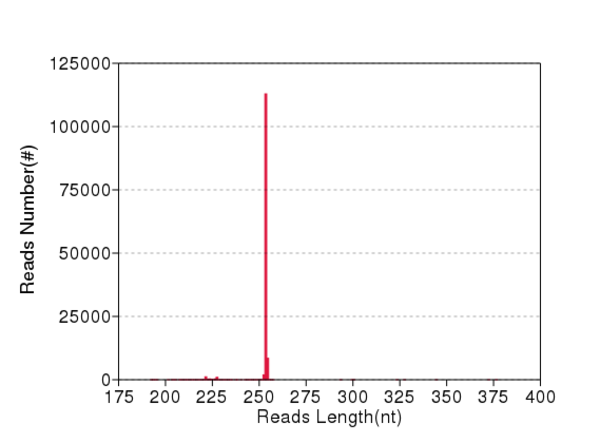

### CephCM.QD.png

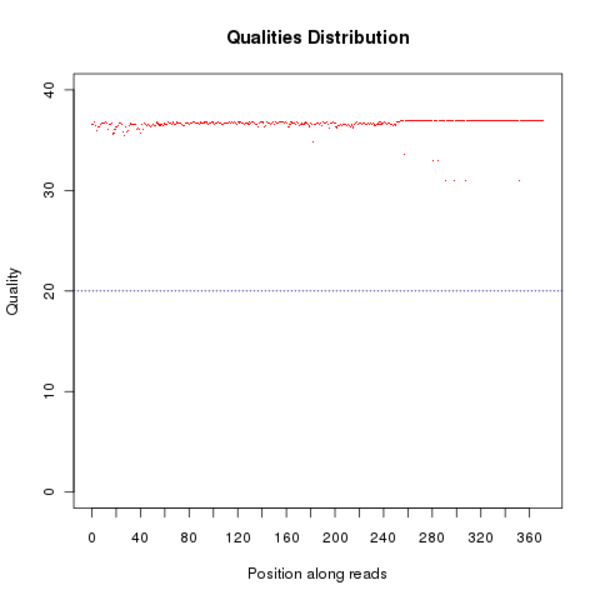

### ConstrCM.Error.JPEG

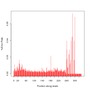

### ConstrCM.Error.png

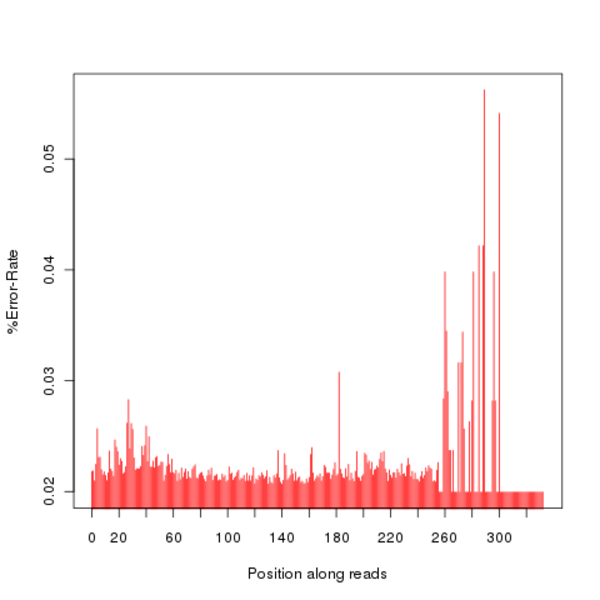

### ConstrCM.GC.JPEG

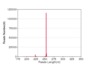

### ConstrCM.QD.JPEG

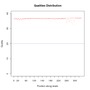

### ConstrCM.QD.png

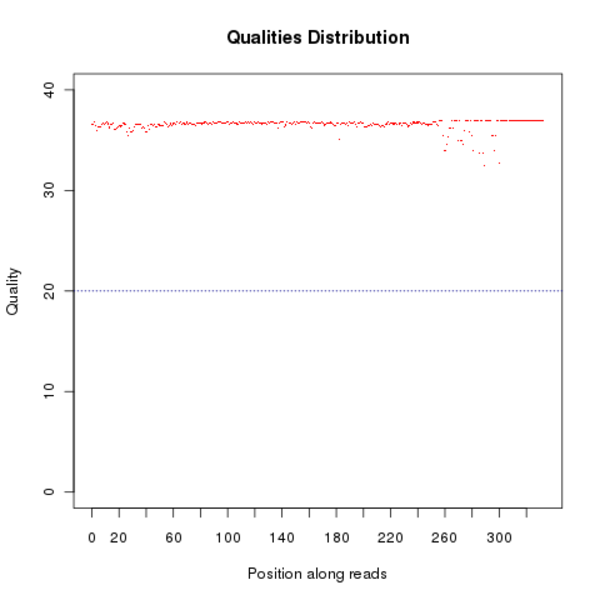

### CoptoBR.Error.JPEG

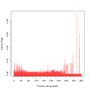

### CoptoBR.GC.JPEG

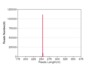

### Cubiter1CM.Error.png

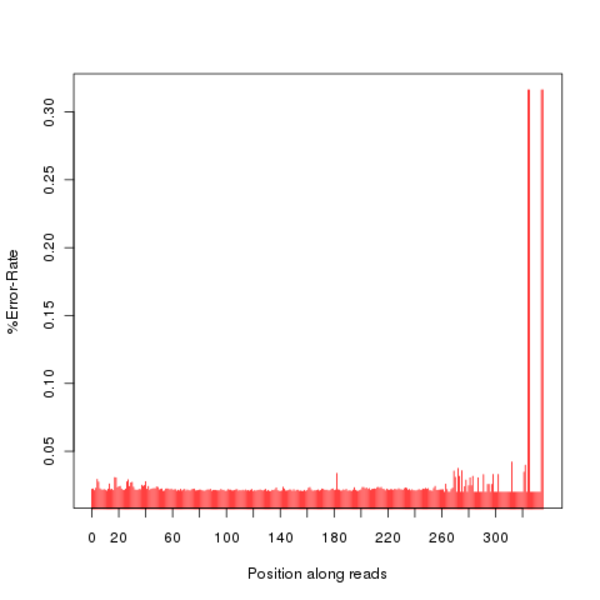

### Cubiter1CM.GC.png

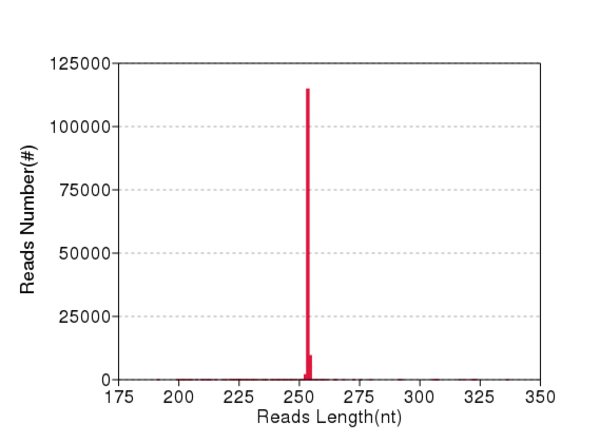

### Cubiter1ML.Error.JPEG

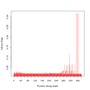

### Cubiter1ML.GC.png

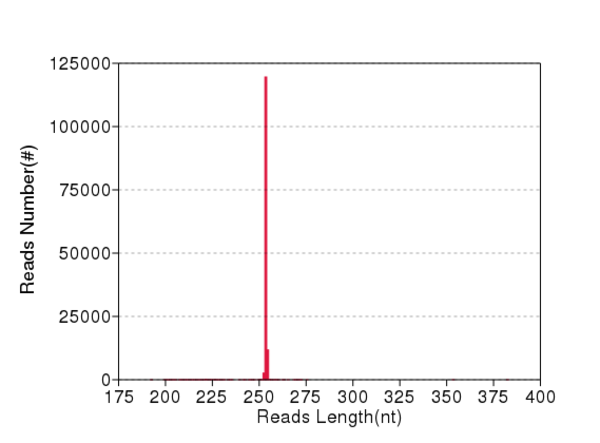

### Cubiter2CM.Error.JPEG

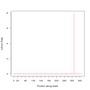

### Cubiter2CM.GC.JPEG

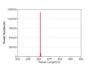

### Cubiter2CM.GC.png

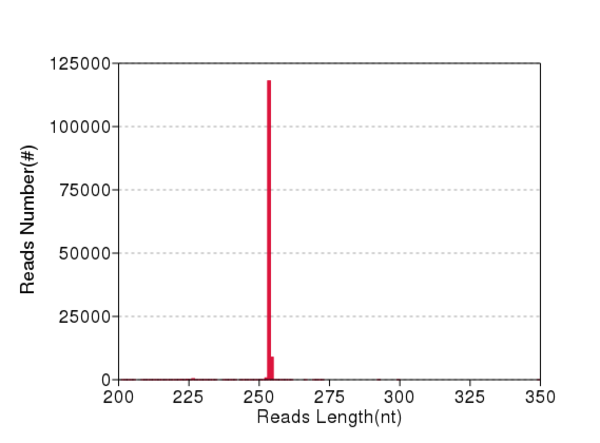

### Cubiter2CM.QD.JPEG

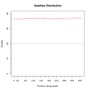

### Cubiter2ML.Error.png

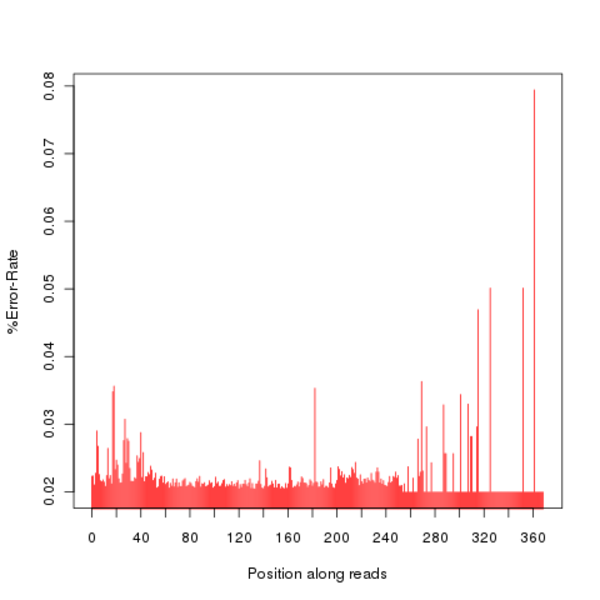

### Cubiter2ML.GC.JPEG

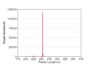

### Cubiter2ML.QD.png

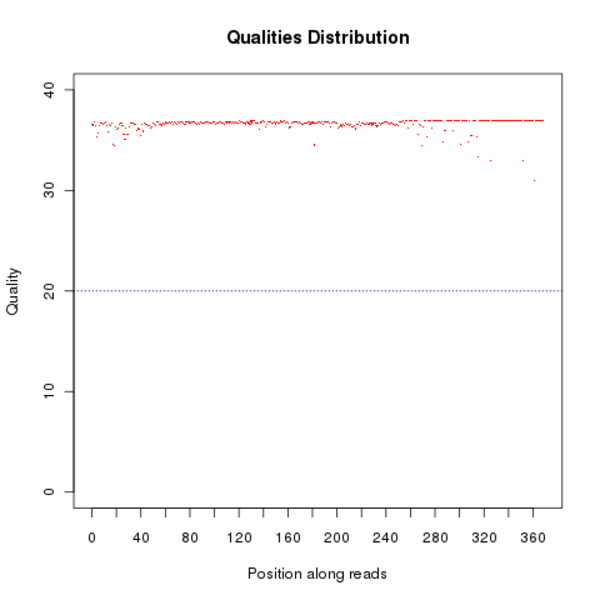

### Cubiter3ML.QD.png

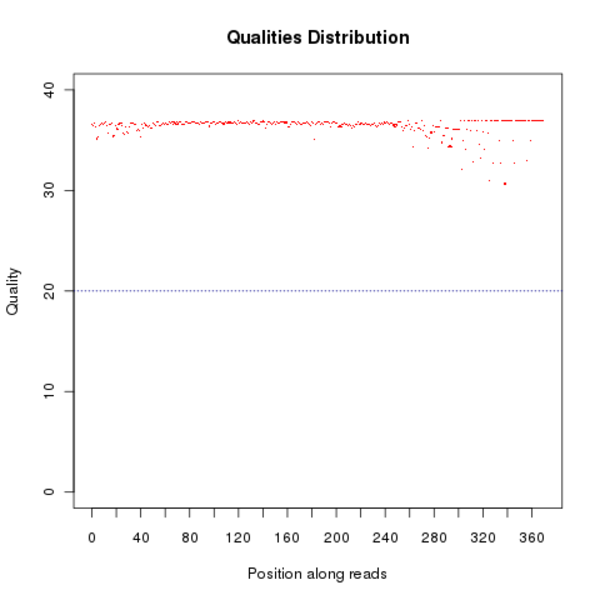

### Cubiter4ML.GC.png

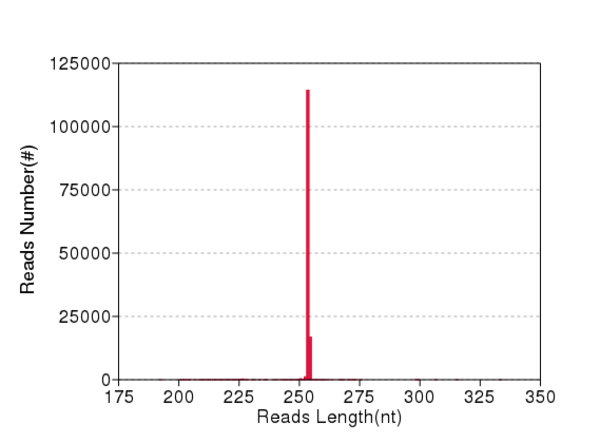

### Cubiter4ML.QD.png

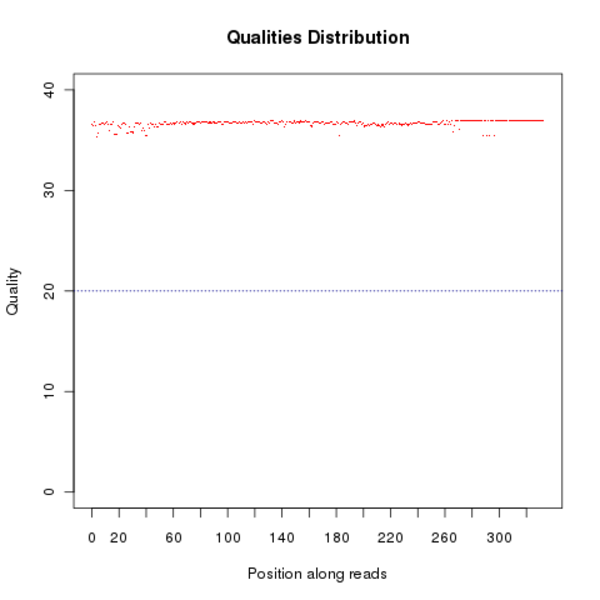

### dna_lib_en.png

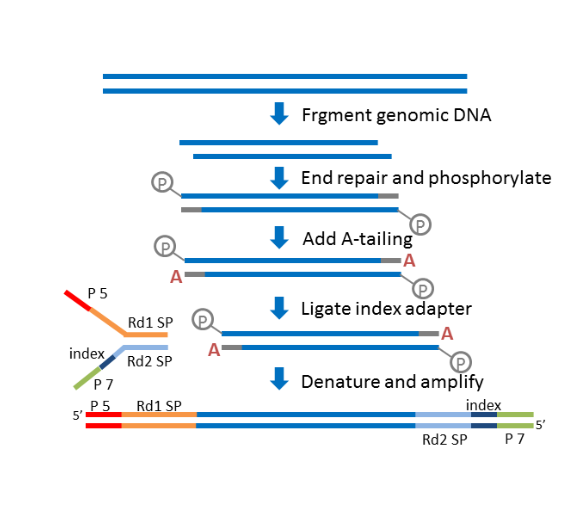

### dna_pipeline_en.png

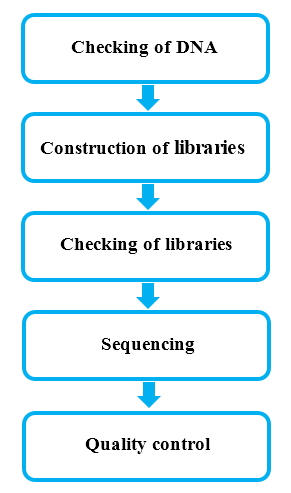

### EmbiraFG.Error.png

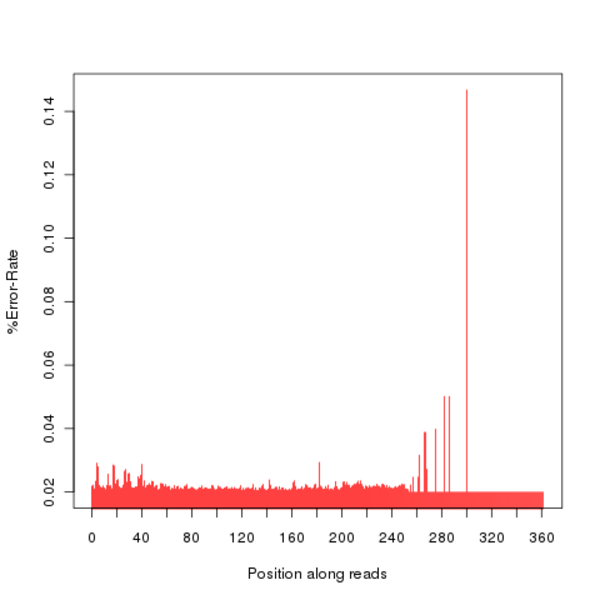

### EmbiraFG.GC.JPEG

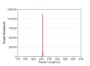

### EmbiraFG.QD.JPEG

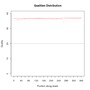

### EmbiraFG.QD.png

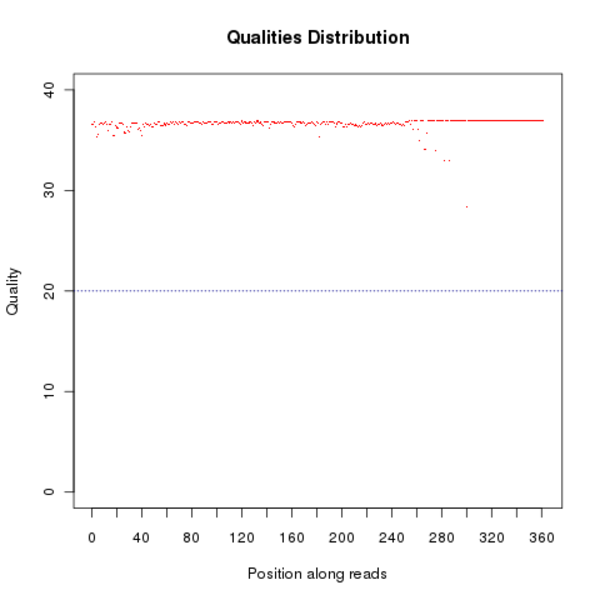

### exp_pipeline_1.png

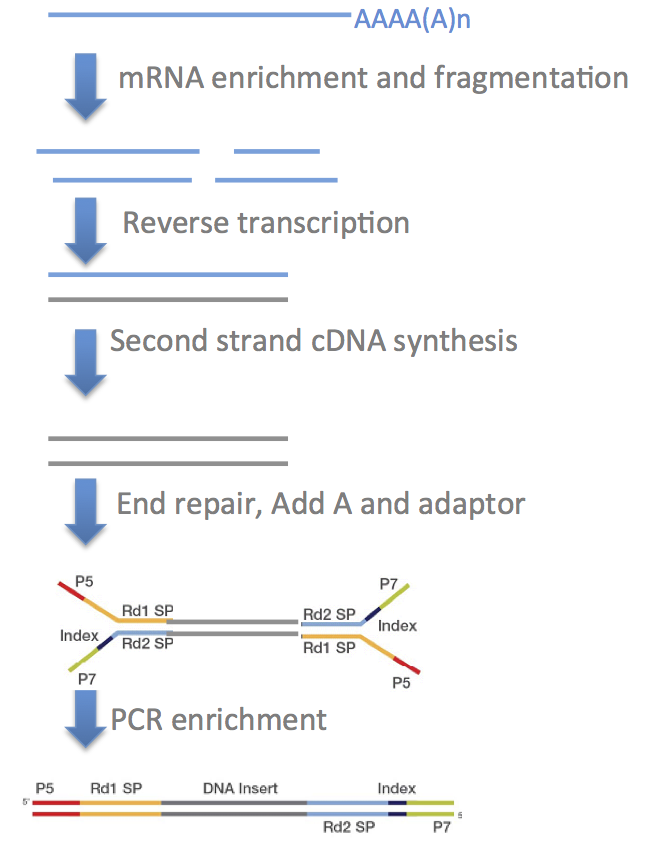

### fancy_close.png

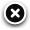

### fancy_loading.png

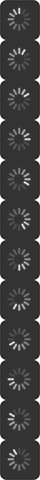

### fancy_nav_left.png

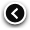

### fancy_nav_right.png

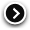

### fancy_shadow_ne.png

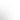

### fancy_shadow_nw.png

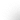

### fancy_shadow_se.png

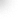

### fancy_shadow_sw.png

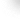

### fancy_title_left.png

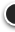

### fancy_title_right.png

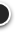

### fancybox.png

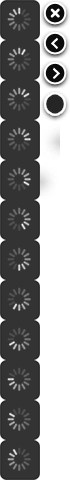

### goTop.jpg

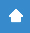

### LabiotFG.Error.JPEG

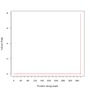

### library_pipeline.png

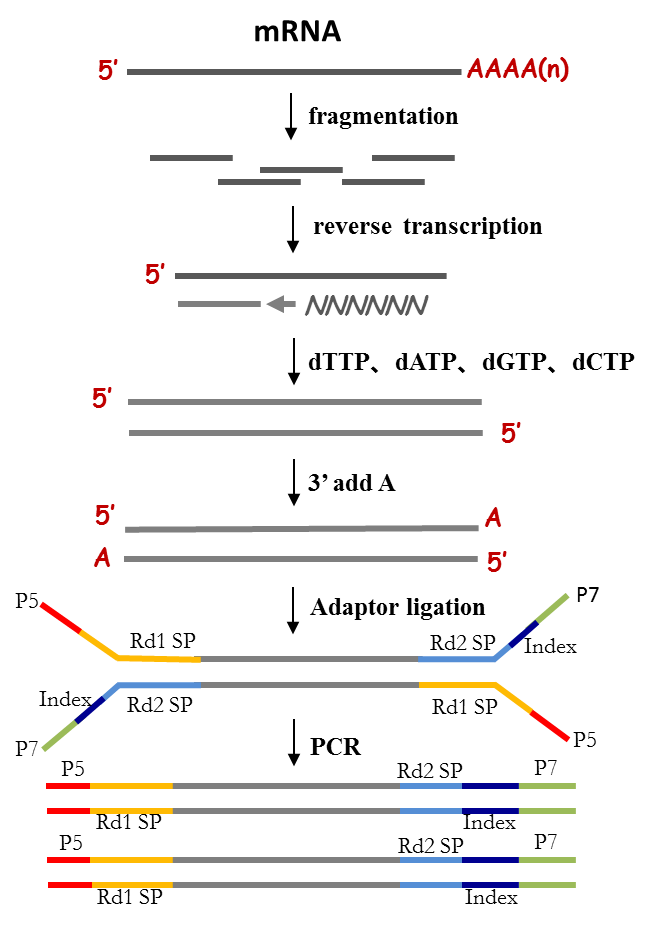

### library_pipeline_strand.png

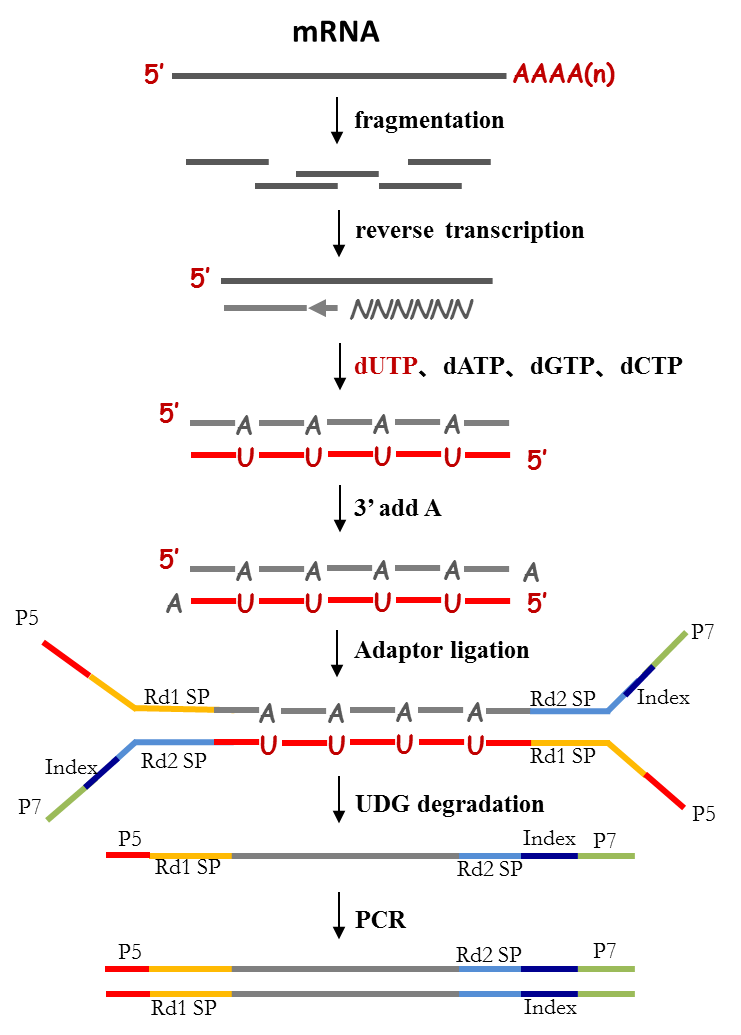

### logo.png

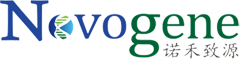

### MicroceroCM.Error.png

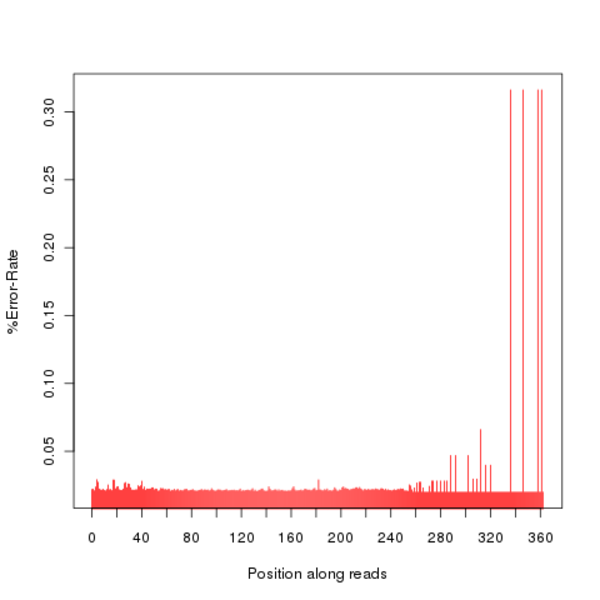

### NasutBR.QD.JPEG

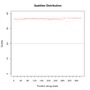

### NasutBR.QD.png

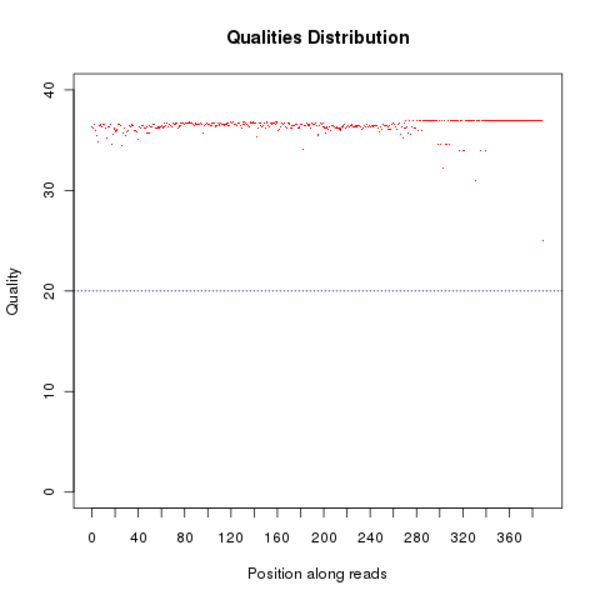

### Nasutluj1CM.GC.png

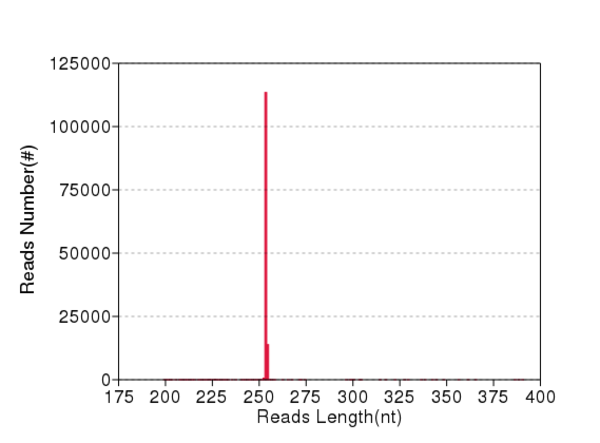

### Nasutluj1CM.QD.JPEG

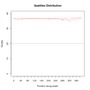

### Nasutluj1CM.QD.png

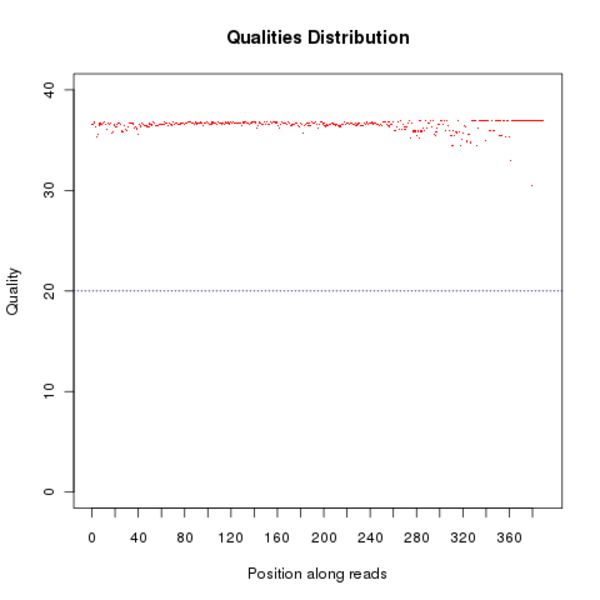

### Nasutluj3CM.Error.JPEG

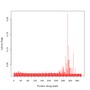

### Nasutluj3CM.Error.png

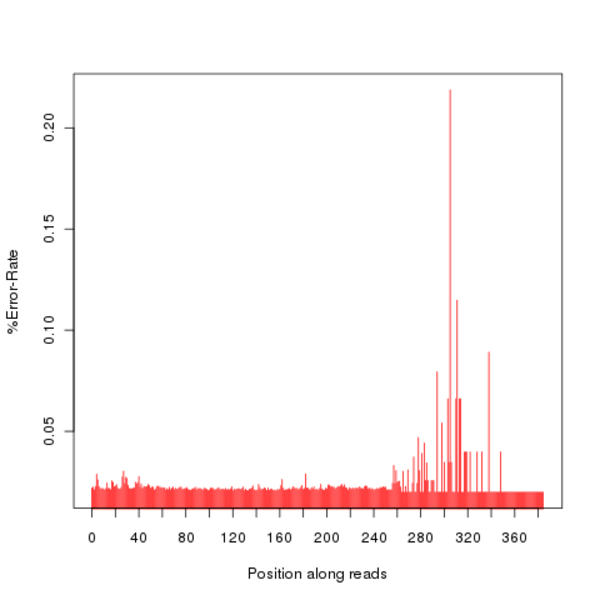

### Nasutluj3CM.QD.JPEG

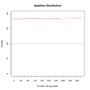

### Nasutluj3CM.QD.png

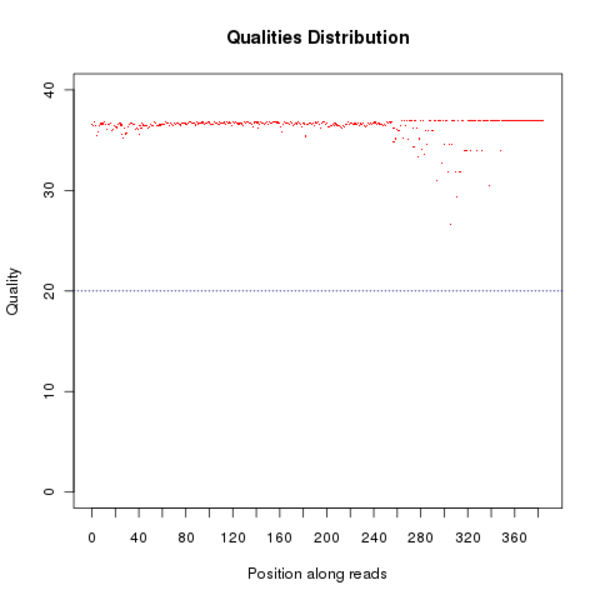

### NasutsimFG.GC.JPEG

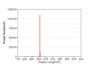

### NasutsimFG.GC.png

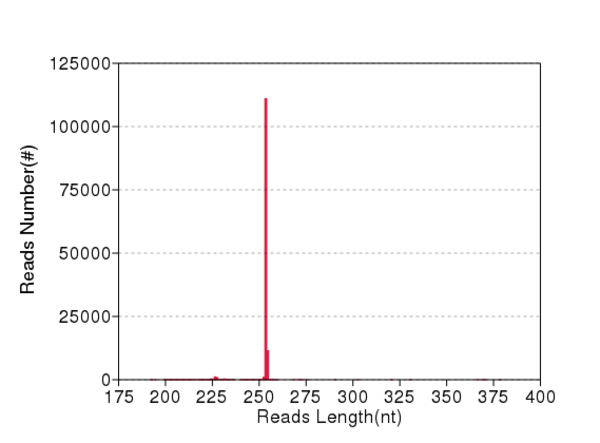

### NasutspCM.Error.png

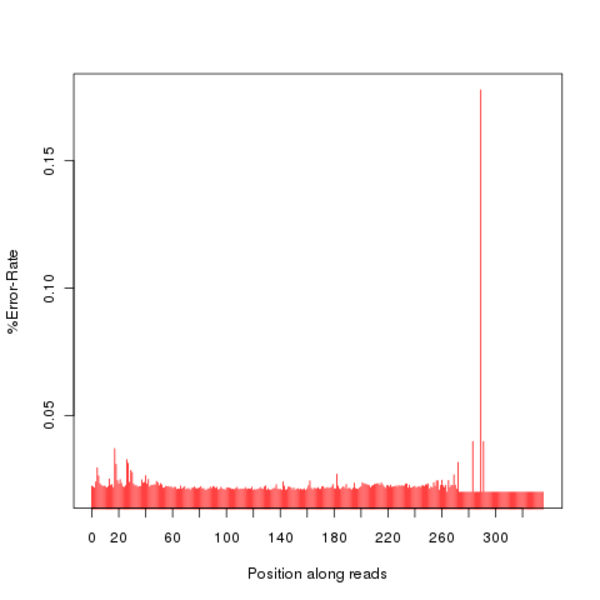

### NasutspCM.GC.JPEG

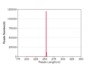

### NasutspCM.QD.JPEG

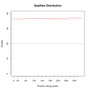

### NasutspCM.QD.png

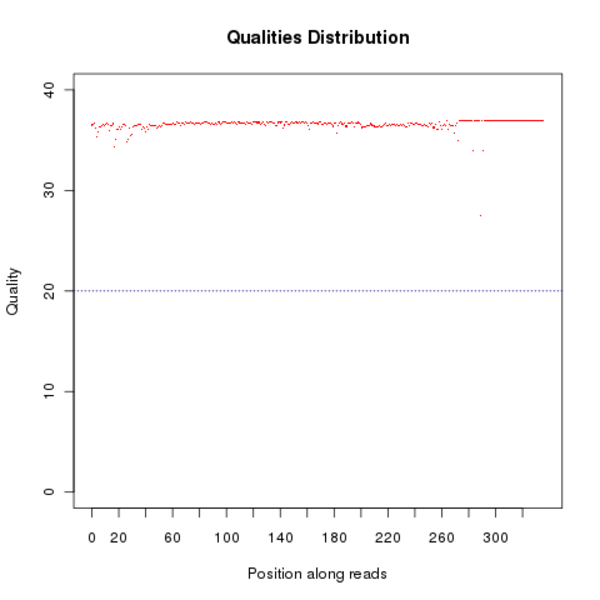

### NasutspFG.GC.JPEG

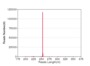

### NasutspFG.GC.png

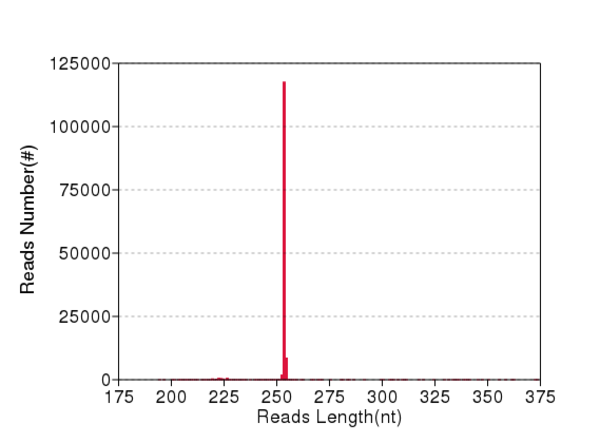

### Negative.GC.JPEG

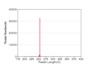

### Negative.QD.png

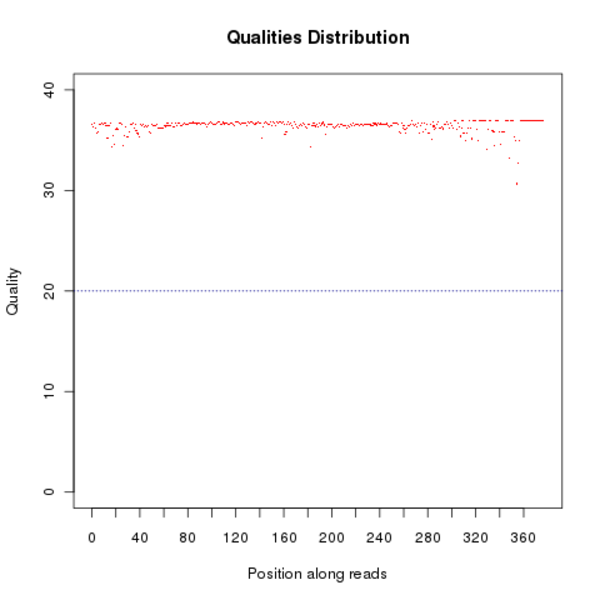

### SilvestFG.Error.JPEG

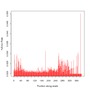

### SilvestFG.GC.JPEG

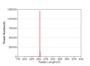

### SilvestFG.GC.png

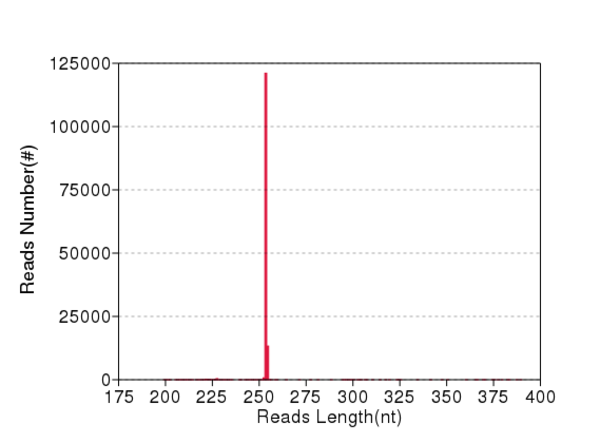

### SilvestFG.QD.png

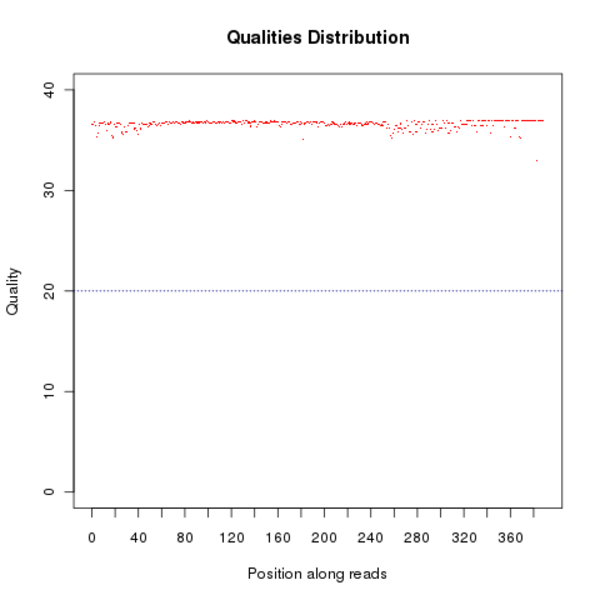

### tags_pipeline_en.png

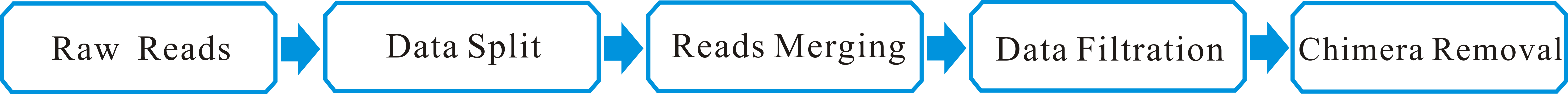

### Taracua0FG19.Error.png

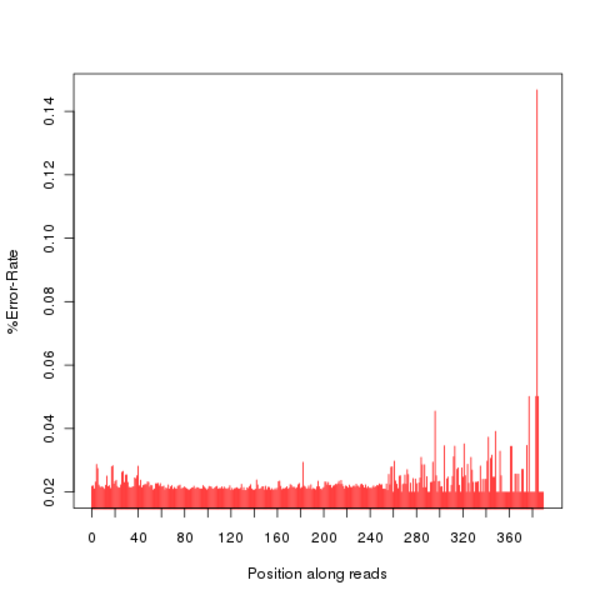

### Taracua0FG19.GC.JPEG

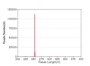

### Taracua0FG19.QD.png

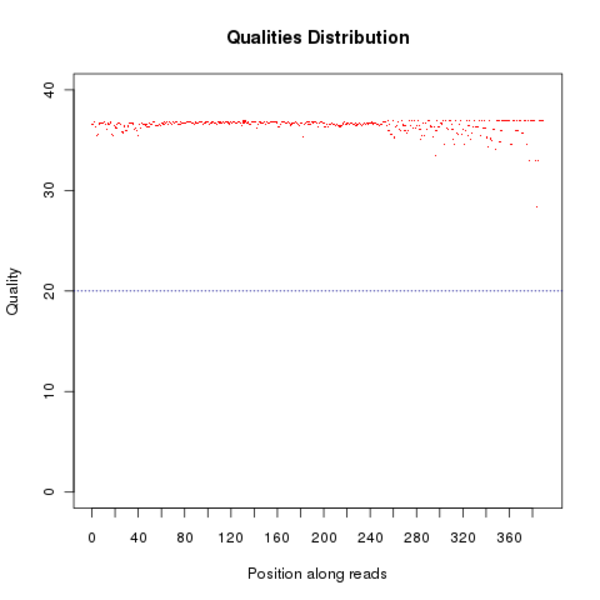

### Taracua4FG19.Error.png

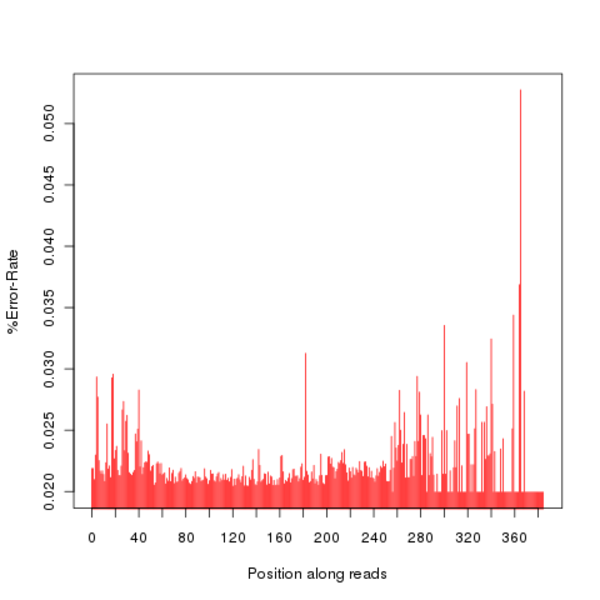

### Taracua4FG19.QD.JPEG

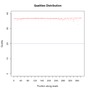

### TaracuaFG20.Error.JPEG

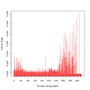

### TaracuaFG20.Error.png

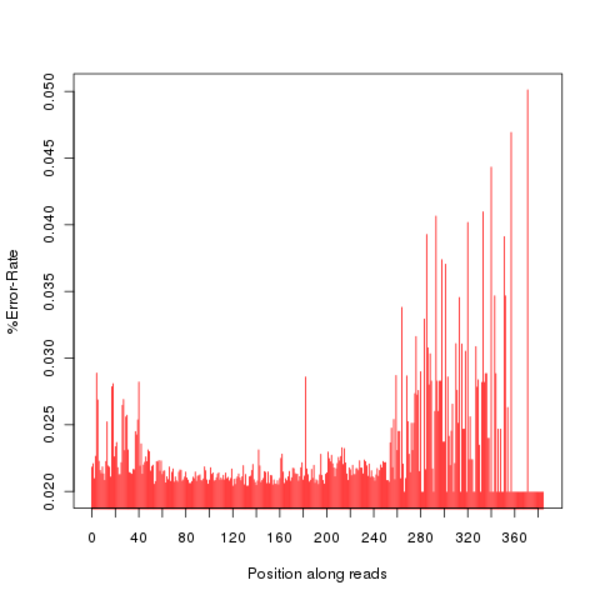

### TaracuaFG20.GC.png

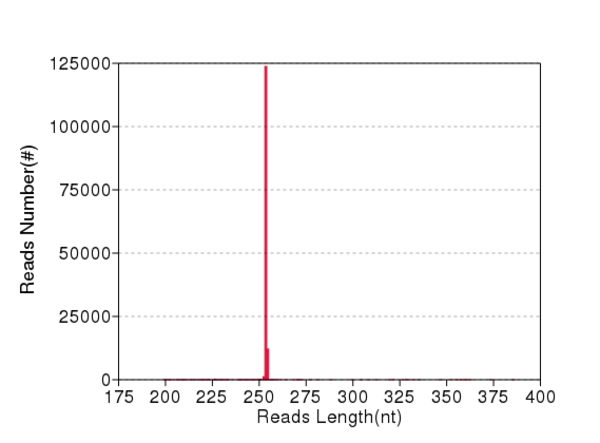

### TaracuaFG20.QD.JPEG

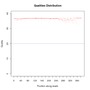

### TermesFG.Error.png

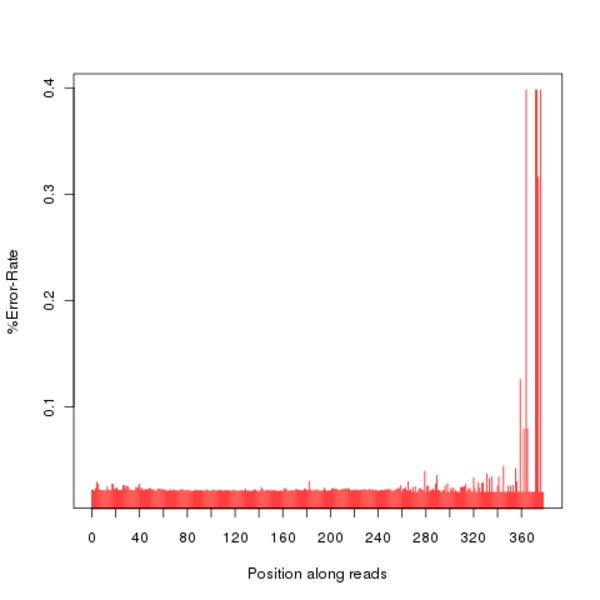

### TermesFG.GC.png

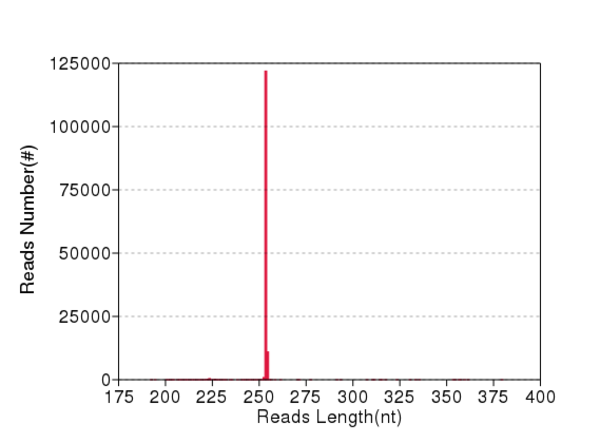

### TermesFG.QD.JPEG

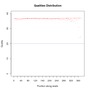
